# Deep estimation of the intensity and timing of selection from ancient genomes

**DOI:** 10.1101/2023.07.27.550703

**Authors:** Guillaume Laval, Etienne Patin, Lluis Quintana-Murci, Gaspard Kerner

## Abstract

Leveraging past allele frequencies has proven to be key to identify the impact of natural selection across time. However, this approach often suffers from imprecise estimations of the intensity (*s*) and timing (*T*) of selection particularly when ancient samples are scarce in specific epochs. Here, we aimed at bypassing the computation of past allele frequencies by implementing new convolutional neural networks (CNNs) algorithms that directly use ancient genotypes sampled across time to refine the estimations of selection parameters. Using computer simulations, we first show that genotype-based CNNs consistently outperform an approximate Bayesian computation (ABC) approach based on past allele frequency trajectories, regardless of the selection model assumed and of the amount of ancient genotypes available. When applying this method to empirical data from modern and ancient Europeans, we confirmed the reported excess of selection events in post-Neolithic Europe, independently of the continental subregion studied. Furthermore, we substantially refined the ABC-based estimations of *s* and *T* for a set of positively-and negatively-selected variants recently identified, including iconic cases of positive selection and experimentally validated disease-risk variants. Thanks to our CNN predictions we provide support to the history of recent and strong selection in northern Europe associated to the Black Death pandemic and confirm the heavy burden recently imposed by tuberculosis in Europe. These findings collectively support that detecting the imprints of natural selection on ancient genomes are crucial for unraveling the past history of severe human diseases.

## INTRODUCTION

The comparison of allele frequencies derived from ancient DNA (aDNA) data at various time periods has enabled the direct identification of the impact of natural selection over time. Studies using such an approach have recently provided new insight into genes and functions involved in human adaptation related to various phenotypes (CHILDEBAYEVA *et al*. 2022; JU and MATHIESON 2021; KEY *et al*. 2016; LINDO *et al*. 2016; MATHIESON 2020; MATHIESON *et al*. 2018; MATHIESON *et al*. 2015; MATHIESON and MATHIESON 2018). These methods also quantify and date the start of the relative advantage (positive selection) or disadvantage (negative selection) conferred by a particular allele in terms of reproductive success, that is, the strength (*s*) and the time (*T*) of onset of selection, respectively. For example, by combining allele frequency trajectories derived from 2,879 ancient and 503 modern genomes and an ancestry-aware approximate Bayesian computation (ABC) framework (BEAUMONT and RANNALA 2004; BEAUMONT *et al*. 2002), a recent study explored the occurrence and selection onset of variants subject to positive or negative time-dependent selection over the last 10,000 years of European history (KERNER *et al*. 2023). Interestingly, the majority of the selection events detected postdated the beginning of the Bronze Age, <4,500 years ago (ya), suggesting that the bulk of human adaptation during the last 10 millennia occurred after the end of the Neolithic period. However, these estimations suffer from having large confidence intervals (CIs), highlighting the need of developing new, more powerful methods.

Methods leveraging past allele frequencies such as ABC or maximum likelihood (KERNER *et al*. 2021; Kerner *et al*. 2023; Key *et al*. 2016; Mathieson 2020; Mathieson *et al*. 2015; Mathieson and MATHIESON 2018), here denoted as frequency-based methods, can provide imprecise estimations of selection parameters when sample sizes used to estimate allele frequencies are small or fail to accurately represent specific epochs. In extreme cases, ABC CIs for the age of selection can nearly cover the entire prior distribution (KERNER *et al*. 2023). Because the use of finer epochs (with lower sample sizes) in ABC is also challenging due to the inclusion of high numbers of summary statistics in the ABC model, an intuitive solution is to directly leverage the ancient individual genotypes, bypassing the computation of allele frequencies in arbitrarily-defined epochs. We thus reasoned that images showing the evolution of individual genotypes across time should contain all information needed to accurately predict past selection parameters and that fine-grained allele frequency trajectories are hidden features of such images, which can be captured by convolutional neural networks (CNNs) (Figure 1). Such deep learning algorithms (CHENG *et al*. 2018; GU *et al*. 2018) have been applied to modern genotype data to infer population genetic parameters (CHAN *et al*. 2018; FLAGEL *et al*. 2019; KERN and SCHRIDER 2018), including the selection coefficient (TORADA *et al*. 2019).

**Figure 1.**
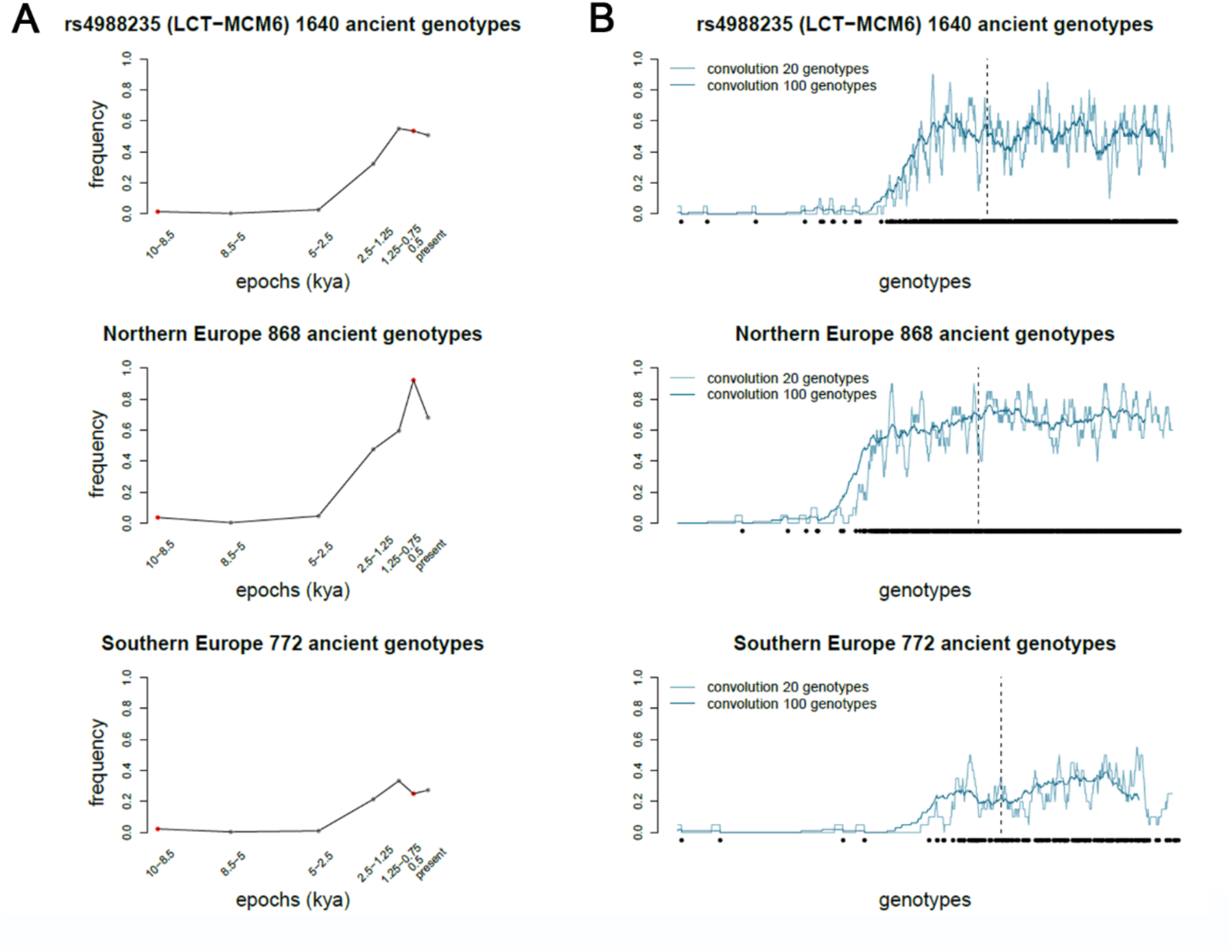
Allele frequency trajectories and genotype-based 1D images for the lactase persistence variant. Allele frequency trajectories (A) and genotype-based 1D images (B) used to perform the estimation of selection parameters by ABC and CNN, respectively. (A) Red points indicate the epochs excluded from the ABC estimation, due to the low sample size of ancient individuals. (B) The input 1D images of the CNNs are shown in the bottom. Black and white points indicate the derived and ancestral alleles, respectively. The black, vertical dashed line separates ancient pseudo-haploid genotype data (left) from modern diploid genotype data (right). Ancient genotypes are sorted by radiocarbon ages of the corresponding individuals, from the oldest on the left to the youngest on the right. The blue curves represent the convolution performed by the first 1D convolution layer in the CNNs. The convolution revealed a fine-grained derived allele frequency trajectory hidden in the image, since a convolution correspond to a sliding average of pixel values (0 for white or 1 for black) using sliding windows of 20 or 100 genotypes. Note the equivalence with allele frequencies (A), as expected since the average of black pixels in a window corresponds to the frequency of the derived allele (see the limit value being the frequency in the 1000G Europeans). (A-B) Top, middle and bottom panels show allele frequency trajectories and 1D images obtained for the lactase persistence variant using all Europeans, northern Europeans, or southern Europeans, respectively.

Here, we present new CNN architectures that directly use individual genotypes, denoted as genotype-based CNN algorithms, seeking to jointly estimate the strength (*s*) and the age (*T*) of selection, under additive models of both positive (MAYNARD SMITH and HAIGH 1974; PRITCHARD *et al*. 2010; STEPHAN *et al*. 1992) and negative selection (CHARLESWORTH *et al*. 1993). These algorithms leverage one-dimension (1D) input images of genotypes of individuals sampled at various generations in the past and sorted by their radiocarbon ages. Using computer simulations, we first show that genotype-based CNNs outperform ABC based on allele frequency trajectories. Using empirical data from modern and ancient Europeans, we found highly consistent, yet substantially improved estimations of *s* and *T* for a set of recently detected candidate positively-and negatively-selected variants (KERNER *et al*. 2023). Illustrating improved accuracy compared to ABC, the CIs of *s* and *T* estimates are reduced by ∼20-40% in average across all selected variants when training our CNN on northern and southern Europeans separately. Our analysis replicates the previously reported significantly higher frequency of post-Neolithic adaptation events relative to the pre-Bronze Age period in Europe (KERNER *et al*. 2023), independently of the nature of selection (positive or negative) and the continental subregion considered. Finally, as a case study, our analysis supports a history of recent and strong selection in northern Europe at the *ERAP2* rs2549794 variant, which has been recently associated to the Black Death pandemic (KLUNK *et al*. 2022).

## MATERIALS AND METHODS

### Ancient and modern data

We used 2,632 aDNA genomes (i) originating from burial sites in western Eurasia (-9< longitude (°) <42.5 and 36< latitude (°) <70.1), (ii) genotyped for 1,233,013 polymorphic sites by ‘1240k capture’, (MATHIESON *et al*. 2015) (iii) showing genotype calls for at least 50,000 polymorphisms, and (iv) retrieved from the V44.3: January 2021 release at https://reich.hms.harvard.edu/allen-ancient-dna-resource-aadr-downloadable-genotypes-present-day-and-ancient-dna-data. 95 samples that were either annotated as duplicated or found to correspond to first-degree relatives of at least one other individual in the dataset were removed (KERNER *et al*. 2023). We used all remaining ancient individuals, regardless of whether they grouped into one culturally defined epoch, including thus 163 ancient genomes previously discarded in (KERNER *et al*. 2023) (90 with estimated ages >8,500 ya, and 73 between 750 and 250 ya). All ancient individuals were treated as pseudo-haploids, i.e., hemizygotes for either the reference or the alternative allele (the same treatment was applied to the simulated ancient individual used for training the CNNs, see below). We used the 503 Europeans from the 1000 Genomes (1000G) project data (AUTON *et al*. 2015), from the IBS, TSI, FIN, GBR and CEU populations, as modern data. Similarly to modern individuals in the empirical data, simulated modern individuals used for training the CNNs were diploid (see below).

### Variants analyzed

We analyzed the 89 variants with the strongest signals of positive selection (FAN *et al*. 2016; JEONG and DI RIENZO 2014; VITTI *et al*. 2013) at each of the 89 candidate genomic loci found to be under positive selection in a previously conducted genome-wide selection scan, see Table S2 in (KERNER *et al*. 2023), and the variant rs2549794 (*ERAP2*) previously identified as a strong candidate for positive selection due to the Black Death epidemic in northern Europeans (KLUNK *et al*. 2022). Of note, the 3 other variants evidenced by (KLUNK *et al*. 2022) were not tested, since they were absent from the ‘1240k capture’ dataset used in the present study. We also included in this analysis 50 missense variants at conserved positions (GERP score >4) previously found to be under negative selection (KERNER *et al*. 2023). Only variants for which the ancestral allele was annotated in the 1000G data and for which the derived allele was present in at least five aDNA samples were analyzed. These variants were previously controlled for potential artifacts due to undetected technical problems in the ‘1240k capture’ dataset, by comparing the allele frequencies obtained with those from shotgun sequencing data as described in (KERNER *et al*. 2023).

### 1D images and Convolutional Neural Network implemented

We implemented various CNN algorithms with keras (Tensorflow as backdoor) able to jointly estimate *s* and *T* only using the genotypes of modern and ancient individuals sampled at various generation in the past. The genotype *g_i_*_,*j*_ of the variant *i* and individual *j* is defined as a vector of indictor variables (0 and 1 for ancestral and derived alleles) of dimension (1 × *d_i_*_,*j*_), *d_i_*_,*j*_ being the ploidy equal to 1 or 2 for haploid or diploid genotypes respectively. The CNN algorithms leverage individual genotype data thanks to the use of 1D grey scale images built from the list of genotypes *g_i_*_,*j*_ observed among modern and ancient genomes used. A 1D images is encoded from *g_i_*_,*j*_ ordered by the age of ancient individuals, a pixel by allele with black or white color for the derived or ancestral allele respectively. We used the radiocarbon age as proxy of the age of each ancient individual as classically done. Ancient genotypes *g_i_*_,*j*_ were treated as pseudo-haploid data (*d_i_*_,*j*_ = 1) and encoded by one black or white pixel for the derived or ancestral alleles, respectively. Modern 1000G genotypes *g_i_*_,*j*_ were treated as diploid data (*d_i_*_,*j*_ = 2) and encoded by two black or white pixels, the order of pixels being randomized for heterozygotes.

In contrast to fully connected neural networks, the CNNs architecture consists of consecutive sets of hidden neuron layers of different types, the convolutional and pooling layers, followed by a set of fully connected layers that carry out the image classification (KORFMANN *et al*. 2023). The CNN algorithms are canonical feed forward neural networks with a first 1D convolution layer, which applies a convolution to the input image using a kernel (also known as filter; a matrix of learnable parameters (KORFMANN *et al*. 2023)) to produce a feature map, which is then fed to a 1D maxpooling layer. This layer is used to reduce the dimensions of the feature map and capture coarse grained features (features dissociated from their positions in the input image) (KORFMANN *et al*. 2023). We added three additional 1D convolutions and a last 1D maxpooling layer to capture more fine-grained features of the input image. A flatten layer is used to convert the feature map into a vector that is fed to several fully connected (dense) layers. The CNN architecture used in this study is thus inspired by the popular AlexNet model (KRIZHEVSKY *et al*. 2017). During training the parameters of the CNNs are adjusted with backpropagation of the gradient of the cross-entropy loss of function. Two output layers of 100 neurons each are used to obtain joint estimations of *s* and *T* through the softmax function, providing the likelihood of each parameter value/bin (*s* and *T* were discretized in 100 bins each; arithmetic precision at the second digit for floats). The maximum likelihood values were obtained with the argmax function and posterior distributions were obtained by resampling according to the prior and likelihood distributions (TORADA *et al*. 2019). Average of the posteriors were used as point estimates and the 95%CIs were computed from the posterior distributions. We implemented various CNN architectures. These architectures, i.e., the numbers of layers, the numbers of neurons, the hyper parameters (numbers of kernels and their dimensions) and the activation functions, of the various CNNs tested in this study are given in the Supplementary Figure S1.

### Training the CNN algorithms

In order to deal with the variability in genotype calls per SNP (the number of called genotypes can strongly vary across variants due to the low coverage of most ancient samples), the CNN algorithms (Supplementary Figure S1) were trained separately for each genetic variant as follows. For each selected variant *i*, the CNNs are trained using genotypes obtained by computer simulations: the genotypes of modern and ancient individuals sampled at various generations in the past from a European population were simulated under a specified demographic model, with values of the selection parameters *s* and *T*, drawn from uniform distributions (KERNER *et al*. 2023). This simulated training dataset accounts for the empirical proportions of ancestry in each generation and reproduces the nature of the empirical data used: diploid and pseudo-haploid genotypes for modern and ancient individuals respectively (the details of the simulations are presented in the next section below). To adjust the training dataset to the empirical data used for prediction, the number and age of simulated genotypes are matched to that observed by excluding the simulated genotypes corresponding to empirical ancient individuals with no genotype call for the variant under investigation. Each computer simulation thus outputs a vector ***g****_i_* of genotypes ordered by the age of individuals (sampling time), and of dimension (1 × *D_i_*), with *D_i_* = ∑*_i_ d_i_*_,*j*_ the sum of the ploidy across non-missing genotypes *g_i_*_,*j*_ observed among the 503 1000G modern and 2,537 ancient genomes used. These vectors of genotypes ***g****_i_*, being lists of 0s (ancestral allele) or 1s (derived allele), are each encoded into a 1D dimension image (white or black pixel for 0 or 1 respectively, see above). Each 1D image is then labelled with the *s* and *T* values used to perform the computer simulation, and the set of simulated labelled images thus obtained is used to train the CNN in a supervised learning.

For each selected variant under investigation, we thus generated 500,200 simulated 1D images matching the number and age of genotypes present in the empirical 1D image used to predict the intensity and age of selection. The simulated and empirical 1D image are of the same dimension and thus contain the same number of pseudo-haploid ancient genotypes (1 pixel per genotype) and 503 haploid modern genotypes (2 pixels per genotype). The model is trained using 500,000 simulated 1D images during a rather short number of epochs (3 to 5 depending on the CNN topology). Note that the accuracy of the CNN predictions was also evaluated using a classical cross validation procedure performed based on the 200 remaining simulated data not used during the training process (these simulated data used as empirical data for which the true values of *s* and *T* are known, are referred to as pseudo-empirical datasets in the following sections).

### Forward-in-time computer simulations used to produce the training datasets

We used the computer simulations of a mutation evolving under a demographic model previously described by Kerner and colleagues (2023) and performed with SLiM (HALLER and MESSER 2017; HALLER and MESSER 2019). The demographic model we used includes divergence times, migration rates and exponential growth of effective sizes of continental populations (African, West and East Eurasian populations) (BERGSTROM *et al*. 2020; GRAVEL *et al*. 2011), see the Table S10 in (KERNER *et al*. 2023). Because SLiM is a forward-in-time simulator, the computation times depend on both the effective population size *N* and the number of generations *t* considered. Effective population sizes and times were thus rescaled according to *N_e_*/*λ* and *t*/*λ*, with *λ* = 10 (HALLER and MESSER 2017; HALLER and MESSER 2019; HOGGART *et al*. 2007). Importantly, the model accounts for migratory events inferred with aDNA data (SKOGLUND and MATHIESON 2018), including the arrival of Anatolian early farmers in Europe ∼8,500 ya (LAZARIDIS *et al*. 2014; SKOGLUND *et al*. 2012), who admixed with the local Mesolithic hunter-gatherers, and that of populations of Yamnaya culture from the north of the Caucasus ∼4,500 ya (ALLENTOFT *et al*. 2015; HAAK *et al*. 2015). For each simulation, the age of the mutation *T*_age_, the age of selection *T* and the strength of selection, as measured by the selection coefficient *s*, were randomly sampled from uniform prior distributions: *T*_age_ ∼ *U*(1000, 10^6^) ya, *T* ∼ *U*(1000, 10000) ya and s ∼ *U*(-0.05, 0) for negative selection and *s* ∼ *U* (0, 0.1) for positive selection, under an additive model (*h* = 0.5) (HALLER and MESSER 2017; HALLER and MESSER 2019). The age of the mutation was defined as the point at which the mutation was introduced into the model in a randomly chosen population. Simulated individuals were randomly sampled from the simulated population at the generations corresponding to the radiocarbon ages of the observed individuals, so that all sampled ancient individuals, as well as the 503 modern individuals of the 1000G dataset, are simulated. For each simulated ancient individual and each polymorphic site, we randomly sampled one allele to generate pseudo-haploid data, mirroring the observed pseudo-haploid empirical aDNA data used. The simulated genotypes corresponding to empirical ancient individuals with no genotype call for the variant under investigation were excluded and the simulated ancestry proportions at each generation was matched to the mean ancestry proportions of the ancient and modern individuals previously estimated at the individual level using a factor analysis (FRANÇOIS and JAY 2020) and 143,081 high-quality SNPs, as described in (KERNER *et al*. 2023). Simulated present-day European individuals were randomly drawn from the last generation of the simulated population and their genotypes were used as diploid data. Of note, variants were simulated one at a time, so that a single simulated allele frequency trajectory was obtained from each simulation.

### ABC estimation from allele frequency trajectories across time transects

We estimated *s* and *T* with a previously described (KERNER *et al*. 2021; KERNER *et al*. 2023) ABC approach (BEAUMONT *et al*. 2002). Such a frequency-based method leverages a vector ***p****_i_* of dimension (1 × *K*), namely the derived allele frequency trajectory computed across *K* discrete epochs arbitrary defined based on the radiocarbon ages of ancient individuals. In each epoch, the derived allele frequency is computed from *p_i_* = ∑*_j_ G_i_*_,*j*_ / ∑*_j_ d_i_*_,*j*_, with *G_i_*_,*j*_ being equal to 0, 1 or 2 for genotypes carrying 0, 1 or 2 derived alleles depending on the ploidy *d_i_*_,*j*_ (the definition of the ploidy *d_i_*_,*j*_ is indicated above). We used the same 503 modern 1000G Europeans and 2,537 ancient genomes used with CNNs to compute the derived allele frequency trajectory ***p****_i_*. The ancient genomes were grouped in *K* = 5 epochs; the Neolithic (8,500-5,000 ya; *n* =729), the Bronze Age (5,000-2,500 ya; *n* = 893), the Iron Age (2,500-1,250 ya; *n* = 319), the Middle Ages (1,250-750 ya; *n* = 435) and present day times (0 ya; *n* = 503). For each simulated or empirical variant *i*, the genetic information in ancient and modern genomes was thus summarized by a five-dimensional (1 × *K*=5) frequency vector, ***p****_i_* used as the summary statistics to fit the empirical data with the simulated data. Weights depending on sample sizes per epoch are given to each frequency estimate as previously described (KERNER *et al*. 2021). ABC estimations were performed with the simulated data used for training the CNN algorithms with underlying parameters drawn from the uniform prior distributions presented above. ABC posterior distributions, point estimates (i.e., posterior mode) and the 95% CIs were computed from the parameters underlying the 1,000 computer simulations best fitting the empirical data, according to the standard ABC approach (‘abc’ R package, method = “Loclinear”) (BEAUMONT *et al*. 2002; CSILLERY *et al*. 2012). Of note, we adopted for ABC the terminology classically used in machine learning. For convenience, the ABC model fitting (determination of the lowest ABC distances between simulated and empirical data) will be abusively denoted training even if this step is not a formal training as those performed in machine learning approaches.

### Simulation-based evaluation of the ABC and CNN algorithms

The performance of our CNN algorithms and the ABC approach were evaluated on the same pseudo-empirical data, i.e., simulated data not used during the training process but used as empirical data for which the true values of *s* and *T* are known. These pseudo-empirical data were simulated using the aforementioned demographic model to closely reproduce the effects of drift, migration waves, recent admixture and variations across time of the ancestry proportions characterizing recent European history, as described in the section above. They also closely reproduce the features of the empirical data used in this study (number of genotypes, age of simulated ancient individuals and pseudo-haploidy for ancient data). For each selected variant investigated in the study, we used 200 pseudo-empirical data. We adopted the same filtering as in Kerner and colleagues (2023) by focusing on pseudo-empirical data with an observed derived allele frequency higher than 0.025 in at least one of the 5 epochs used for ABC. Based on these pseudo-empirical data, we then assessed the accuracy of both the CNN and ABC estimations by comparing estimated and simulated parameter values, 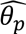 and *θ_p_*, respectively, obtained for the pseudo-empirical data *p*, using classic accuracy indices computed in cross-validation procedures: the linear correlation coefficient *r* computed between 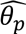 and *θ_p_*, the relative root of the mean square error, *RMSE* (i.e. the root of the *MSE* expressed as a proportion of the true value, 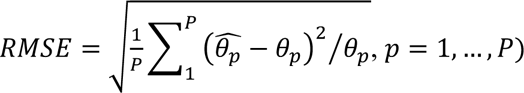, and the proportion of true values within the 90% credible intervals of estimates, 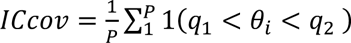 where 1(*C*) is the indicative function (equal to 1 when *C* is true, 0 otherwise) and *q*_1_ and *q*_2,_ the corresponding percentiles of the posterior distributions. To compare the accuracy indices hence obtained (*r* and *RMSE*) between ABC and CNNs we computed the two relative differences, (*r_CNN_* – *r_ABC_*)/*r_ABC_* and (*RMSE_CNN_* – *RESE_ABC_*)*RMSE_ABC_*. For real data, we also computed a similar relative difference between the CIs extent obtained for each selected variant, (*CI_CNN_* – *CI_ABC_*)⁄*CI_ABC_*.

### Data availability

Supplemental material, including Figures S1 to S17, Tables S1 to S4 (Tables S1-S4 are supplied as Excel files) and Files S1 to S5 (Files S1-S5 are supplied as pdf files) can be found with this article online. The data used in this study can be downloaded: at https://reich.hms.harvard.edu/allen-ancient-dna-resource-aadr-downloadable-genotypes-present-day-and-ancient-dna-data (v44.2) for ancient individuals, and at the 1000 Genomes Project web site https://www.internationalgenome.org/ for modern individuals. The code for simulating modern and ancient DNA data, building 1D images for training and prediction, implementing the three different CNNs architectures tested in this study and performing both predictions and cross validation tests based on simulated data can be downloaded at the GitHub code depository site (the site will soon be available). Rather than providing a non-exhaustive analysis of the impact of various demographic scenarios on the estimations, we provide user friendly tools suitable to handle other populations and other data, in order to automatically perform cross validations based on pseudo-empirical datasets simulated under the assumed demographic model, insisting on the importance to systematically perform such cross validations for every variant under investigation. In addition, the topology of the CNNs may have various impacts on the estimations, depending on the demographic scenario assumed. We thus provide user friendly tools to automatically set the CNN architecture (numbers of convolutions and maxpooling layers, parameters of the convolution, numbers of hidden layers and number of neurons per layer). Used in combination with tools allowing cross validations, it will provide an optimal solution to maximize the prediction accuracy given the specificity of the data and populations analyzed.

## RESULTS

### Deep learning improves the estimation of selection parameters

We applied our genotype-based CNN algorithms to jointly estimate *s* and *T* for each of the 89 candidate variants that were recently identified to be under positive selection over the last 10,000 years of European history (KERNER *et al*. 2023). The CNNs, which use as input data a 1D image which encodes a vector of modern and ancient genotypes sampled at various generations in the past and sorted by age, were trained with vectors of simulated genotypes. These genotypes derive from forward-in-time simulations of derived alleles under positive selection using random draws of *s* and *T* (Material and Methods) under a validated European demographic model, reproducing known ancestry variations across ages (KERNER *et al*. 2023). Importantly, the simulated 1D images must perfectly match the number, the age and the ploidy of the empirical genotypes analyzed (Material and Methods). For example, all simulated and empirical 1D images of the lactase persistence variant (rs4988235, *MCM6-LCT* region) must be of equal dimension, which in our case corresponds to each containing 1,640 pseudo-haploid ancient genotypes (1 pixel per genotype) and 503 diploid modern genotypes (2 pixels per genotype). For this example of positive selection (BERSAGLIERI *et al*. 2004; ENATTAH *et al*. 2002; HOLLOX *et al*. 2001; SEGUREL and BON 2017; TISHKOFF *et al*. 2007), the empirical 1D image with derived alleles in black (pixel=1) darkens as generations approach the present because of the rapid increase with time of the derived allele frequency due to selection (Figure 1). As hypothesized, convolutions applied to such an image by the first 1D convolution layer in the CNNs, which is equivalent to the sliding average of pixel values, produce a feature map corresponding to a fine-grained allele frequency trajectory (Figure 1; the empirical 1D images and convolutions for the 89 positively selected variants are shown in File S1). Such allele frequency trajectory with much higher number of data points than those used in ABC are the kind of hidden features captured by the CNNs that should improve the *s* and *T* predictions.

We first assessed the accuracy of *s* and *T* estimations by means of a cross validation procedure based on pseudo-empirical data, also reproducing the complexity of the European demographic history and the features of the empirical data (ploidy, number and ages of ancient genotypes, Material and Methods). Due to the low coverage of ancient samples, the number and the ages of genotypes called drastically vary across all analyzed variants. To account for such variability, we assessed the accuracy of the predictions for each of the 89 variants separately (the number of ancient genotypes available is a potential driver of the estimation accuracy). Similarly, we trained the CNNs for each variant separately, using 89 training sets of 500,000 simulated images adjusted on the number and age of the called genotypes (Material and Methods). Based on the comparisons between simulated and predicted values, we found very similar estimation accuracies for the three tested CNNs (Supplementary Figure S1, Supplementary Figures S2-4). In light of these results, we opted to focus only on the CNN1 architecture (Supplementary Figure S1) with 3 dense layers and filter dimensions of 100 genotypes (sliding average of 100 pixels, Figure 1), which tends to outperform the other two neural networks (note that it has the lowest average *RMSE*, Supplementary Figure S2). A direct comparison with the ABC approach based on the same training dataset (Material and Methods) shows that the CNN1 substantially improves prediction accuracies (Figure 2, Supplementary Figure S2). Values predicted by CNN1 are more strongly correlated with true values relative to ABC (correlation coefficients *r* between true and estimated values are ∼5% and ∼30% higher in average for *s* and *T* respectively), and RMSE is ∼25-30% lower on average, for both *s* and *T*, an improvement that is most noticeable for the age of the selection parameter (Figure 2B, Supplementary Figure S2).

**Figure 2.**
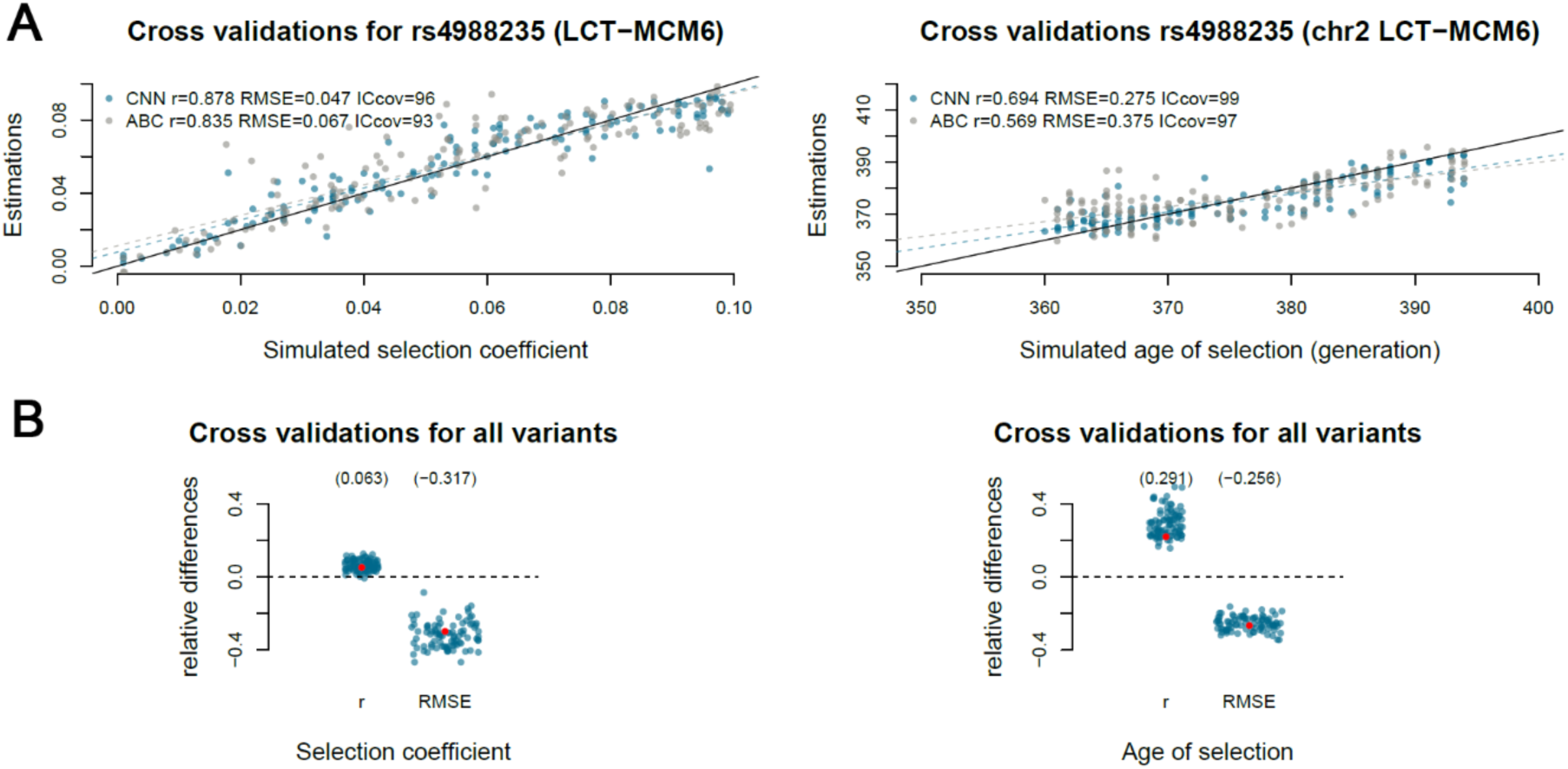
CNN estimation accuracy of the strength and timing of positive selection. (A) Simulation-based cross validations for the estimation by ABC and CNN of the selection coefficient (left panel) and the age of onset of selection (right panel), for the European lactase persistence variant (rs4988235). Dashed colored lines represent the regression line between true and predicted values obtained for each method. Black solid lines represent the identity line. The linear correlation coefficient *r*, the relative root of the mean square error *RMSE* and the proportion of true values within the credible intervals of estimates *ICcov*, are indicated on each panel. (B) Simulation-based cross validations for the estimation by ABC and CNN of the selection coefficient (left panel) and the age of onset of selection (right panel), performed for each of the 89 positively selected variants investigated in this study. Points indicate the relative difference between accuracy indices obtained with CNN and ABC for each selected variant. A positive and negative value for *r* and *RMSE* obtained with higher *r* and lower *RMSE* for CNN respectively, indicates that CNN predictions better correlate with true values, with a lower estimation variance than ABC (lower dispersal of the CNN estimates around the true values than ABC). Red points indicate the relative differences obtained for the lactase persistence allele. See also Supplementary Table S1 for detailed numeric values. Numbers indicated in brackets are the averaged *r* and *MSE* computed across the 89 positively selected variants.

To test whether our predictions are affected by errors in radiocarbon dating, we performed new cross validations based on pseudo-empirical data where we added a uniform random variable ±*δ*∼*U*(200) *ya* to the true sampling time and sorted the ancient individuals in the images accordingly (Supplementary Figure S5). These results indicate that both CNN and ABC methods are minimally affected in case of errors in radiocarbon dating. Altogether, the estimation of selection parameters by CNNs exhibits lower bias and reduced estimation variance compared to a frequency-based ABC method. This analysis shows that constraints imposed by the use of frequency-based approaches, e.g., the number and duration of past epochs, can be efficiently circumvented, with a substantial gain in accuracy, by the use of genotype-based approaches.

### Estimating the intensity and onset of selection events

Because we found *T* estimations to be more precise with CNN than with ABC, we applied the CNN1 architecture to empirical data in order to reevaluate the previous ABC-based finding that the bulk of genetic adaptations in Europe primarily occurred after the start of the Bronze Age (KERNER *et al*. 2023). Accordingly, among the 89 positively selected variants, we found that 79% (76% using ABC) of our point estimates of *T* were lower than 4,500 ya (Figure 3A), confirming the large excess of recent selection events (KERNER *et al*. 2023). When comparing CNN and ABC estimations obtained at each variant, we found a large overlap between predictions (point estimates of one method are contained in the CI of the other in ∼95% and ∼100% of cases for *s* and *T* respectively, Supplementary Table S1). Yet, deep learning tends to predict lower *s* values than ABC, in line with the lower upward bias of CNNs relative to ABC for *s* < 0.05 (Figure 2A). Although this suggests weaker selection than previously estimated (KERNER *et al*. 2023), our analysis replicated 89% of selection signals (Supplementary Table S1). The CIs around point estimates of *s* obtained with CNN and ABC are of similar order of magnitude in average (Figure 3B), suggesting that allele frequency trajectories using five data points are sufficient to well fit the overall rate of change in frequency and obtain informative ABC estimations with sharp posteriors of *s* (Figure 3C,D).

**Figure 3.**
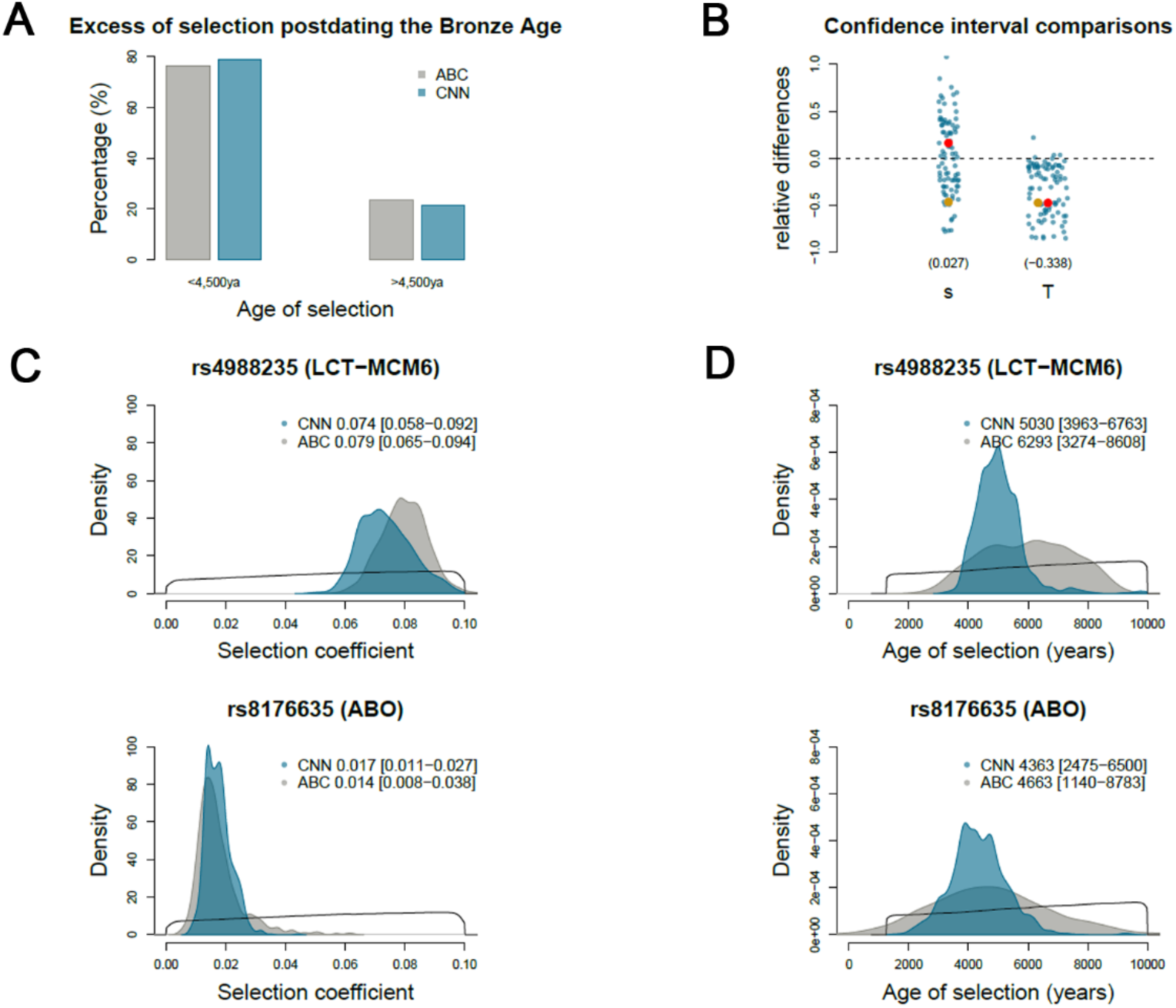
CNN estimation of the selection coefficient and the age of selection for 89 positively selected variants in Europe. (A) Percentage of variants positively selected predating (right) and postdating (left) the beginning of the Bronze Age. (B) Relative difference between the CI ranges obtained by CNN and ABC, for each selected variant. A negative value indicates that CNN provides a lower confidence interval than ABC. Posterior distributions of the selection coefficient (C) and the age of selection (D) obtained for two selected variants, the *MCM6/LCT* and the *ABO* variants. The priors are indicated with solid lines. (B) Red and gold points indicate the relative difference between the CI ranges for the two corresponding variants, respectively. See also Supplementary Table S1 for detailed numeric values. Numbers indicated in brackets are the averaged relative difference between the CI ranges computed across the cross-validations performed.

Conversely, genotype-based CNNs substantially improved the estimation of the age of selection, with a reduced uncertainty around point estimates of *T* in virtually all the cases. We obtained ∼30% reduction in the length of CIs in average, with up to an ∼80% reduction in some cases (reductions by more than 70% are sometimes due to negative CI bounds, a numerical issue that often arises in ABC estimations, Figure 3B, Supplementary Table S1). Notably, for some iconic cases of selection, such as the ABO blood group system or the lactase persistence variant, we obtained a two-fold reduction in the length of the CIs (Figure 3C,D). Our CNN predictions for the lactase persistence allele, highly overlapping that of *s* using ABC (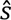*_CNN_* = 0.074 [0.058 − 0.092] *vs* 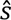*_ABC_* = 0.082 [0.066 − 0.096]) (KERNER *et al*. 2023), substantially improved the estimation of the age of selection (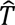*_CNN_* = 5,030 [3,963 − 6,763] *vs* 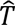*_ABC_* = 6,102 [3,197 − 8,547]) (KERNER *et al*. 2023) in the light of other aDNA-based reports pointing to an onset of the Eurasian lactase persistence allele at the beginning of the Bronze Age (ALLENTOFT *et al*. 2015; BURGER *et al*. 2020; MATHIESON *et al*. 2015). Finally, when re-estimating *T* with ABC but with a higher number of epochs, and thus of summary statistics (12 epochs of ∼1000 years each *vs* 5 in the initial ABC estimations), we still found that accuracy (*r* and *MSE*) was consistently lower than that obtained with CNN1 (Supplementary Figure S6). On real data, these new ABC estimations still showed a large uncertainty around point estimates with CIs being consistently larger than those obtained with CNN1 (Supplementary Figure S7). Collectively, our results confirm that genotype-based CNNs improve ABC predictions of the time of onset of selection.

### Detecting selection at the subcontinental scale

To delineate possible differences in selection events at the local scale, we next investigated the strength and timing of selection for the 89 selected variants in two different regions of Europe. We used the same CNN1 architecture trained on ancient genomes from northern (north of 45°N) and southern (south of 45°N) Europe separately (KERNER *et al*. 2021; MATHIESON and MATHIESON 2018), and on modern genomes of FIN, GBR and CEU 1000G populations, as well as IBS and TSI 1000G populations, respectively. For the lactase persistence variant, for example, the empirical images built with northern and southern European genotypes are reduced by ∼45% and ∼55% relative to the full dataset (Figure 1, other images are shown in Files S2,3). Our analysis based on pseudo-empirical data shows that genotype-based CNNs remain more accurate than ABC, even when applied to a reduced sample of ancient individuals, for both the intensity and age of selection (e.g., a ∼25% reduction of the RMSE; Supplementary Figures S2,8,9). As expected, cross-validation analyses showed that the overall accuracy was diminished when compared to the use of the full dataset (Supplementary Figure S2, Supplementary Tables S2,3).

In both northern and southern Europeans, we observed an excess of selection times postdating the Bronze Age (71% and 73% of point estimates < 4,500 ya, respectively, Supplementary Figure S10), with substantially improved estimations compared to ABC for both the *s* and *T* predictions in virtually all cases (∼20% to ∼40% reduction of CIs of the *s* and *T* estimations in average for the 89 positively selected loci, Supplementary Figure S10, Supplementary Tables S2,3). Whereas ABC failed to accurately estimate timing and intensity of selection for the lactase persistence variant in southern Europe (see the flat posteriors in Figure 4), the CNN consistently estimates the onset of selection at the beginning Bronze Age in both subcontinental regions (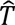*_CNN_* = 5,721 [3,963 − 8,713] and 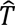*_CNN_* = 5,268 [3,787 − 7,987] in northern and southern Europe respectively, Figure 4). Our predictions highlight a lower intensity of selection in the south (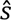*_CNN_* = 0.057 [0.036 − 0.089]) than in the north (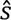*_CNN_* = 0.074 [0.055 − 0.096]), mirroring the signal captured by the empirical 1D images (Figure 1). In agreement with previous analysis based on ancient DNA data (OLALDE *et al*. 2019), our results support that the lactase persistence variant increased due to selection in southern Europe, but did not become as common as in northern Europe, possibly because of different agricultural practices (BEJA-PEREIRA *et al*. 2003). We found other examples of selection episodes with contrasted strengths across subcontinental regions (rs8103030 downstream a long noncoding RNA, CTB−175P5.1, 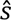*_CNN_* = 0.02 [0.003 − 0.048] and 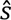*_CNN_* = 0.053 [0.015 − 0.095] in northern and southern Europe respectively, Supplementary Tables S2,3). Interestingly, the rs10765770 variant (*GRM5*) associated with pigmentation (MATHIESON *et al*. 2015) shows the opposite pattern, suggesting two separate selection events of similar strength at two different epochs (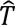*_CNN_*= 8,653 [6,500 − 10,000] ya and 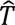*_CNN_* = 5,300 [2,475 − 9,475] ya for the north and south, respectively). This difference was reassessed, due to rather imprecise predictions and posteriors skewed toward the upper prior bounds in southern and northern Europe, respectively (see the next section).

**Figure 4.**
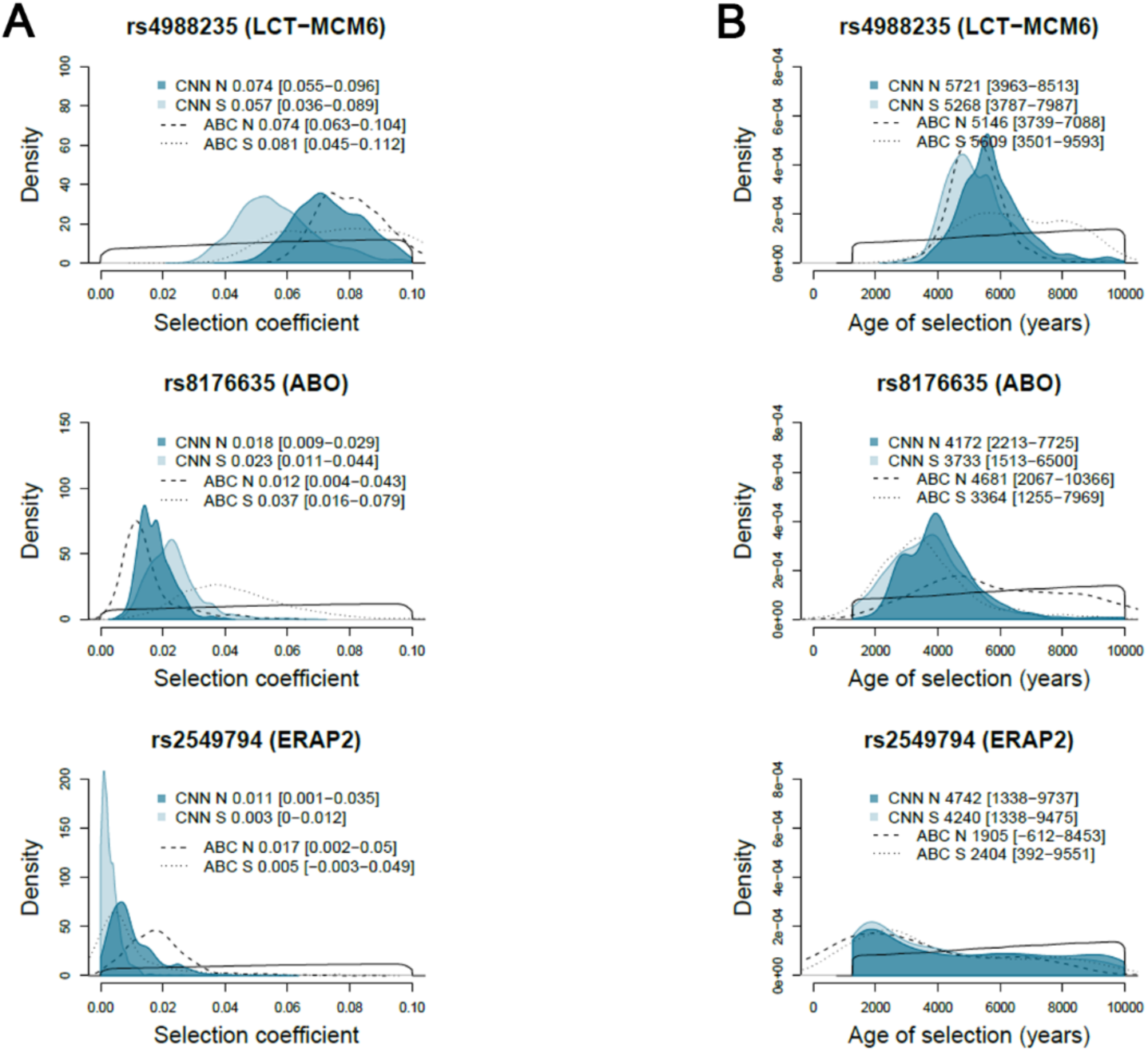
Strength and timing of positive selection in northern and southern Europe. Posteriors distributions of the selection coefficient (A) and the age of selection (B) obtained for three selected variants, the lactase persistence and the *ABO* variants, together with the *ERAP2* variant under selection due to the Black Death pandemic in northern Europe. The priors are indicated with solid lines. N and S stand for northern and southern Europe, respectively. See also Supplementary Tables S2,3 for detailed numeric values.

Finally, known cases of selected variants associated with host defense against pathogens showed similar intensities and ages of selection in northern and southern Europe, including rs8176635 (*ABO*; 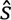*_CNN_* ≈ 0.02, 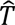*_CNN_* 4,000 ya) (Figure 4) and rs10008492 (*TLR10-TLR1-TLR6* gene cluster; 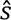*_CNN_* ≈ 0.03, 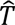*_CNN_* ≈ 8,500 ya,). Consistently with recent findings reporting positive selection due to the Black Death pandemic in northern medieval Europe (KLUNK *et al*. 2022), the variant rs2549794 (*ERAP2*) exhibits a significant signal of selection in northern Europe, with CIs excluding neutrality (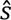*_CNN_* = 0.011 [0.001 − 0.035] and 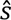*_ABC_* = 0.017 [0.002 − 0.05], Figure 4, File S4). The relatively imprecise prediction of *T* points to selection starting ∼2,000 ya, or ∼1,000 ya when assuming an onset of selection of 2,000 years in the past at most, a period that includes the Justinian plague and the medieval Black Death pandemics in Europe (Supplementary Figure S11). To confirm, we analyzed the splice region SNP rs2248374, a variant directly controlling the ERAP2 expression (VENEMA *et al*. 2023). Independently of the continental subregion studied and the methods used, we found significant signals of positive selection targeting the allele producing full-size ERAP2 protective against Black Death, with CNN predictions also pointing to recent selection starting ∼2,000 ya (Supplementary Figure S12).

### Robustness to training

To check whether our CNN method is sensitive to model training and prior parameter ranges, we re-estimated the *s* and *T* parameters for all the 89 positively-selected variants by training the model with simulations where the uniform prior of *T* was extended to 30,000 ya, and checked if the predictions of *s* and *T* were affected. This analysis showed that 95% of the initial point estimates of the intensity of selection are contained in the CIs of the new set of predictions (Supplementary Figure S13). We then focused on two variants associated with skin pigmentation, in *GRM5* and *SLC45A2*, for which the age of selection is expected to predate 10,000 years. For rs185146 (*SLC45A2*), selection was previously dated to 10,680 to 36,070 ya (LOPEZ *et al*. 2014), although our initial predictions indicated more recent selection (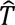*_CNN_* = 5,156 [4,225 − 6,237] ya and 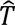*_CNN_* = 3,947 [2,737 − 5,537] ya in northern and southern Europe respectively). For rs10765770 (*GRM5*), the CNN posteriors were skewed toward the upper prior bound in northern Europe, suggesting a selection episode pre-dating 10,000 years (Supplementary Figure S14). As expected, the prediction of the age of selection varied for rs10765770 (*GRM5*), with the new analysis indicating selection predating the Neolithic. In contrast, our first prediction was confirmed for rs185146 (*SLC45A2*), mainly in southern Europe (Supplementary Figure S15). Independently of the priors used, for *SLC45A2* our results support a recent episode of selection perhaps on top of an older selection event that started before 10,000 ya, (LOPEZ *et al*. 2014), as suggested by the sudden increases in frequency over a monotonous growth that can be visualized in the genotype-based 1D images (red arrows in Supplementary Figure S16). Importantly, in all cases, the *s* predicted remained similar, ∼0.02 and ∼0.05 for *GRM5* and *SLC45A2* respectively, suggesting that CNN, as also ABC, is robust to model training, even when *T* predictions were imprecise.

### Footprints of time-dependent negative selection

Because ancient DNA data can also be used to estimate the intensity and age of negative selection (KERNER *et al*. 2021; KERNER *et al*. 2023), we trained the CNN algorithm (CNN1, Supplementary Figure S1) with simulated images of variants with derived alleles under negative selection. Specifically, we aimed at predicting the selection parameters for a set of 50 missense variants at conserved positions (GERP score >4) that we previously found to be deleterious in terms of reproductive success (KERNER *et al*. 2023). The procedure used to generate 1D images of empirical and simulated genotypes was the same as described before (1D images used for the negatively selected variants are shown in File S5). In the case of negative selection, consistent with a significant decrease in derived allele frequency over generations, the image is expected to become lighter with time due to the loss of the derived allele. Before predicting selection parameters on real variants, we first checked whether the trained CNN1 accurately predicts parameters underlying simulated pseudo-empirical 1D images by applying the same cross-validation procedure use for positive-selected variants to each negatively-selected variant (Supplementary Figure S17, Supplementary Table S4). With respect to ABC, the CNN1 algorithm improved again the estimation of selection parameters, with similar or higher correlations between estimates and true values, together with a ∼10% and ∼20% reduction in RMSEs for *s* and *T*, respectively (Supplementary Figure S17). For example, the cross validations based on simulated data performed for the tuberculosis risk variant rs34536443 (*TYK2*) showed that CNN estimation of *T* is less biased and the estimation variance is smaller for both *s* and *T* (Supplementary Figure S17). These results indicate, as for positive selection parameters, that leveraging individual genotypes improves parameter estimation under models of time-dependent selection.

When analyzing the 50 negatively-selected variants, we confirmed the excess of recent selection postdating the start of the Bronze Age (KERNER *et al*. 2023) (Figure 5). We also found a perfect overlap of point estimates for both the strength and the timing of selection, with 100% of point estimates obtained in one method contained in the CI of the other (Supplementary Table S4). Importantly, as for positive selection, the genotype-based CNN1 reduced the CIs of the age of selection compared to ABC and consequently, the estimations of *T* were substantially improved, with a reduction of the CIs by ∼20% on average. Likewise, we replicated the signals of selection for the six negatively selected variants identified in immunity genes, rs34536443 (*TYK2*), rs3803716 (*TNRC6A*), rs12146727 (*C1S*), rs2232607 (*LBP*), rs11209026 (*IL23R*) and rs3775291 (*TLR3*) (Figure 5, Supplementary Table S4). For example, the tuberculosis risk variant rs34536443 (*TYK2*) shows a significant signal of negative selection, consistent with ABC estimations, (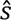*_CNN_* = 0.015 [0.002 − 0.035] and 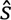*_ABC_* = 0.013 [0.004 − 0.036]) over the last 2,000 years (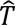*_CNN_* = 2,917 [1,338 − 6,500] ya and 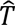*_ABC_*= 1,805 [324 − 8,402] ya) (Figure 5).

**Figure 5.**
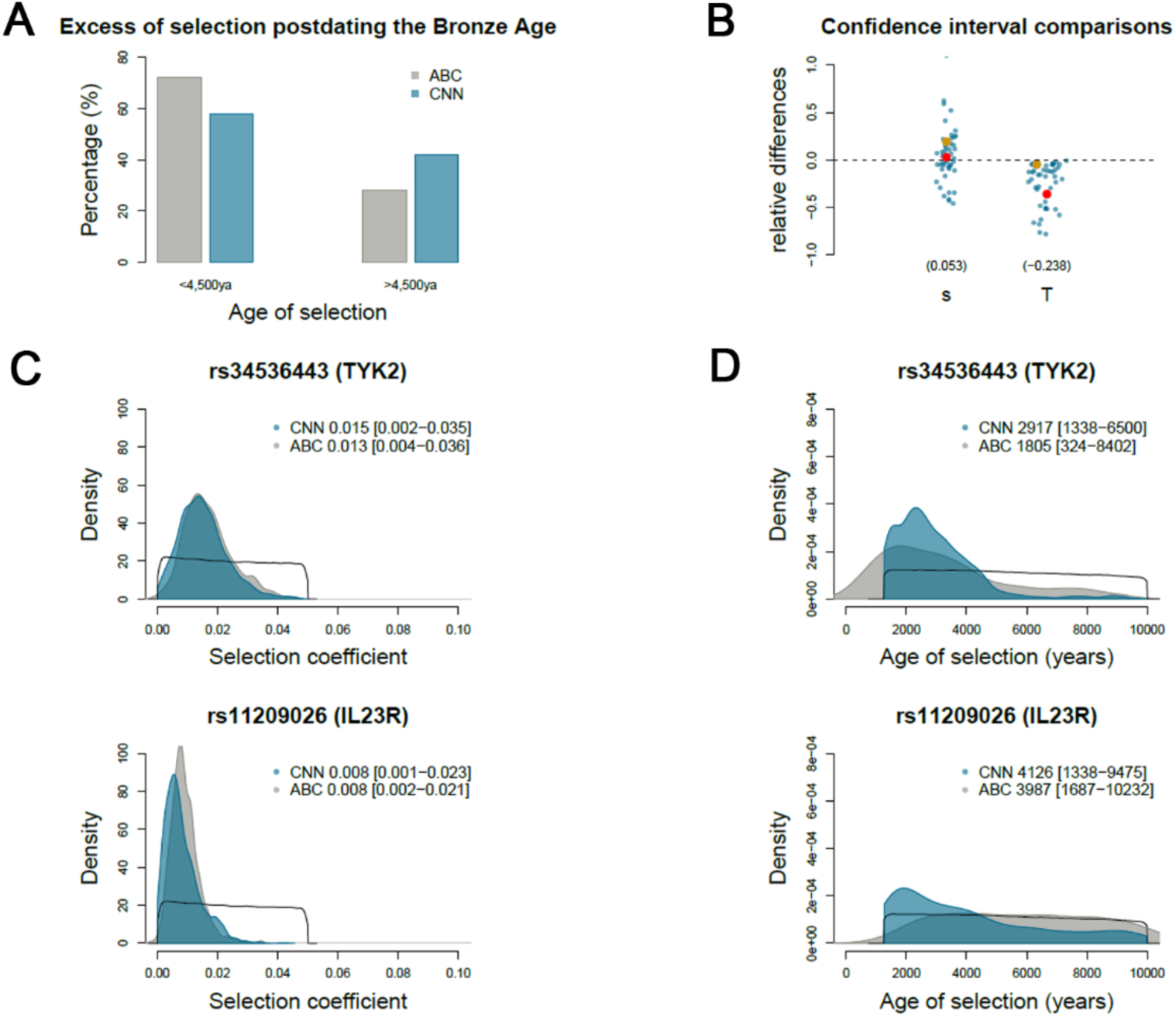
CNN estimation of the selection coefficient and the age of selection for 50 negatively selected variants in Europe. (A) Percentage of variants negatively selected predating (right) and postdating (left) the beginning of the Bronze Age (B) Relative difference between the CI ranges obtained by CNN and ABC, for each selected variant. Posteriors distribution of the selection coefficient (C) and the age of selection (D) obtained for two selected variants, the *TYK2* and the *IL23R* variants. The priors are indicated with solid lines. (B) Red and gold points indicate the relative difference between the CI ranges for the two corresponding variants, respectively. See also Supplementary Table S4 for detailed numeric values. Numbers indicated in brackets are the averaged relative difference between the CI ranges computed across the cross-validations performed.

## DISCUSSION

Here, we present new CNN algorithms that we systematically compared to an ABC frequency-based approach for the estimation of key selection parameters. Using pseudo-empirical data reproducing aDNA data, simulated under complex demography including variation of effective population sizes, episodes of intense migrations and variation in time of individual ancestry, we show that genotyped-based CNNs provide predicted values that are closer to true values (Supplementary Figure S2). When applied to real data, CNN also provides decreased uncertainty around point estimates for both the intensity and time of onset of selection (Figures 3-5). Notably, CNNs do not provide point estimates or CIs outside the bounds of the prior distributions, which is a technical issue commonly encountered in ABC estimation (KERNER *et al*. 2023).

Genotype-based CNNs overcome two important limitations of ABC or similar frequency-based approaches. First, it minimizes loss of information by reincorporating in the CNN predictions 163 ancient individuals from two epochs for which sample sizes were low (KERNER *et al*. 2023). Second, it improves the inference of the first stages of selection, i.e., the time at which selection starts: while allele frequency trajectories assessed with few data points are sufficient to fit the overall rate of frequency change, informing the intensity of selection, such a low number of points prevents accurate inference of the first stages of selection. For example, simulated selection events occurring in consecutive epochs, and potentially spaced by several thousands of years, may similarly fit the data during the ABC training, preventing an accurate estimation of the time of the onset of selection.

Conversely, CNN algorithms leverage the fine-grained frequency trajectories hidden in the genotype-based images, resulting in more accurate predictions of both the intensity and the timing of selection. Third, our CNN results not only confirmed previously reported selection signals in Europe but also refined the predictions of selection parameters when using reduced numbers of individuals. For example for the European lactase persistence variant, CNNs evidenced a reduced intensity of selection in southern Europe (*s* = 0.057 *vs*. 0.074) while ABC estimated higher intensity of selection than in the north in disagreement with the data (see allele frequency trajectories and empirical 1D images in Figure 1).

Our study confirmed the importance of the post-Neolithic period in the adaptive history of Europeans, as most positive and negative selection events postdate the beginning of the Bronze Age (KERNER *et al*. 2023), both in northern and southern Europe. This result support an increase in genetic adaptations following the Neolithic period, likely due to population growth, which promoted the emergence of selected mutations and increased selection efficacy. Our predictions for the ABO blood group system (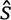*_CNN_* ≈ 4,000 ya with similar intensities in northern and southern Europe, (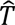*_CNN_* ≈ 0.02) confirm that the frequency of the A, AB and, particularly, B groups has increased through positive selection over the last 4,000 years (KERNER *et al*. 2023), and support the notion that past epidemics are responsible for major changes in ABO blood groups in different parts of the world (ABEGAZ 2021). Interestingly, our analysis support recent data reporting a positive selection signal at the rs2549794 (*ERAP2*) due to the second Black Death pandemic in northern Europe (KLUNK *et al*. 2022). However, the intensity of selection estimated is much lower compared to the previous finding independently of the priors on *T* used (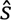 = 0.39 *vs* 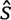*_CNN_* = 0.017 and also 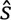*_ABC_* = 0.022 assuming onset of selection until 2,000 years in the past at most, Supplementary Figure S11). These discrepancies are likely explained by our parsimonious assumption of a constant *s* from the onset of selection, whereas plague epidemics were more sporadic and of very high intensity during short time periods. A selection coefficient close to 0.02 assuming constant selection over the last 2,000 years is consistent with a selection coefficient close to 0.4 (= 0.02 × 2000⁄100) assuming selection during a 100-year-long pandemic, suggesting that higher sample sizes in the Middle Ages are necessary to refine the estimations of the intensity of selection due the European Black Death pandemic.

In the context of disease risk, we confirmed that ancient DNA data is also valuable to estimate the intensity and age of negative selection at risk variants (KERNER *et al*. 2021; KERNER *et al*. 2023). Using CNN, we replicate the signals of selection of the six negatively selected variants identified in immunity genes in a previous work, three of them experimentally validated (KERNER *et al*. 2023). We refined the estimations of the variant rs34536443 (*TYK2*), previously shown to underlie clinical TB (BOISSON-DUPUIS *et al*. 2018; KERNER *et al*. 2021; KERNER *et al*. 2019), confirming that it evolved under time-dependent negative selection over the last 2,000 years, probably due to an increase in the pressures imposed by *Mycobacterium tuberculosis*. The present analysis thus confirmed that TB has recently imposed a heavy burden in Europeans (KERNER *et al*. 2021), showing again that imprints of natural selection left on ancient genomes are key to dissect the past history of severe, human diseases.

In conclusion, the genotype-based CNN algorithms presented here are suitable to conduct a detailed analysis of natural selection on candidate variants, taking into account aDNA data features and the demography of the studied populations. However, our approach suffers from some limitations. First, our simulated training datasets assume a constant selection coefficient (*s*) from the start of selection and population continuity since the Bronze Age. These limitations, inherited from the simulation model previously used (KERNER *et al*. 2023), may be circumvented by training the CNNs in more complex simulation scenarios. Second, variations in genotype call rate among different variants require separate model training for each variant, making it challenging to scale CNN analyses on a genome-wide level (∼2h per variant for model training). In such cases, ABC can be conducted genome-wide to detect selected variants, with CNN applied to refine the estimation on the strongest candidates. Alternatively, analyses of high-coverage ancient genomes, combined with genotype imputation, may allow the CNN model to be applied at the genome-wide scale through a single training process. In such situations, CNN predictions require less computation time compared to ABC and can be easily employed to estimate the selection coefficient and CI of millions of variants, enabling a detailed detection of natural selection over the past millennia of human history.

## Supporting information

1D images for the 89 positively selected variants

1D images for the 89 positively selected variants in northern Europe

1D images for the 89 positively selected variants in southern Europe

1D images for the positively selected variants associated to Black Death pandemic (ERAP2)

1D images for the 50 negatively selected variants

## ACKNOWLEDGMENTS

We acknowledge the help of the HPC Core Facility of Institut Pasteur for this work. This work was supported by the *Institut Pasteur*, the *Collège de France*, the *Centre Nationale de la Recherche Scientifique* (CNRS), the *Agence Nationale de la Recherche* (ANR) grants LIFECHANGE (ANR 17 CE12 0018 02), CNSVIRGEN (ANR-19-CE15-0009-02) and MORTUI (ANR-19-CE35-0005), the French Government’s *Investissement d’Avenir* program, *Laboratoires d’Excellence* “Integrative Biology of Emerging Infectious Diseases” (ANR-10- LABX-62-IBEID) and “*Milieu Intérieur*” (ANR-10-LABX-69-01), the *Fondation pour la Recherche Médicale* (Equipe FRM DEQ20180339214), the *Fondation Allianz-Institut de France*, and the *Fondation de France* (no. 00106080). G.K. is supported by a Pasteur-Roux-Cantarini fellowship.

## AUTHOR CONTRIBUTIONS

G.L conceived and designed the study. G.L. was the lead analyst, with important contributions from G.K., and E.P. G.L. wrote the manuscript with important contributions from E.P., G.K. and L.Q.-M.

## DECLARATION OF INTERESTS

The authors have no competing interests to declare.

## INCLUSION AND DIVERSITY

We support inclusive, diverse, and equitable conduct of research.

## Notes

### Competing Interest Statement

The authors have declared no competing interest.

