## Supplemental Figures S1-S17 for "Deep estimation of the intensity and timing of selection from ancient genomes"

### Figure S1. CNN architectures

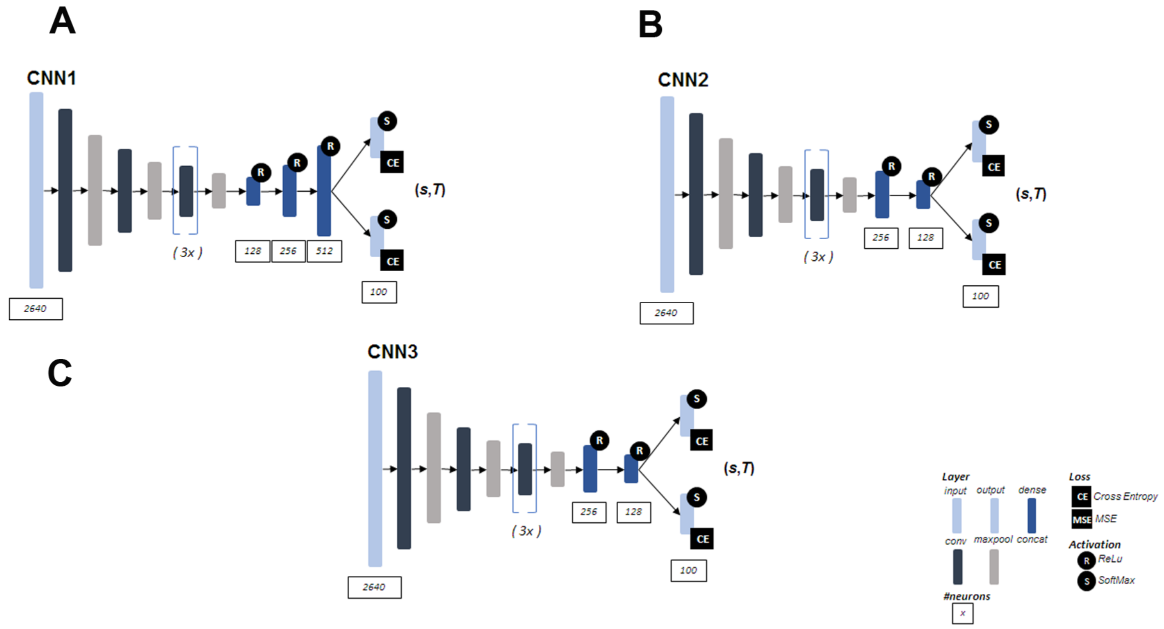


Convolutional neural networks tested in this study. The CNN1 (A) with kernel size in the first convolution layer of 100 pixels (or alleles) differs by the number and size of the dense hidden layers. CNN2 (B) and CNN3 (C) with the same architecture differ by the kernel size in the first convolution layer, 100 and 20 pixels for CNN3 and CNN3 respectively (convolutions with 20 and 100 alleles are shown in Figure 1 and in Files S1-5).

**Figure S2. Cross validations for the positively and negatively selected variants**

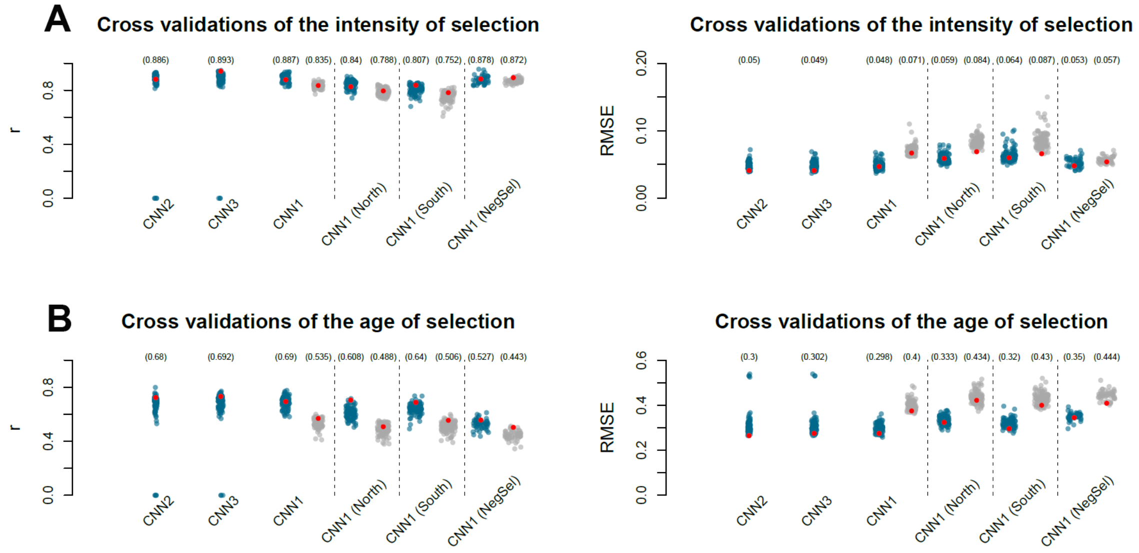

Accuracy of the CNN predictions performed for the 89 positively selected and the 50 negatively selected variants analyzed in this study. Cross validations for the selection coefficient (A) and the age of selection (B) based on pseudo-empirical datasets simulated under the European demographic history and reproducing the empirical data used (ploidy, number and age of ancient genotypes observed at each variant). Two accuracy indices are displayed for each variant: the linear correlation coefficient *r* between true and predicted values (left panels) and the relative root of the mean square error *RMSE* (right panels), each computed using 200 pseudo-empirical datasets. Blue points are the accuracy indices computed for each variant (the numbers indicated in brackets are the averages computed across variants). The CNN1, CNN2, CNN3, CNN1 (North) and CNN1 (South) labels indicate the cross validations for the 89 positively selected variants. The CNN1, CNN2 and CNN3 labels show the accuracy of the CNN predictions obtained with the corresponding algorithms and all modern and ancient Europeans used in this study (the corresponding ABC *r* and *RMSE* are displayed in grey for comparison). The CNN1 (North) and CNN1 (South) labels show the accuracy obtained with CNN1 predictions based on northern and southern Europeans respectively (the corresponding ABC *r* and *RMSE* are displayed in grey in each case). The CNN1 (NegSel) labels shows the cross validations performed for each of the 50 negatively selected variants analyzed in this study. Compared to ABC, higher *r* together with lower *RMSE* indicate a better accuracy with CNN predictions closer to true values (see the red points highlighting the accuracy indices obtained for the European lactase persistence variant rs4988235, MCM6-LCT region, in case of positive selection, and for the tuberculosis risk variant rs34536443, *TYK2* gene, in case of negative selection). Note that, in some cases the CNN2 and CNN3 architectures with two fully connected layers did not converge (see the correlation coefficients *r* between true and estimated values equal to 0 in A and B). These cases were excluded from the computation of the averaged *r* and *MSE* indicated in brackets.

**Figure S3. Predictions of the strength and timing of positive selection based on the CNN2 architecture**

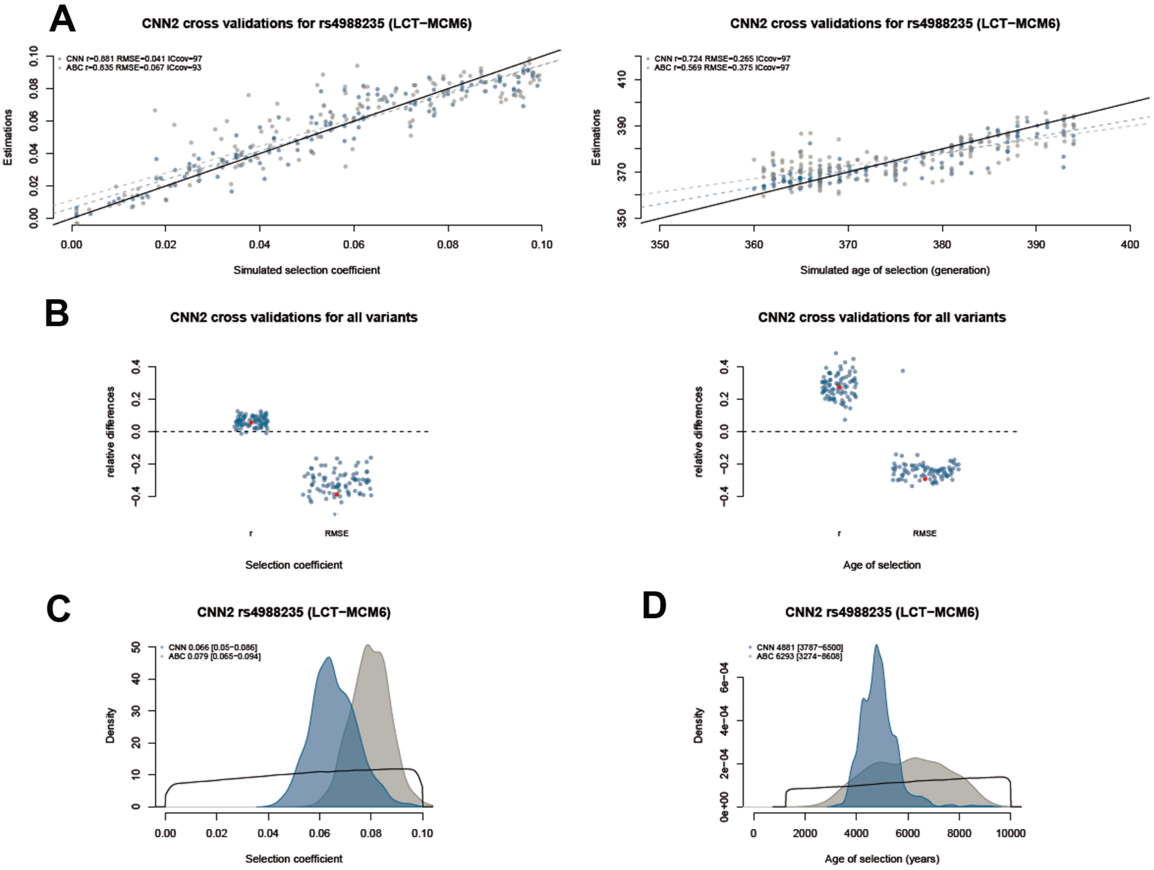


Simulation-based cross validations for the selection coefficient (left panel) and the age of selection (right panel) performed for the European lactase persistence variant (A) and for each of the 89 positively selected variants investigated in this study (B). Posteriors distributions of the selection coefficient (C) and the age of selection (D) obtained for the lactase persistence variant. See the legends of Figures 2 and 3.

**Figure S4. Predictions of the strength and timing of positive selection based on the CNN3 architecture**

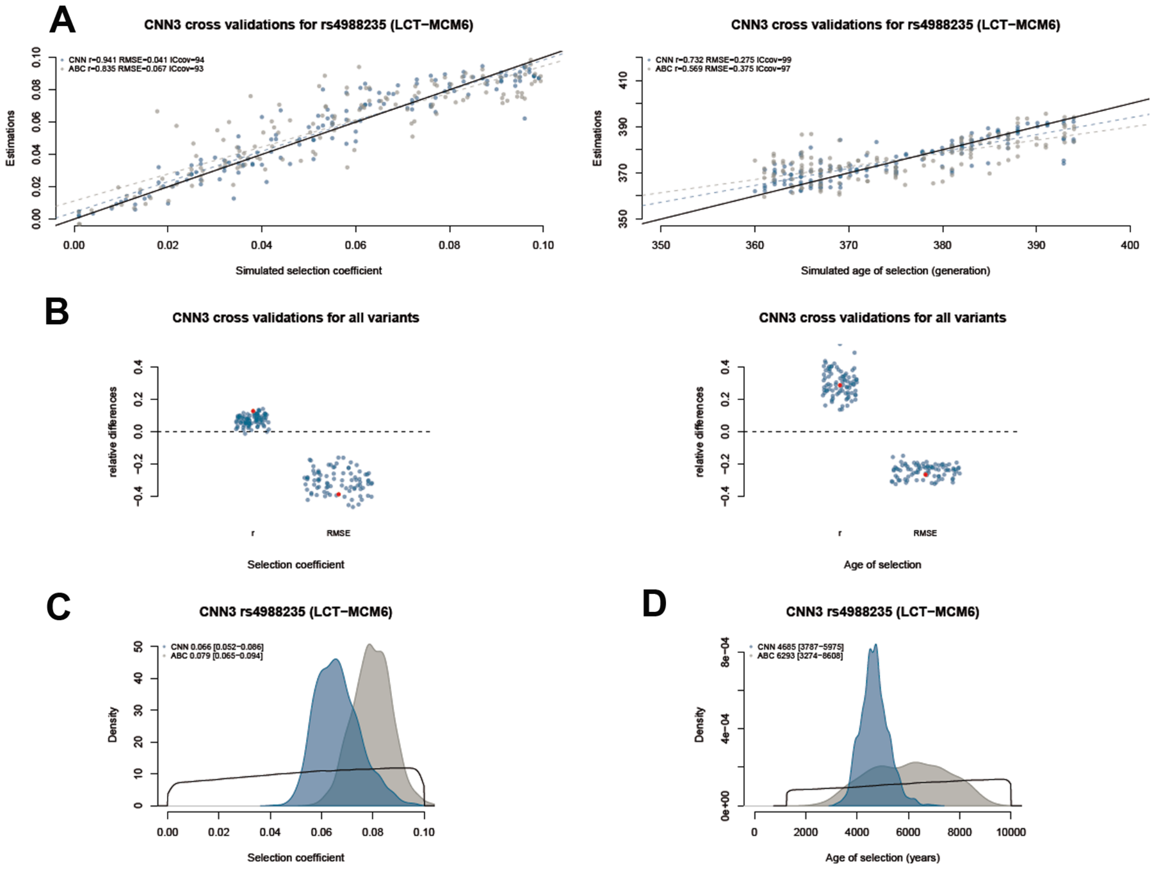


See the legend of Figure S3.

#### Figure S5. Sensitivity to radiocarbon dating errors

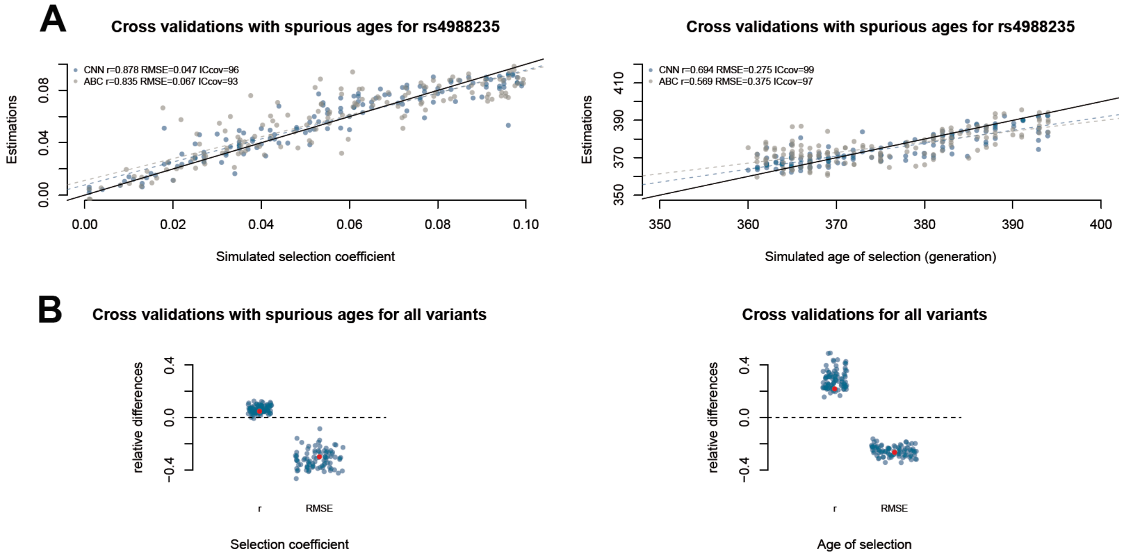


Simulation-based cross validations for the selection coefficient (left panel) and the age of selection (right panel) performed for the European lactase persistence variant (A) and for each of the 89 positively selected variants investigated in this study (B). The simulated ages in pseudo-empirical data were changed by adding a uniform random variable $\pm\delta \sim U(200) ya$. The ancient individuals were permutated in the 1D simulated images accordingly. See the legend of Figure 2.

**Figure S6. Accuracy of ABC predictions based on higher number of epochs**

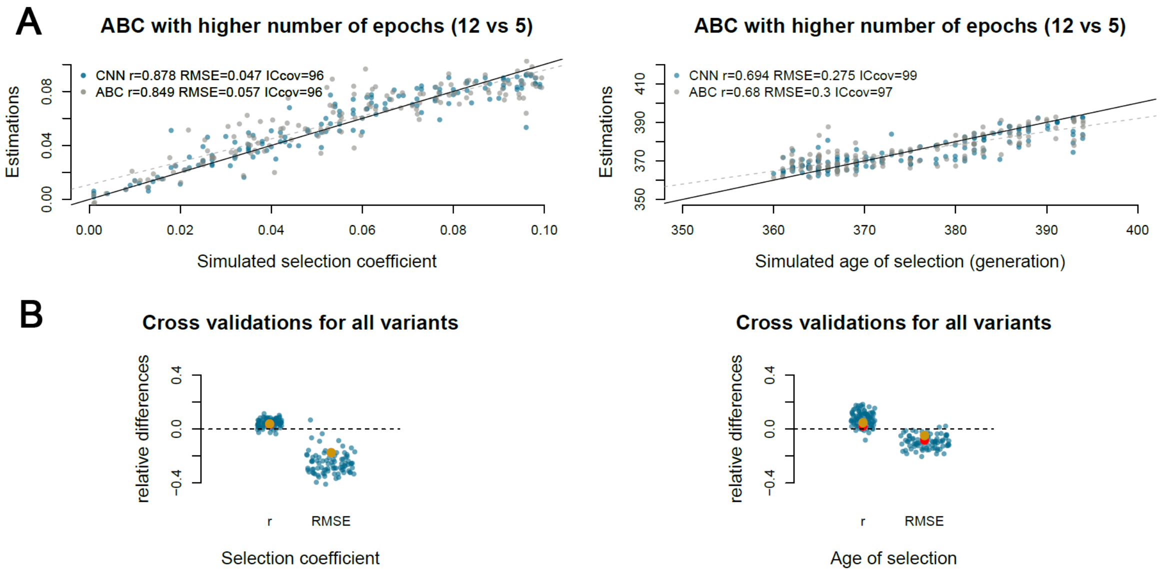


(A) Simulation-based cross validations for the estimation by ABC performed with higher number of epochs for the European lactase persistence variant (12 epochs of ~1000 years each *vs* 5 in the initial ABC estimations). The CNN results are those shown in Figure 2. (B) Simulation-based cross validations performed for each of the 89 positively selected variants investigated in this study. The comparisons between the ABC with higher number of epochs and CNN are done with the CNN results used in Figure 2. See the legend of Figure 2.

**Figure S7. ABC predictions based on higher number of epochs for the 89 positively selected variants in Europe**

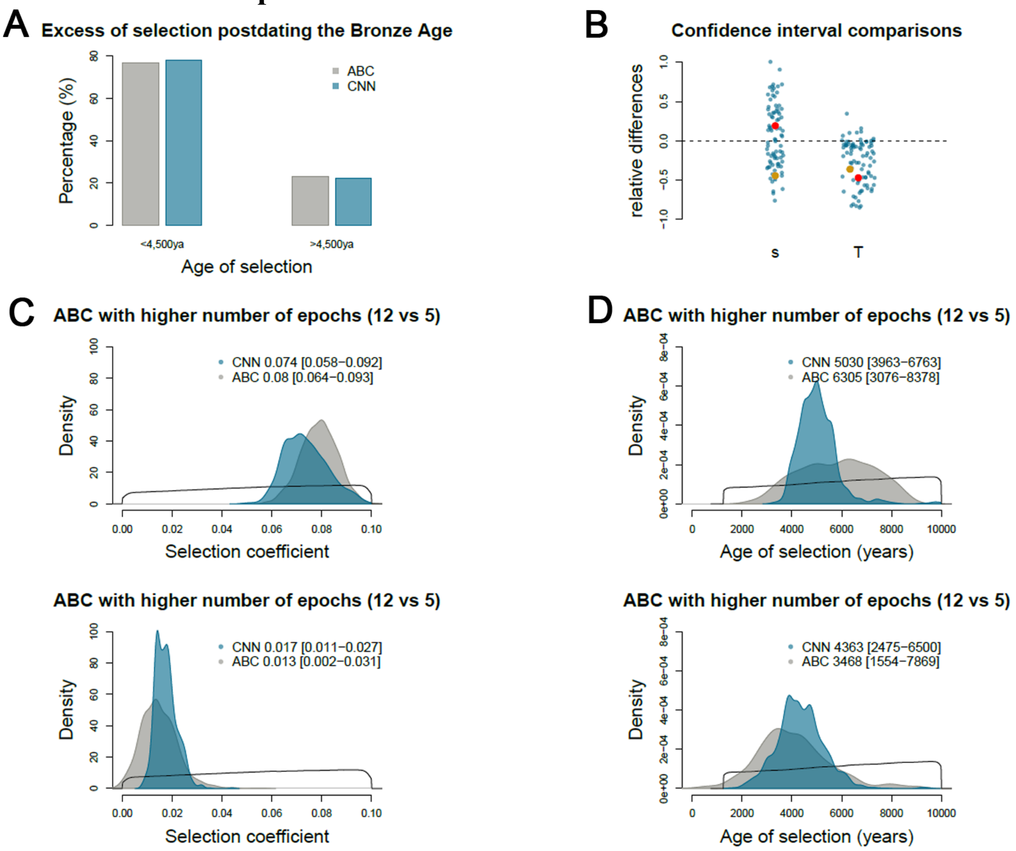


ABC performed with higher number of epochs (12 epochs of ~1000 years each *vs* 5 in the initial ABC estimations) (A) Percentage of variants positively selected predating (right) and postdating (left) the beginning of the Bronze Age. (B) Relative difference between the CI ranges obtained by CNN and ABC, for each selected variant. A negative value indicates that CNN provides a lower confidence interval than ABC. Posterior distributions of the selection coefficient (C) and the age of selection (D) obtained for two selected variants, the MCM6/LCT and the ABO variants. The CNN results are those shown in Figure 3. See the legend of Figure 3.

**Figure S8. Estimation accuracy of the strength and timing of positive selection in northern Europe**

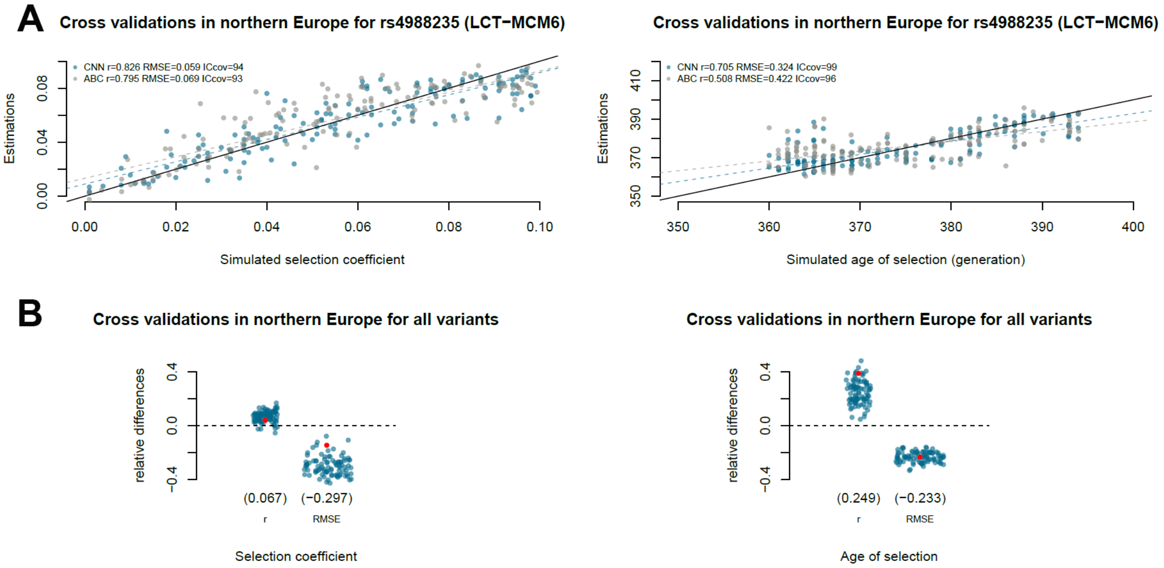


CNN and ABC predictions based on ancient and modern individuals located in northern Europe. Simulation-based cross validations for the selection coefficient (left panel) and the age of selection (right panel) performed for the European lactase persistence variant (A) and for each of the 89 positively selected variants investigated in this study (B). See the legend of Figure 2.

**Figure S9. Estimation accuracy of the strength and timing of positive selection in southern Europe**

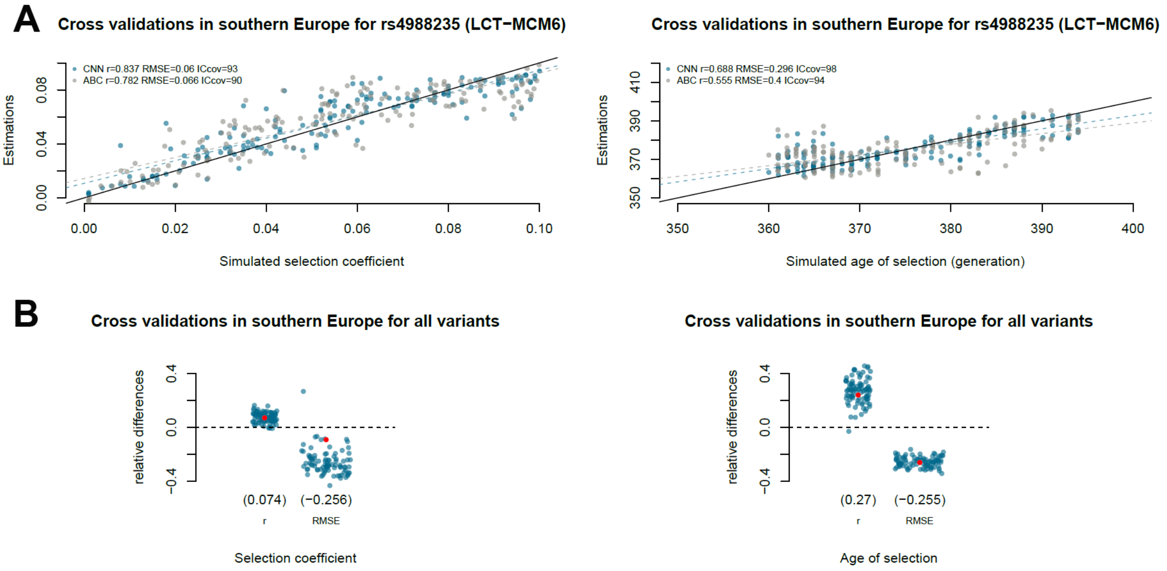


CNN and ABC predictions based on ancient and modern individuals located in southern Europe. Simulation-based cross validations for the selection coefficient (left panel) and the age of selection (right panel) performed for the European lactase persistence variant (A) and for each of the 89 positively selected variants investigated in this study (B). See the legend of Figure 2.

**Figure S10. Post Bronze Age positive selection in northern and southern Europe**

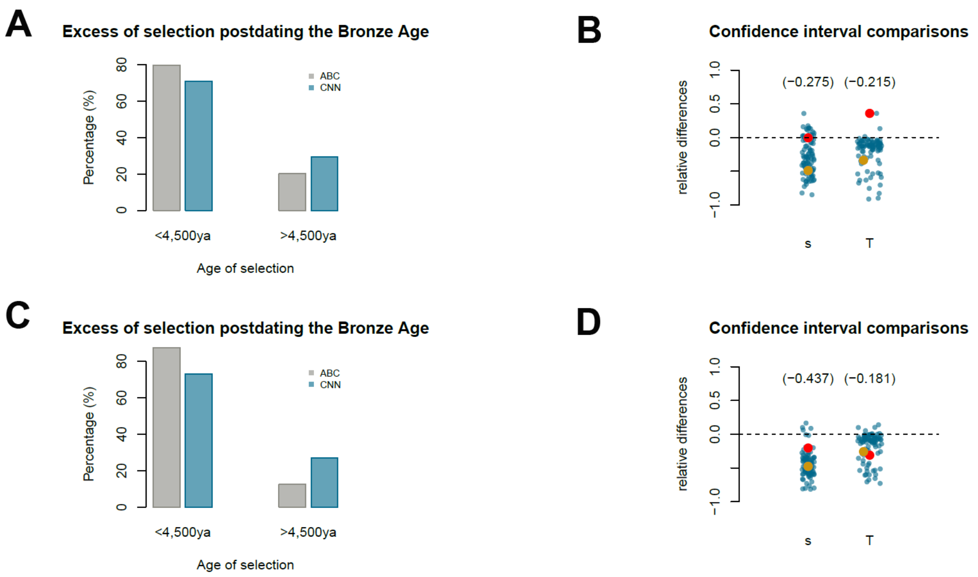


CNN and ABC predictions obtained in northern (A,B) and southern (C,D) Europe. (A,C) Percentage of the positive selection predating (right) and postdating (left) the beginning of Bronze Age. (B,D) Plots of the relative difference between ICs obtained with CNN and ABC for each of the 89 positively selected variants investigated in this study. See the legend of Figure 3.

**Figure S11 Predictions for rs2549794 (ERAP2) assuming more recent selection in northern Europe**

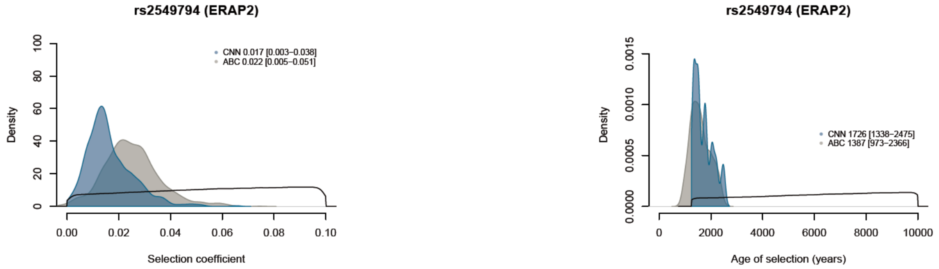


CNN and ABC predictions obtained assuming constant selection up to 2,000 years at most, i.e., the simulated ages of selection exceeding 2,000 ya in the prior distribution were excluded from the CNN and ABC training. See the legend of Figure 2.

**Figure S12 Predictions of the strength and age of positive selection targeting rs2248374 (ERAP2)**

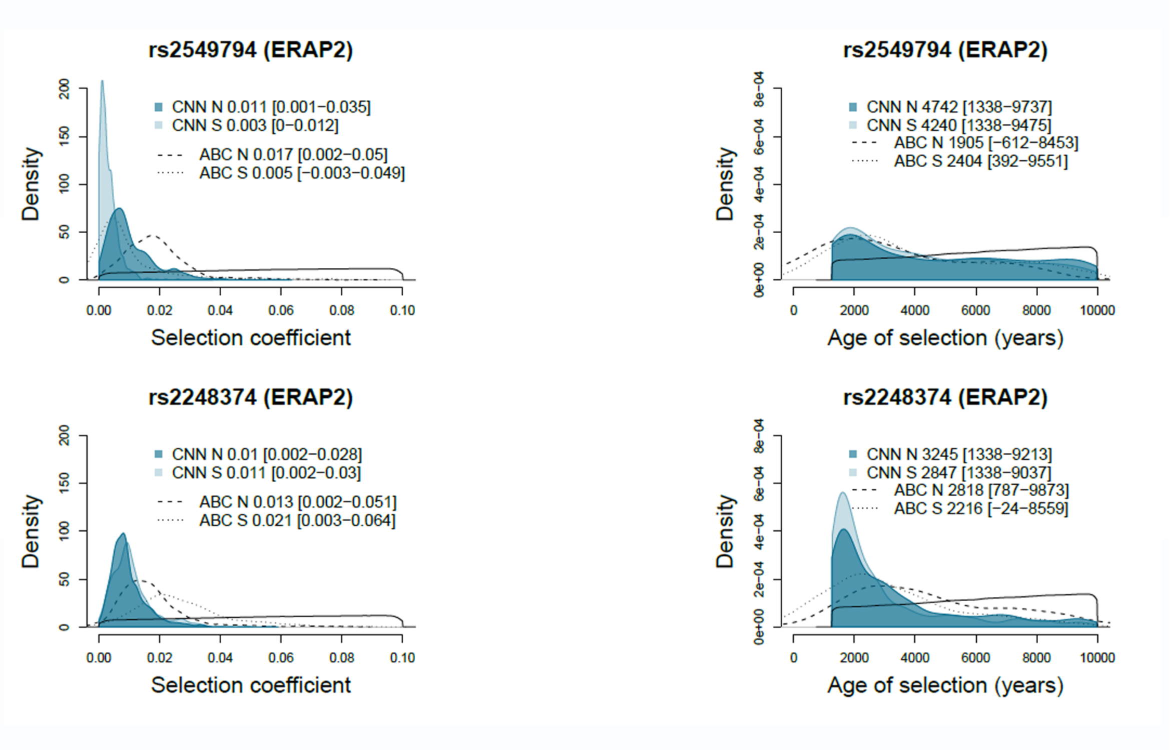


(Bottom panels) Posteriors distribution of the selection coefficient and the age of selection obtained for rs2248374 (ERAP2), a splice region SNP directly controlling the ERAP2 expression. (Top panels) Posteriors distribution obtained for rs2549794 (ERAP2) shown in Figure 4 and displayed here for comparison. See the legend of Figure 4.

**Figure S13. Robustness to training**

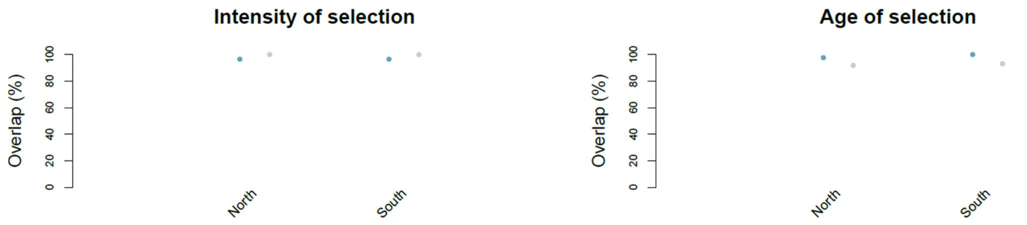


Percent of overlapping estimations across the 89 positively selected variants. We re-estimated the parameters using a new model training with an extended uniform prior of *T* up to 30,000ya. The estimates are overlapping when the initial estimates are contained in the CIs of the new estimations. Blue and grey points correspond to the CNN and ABC predictions.

**Figure S14. Predictions of the strength and timing of positive selection targeting rs10765770 (GRM5, associated with pigmentation)**

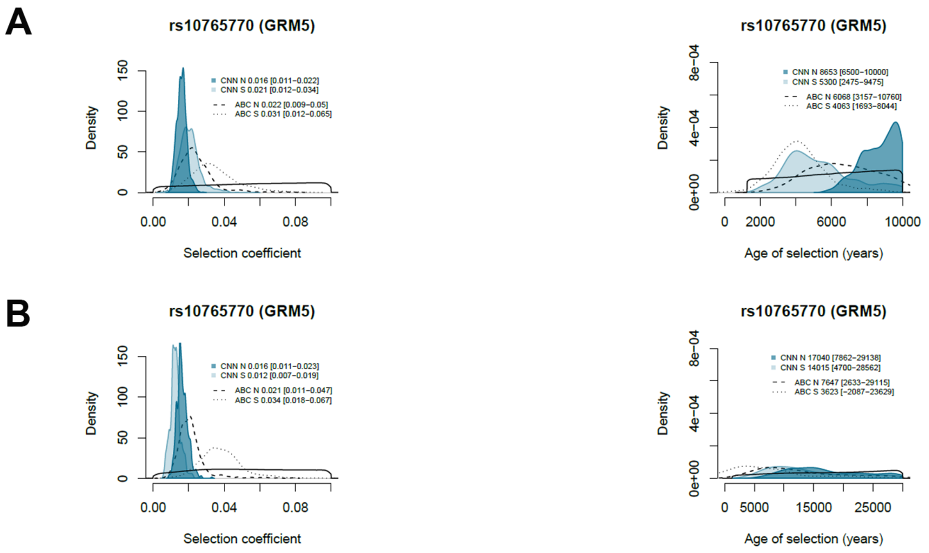


Posteriors distribution of the selection coefficient (left panel) and the age of selection (right panel) obtained for rs10765770 (GRM5, associated with pigmentation). The CNN1 model was trained with simulated age of selection up to 10,000ya (A) and 30,000ya (B). See the legend of Figure 4.

**Figure S15. Predictions of the strength and timing of positive selection targeting rs185146 (SLC45A2, associated with pigmentation)**

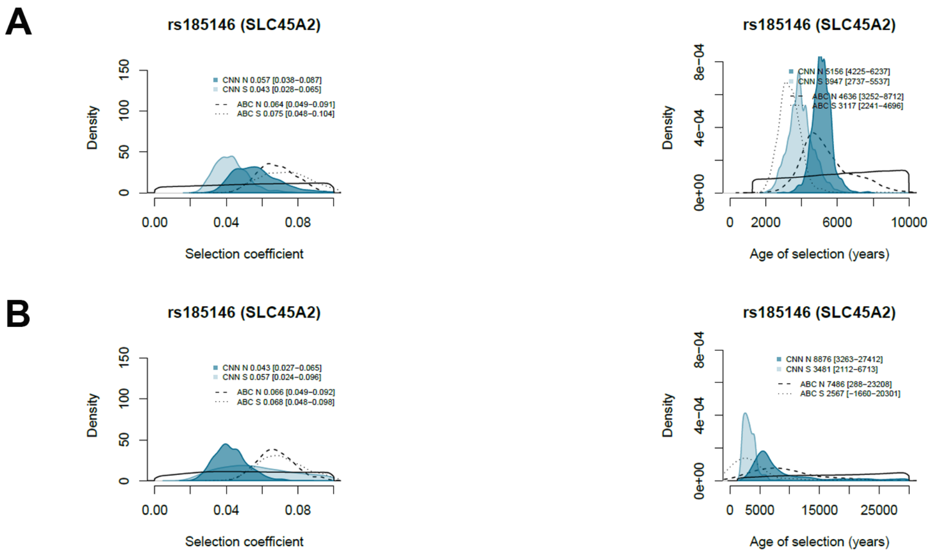


Posteriors distribution of the selection coefficient (left panel) and the age of selection (right panel) obtained for rs185146 (SLC45A2, associated with pigmentation). The CNN1 model was trained with simulated age of selection up to 10,000ya (A) and 30,000ya (B). See the legend of Figure 4.

**Figure S16. Allele frequency trajectories and genotype-based 1D images for rs185146 (SLC45A2)**

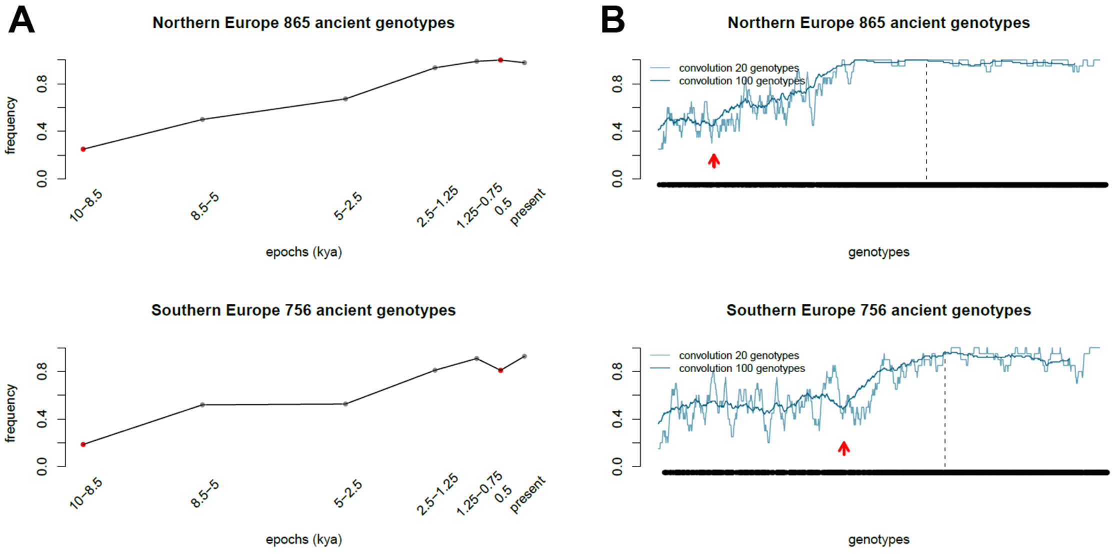


Allele frequency trajectories (A) and 1D images (B) used to perform the ABC and CNN estimations. The red arrows show a sudden increase in frequencies in genotype-based 1D images (B). Note that the different locations of these two arrows does not necessarily reflect difference in age of selection but are also explained by the distributions of radiocarbon ages in northern and southern Europe. See the legend of Figure 1.

**Figure S17. Estimation accuracy of the strength and timing of negative selection**

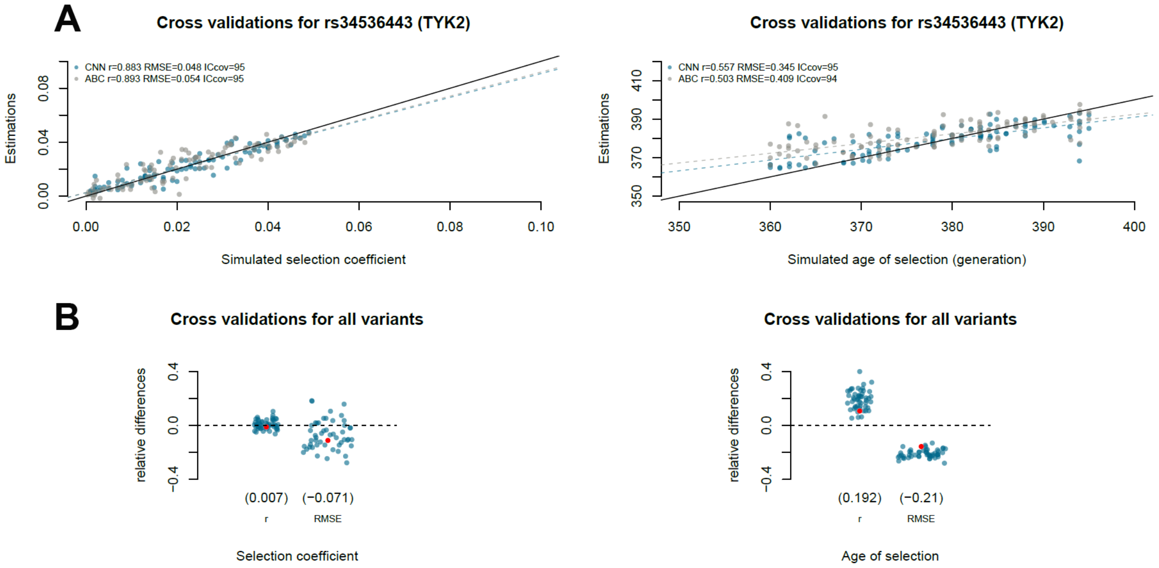


Simulation-based cross validations for the selection coefficient (left panel) and the age of selection (right panel) performed for the tuberculosis risk variant rs2549794 (TYK2) (A) and for each of the 50 negatively selected variants investigated in this study (B). See the legend of Figure 2.

**Table S1. Accuracy indices and predictions for the 89 positively selected variants**

**Table S2. Accuracy indices and predictions for the 89 positively selected variants in northern Europe**

**Table S3. Accuracy indices and predictions for the 89 positively selected variants in southern Europe**

**Table S4. Accuracy indices and predictions for the 50 negatively selected variants**

Note: In ach file the overlap of point estimates are given by three fields overlap_s, overlap_t and overlap_BA. overlap_s = 1 and overlap_t = 1 when the point estimate of one method is contained in the CI of the other for *s* and *T* respectively, overlap_s = 0 and overlap_t = 0 otherwise. overlap_BA = 1 when the two point estimates of *T* are both younger or older than 4,500 ya, overlap_BA = 0 otherwise. The average by fields (x100) give the percent of overlap in each case.

Tables S1-4 are supplied as Excel files

**File S1. 1D images for the 89 positively selected variants**

**File S2. 1D images for the 89 positively selected variants in northern Europe**

**File S3. 1D images for the 89 positively selected variants in southern Europe**

**File S4. 1D images for the positively selected variants associated to Black Death pandemic (ERAP2)**

**File S5. 1D images for the 50 negatively selected variants**

Files S1-5 are supplied as pdf files
