## Supplementary material for "Deep estimation of the intensity and timing of selection from ancient genomes": 1D images for the 89 positively selected variants

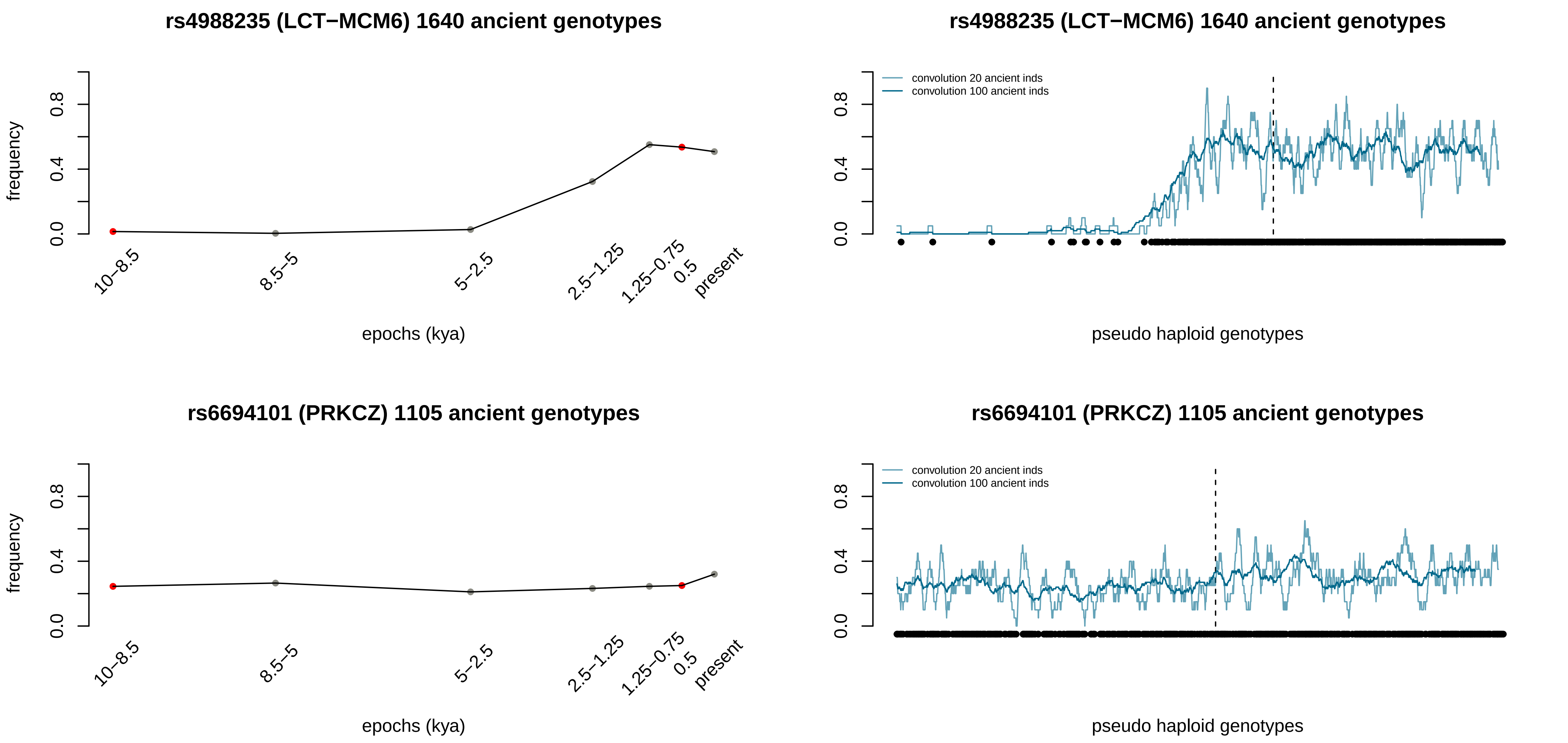
rs4988235 (LCT−MCM6) 1640 ancient genotypes
frequency
0.0
0.4
0.8
10–8.5
8.5–5
5–2.5
2.5–1.25
1.25–0.75
0.5
present
epochs (kya)
rs4988235 (LCT−MCM6) 1640 ancient genotypes
convolution 20 ancient inds
convolution 100 ancient inds
pseudo haploid genotypes
rs6694101 (PRKCZ) 1105 ancient genotypes
frequency
epochs (kya)
rs6694101 (PRKCZ) 1105 ancient genotypes
convolution 20 ancient inds
convolution 100 ancient inds
pseudo haploid genotypes

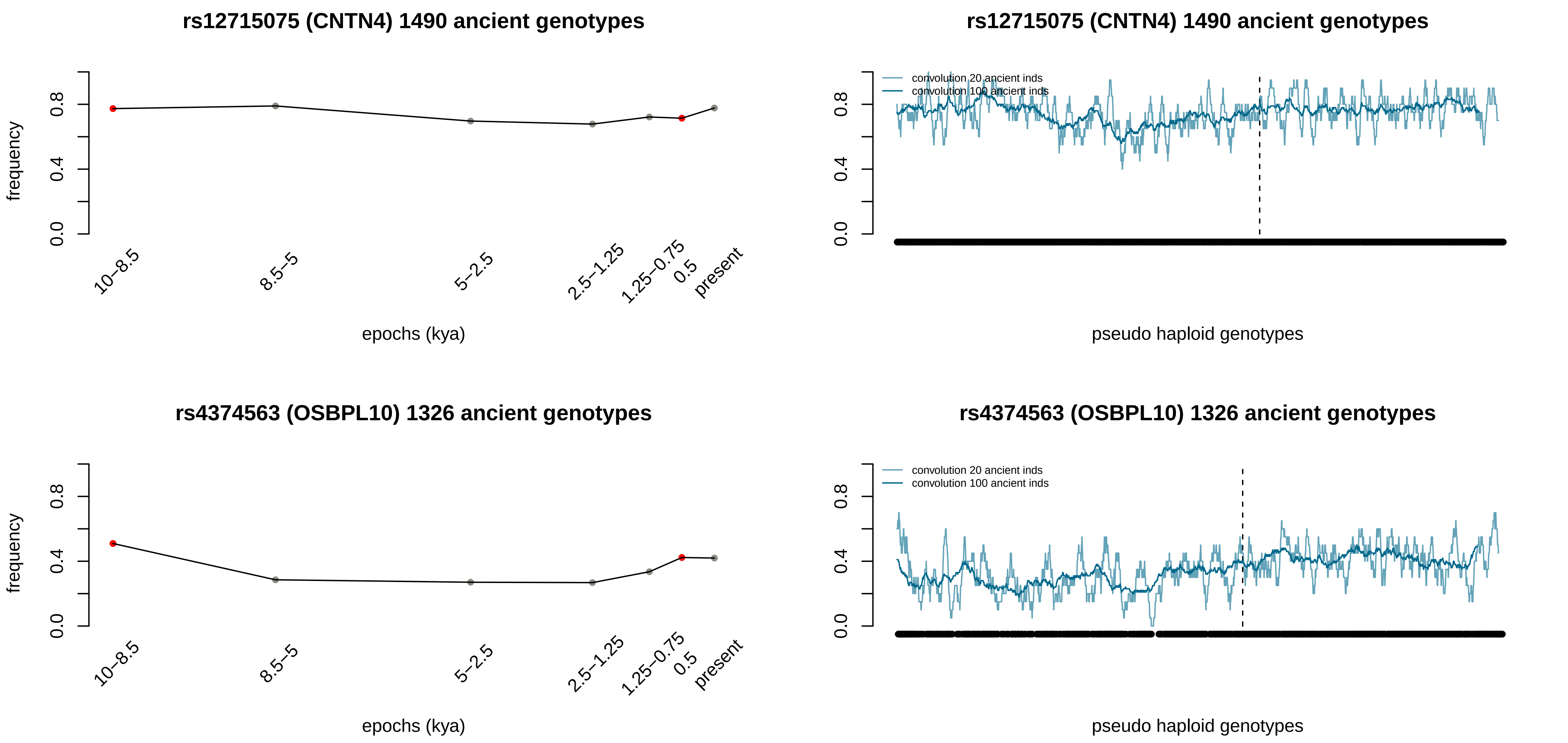
rs12715075 (CNTN4) 1490 ancient genotypes
frequency
0.0
0.4
0.8
10–8.5
8.5–5
5–2.5
2.5–1.25
1.25–0.75
0.5
present
epochs (kya)
rs12715075 (CNTN4) 1490 ancient genotypes
convolution 20 ancient inds
convolution 100 ancient inds
0.0
0.4
0.8
pseudo haploid genotypes
rs4374563 (OSBPL10) 1326 ancient genotypes
frequency
0.0
0.4
0.8
10–8.5
8.5–5
5–2.5
2.5–1.25
1.25–0.75
0.5
present
epochs (kya)
rs4374563 (OSBPL10) 1326 ancient genotypes
convolution 20 ancient inds
convolution 100 ancient inds
0.0
0.4
0.8
pseudo haploid genotypes

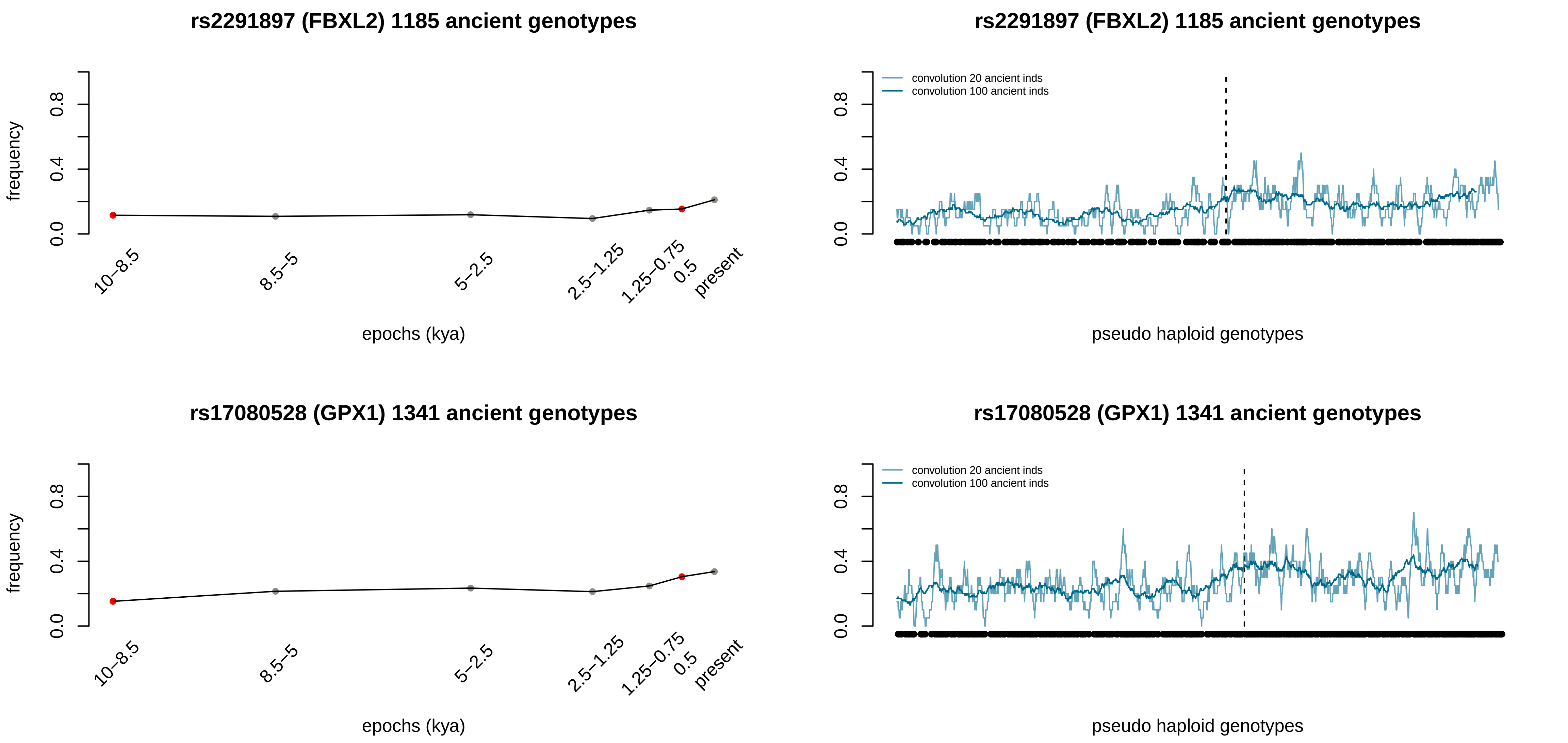

rs2291897 (FBXL2) 1185 ancient genotypes
frequency
0.0
0.4
0.8
10–8.5
8.5–5
5–2.5
2.5–1.25
1.25–0.75
0.5
present
epochs (kya)
rs2291897 (FBXL2) 1185 ancient genotypes
convolution 20 ancient inds
convolution 100 ancient inds
pseudo haploid genotypes
rs17080528 (GPX1) 1341 ancient genotypes
frequency
epochs (kya)
rs17080528 (GPX1) 1341 ancient genotypes
convolution 20 ancient inds
convolution 100 ancient inds
pseudo haploid genotypes

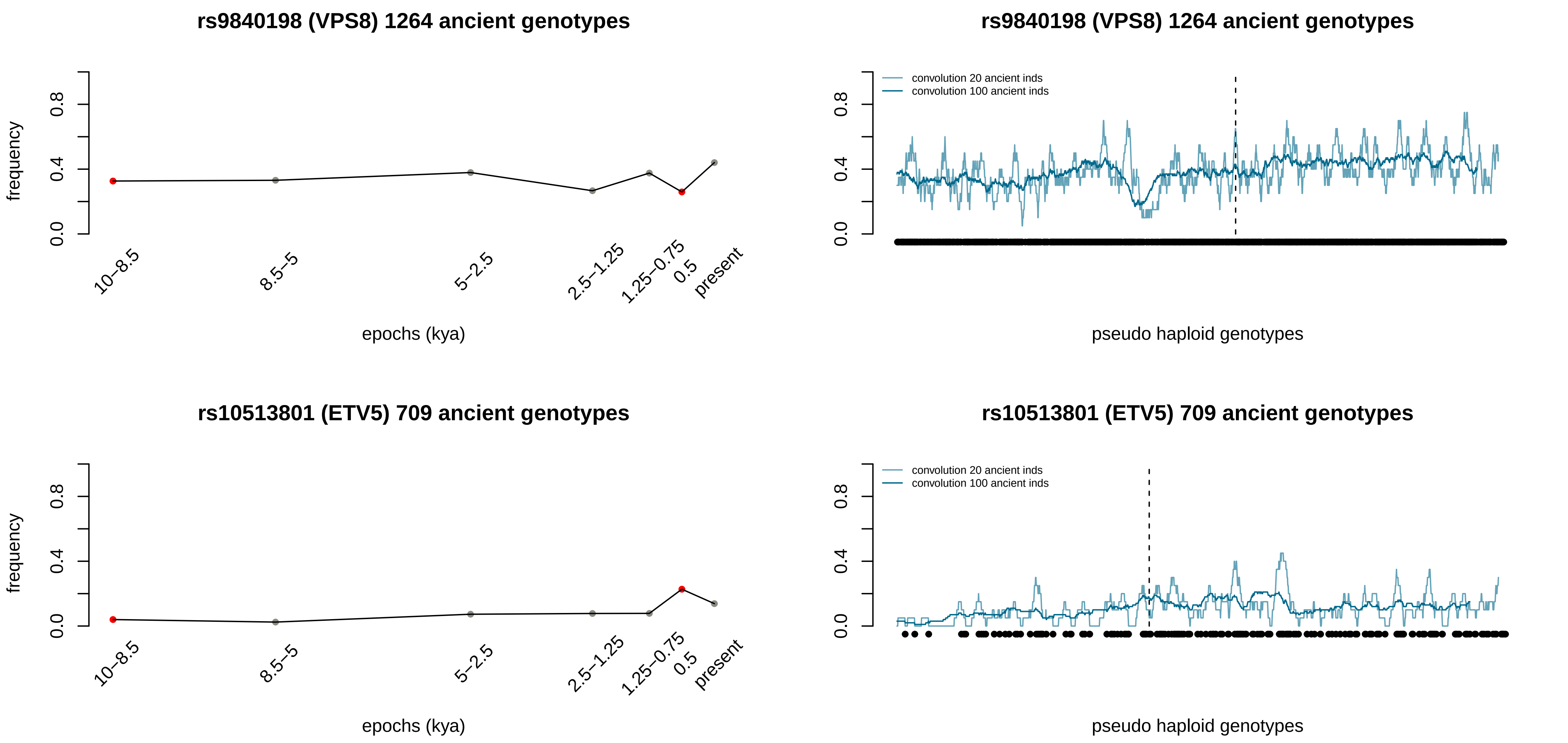

rs9840198 (VPS8) 1264 ancient genotypes
frequency
0.0
0.4
0.8
10–8.5
8.5–5
5–2.5
2.5–1.25
1.25–0.75
0.5
present
epochs (kya)
rs9840198 (VPS8) 1264 ancient genotypes
convolution 20 ancient inds
convolution 100 ancient inds
pseudo haploid genotypes
rs10513801 (ETV5) 709 ancient genotypes
frequency
epochs (kya)
rs10513801 (ETV5) 709 ancient genotypes
convolution 20 ancient inds
convolution 100 ancient inds
pseudo haploid genotypes

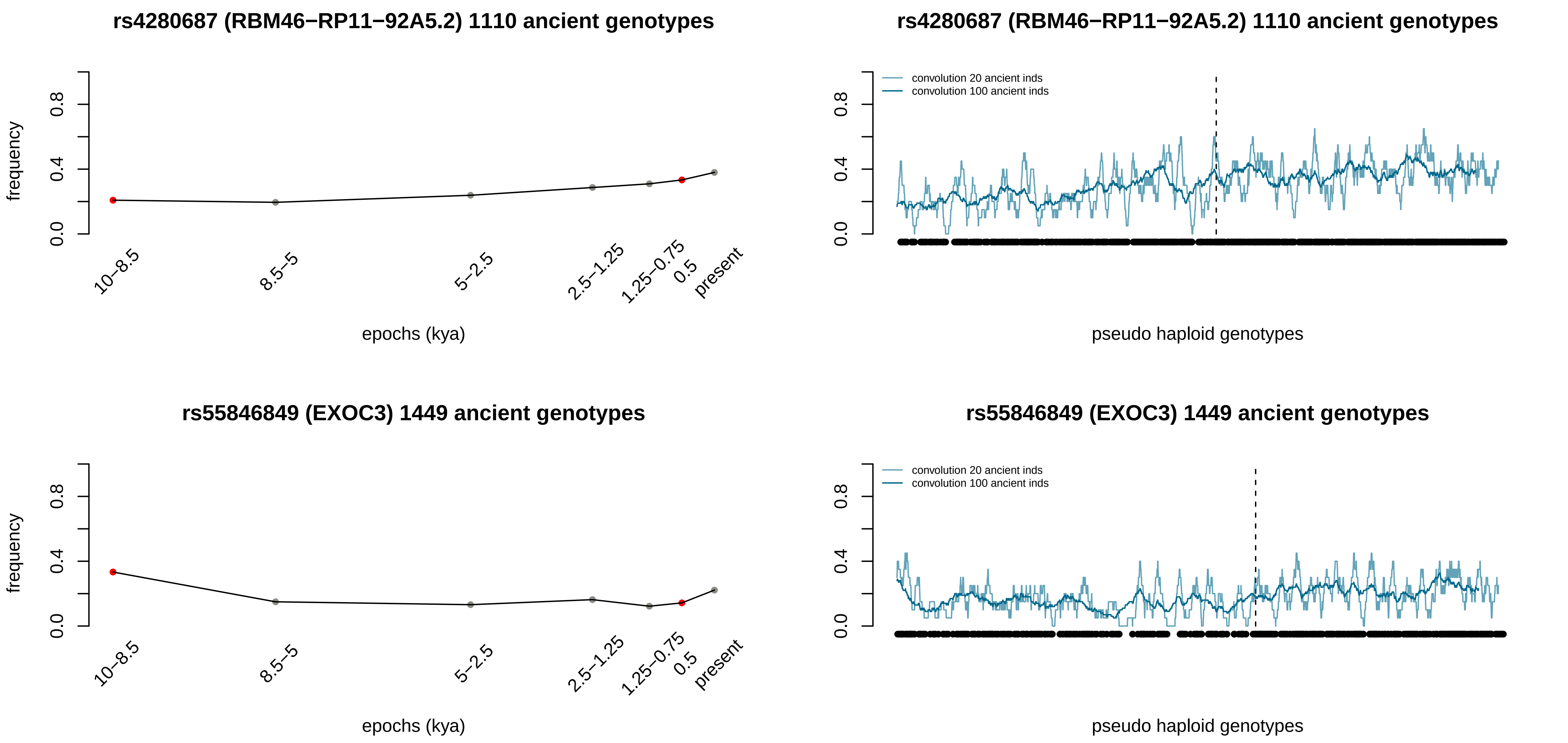
rs4280687 (RBM46−RP11−92A5.2) 1110 ancient genotypes
frequency
0.0
0.4
0.8
10−8.5
8.5−5
5−2.5
2.5−1.25
1.25−0.75
0.5
present
epochs (kya)
rs4280687 (RBM46−RP11−92A5.2) 1110 ancient genotypes
convolution 20 ancient inds
convolution 100 ancient inds
pseudo haploid genotypes
rs55846849 (EXOC3) 1449 ancient genotypes
frequency
epochs (kya)
rs55846849 (EXOC3) 1449 ancient genotypes
convolution 20 ancient inds
convolution 100 ancient inds
pseudo haploid genotypes

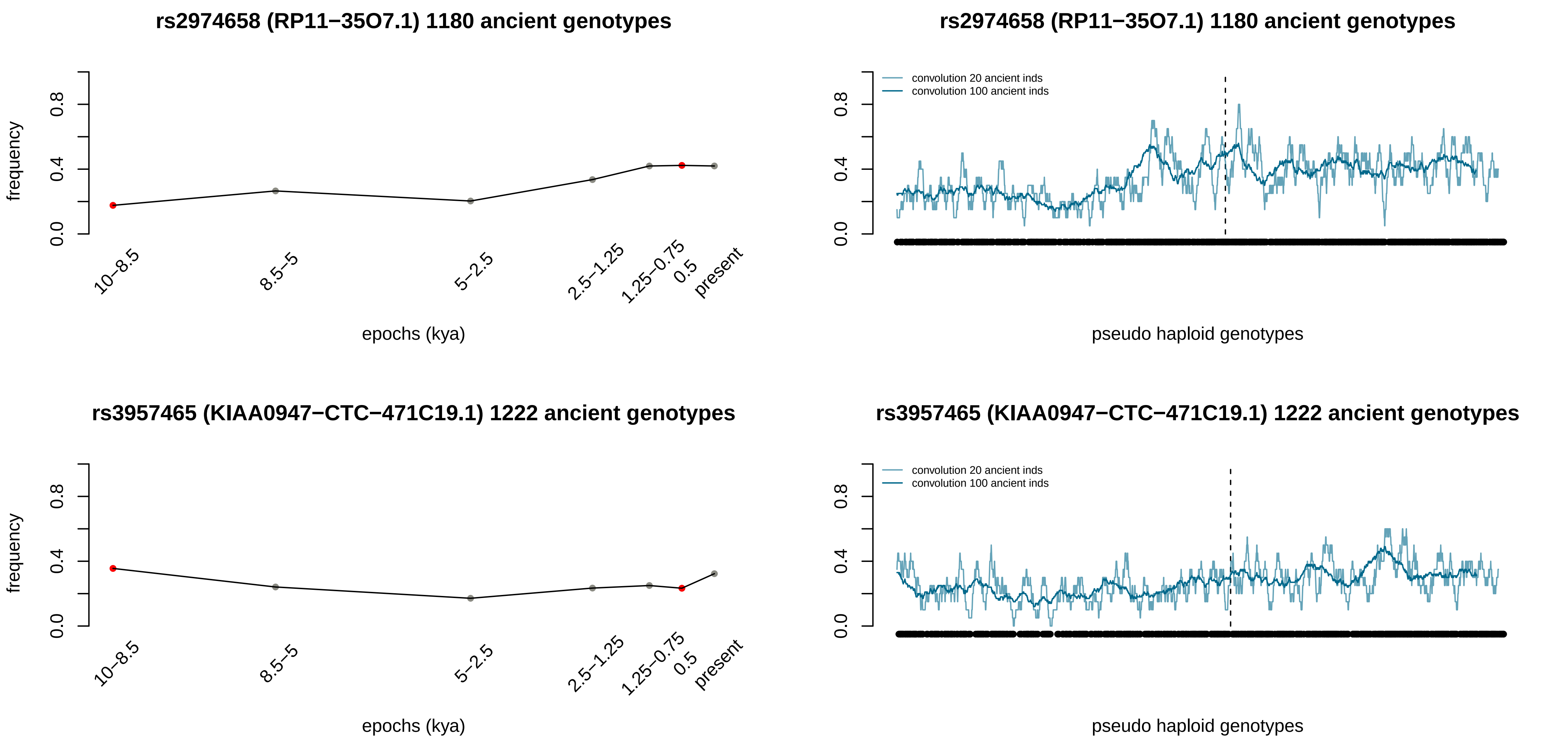
rs2974658 (RP11−35O7.1) 1180 ancient genotypes
frequency
0.0
0.4
0.8
10−8.5
8.5−5
5−2.5
2.5−1.25
1.25−0.75
0.5
present
epochs (kya)
rs2974658 (RP11−35O7.1) 1180 ancient genotypes
convolution 20 ancient inds
convolution 100 ancient inds
pseudo haploid genotypes
rs3957465 (KIAA0947−CTC−471C19.1) 1222 ancient genotypes
frequency
epochs (kya)
rs3957465 (KIAA0947−CTC−471C19.1) 1222 ancient genotypes
convolution 20 ancient inds
convolution 100 ancient inds
pseudo haploid genotypes

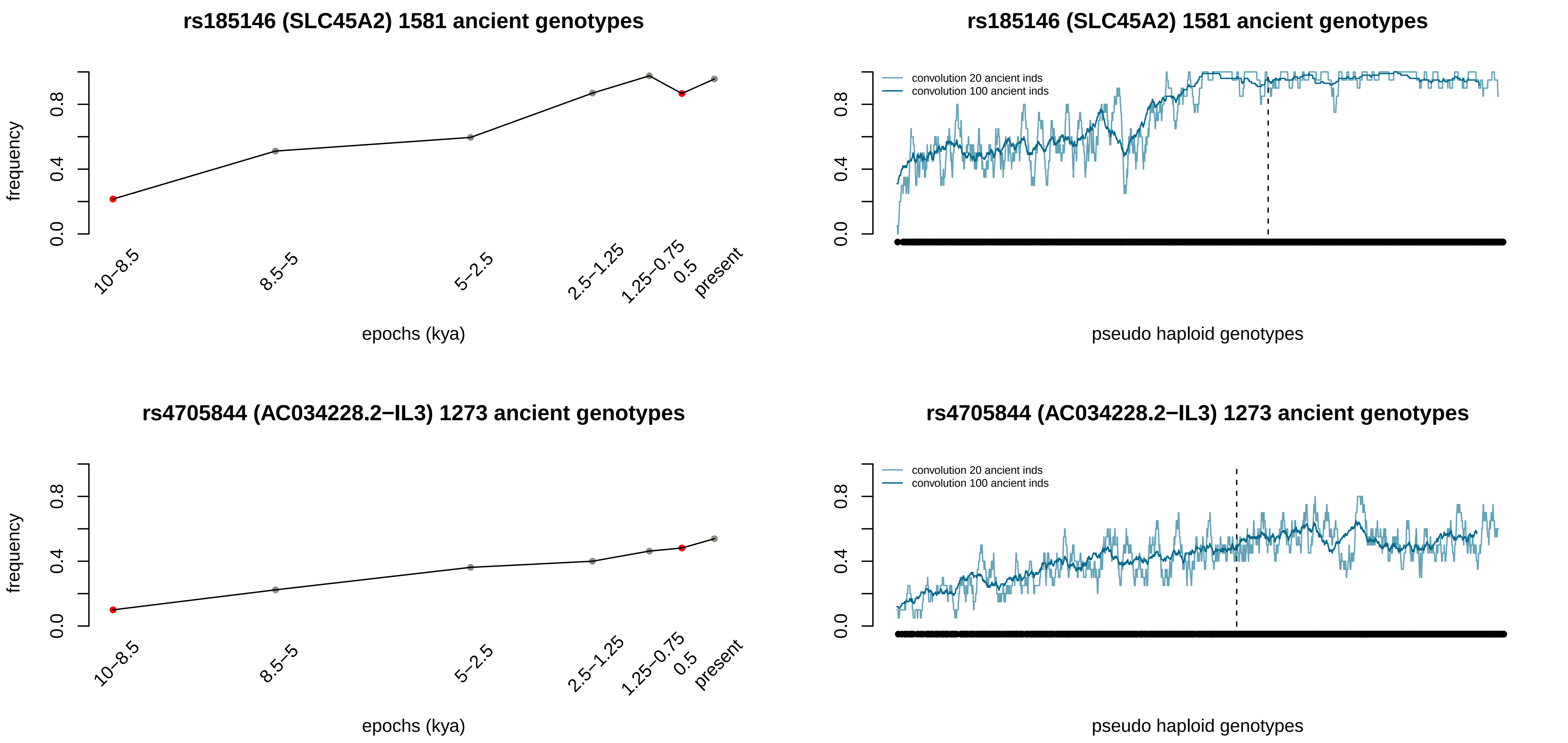
rs185146 (SLC45A2) 1581 ancient genotypes
frequency
0.0
0.4
0.8
10−8.5
8.5−5
5−2.5
2.5−1.25
1.25−0.75
0.5
present
epochs (kya)
rs185146 (SLC45A2) 1581 ancient genotypes
convolution 20 ancient inds
convolution 100 ancient inds
pseudo haploid genotypes
rs4705844 (AC034228.2−IL3) 1273 ancient genotypes
frequency
epochs (kya)
rs4705844 (AC034228.2−IL3) 1273 ancient genotypes
convolution 20 ancient inds
convolution 100 ancient inds
pseudo haploid genotypes

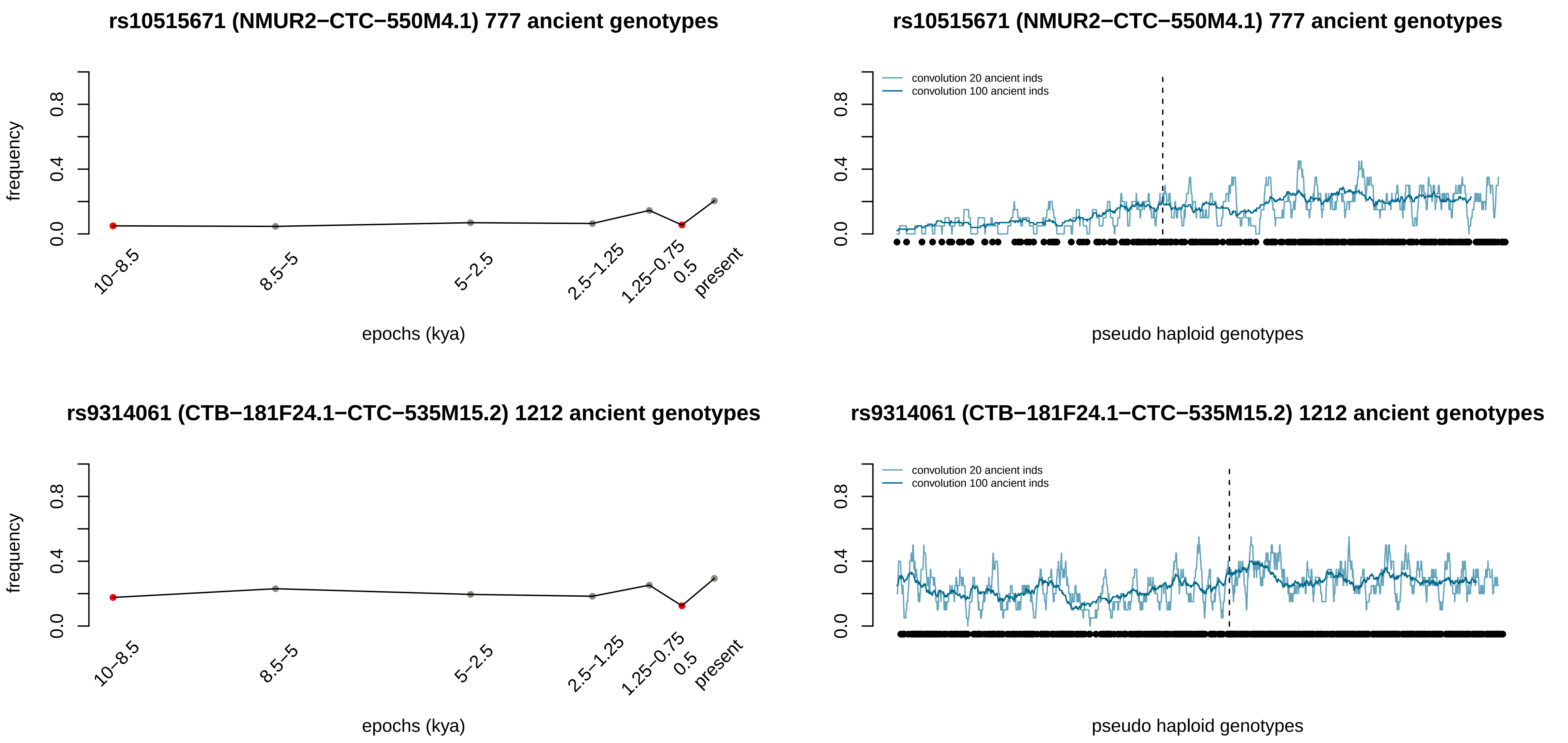
rs10515671 (NMUR2−CTC−550M4.1) 777 ancient genotypes
frequency
0.0
0.4
0.8
10−8.5
8.5−5
5−2.5
2.5−1.25
1.25−0.75
0.5
present
epochs (kya)
rs10515671 (NMUR2−CTC−550M4.1) 777 ancient genotypes
convolution 20 ancient inds
convolution 100 ancient inds
pseudo haploid genotypes
rs9314061 (CTB−181F24.1−CTC−535M15.2) 1212 ancient genotypes
frequency
epochs (kya)
rs9314061 (CTB−181F24.1−CTC−535M15.2) 1212 ancient genotypes
convolution 20 ancient inds
convolution 100 ancient inds
pseudo haploid genotypes

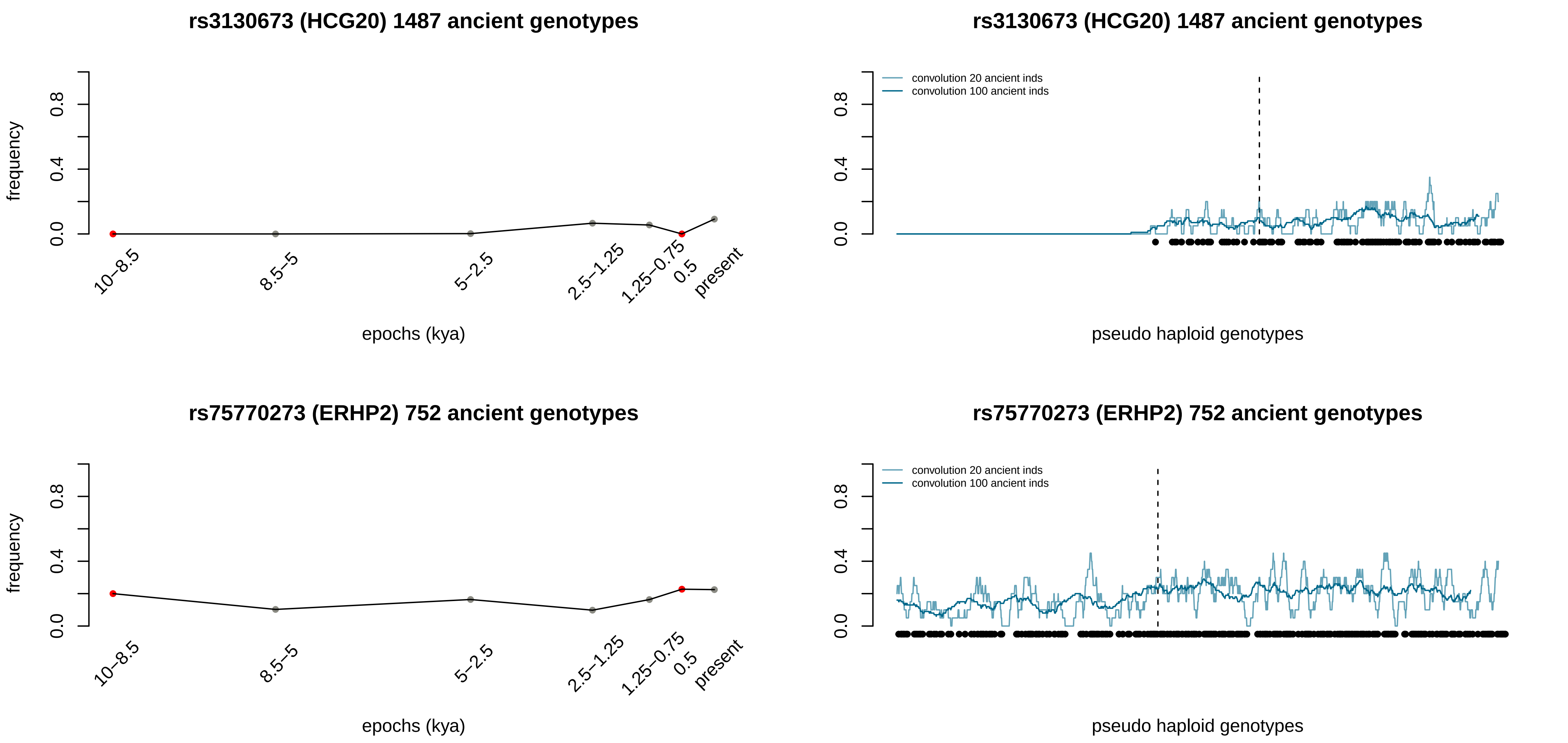
rs3130673 (HCG20) 1487 ancient genotypes
frequency
0.0
0.4
0.8
10–8.5
8.5–5
5–2.5
2.5–1.25
1.25–0.75
0.5
present
epochs (kya)
rs3130673 (HCG20) 1487 ancient genotypes
convolution 20 ancient inds
convolution 100 ancient inds
pseudo haploid genotypes
rs75770273 (ERHP2) 752 ancient genotypes
frequency
epochs (kya)
rs75770273 (ERHP2) 752 ancient genotypes
convolution 20 ancient inds
convolution 100 ancient inds
pseudo haploid genotypes

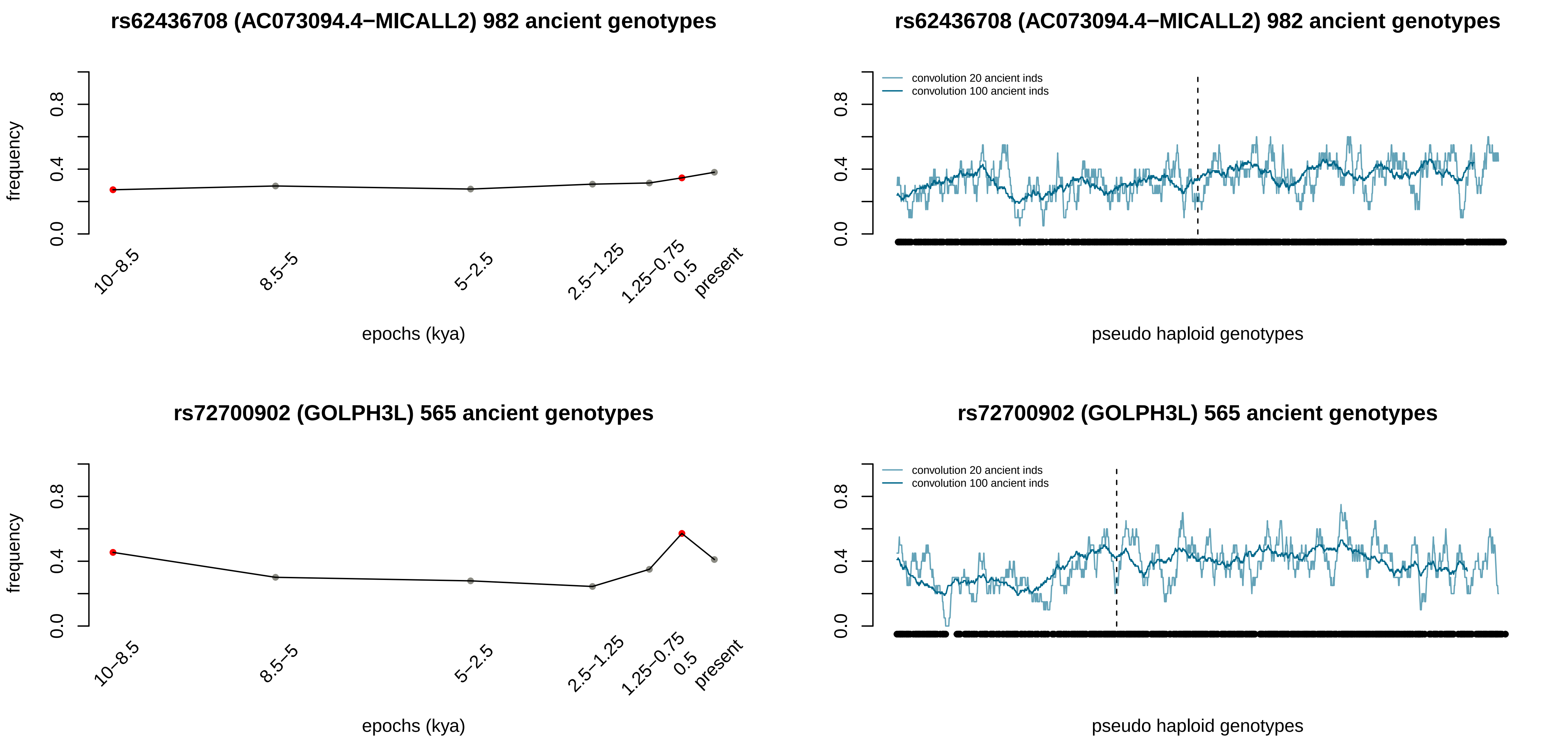
rs62436708 (AC073094.4−MICALL2) 982 ancient genotypes
frequency
0.0
0.4
0.8
10–8.5
8.5–5
5–2.5
2.5–1.25
1.25–0.75
0.5
present
epochs (kya)
rs62436708 (AC073094.4−MICALL2) 982 ancient genotypes
convolution 20 ancient inds
convolution 100 ancient inds
pseudo haploid genotypes
rs72700902 (GOLPH3L) 565 ancient genotypes
frequency
epochs (kya)
rs72700902 (GOLPH3L) 565 ancient genotypes
convolution 20 ancient inds
convolution 100 ancient inds
pseudo haploid genotypes

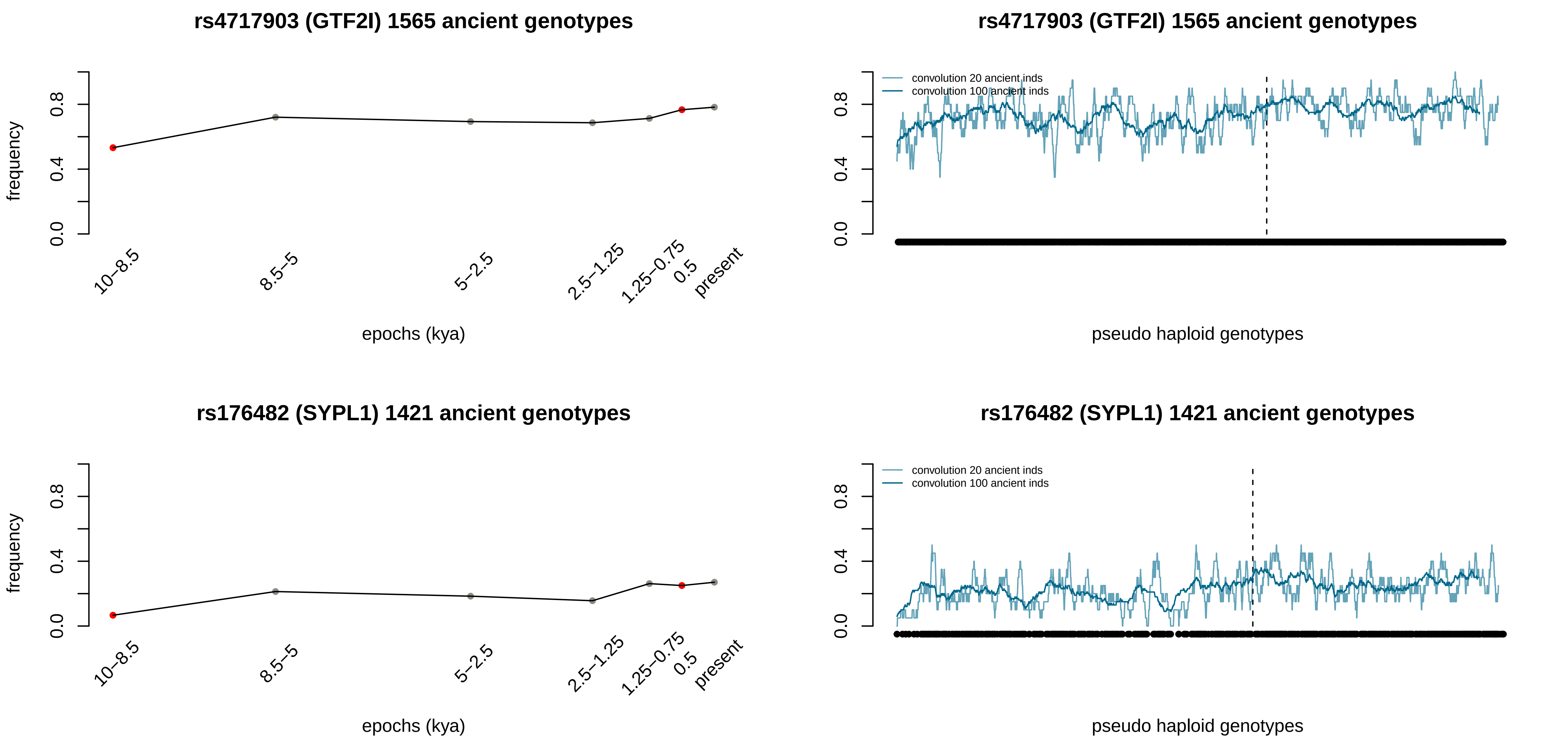

rs4717903 (GTF2I) 1565 ancient genotypes
frequency
0.0
0.4
0.8
10–8.5
8.5–5
5–2.5
2.5–1.25
1.25–0.75
0.5
present
epochs (kya)
rs4717903 (GTF2I) 1565 ancient genotypes
convolution 20 ancient inds
convolution 100 ancient inds
pseudo haploid genotypes
rs176482 (SYPL1) 1421 ancient genotypes
frequency
epochs (kya)
rs176482 (SYPL1) 1421 ancient genotypes
convolution 20 ancient inds
convolution 100 ancient inds
pseudo haploid genotypes

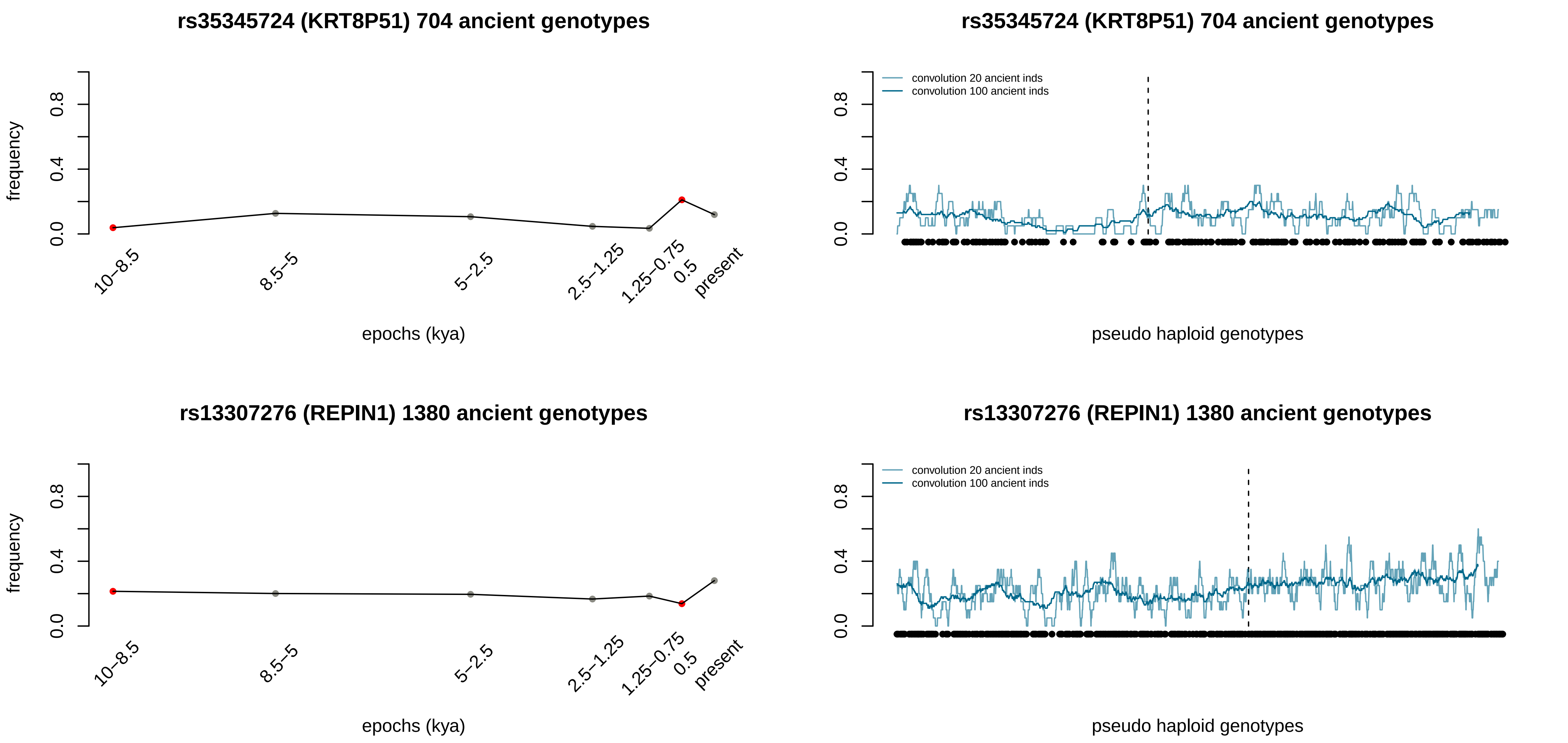

rs35345724 (KRT8P51) 704 ancient genotypes
frequency
0.0
0.4
0.8
10–8.5
8.5–5
5–2.5
2.5–1.25
1.25–0.75
0.5
present
epochs (kya)
rs35345724 (KRT8P51) 704 ancient genotypes
convolution 20 ancient inds
convolution 100 ancient inds
pseudo haploid genotypes
rs13307276 (REPIN1) 1380 ancient genotypes
frequency
epochs (kya)
rs13307276 (REPIN1) 1380 ancient genotypes
convolution 20 ancient inds
convolution 100 ancient inds
pseudo haploid genotypes

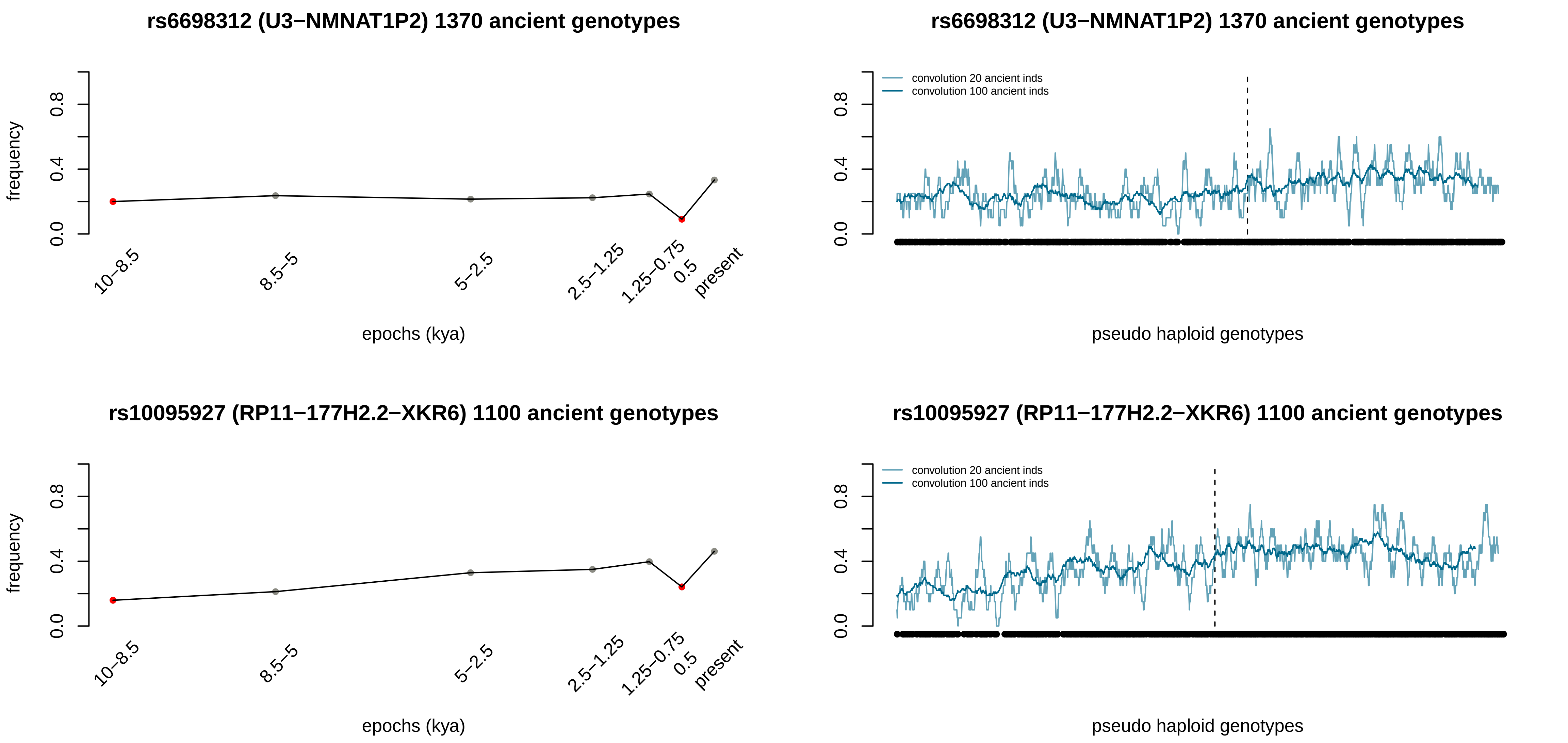
rs6698312 (U3−NMNAT1P2) 1370 ancient genotypes
frequency
epochs (kya)
10−8.5
8.5−5
5−2.5
2.5−1.25
1.25−0.75
0.5
present
convolution 20 ancient inds
convolution 100 ancient inds
pseudo haploid genotypes
rs10095927 (RP11−177H2.2−XKR6) 1100 ancient genotypes
0.0
0.4
0.8

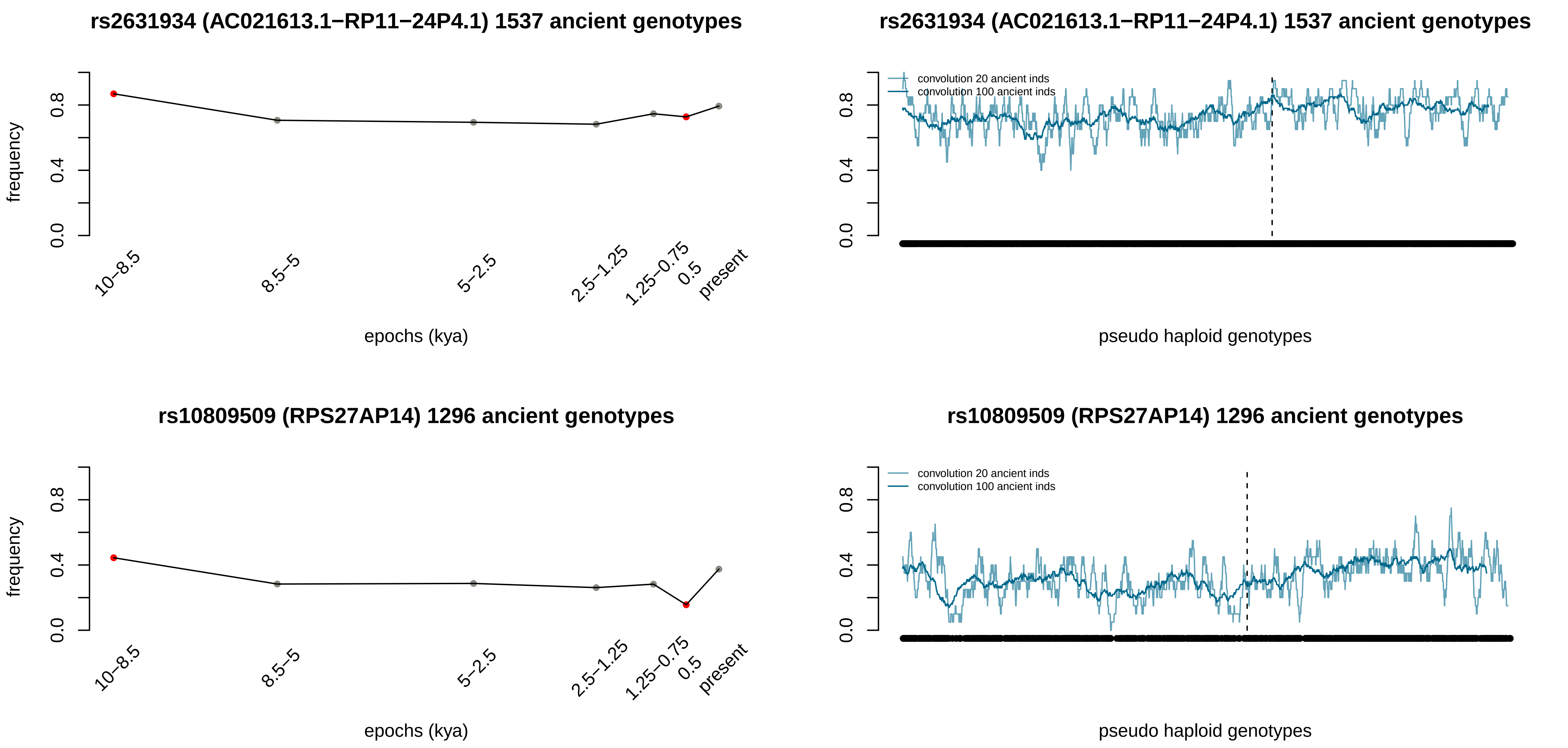

rs2631934 (AC021613.1−RP11−24P4.1) 1537 ancient genotypes
frequency
0.0
0.4
0.8
10−8.5
8.5−5
5−2.5
2.5−1.25
1.25−0.75
0.5
present
epochs (kya)
rs2631934 (AC021613.1−RP11−24P4.1) 1537 ancient genotypes
convolution 20 ancient inds
convolution 100 ancient inds
pseudo haploid genotypes
rs10809509 (RPS27AP14) 1296 ancient genotypes
frequency
epochs (kya)
rs10809509 (RPS27AP14) 1296 ancient genotypes
convolution 20 ancient inds
convolution 100 ancient inds
pseudo haploid genotypes

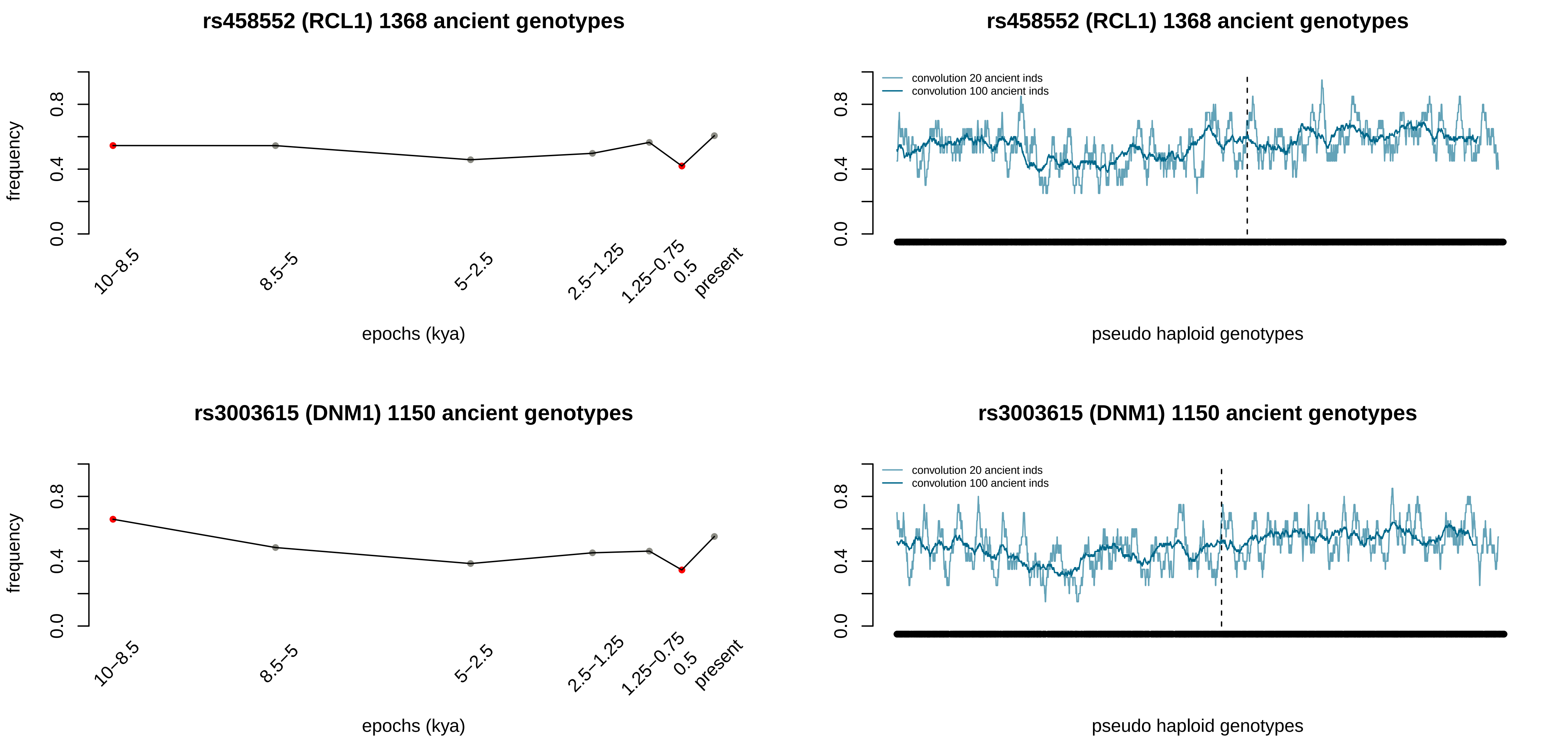

rs458552 (RCL1) 1368 ancient genotypes
frequency
0.0
0.4
0.8
10–8.5
8.5–5
5–2.5
2.5–1.25
1.25–0.75
0.5
present
epochs (kya)
rs458552 (RCL1) 1368 ancient genotypes
convolution 20 ancient inds
convolution 100 ancient inds
pseudo haploid genotypes
rs3003615 (DNM1) 1150 ancient genotypes
frequency
epochs (kya)
rs3003615 (DNM1) 1150 ancient genotypes
convolution 20 ancient inds
convolution 100 ancient inds
pseudo haploid genotypes

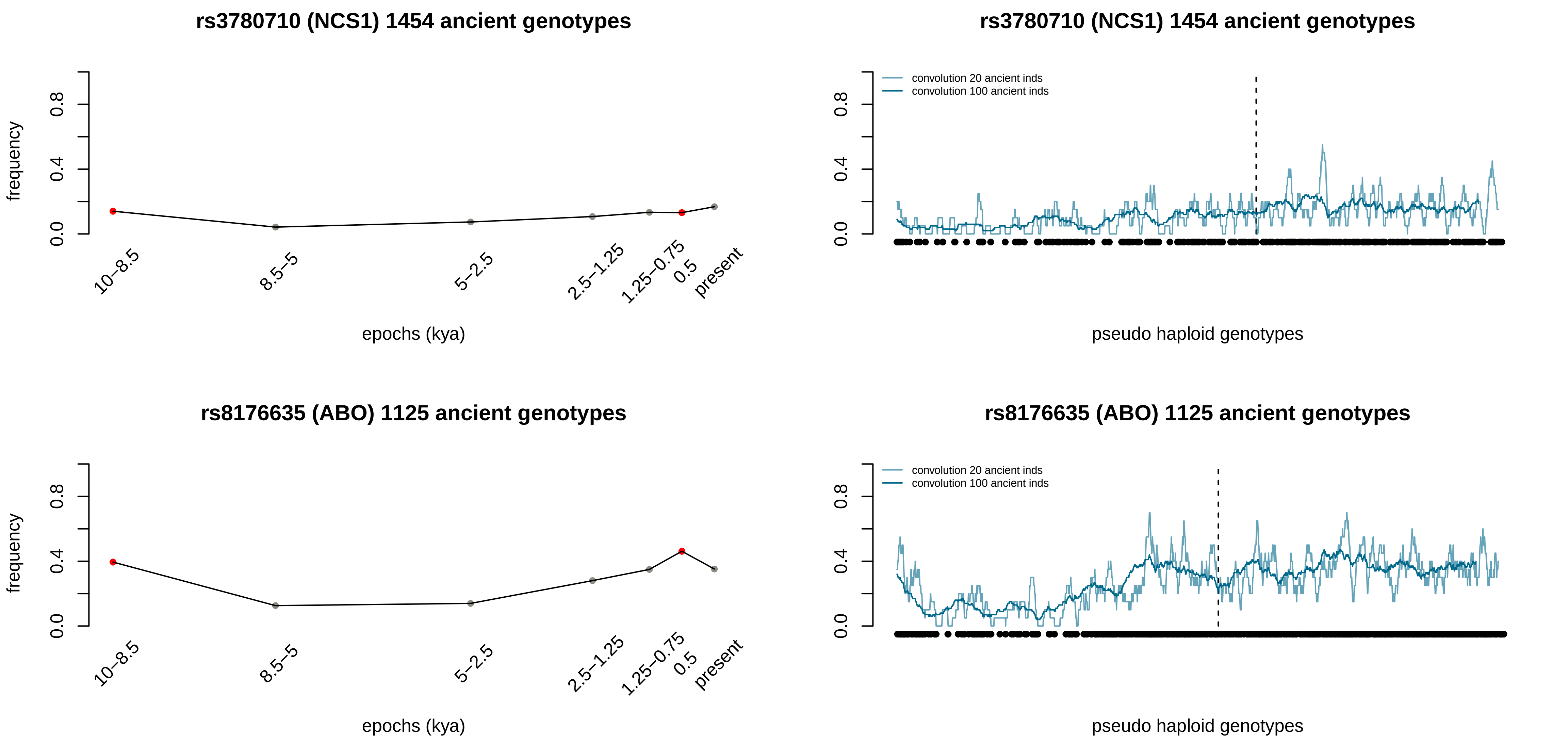
rs3780710 (NCS1) 1454 ancient genotypes
frequency
0.0
0.4
0.8
epochs (kya)
10–8.5
8.5–5
5–2.5
2.5–1.25
1.25–0.75
0.5
present
rs3780710 (NCS1) 1454 ancient genotypes
convolution 20 ancient inds
convolution 100 ancient inds
pseudo haploid genotypes
rs8176635 (ABO) 1125 ancient genotypes
frequency
epochs (kya)
rs8176635 (ABO) 1125 ancient genotypes
convolution 20 ancient inds
convolution 100 ancient inds
pseudo haploid genotypes

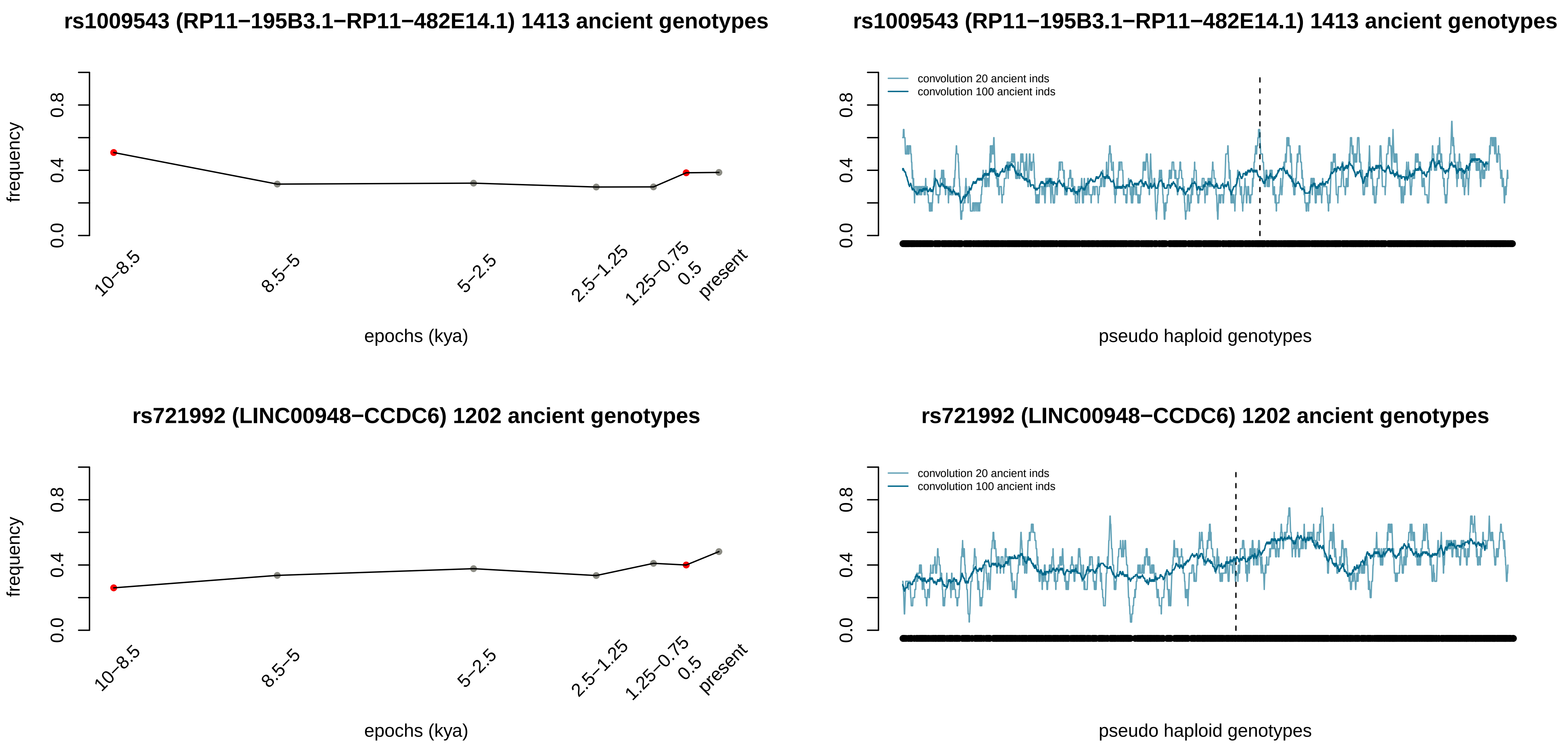
rs1009543 (RP11−195B3.1−RP11−482E14.1) 1413 ancient genotypes
frequency
0.0
0.4
0.8
10−8.5
8.5−5
5−2.5
2.5−1.25
1.25−0.75
0.5
present
epochs (kya)
rs1009543 (RP11−195B3.1−RP11−482E14.1) 1413 ancient genotypes
convolution 20 ancient inds
convolution 100 ancient inds
pseudo haploid genotypes
rs721992 (LINC00948−CCDC6) 1202 ancient genotypes
frequency
epochs (kya)
rs721992 (LINC00948−CCDC6) 1202 ancient genotypes
convolution 20 ancient inds
convolution 100 ancient inds
pseudo haploid genotypes

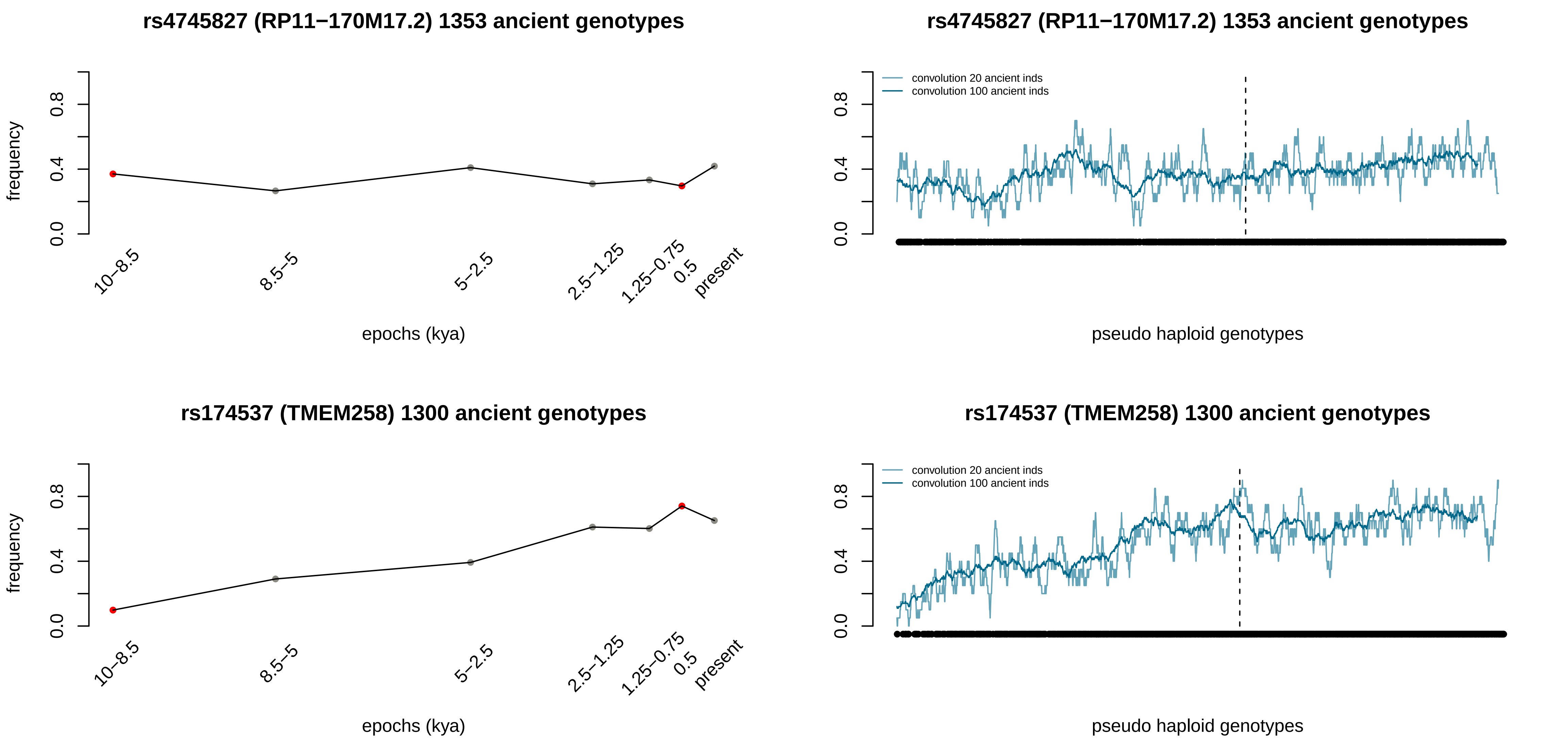
rs4745827 (RP11−170M17.2) 1353 ancient genotypes
frequency
0.0
0.4
0.8
10–8.5
8.5–5
5–2.5
2.5–1.25
1.25–0.75
0.5
present
epochs (kya)
rs4745827 (RP11−170M17.2) 1353 ancient genotypes
convolution 20 ancient inds
convolution 100 ancient inds
pseudo haploid genotypes
rs174537 (TMEM258) 1300 ancient genotypes
frequency
epochs (kya)
rs174537 (TMEM258) 1300 ancient genotypes
convolution 20 ancient inds
convolution 100 ancient inds
pseudo haploid genotypes

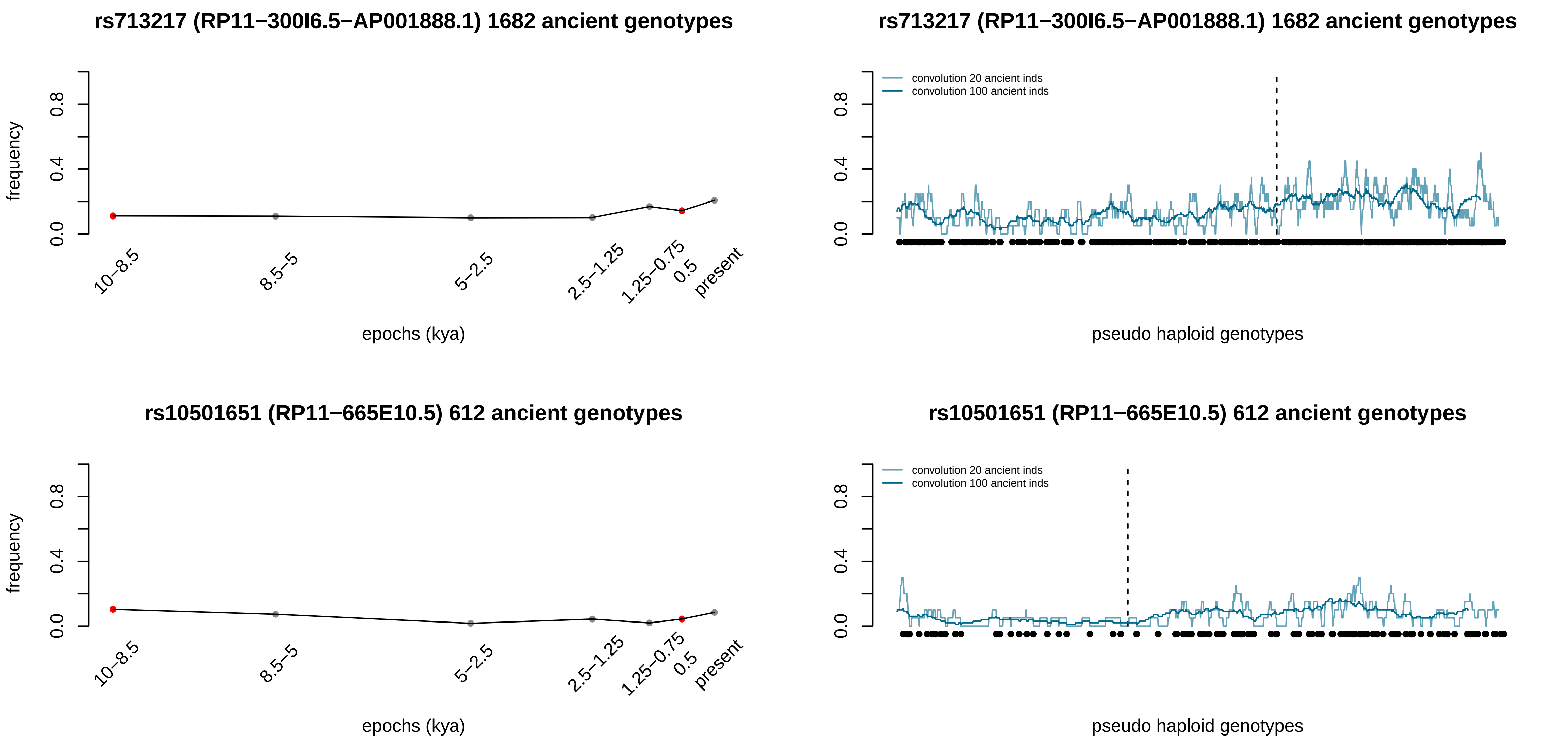
rs713217 (RP11−300I6.5−AP001888.1) 1682 ancient genotypes
frequency
epochs (kya)
10−8.5
8.5−5
5−2.5
2.5−1.25
1.25−0.75
0.5
present
convolution 20 ancient inds
convolution 100 ancient inds
pseudo haploid genotypes
rs10501651 (RP11−665E10.5) 612 ancient genotypes

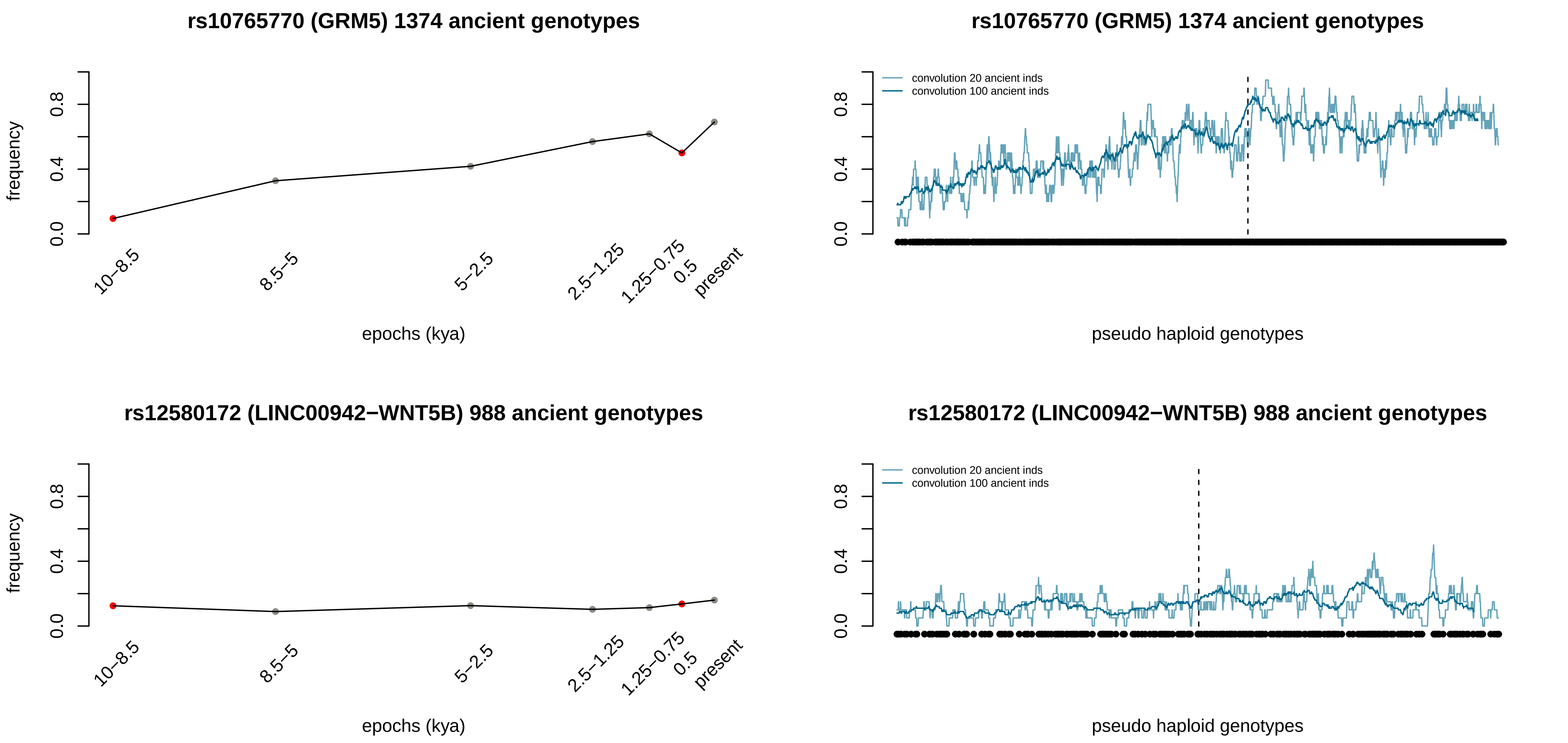
rs10765770 (GRM5) 1374 ancient genotypes
frequency
0.0
0.4
0.8
10−8.5
8.5−5
5−2.5
2.5−1.25
1.25−0.75
0.5
present
epochs (kya)
rs10765770 (GRM5) 1374 ancient genotypes
convolution 20 ancient inds
convolution 100 ancient inds
pseudo haploid genotypes
rs12580172 (LINC00942−WNT5B) 988 ancient genotypes
frequency
epochs (kya)
rs12580172 (LINC00942−WNT5B) 988 ancient genotypes
convolution 20 ancient inds
convolution 100 ancient inds
pseudo haploid genotypes

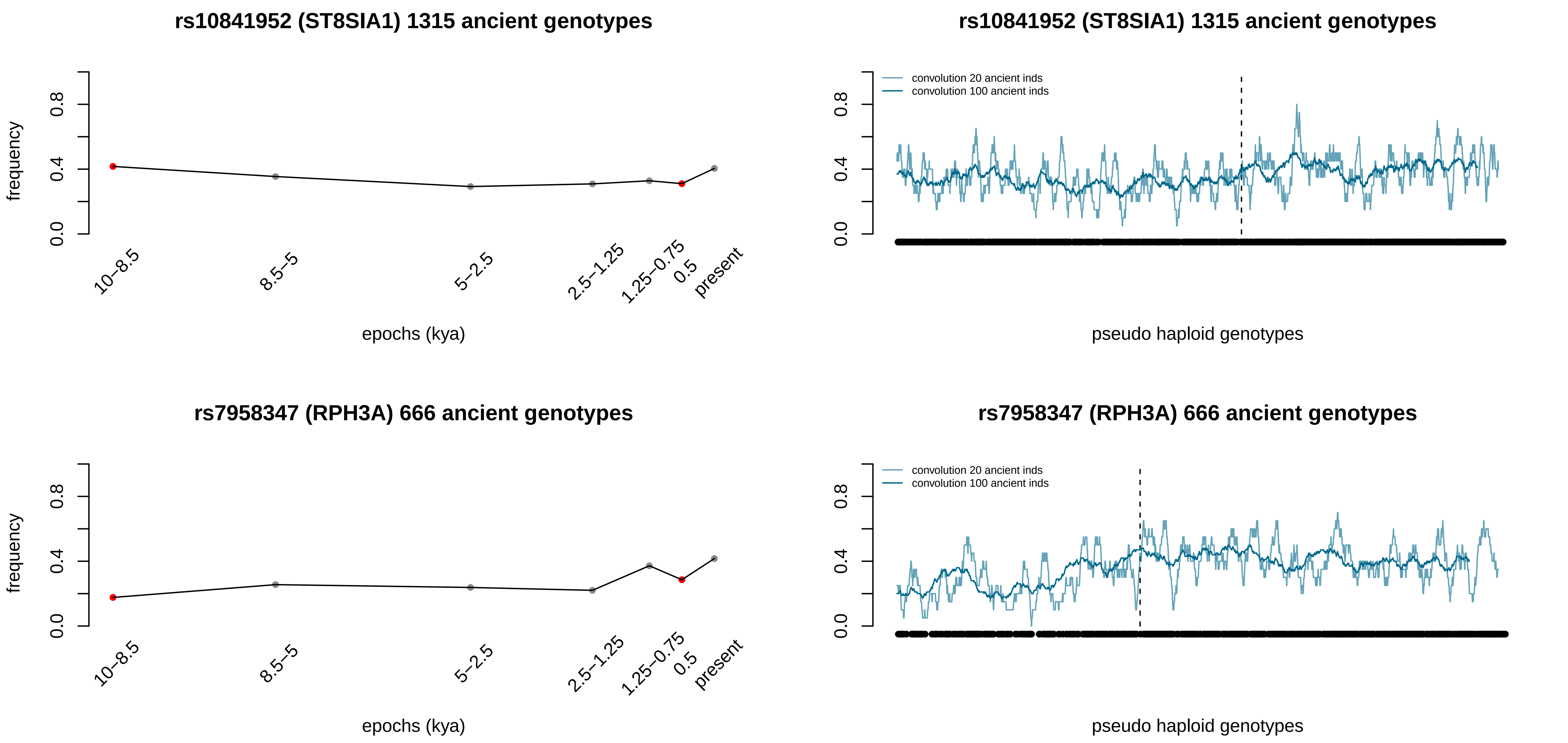
rs10841952 (ST8SIA1) 1315 ancient genotypes
frequency
0.0
0.4
0.8
10–8.5
8.5–5
5–2.5
2.5–1.25
1.25–0.75
0.5
present
epochs (kya)
rs10841952 (ST8SIA1) 1315 ancient genotypes
convolution 20 ancient inds
convolution 100 ancient inds
pseudo haploid genotypes
rs7958347 (RPH3A) 666 ancient genotypes
frequency
epochs (kya)
rs7958347 (RPH3A) 666 ancient genotypes
convolution 20 ancient inds
convolution 100 ancient inds
pseudo haploid genotypes

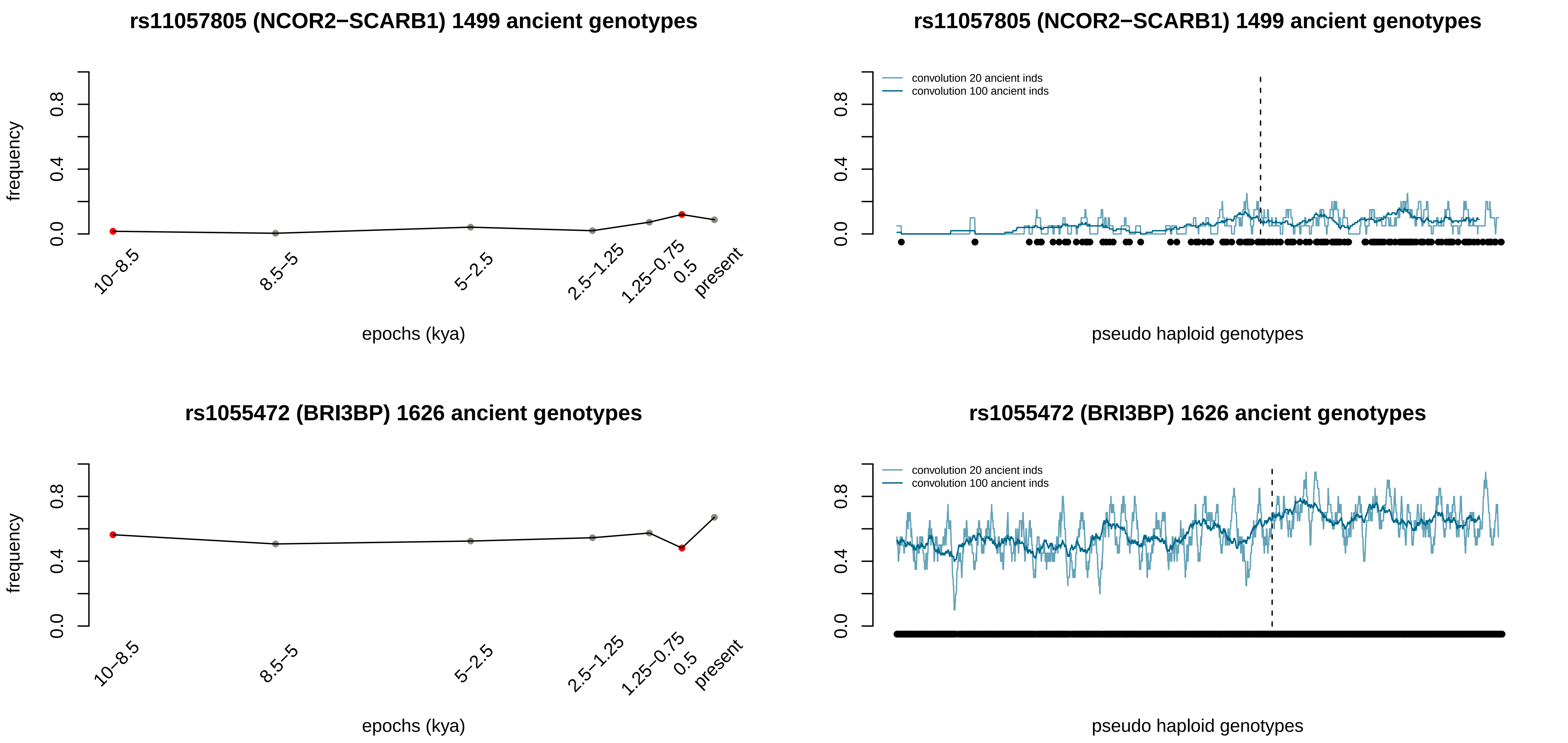

rs11057805 (NCOR2−SCARB1) 1499 ancient genotypes
frequency
0.0
0.4
0.8
10–8.5
8.5–5
5–2.5
2.5–1.25
1.25–0.75
0.5
present
epochs (kya)
rs11057805 (NCOR2−SCARB1) 1499 ancient genotypes
convolution 20 ancient inds
convolution 100 ancient inds
pseudo haploid genotypes
rs1055472 (BRI3BP) 1626 ancient genotypes
frequency
epochs (kya)
rs1055472 (BRI3BP) 1626 ancient genotypes
convolution 20 ancient inds
convolution 100 ancient inds
pseudo haploid genotypes

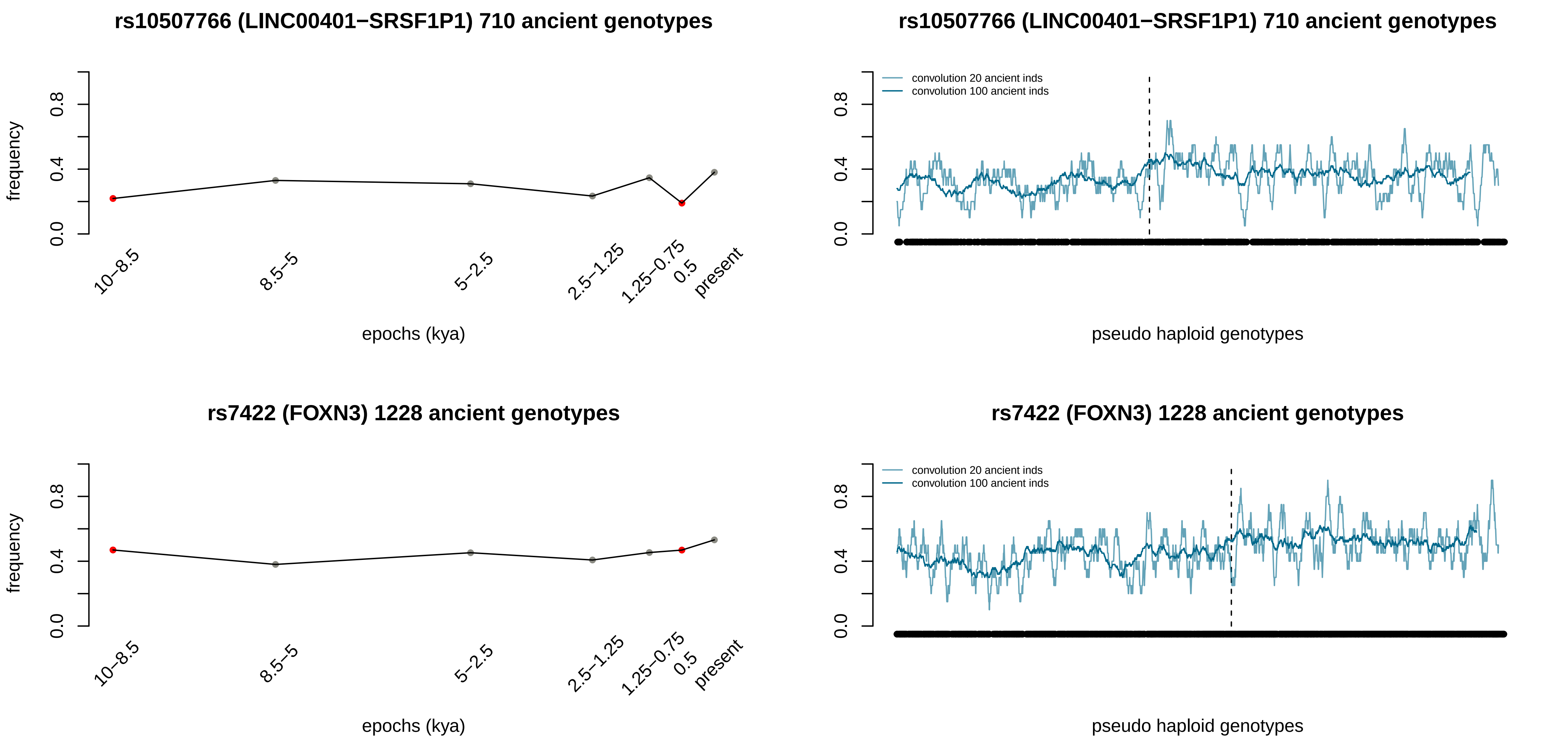
rs10507766 (LINC00401−SRSF1P1) 710 ancient genotypes
frequency
0.0
0.4
0.8
10−8.5
8.5−5
5−2.5
2.5−1.25
1.25−0.75
0.5
present
epochs (kya)
rs10507766 (LINC00401−SRSF1P1) 710 ancient genotypes
convolution 20 ancient inds
convolution 100 ancient inds
0.0
0.4
0.8
pseudo haploid genotypes
rs7422 (FOXN3) 1228 ancient genotypes
frequency
0.0
0.4
0.8
10−8.5
8.5−5
5−2.5
2.5−1.25
1.25−0.75
0.5
present
epochs (kya)
rs7422 (FOXN3) 1228 ancient genotypes
convolution 20 ancient inds
convolution 100 ancient inds
0.0
0.4
0.8
pseudo haploid genotypes

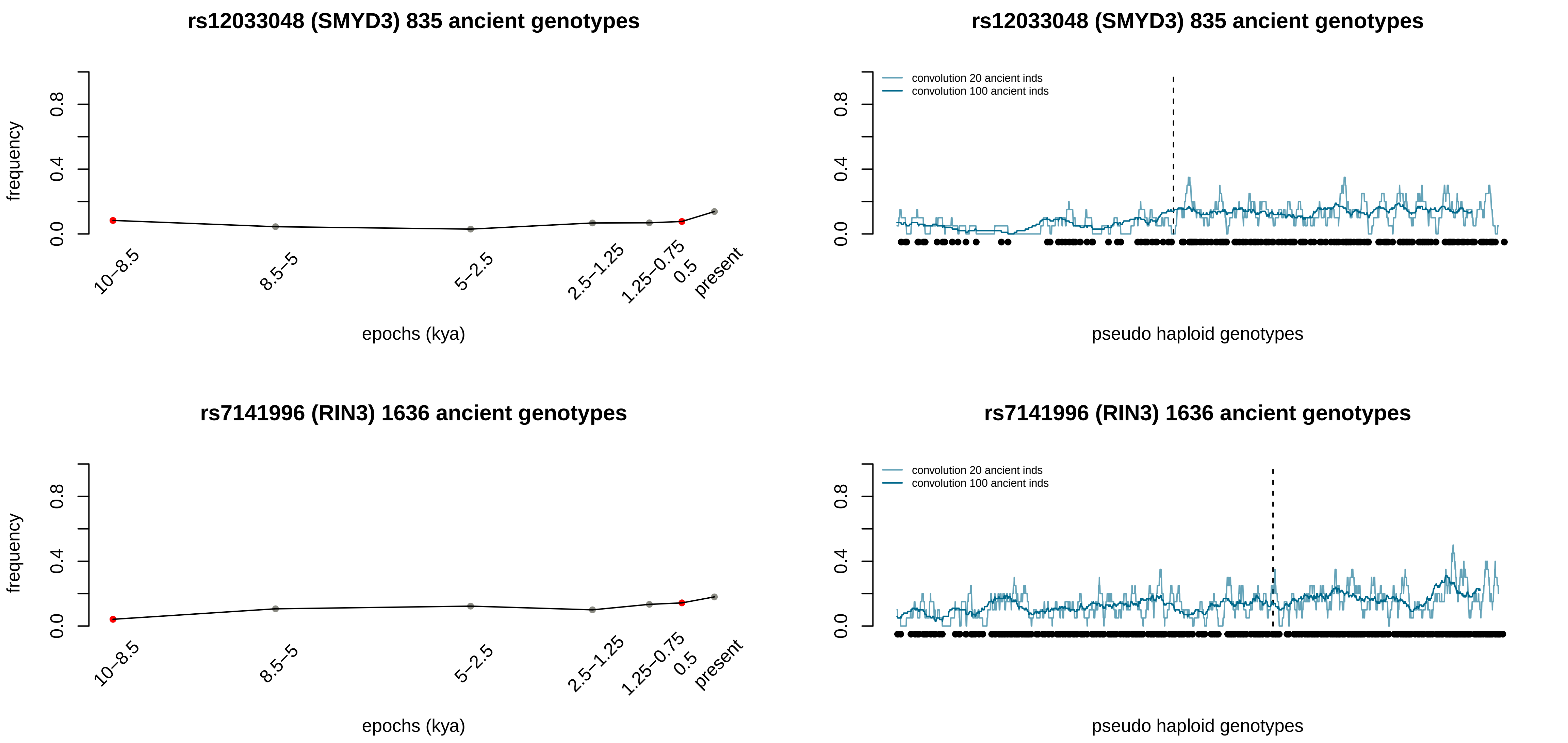
rs12033048 (SMYD3) 835 ancient genotypes
frequency
0.0
0.4
0.8
10–8.5
8.5–5
5–2.5
2.5–1.25
1.25–0.75
0.5
present
epochs (kya)
rs12033048 (SMYD3) 835 ancient genotypes
convolution 20 ancient inds
convolution 100 ancient inds
pseudo haploid genotypes
rs7141996 (RIN3) 1636 ancient genotypes
frequency
epochs (kya)
rs7141996 (RIN3) 1636 ancient genotypes
convolution 20 ancient inds
convolution 100 ancient inds
pseudo haploid genotypes

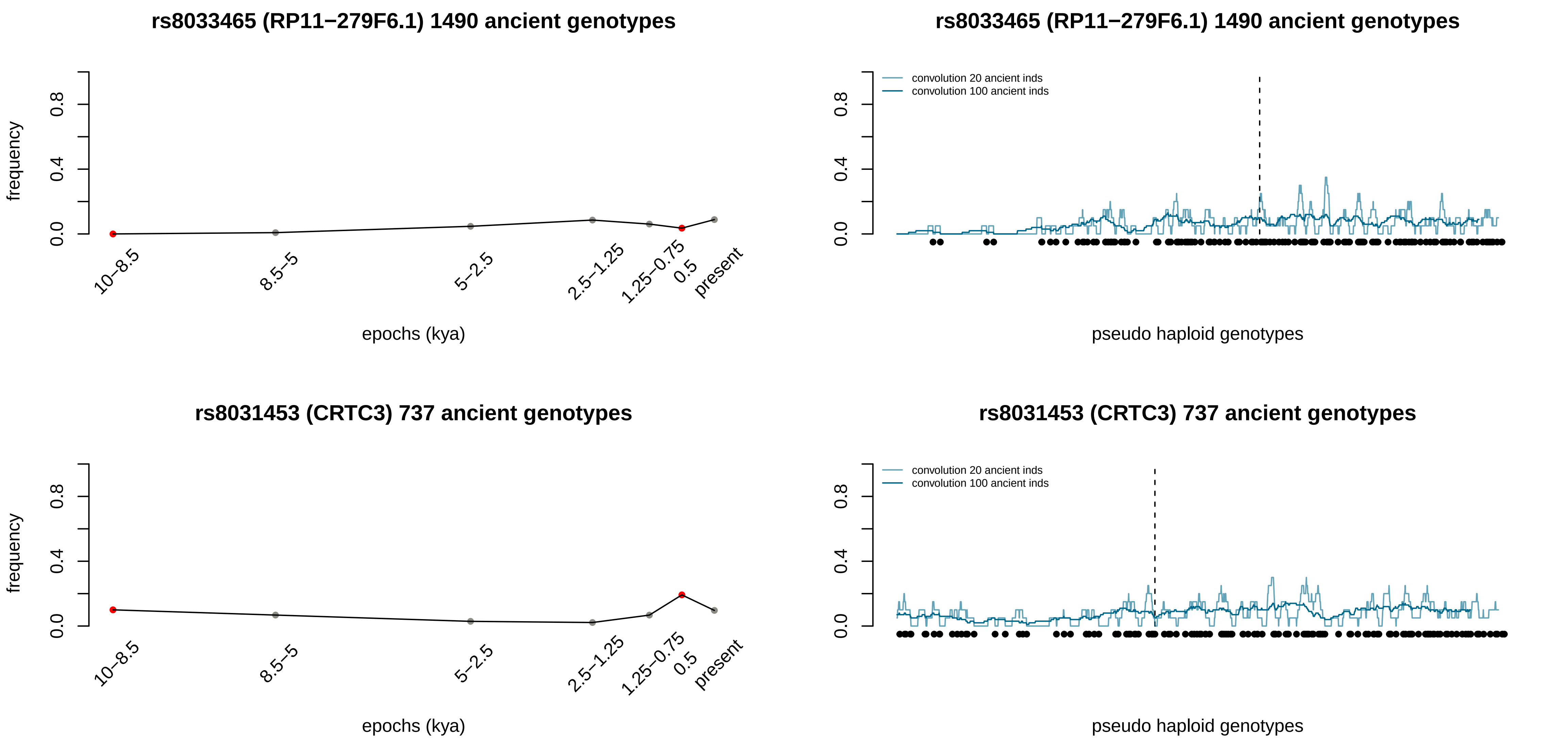
rs8033465 (RP11−279F6.1) 1490 ancient genotypes
frequency
0.0
0.4
0.8
10−8.5
8.5−5
5−2.5
2.5−1.25
1.25−0.75
0.5
present
epochs (kya)
rs8033465 (RP11−279F6.1) 1490 ancient genotypes
convolution 20 ancient inds
convolution 100 ancient inds
pseudo haploid genotypes
rs8031453 (CRTC3) 737 ancient genotypes
frequency
epochs (kya)
rs8031453 (CRTC3) 737 ancient genotypes
convolution 20 ancient inds
convolution 100 ancient inds
pseudo haploid genotypes

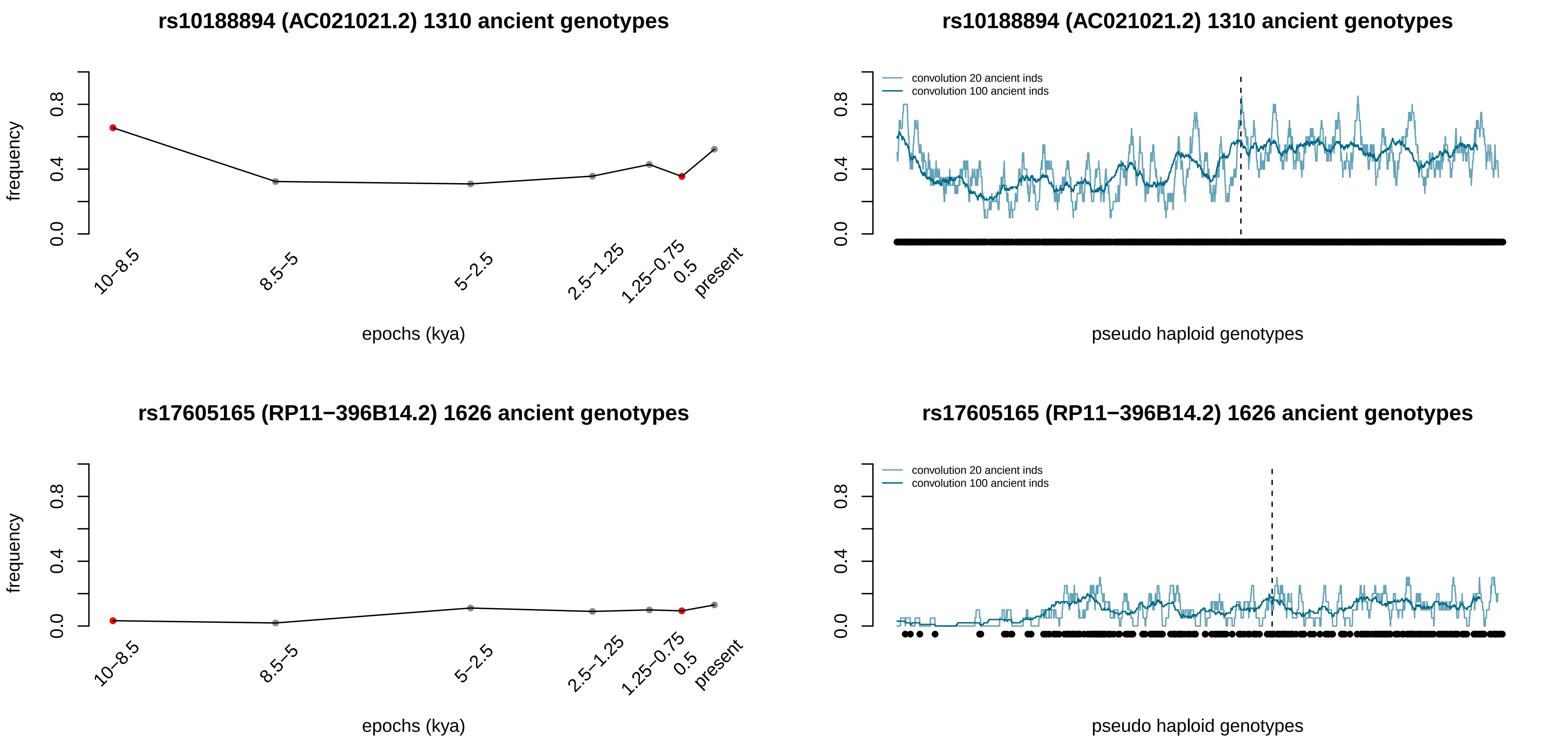
rs10188894 (AC021021.2) 1310 ancient genotypes
frequency
0.0
0.4
0.8
10–8.5
8.5–5
5–2.5
2.5–1.25
1.25–0.75
0.5
present
epochs (kya)
rs10188894 (AC021021.2) 1310 ancient genotypes
convolution 20 ancient inds
convolution 100 ancient inds
0.0
0.4
0.8
pseudo haploid genotypes
rs17605165 (RP11−396B14.2) 1626 ancient genotypes
frequency
0.0
0.4
0.8
10–8.5
8.5–5
5–2.5
2.5–1.25
1.25–0.75
0.5
present
epochs (kya)
rs17605165 (RP11−396B14.2) 1626 ancient genotypes
convolution 20 ancient inds
convolution 100 ancient inds
0.0
0.4
0.8
pseudo haploid genotypes

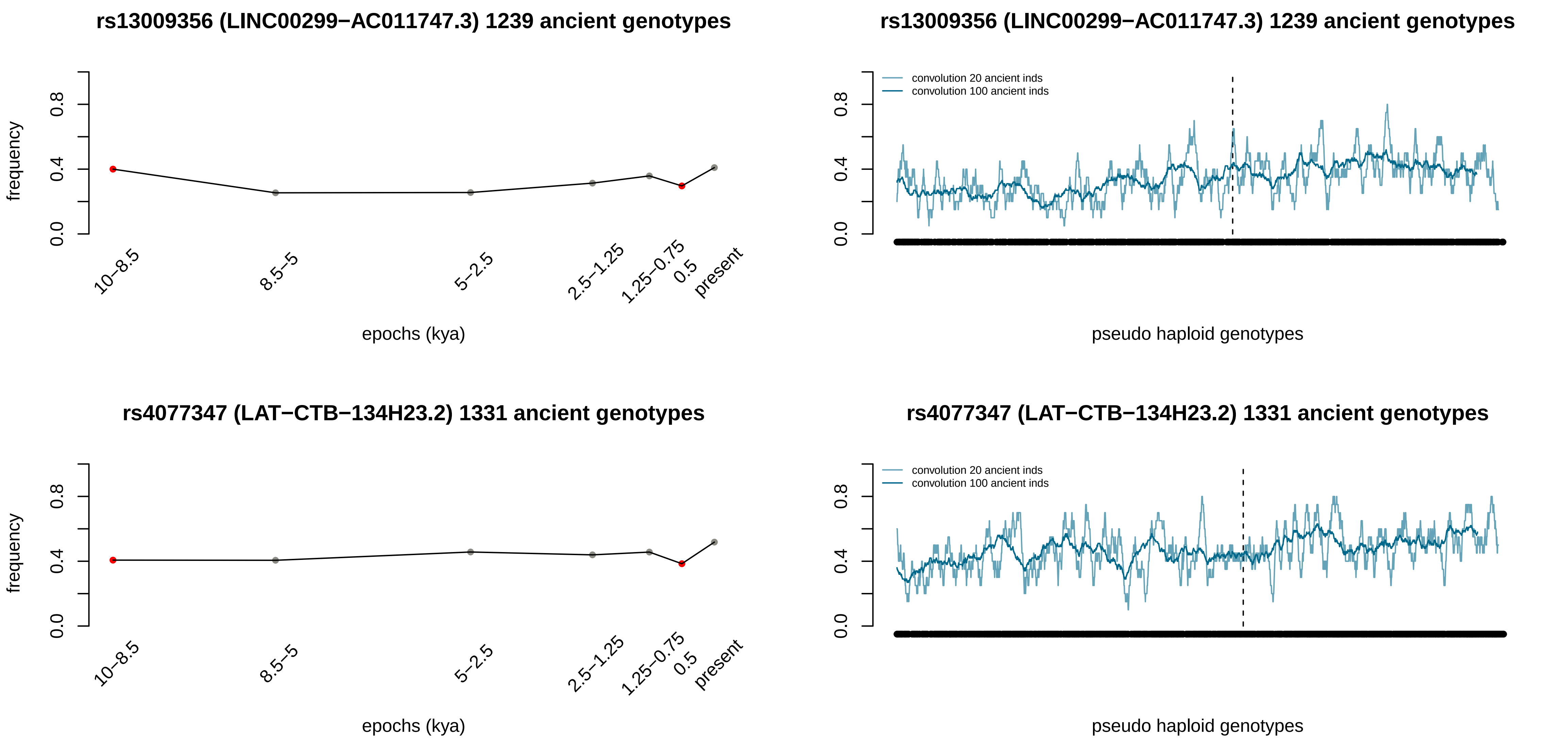
rs13009356 (LINC00299−AC011747.3) 1239 ancient genotypes
frequency
0.0
0.4
0.8
10−8.5
8.5−5
5−2.5
2.5−1.25
1.25−0.75
0.5
present
epochs (kya)
rs13009356 (LINC00299−AC011747.3) 1239 ancient genotypes
convolution 20 ancient inds
convolution 100 ancient inds
pseudo haploid genotypes
rs4077347 (LAT−CTB−134H23.2) 1331 ancient genotypes
frequency
epochs (kya)
rs4077347 (LAT−CTB−134H23.2) 1331 ancient genotypes
convolution 20 ancient inds
convolution 100 ancient inds
pseudo haploid genotypes

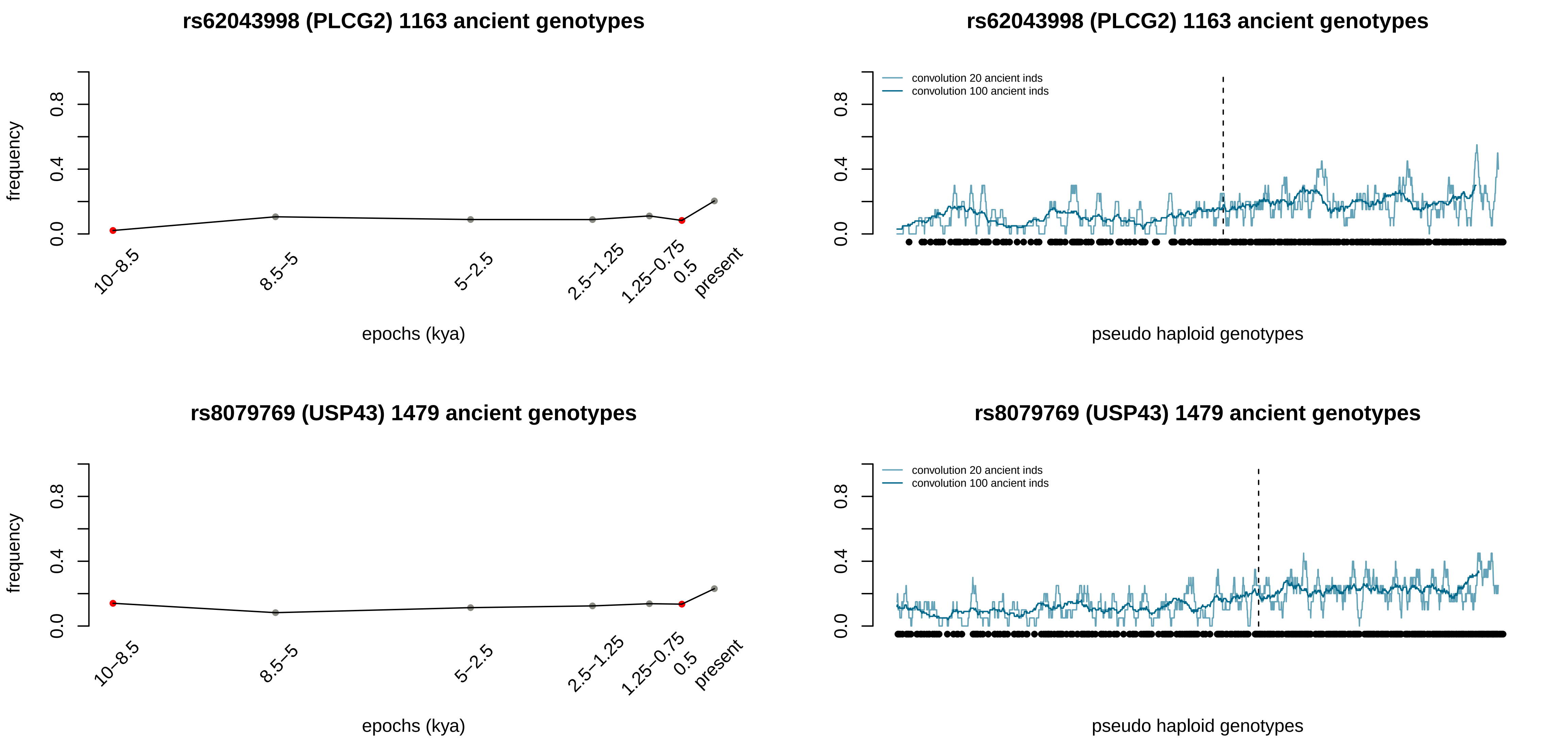
rs62043998 (PLCG2) 1163 ancient genotypes
frequency
0.0
0.4
0.8
10–8.5
8.5–5
5–2.5
2.5–1.25
1.25–0.75
0.5
present
epochs (kya)
rs62043998 (PLCG2) 1163 ancient genotypes
convolution 20 ancient inds
convolution 100 ancient inds
pseudo haploid genotypes
rs8079769 (USP43) 1479 ancient genotypes
frequency
epochs (kya)
rs8079769 (USP43) 1479 ancient genotypes
convolution 20 ancient inds
convolution 100 ancient inds
pseudo haploid genotypes

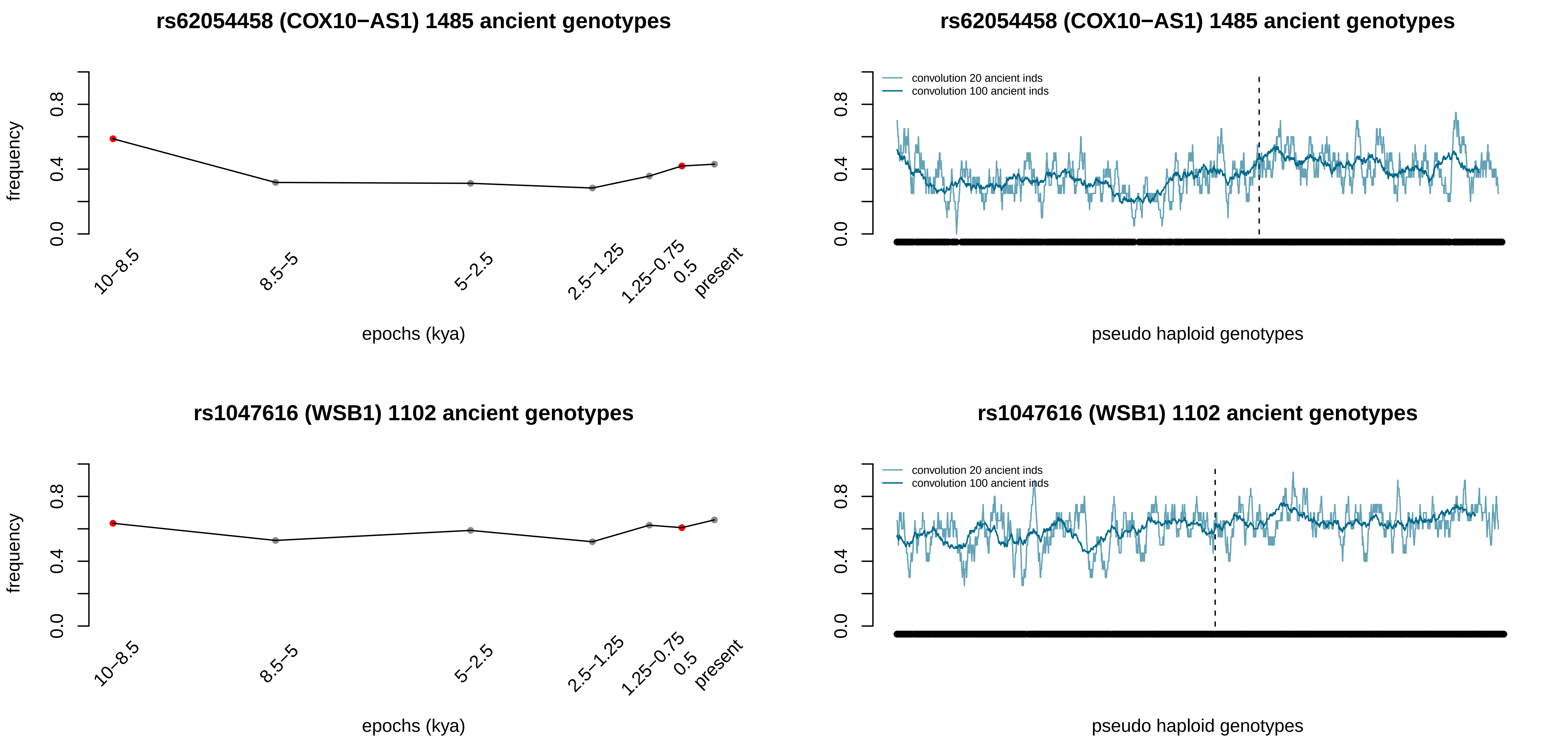
rs62054458 (COX10−AS1) 1485 ancient genotypes
frequency
0.0
0.4
0.8
10−8.5
8.5−5
5−2.5
2.5−1.25
1.25−0.75
0.5
present
epochs (kya)
rs62054458 (COX10−AS1) 1485 ancient genotypes
convolution 20 ancient inds
convolution 100 ancient inds
pseudo haploid genotypes
rs1047616 (WSB1) 1102 ancient genotypes
frequency
epochs (kya)
rs1047616 (WSB1) 1102 ancient genotypes
convolution 20 ancient inds
convolution 100 ancient inds
pseudo haploid genotypes

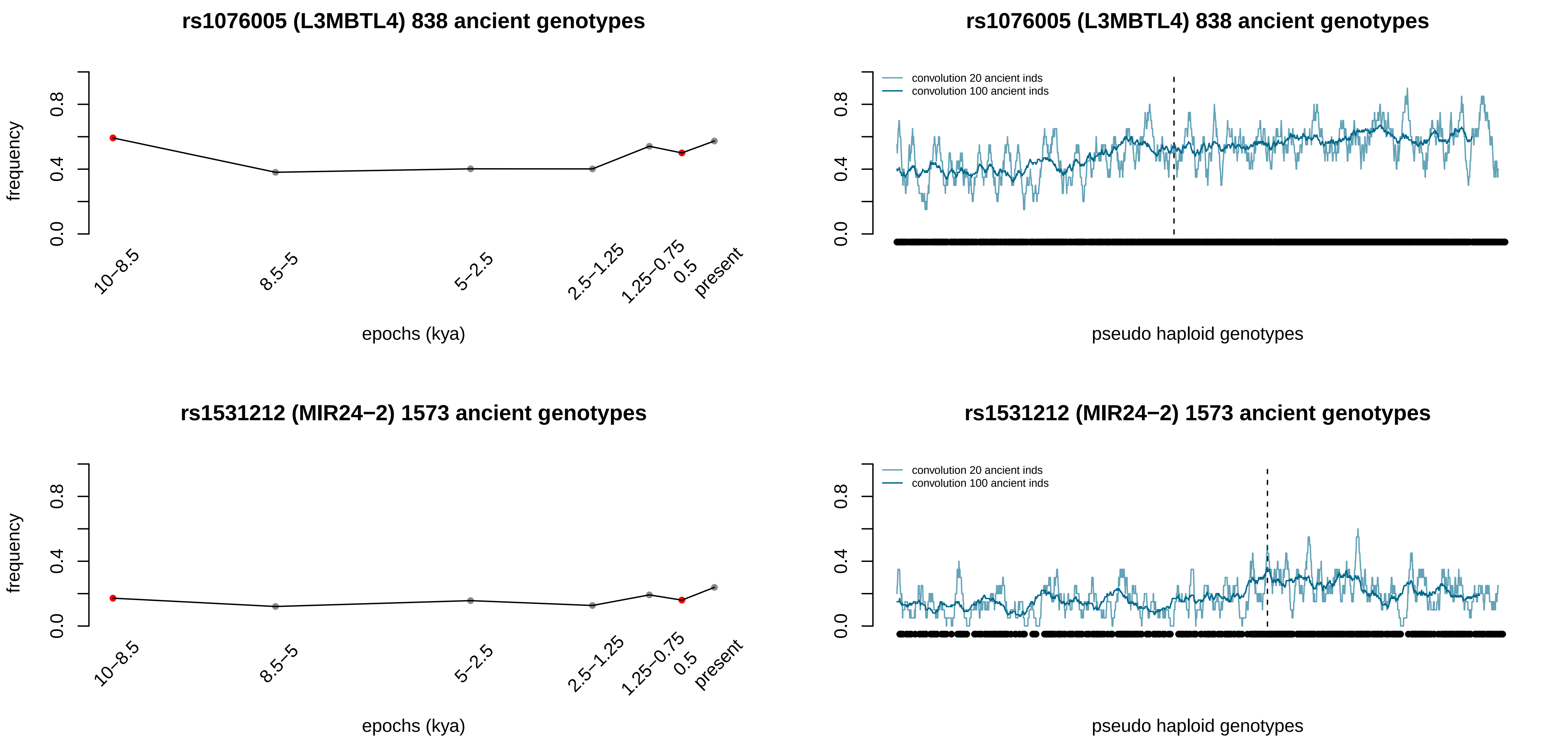
rs1076005 (L3MBTL4) 838 ancient genotypes
frequency
0.0
0.4
0.8
10–8.5
8.5–5
5–2.5
2.5–1.25
1.25–0.75
0.5
present
epochs (kya)
rs1076005 (L3MBTL4) 838 ancient genotypes
convolution 20 ancient inds
convolution 100 ancient inds
pseudo haploid genotypes
rs1531212 (MIR24−2) 1573 ancient genotypes
frequency
epochs (kya)
rs1531212 (MIR24−2) 1573 ancient genotypes
convolution 20 ancient inds
convolution 100 ancient inds
pseudo haploid genotypes

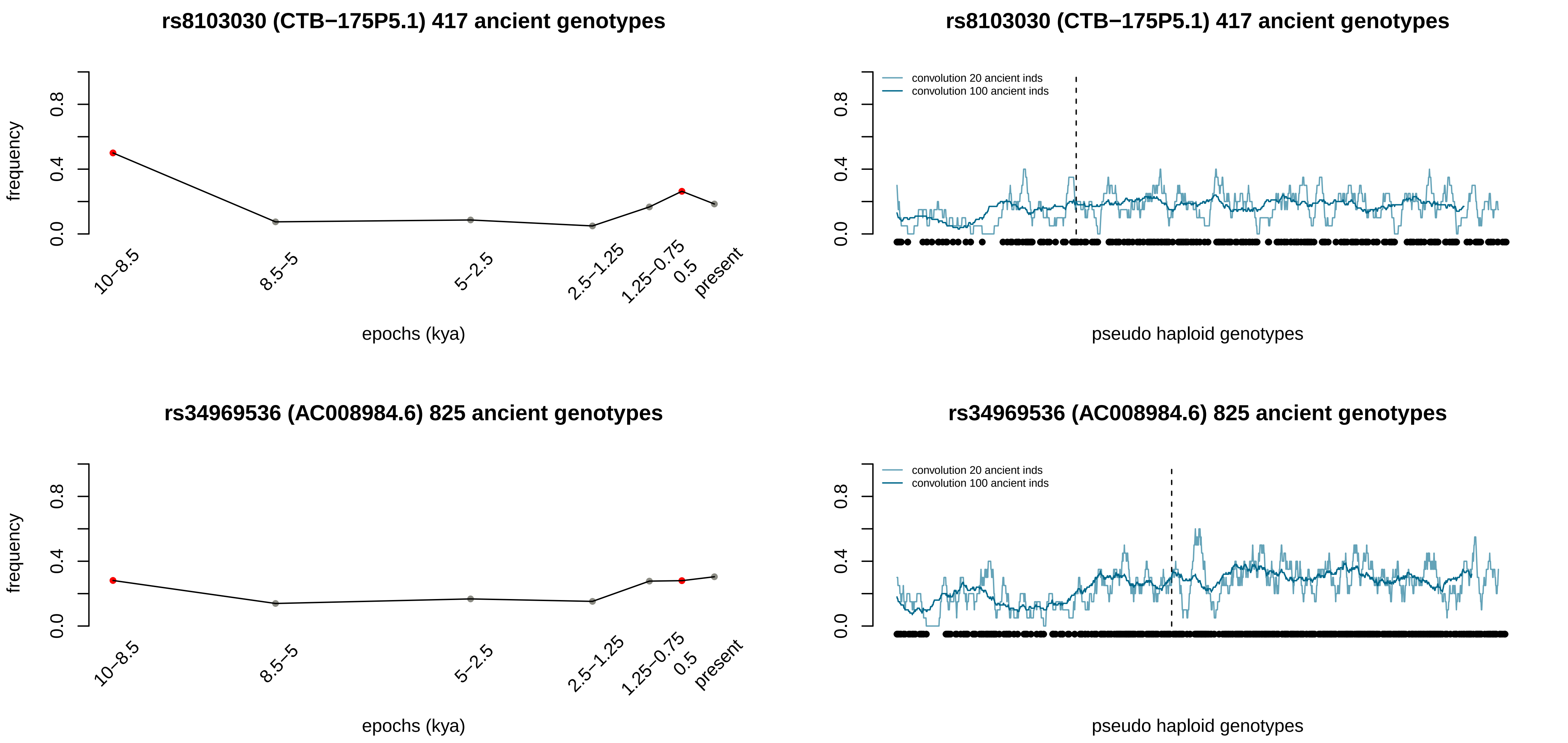

rs8103030 (CTB−175P5.1) 417 ancient genotypes
frequency
0.0
0.4
0.8
10–8.5
8.5–5
5–2.5
2.5–1.25
1.25–0.75
0.5
present
epochs (kya)
rs8103030 (CTB−175P5.1) 417 ancient genotypes
convolution 20 ancient inds
convolution 100 ancient inds
pseudo haploid genotypes
rs34969536 (AC008984.6) 825 ancient genotypes
frequency
epochs (kya)
rs34969536 (AC008984.6) 825 ancient genotypes
convolution 20 ancient inds
convolution 100 ancient inds
pseudo haploid genotypes

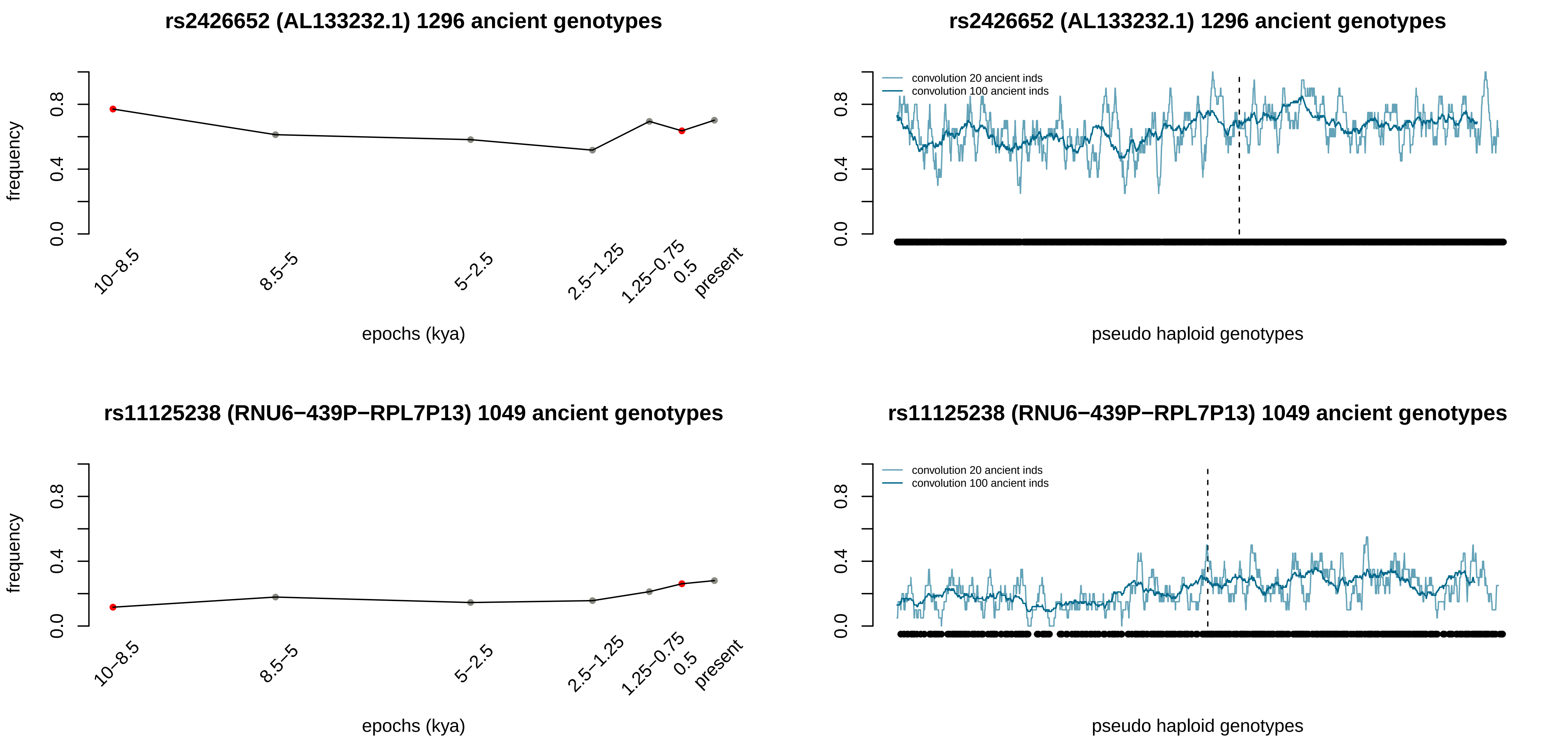

rs2426652 (AL133232.1) 1296 ancient genotypes
frequency
0.0
0.4
0.8
10−8.5
8.5−5
5−2.5
2.5−1.25
1.25−0.75
0.5
present
epochs (kya)
rs2426652 (AL133232.1) 1296 ancient genotypes
convolution 20 ancient inds
convolution 100 ancient inds
pseudo haploid genotypes
rs11125238 (RNU6−439P−RPL7P13) 1049 ancient genotypes
frequency
epochs (kya)
rs11125238 (RNU6−439P−RPL7P13) 1049 ancient genotypes
convolution 20 ancient inds
convolution 100 ancient inds
pseudo haploid genotypes

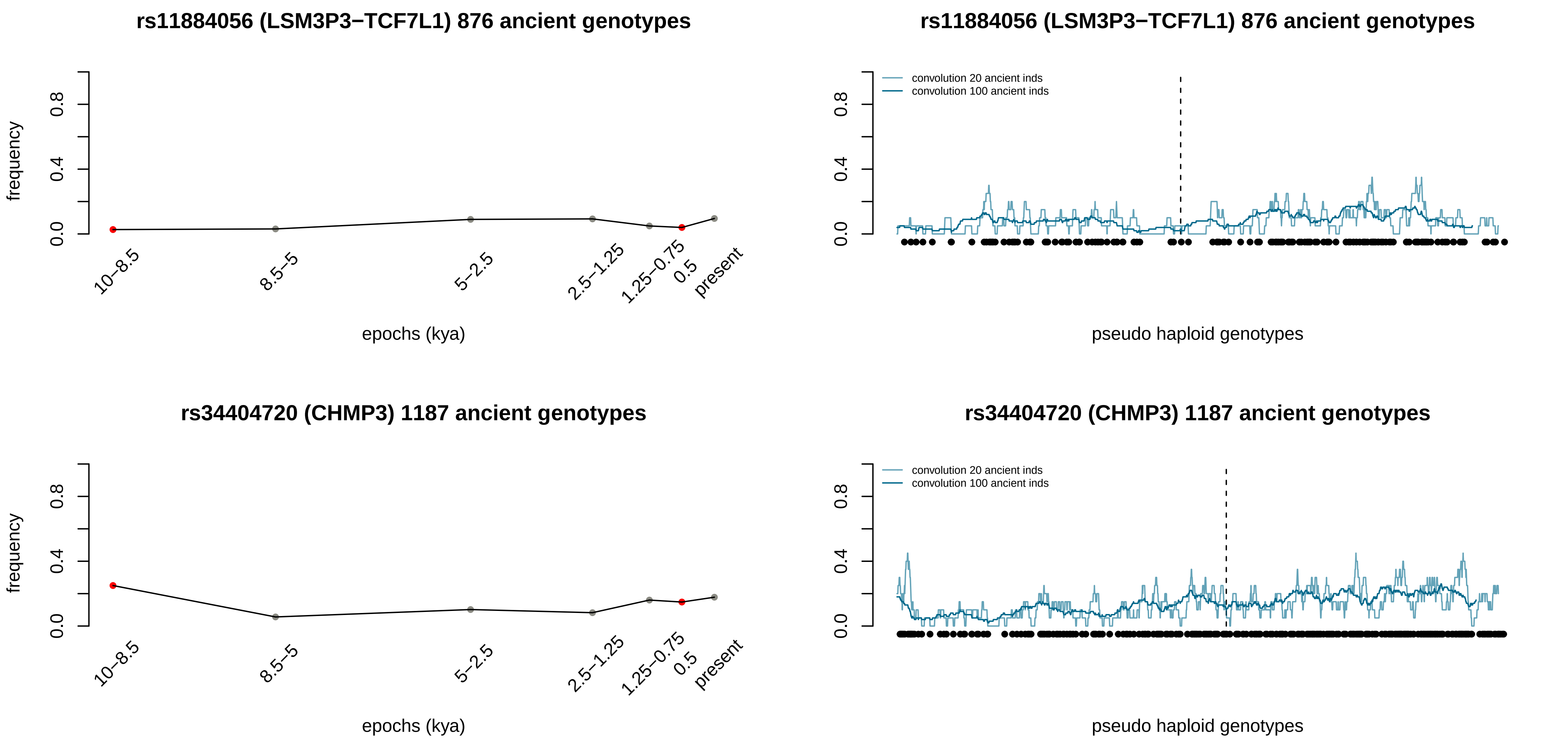
rs11884056 (LSM3P3−TCF7L1) 876 ancient genotypes
frequency
0.0
0.4
0.8
10–8.5
8.5–5
5–2.5
2.5–1.25
1.25–0.75
0.5
present
epochs (kya)
rs11884056 (LSM3P3−TCF7L1) 876 ancient genotypes
convolution 20 ancient inds
convolution 100 ancient inds
pseudo haploid genotypes
rs34404720 (CHMP3) 1187 ancient genotypes
frequency
epochs (kya)
rs34404720 (CHMP3) 1187 ancient genotypes
convolution 20 ancient inds
convolution 100 ancient inds
pseudo haploid genotypes

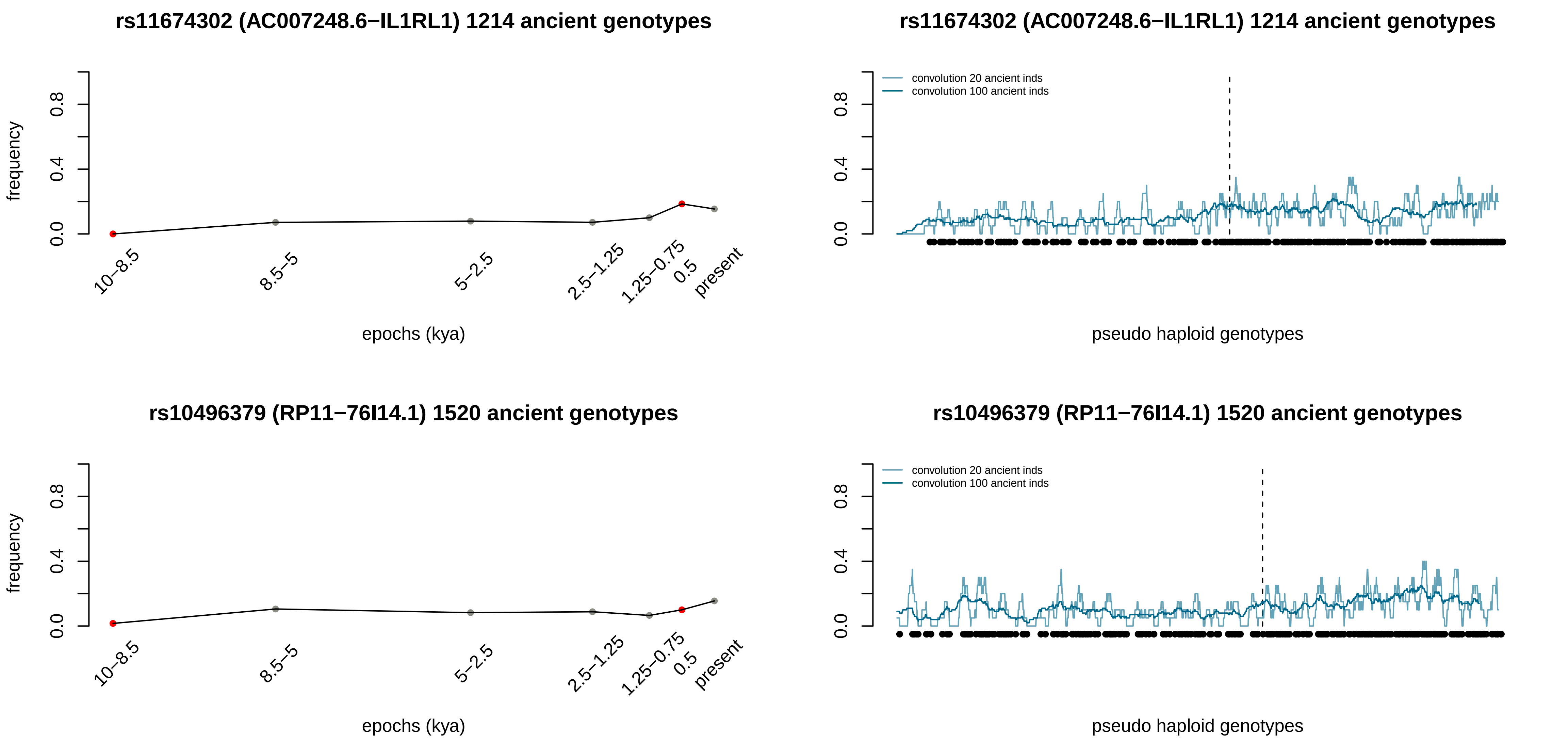
rs11674302 (AC007248.6−IL1RL1) 1214 ancient genotypes
frequency
0.0
0.4
0.8
10−8.5
8.5−5
5−2.5
2.5−1.25
1.25−0.75
0.5
present
epochs (kya)
rs11674302 (AC007248.6−IL1RL1) 1214 ancient genotypes
convolution 20 ancient inds
convolution 100 ancient inds
pseudo haploid genotypes
rs10496379 (RP11−76I14.1) 1520 ancient genotypes
frequency
epochs (kya)
rs10496379 (RP11−76I14.1) 1520 ancient genotypes
convolution 20 ancient inds
convolution 100 ancient inds
pseudo haploid genotypes

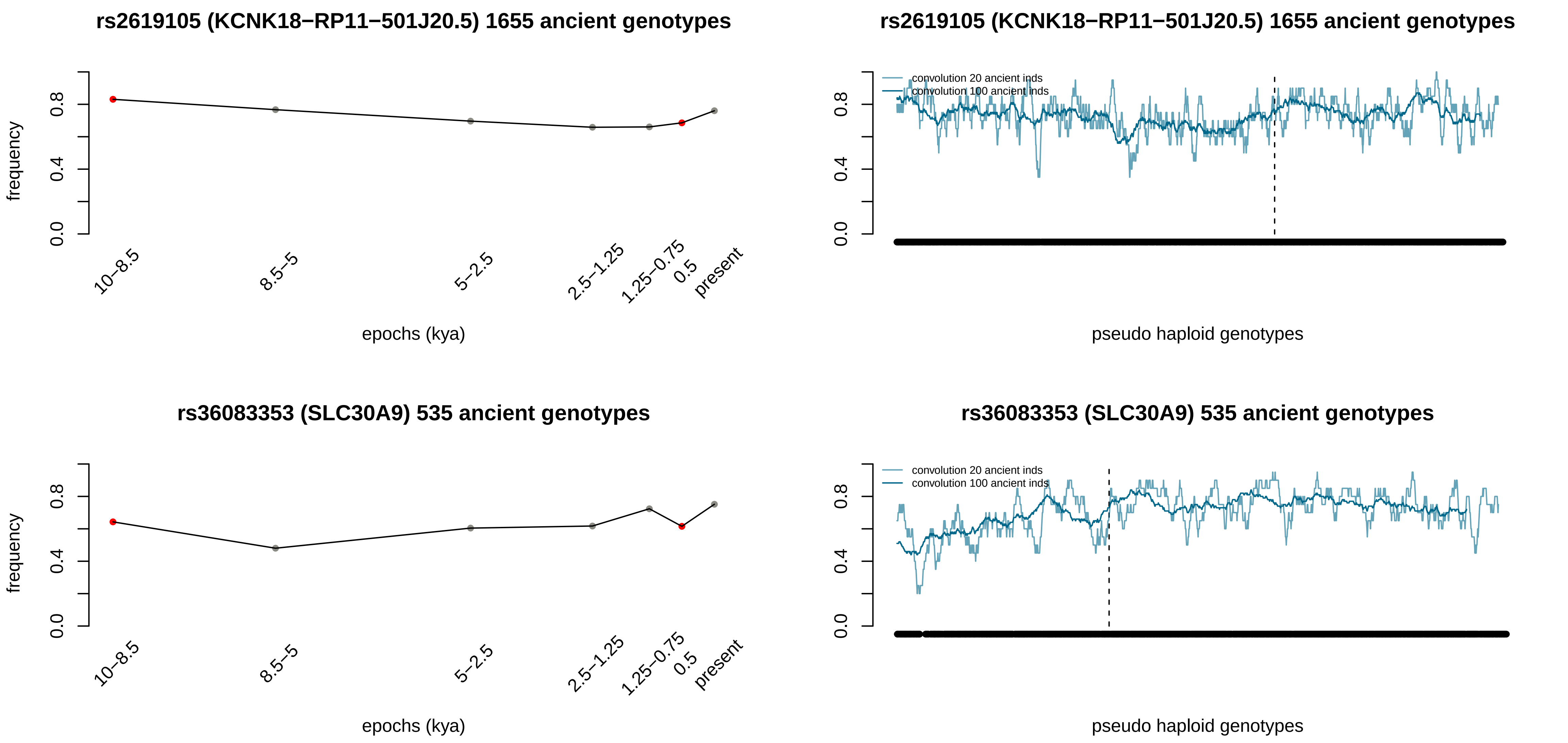

rs2619105 (KCNK18−RP11−501J20.5) 1655 ancient genotypes
frequency
0.0
0.4
0.8
10−8.5
8.5−5
5−2.5
2.5−1.25
1.25−0.75
0.5
present
epochs (kya)
rs2619105 (KCNK18−RP11−501J20.5) 1655 ancient genotypes
convolution 20 ancient inds
convolution 100 ancient inds
pseudo haploid genotypes
rs36083353 (SLC30A9) 535 ancient genotypes
frequency
epochs (kya)
rs36083353 (SLC30A9) 535 ancient genotypes
convolution 20 ancient inds
convolution 100 ancient inds
pseudo haploid genotypes

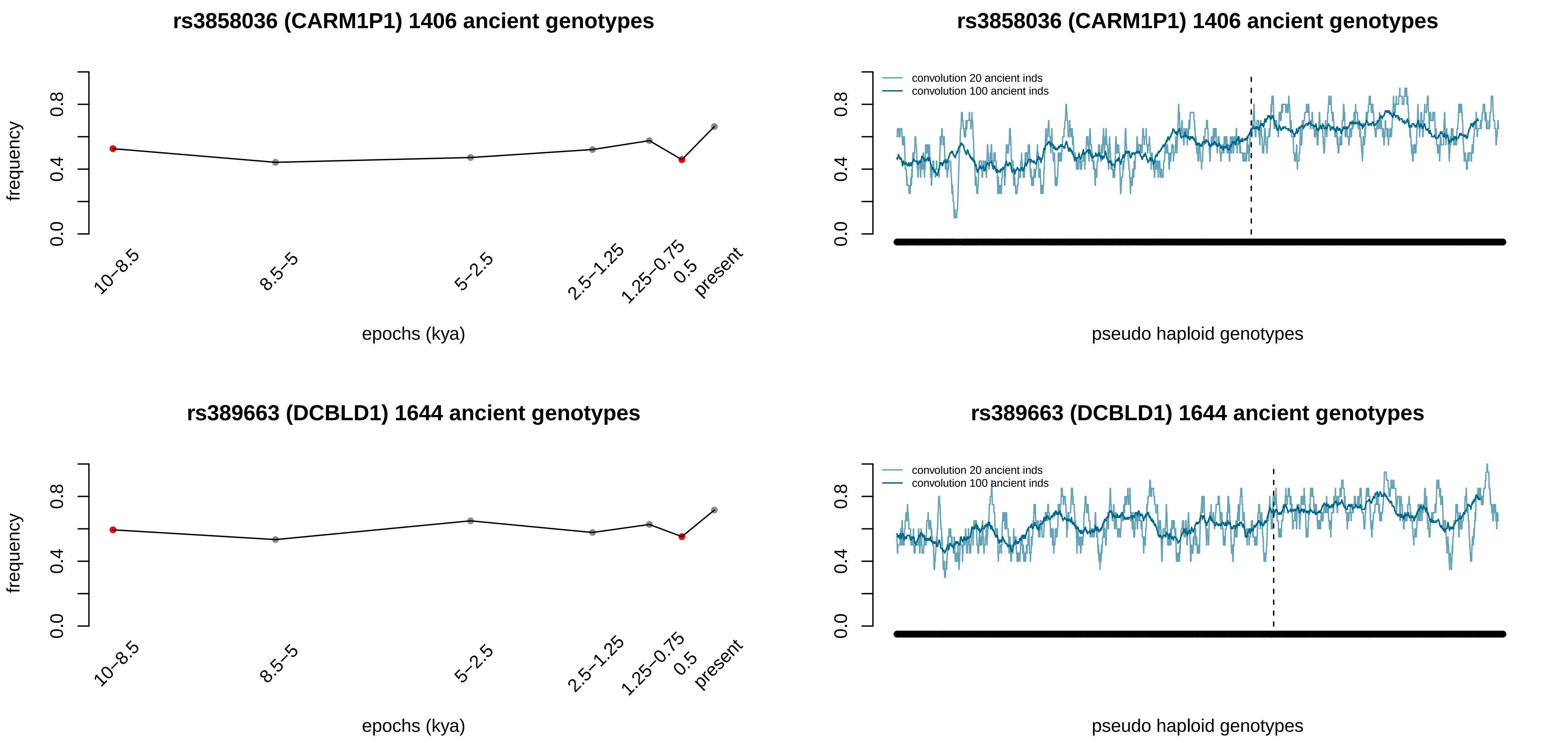

rs3858036 (CARM1P1) 1406 ancient genotypes
frequency
0.0
0.4
0.8
10–8.5
8.5–5
5–2.5
2.5–1.25
1.25–0.75
0.5
present
epochs (kya)
rs3858036 (CARM1P1) 1406 ancient genotypes
convolution 20 ancient inds
convolution 100 ancient inds
pseudo haploid genotypes
rs389663 (DCBLD1) 1644 ancient genotypes
frequency
epochs (kya)
rs389663 (DCBLD1) 1644 ancient genotypes
convolution 20 ancient inds
convolution 100 ancient inds
pseudo haploid genotypes

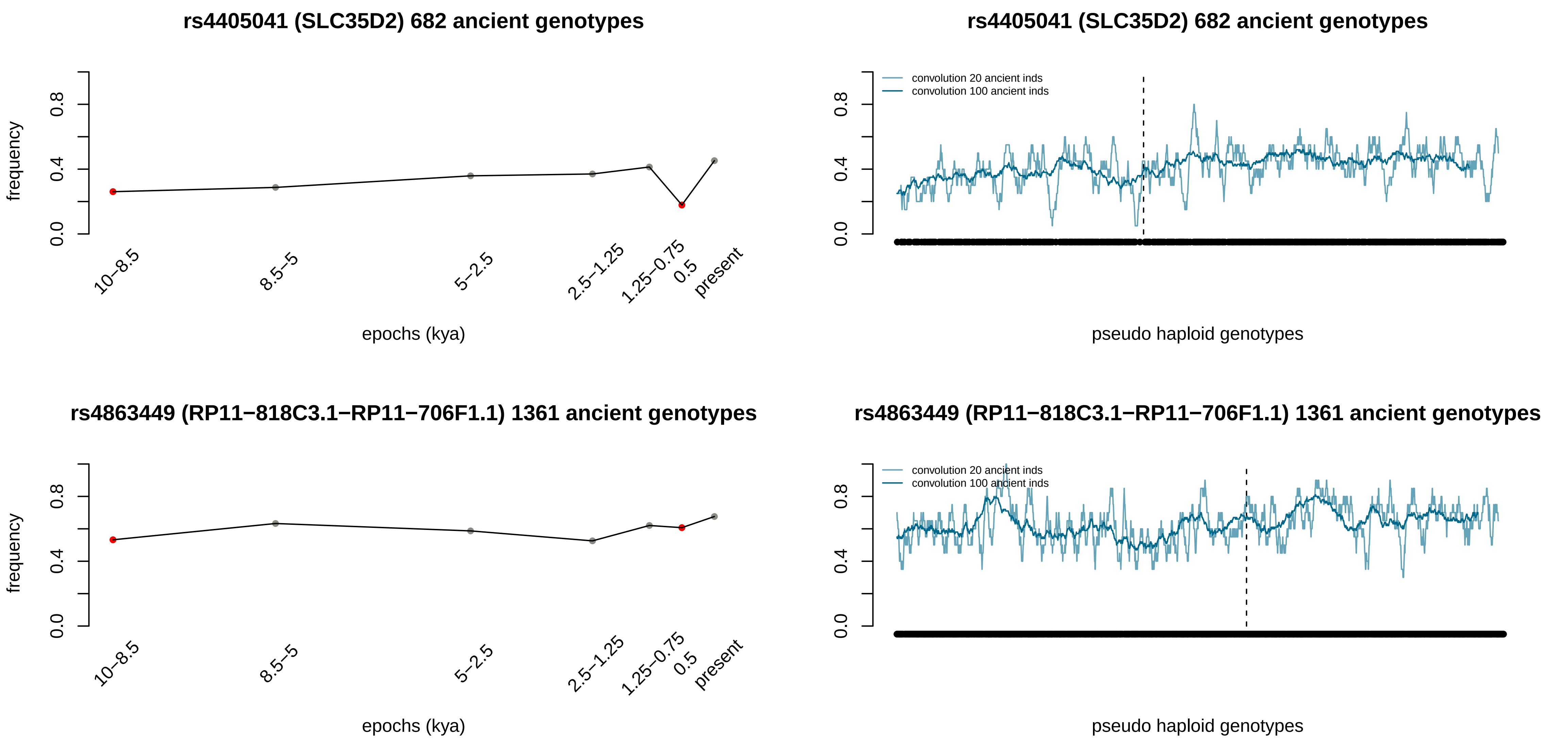
rs4405041 (SLC35D2) 682 ancient genotypes
frequency
0.0
0.4
0.8
10−8.5
8.5−5
5−2.5
2.5−1.25
1.25−0.75
0.5
present
epochs (kya)
rs4405041 (SLC35D2) 682 ancient genotypes
convolution 20 ancient inds
convolution 100 ancient inds
pseudo haploid genotypes
rs4863449 (RP11−818C3.1−RP11−706F1.1) 1361 ancient genotypes
frequency
epochs (kya)
rs4863449 (RP11−818C3.1−RP11−706F1.1) 1361 ancient genotypes
convolution 20 ancient inds
convolution 100 ancient inds
pseudo haploid genotypes

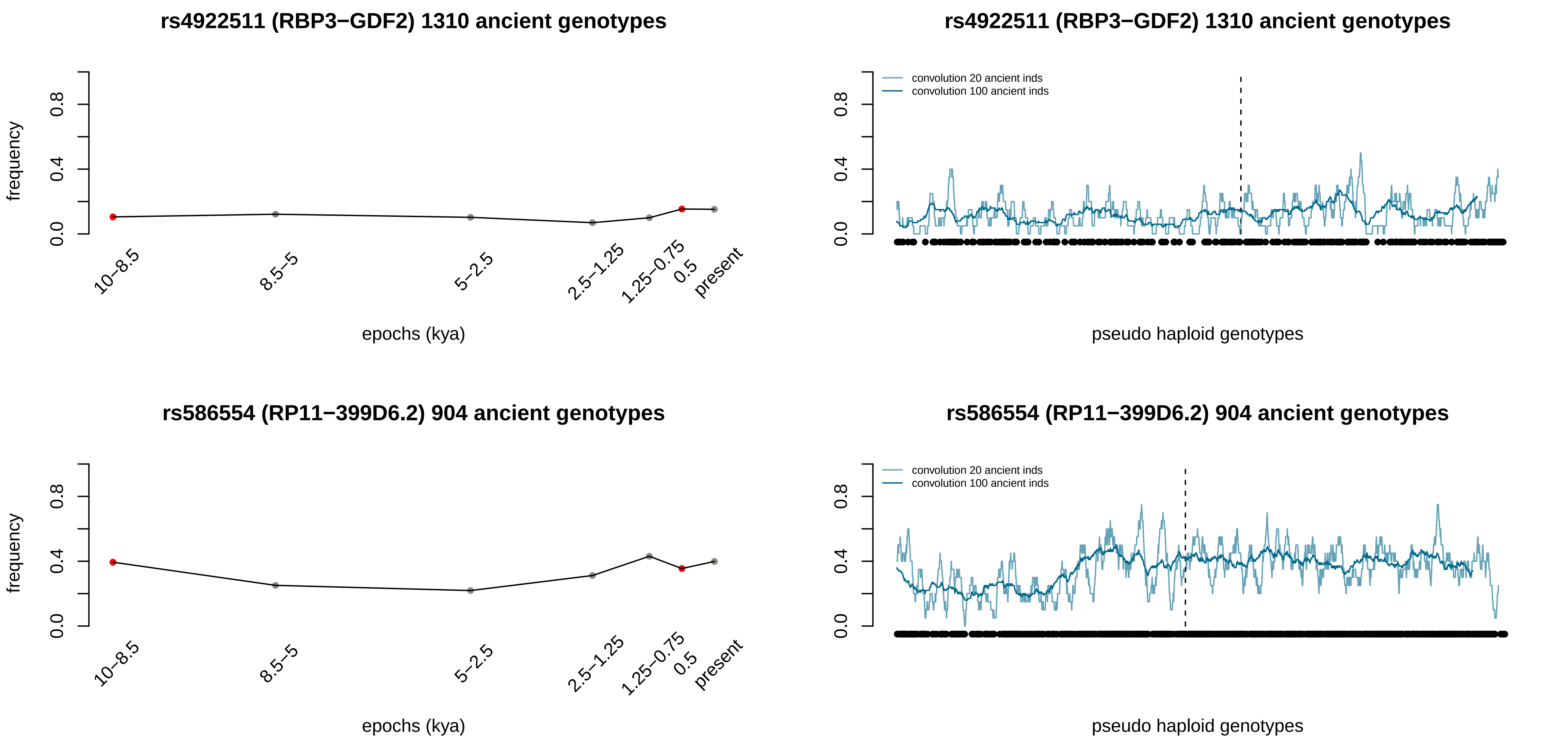
rs4922511 (RBP3−GDF2) 1310 ancient genotypes
frequency
0.0
0.4
0.8
10−8.5
8.5−5
5−2.5
2.5−1.25
1.25−0.75
0.5
present
epochs (kya)
rs4922511 (RBP3−GDF2) 1310 ancient genotypes
convolution 20 ancient inds
convolution 100 ancient inds
0.0
0.4
0.8
pseudo haploid genotypes
rs586554 (RP11−399D6.2) 904 ancient genotypes
frequency
0.0
0.4
0.8
10−8.5
8.5−5
5−2.5
2.5−1.25
1.25−0.75
0.5
present
epochs (kya)
rs586554 (RP11−399D6.2) 904 ancient genotypes
convolution 20 ancient inds
convolution 100 ancient inds
0.0
0.4
0.8
pseudo haploid genotypes

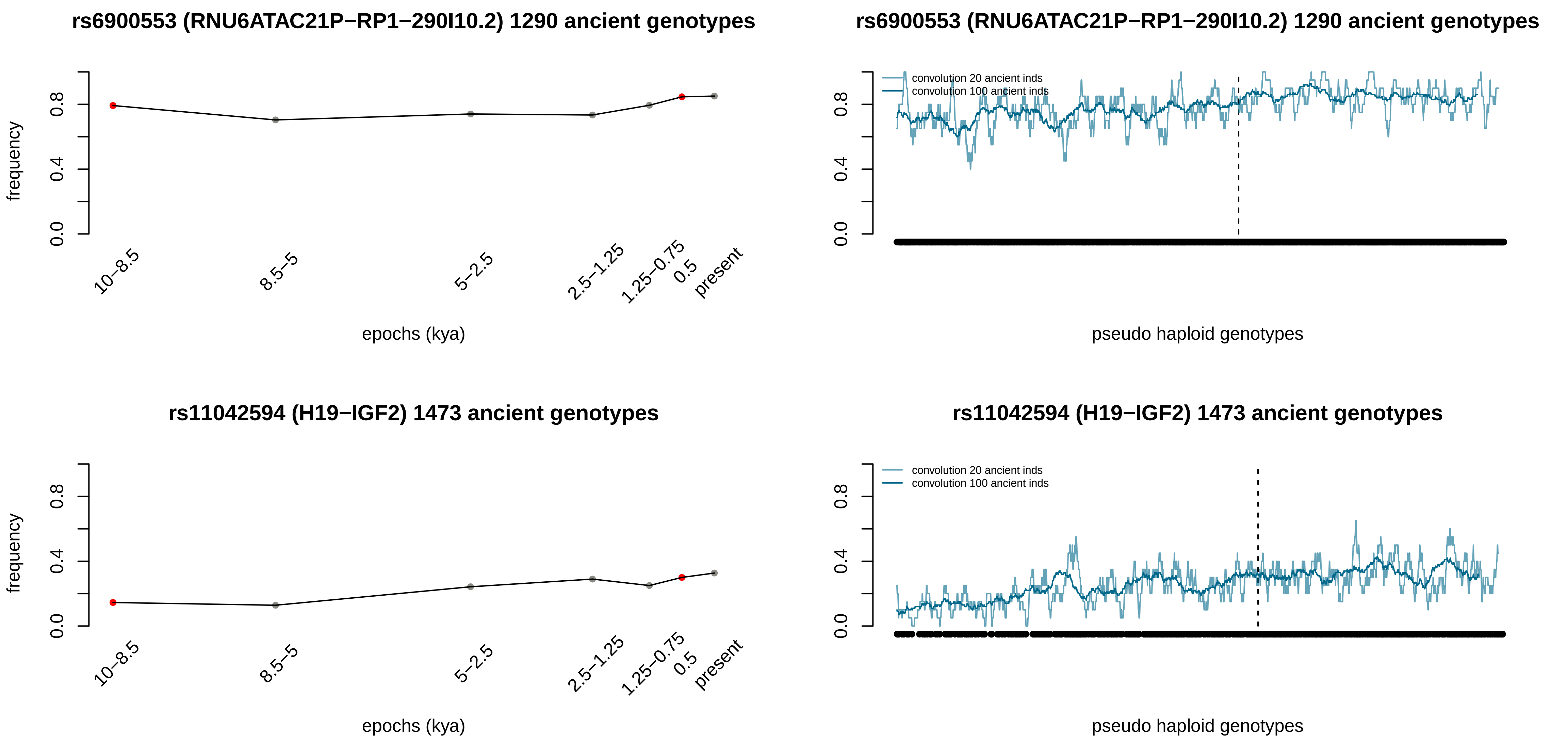
rs6900553 (RNU6ATAC21P−RP1−290I10.2) 1290 ancient genotypes
frequency
0.0
0.4
0.8
10−8.5
8.5−5
5−2.5
2.5−1.25
1.25−0.75
0.5
present
epochs (kya)
rs6900553 (RNU6ATAC21P−RP1−290I10.2) 1290 ancient genotypes
convolution 20 ancient inds
convolution 100 ancient inds
0.0
0.4
0.8
pseudo haploid genotypes
rs11042594 (H19−IGF2) 1473 ancient genotypes
frequency
0.0
0.4
0.8
10−8.5
8.5−5
5−2.5
2.5−1.25
1.25−0.75
0.5
present
epochs (kya)
rs11042594 (H19−IGF2) 1473 ancient genotypes
convolution 20 ancient inds
convolution 100 ancient inds
0.0
0.4
0.8
pseudo haploid genotypes

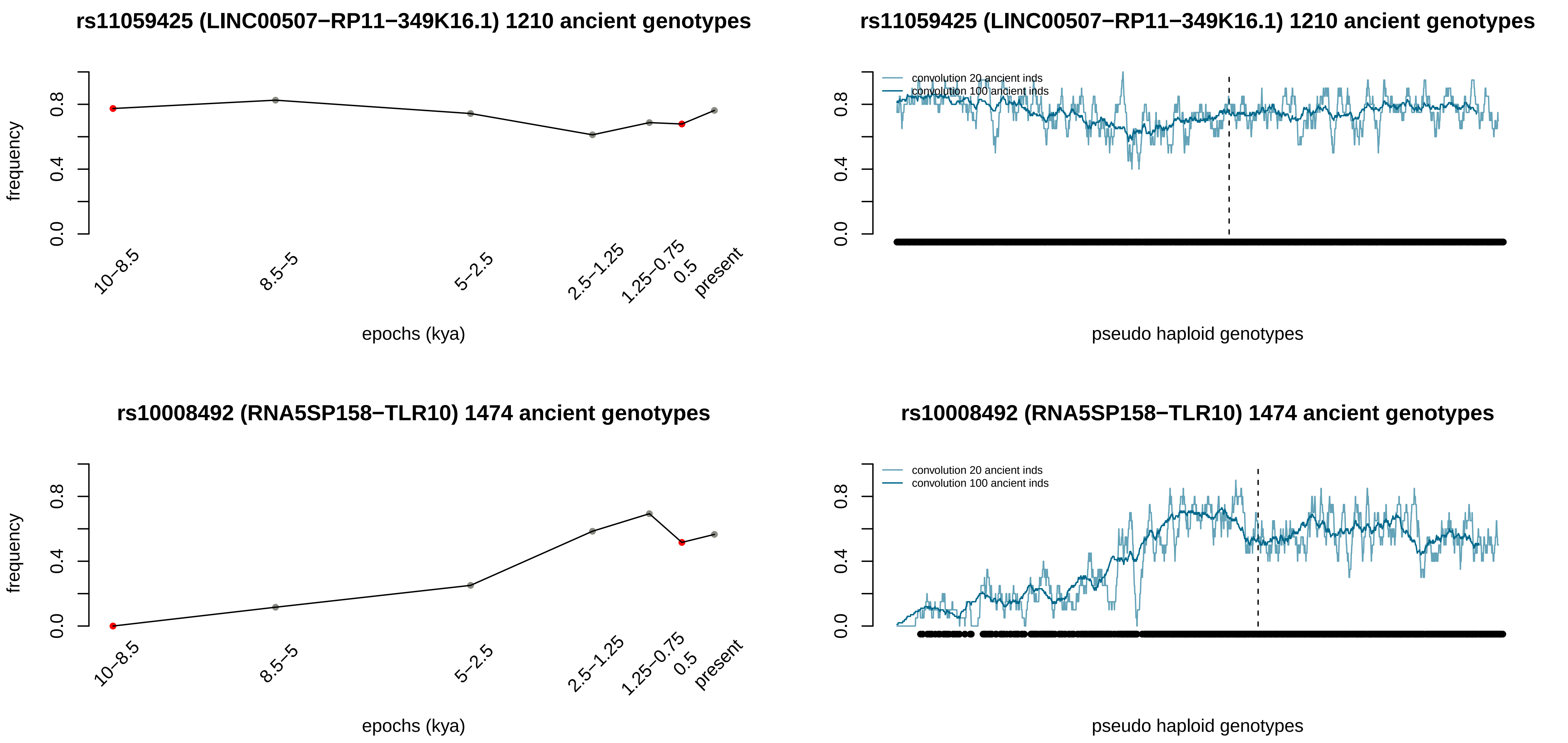
rs11059425 (LINC00507−RP11−349K16.1) 1210 ancient genotypes
frequency
0.0
0.4
0.8
10−8.5
8.5−5
5−2.5
2.5−1.25
1.25−0.75
0.5
present
epochs (kya)
rs11059425 (LINC00507−RP11−349K16.1) 1210 ancient genotypes
convolution 20 ancient inds
convolution 100 ancient inds
pseudo haploid genotypes
rs10008492 (RNA5SP158−TLR10) 1474 ancient genotypes
frequency
epochs (kya)
rs10008492 (RNA5SP158−TLR10) 1474 ancient genotypes
convolution 20 ancient inds
convolution 100 ancient inds
pseudo haploid genotypes

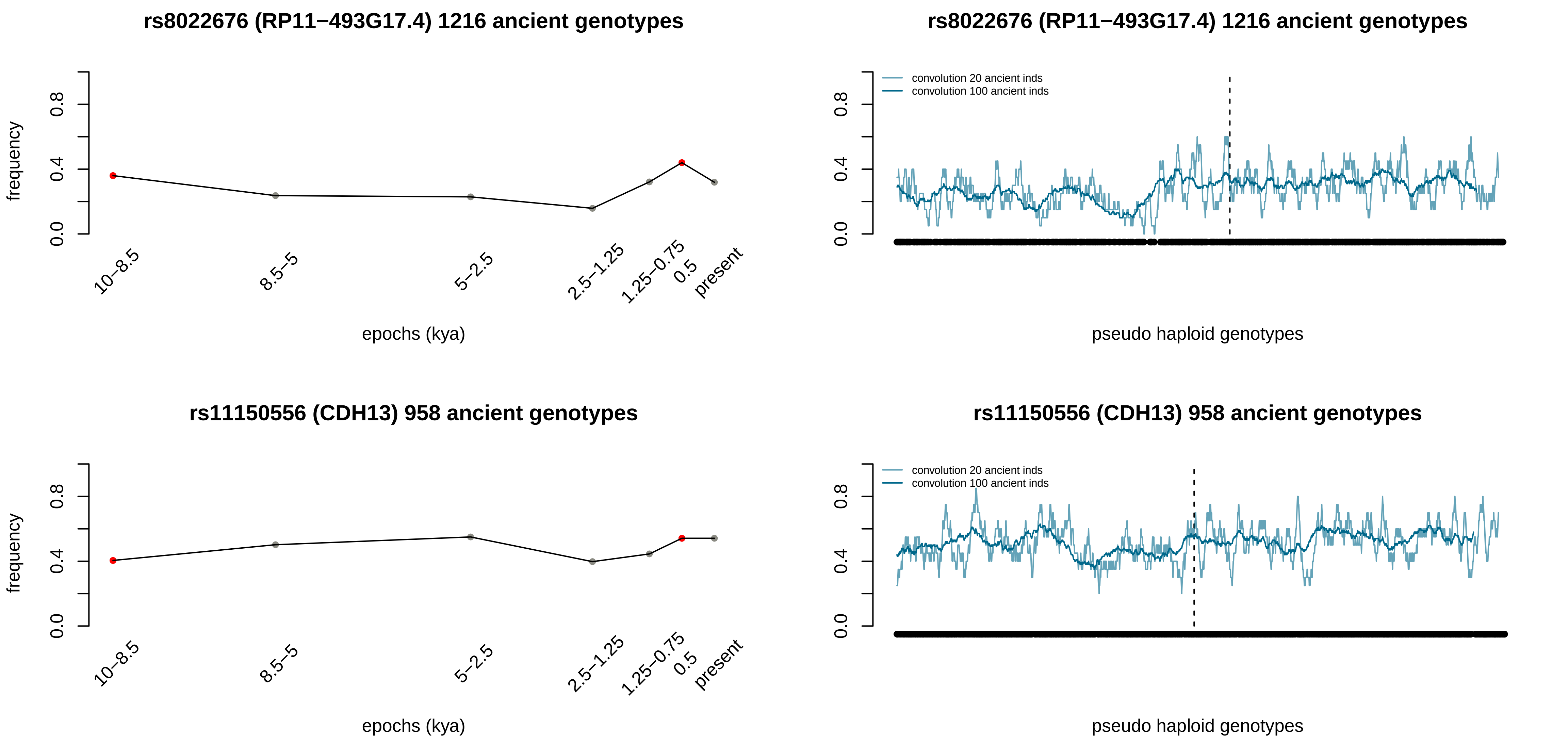
rs8022676 (RP11−493G17.4) 1216 ancient genotypes
frequency
0.0
0.4
0.8
10−8.5
8.5−5
5−2.5
2.5−1.25
1.25−0.75
0.5
present
epochs (kya)
rs8022676 (RP11−493G17.4) 1216 ancient genotypes
convolution 20 ancient inds
convolution 100 ancient inds
pseudo haploid genotypes
rs11150556 (CDH13) 958 ancient genotypes
frequency
epochs (kya)
rs11150556 (CDH13) 958 ancient genotypes
convolution 20 ancient inds
convolution 100 ancient inds
pseudo haploid genotypes

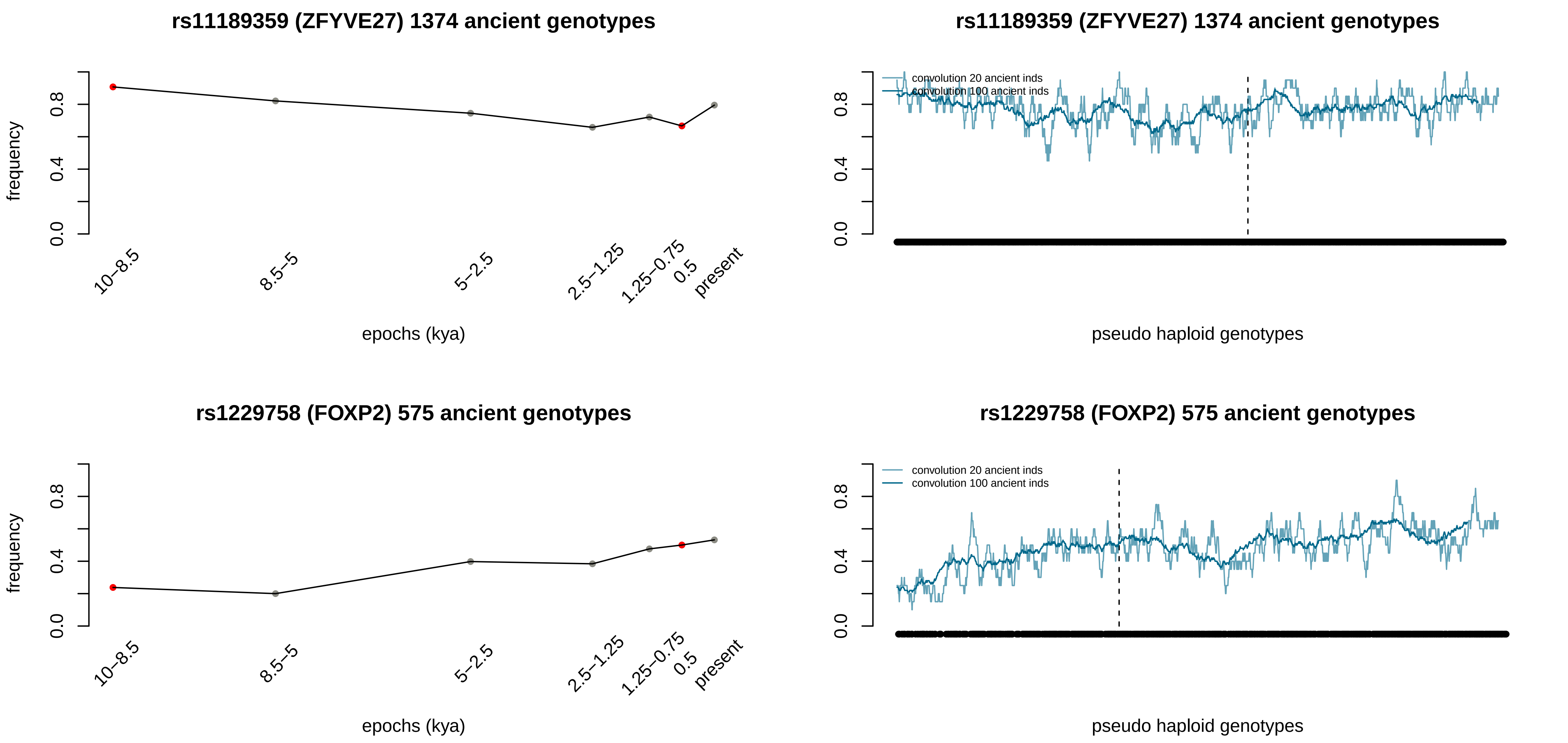
rs11189359 (ZFYVE27) 1374 ancient genotypes
frequency
0.0
0.4
0.8
10–8.5
8.5–5
5–2.5
2.5–1.25
1.25–0.75
0.5
present
epochs (kya)
rs11189359 (ZFYVE27) 1374 ancient genotypes
convolution 20 ancient inds
convolution 100 ancient inds
pseudo haploid genotypes
rs1229758 (FOXP2) 575 ancient genotypes
frequency
epochs (kya)
rs1229758 (FOXP2) 575 ancient genotypes
convolution 20 ancient inds
convolution 100 ancient inds
pseudo haploid genotypes

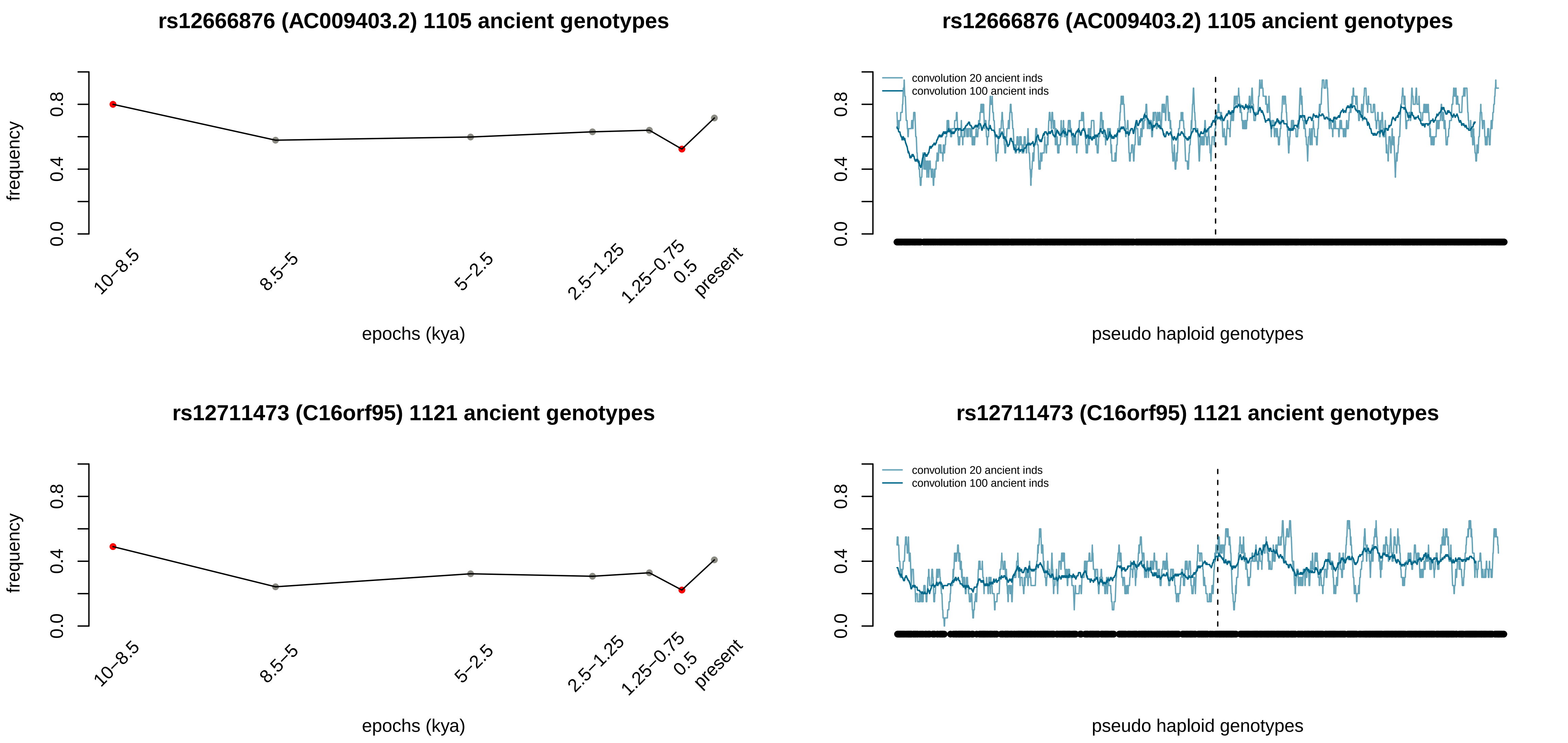
rs12666876 (AC009403.2) 1105 ancient genotypes
frequency
0.0
0.4
0.8
10–8.5
8.5–5
5–2.5
2.5–1.25
1.25–0.75
0.5
present
epochs (kya)
rs12666876 (AC009403.2) 1105 ancient genotypes
convolution 20 ancient inds
convolution 100 ancient inds
pseudo haploid genotypes
rs12711473 (C16orf95) 1121 ancient genotypes
frequency
epochs (kya)
rs12711473 (C16orf95) 1121 ancient genotypes
convolution 20 ancient inds
convolution 100 ancient inds
pseudo haploid genotypes
