## Supplementary material for "Deep estimation of the intensity and timing of selection from ancient genomes": 1D images for the 89 positively selected variants in northern Europe

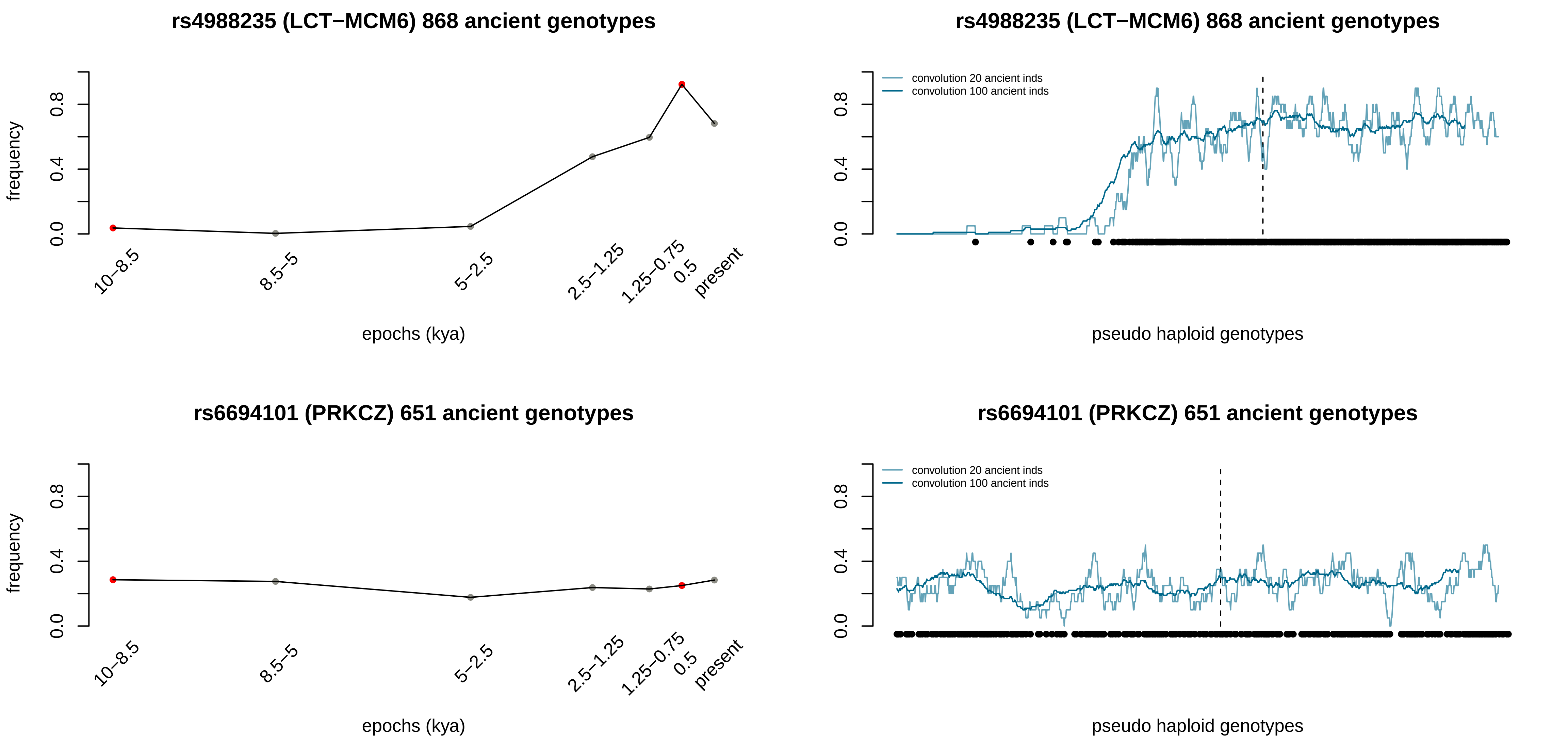

rs4988235 (LCT−MCM6) 868 ancient genotypes
frequency
0.0
0.4
0.8
10−8.5
8.5−5
5−2.5
2.5−1.25
1.25−0.75
0.5
present
epochs (kya)
rs4988235 (LCT−MCM6) 868 ancient genotypes
convolution 20 ancient inds
convolution 100 ancient inds
pseudo haploid genotypes
rs6694101 (PRKCZ) 651 ancient genotypes
frequency
epochs (kya)
rs6694101 (PRKCZ) 651 ancient genotypes
convolution 20 ancient inds
convolution 100 ancient inds
pseudo haploid genotypes

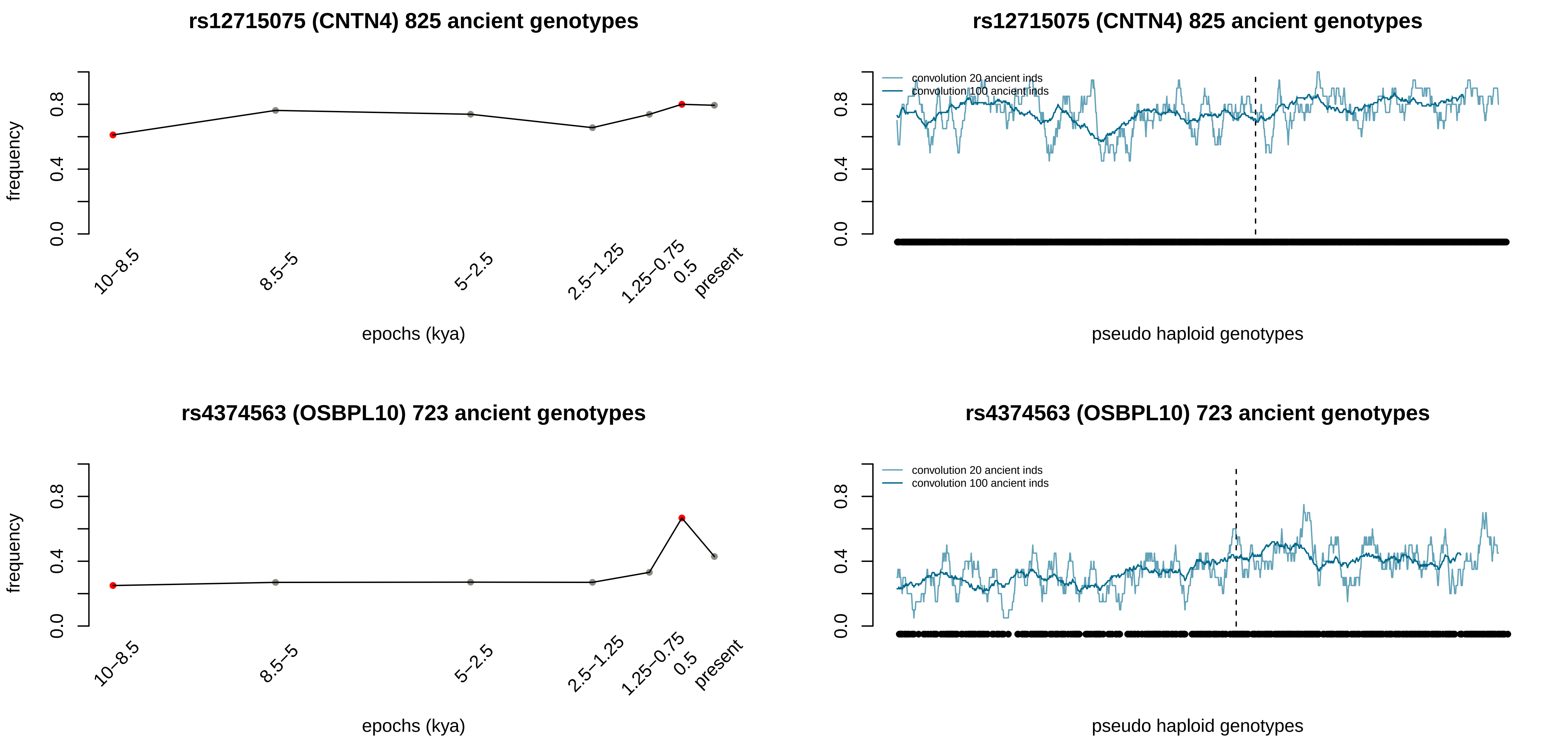

rs12715075 (CNTN4) 825 ancient genotypes
frequency
0.0
0.4
0.8
10–8.5
8.5–5
5–2.5
2.5–1.25
1.25–0.75
0.5
present
epochs (kya)
rs12715075 (CNTN4) 825 ancient genotypes
convolution 20 ancient inds
convolution 100 ancient inds
pseudo haploid genotypes
rs4374563 (OSBPL10) 723 ancient genotypes
frequency
epochs (kya)
rs4374563 (OSBPL10) 723 ancient genotypes
convolution 20 ancient inds
convolution 100 ancient inds
pseudo haploid genotypes

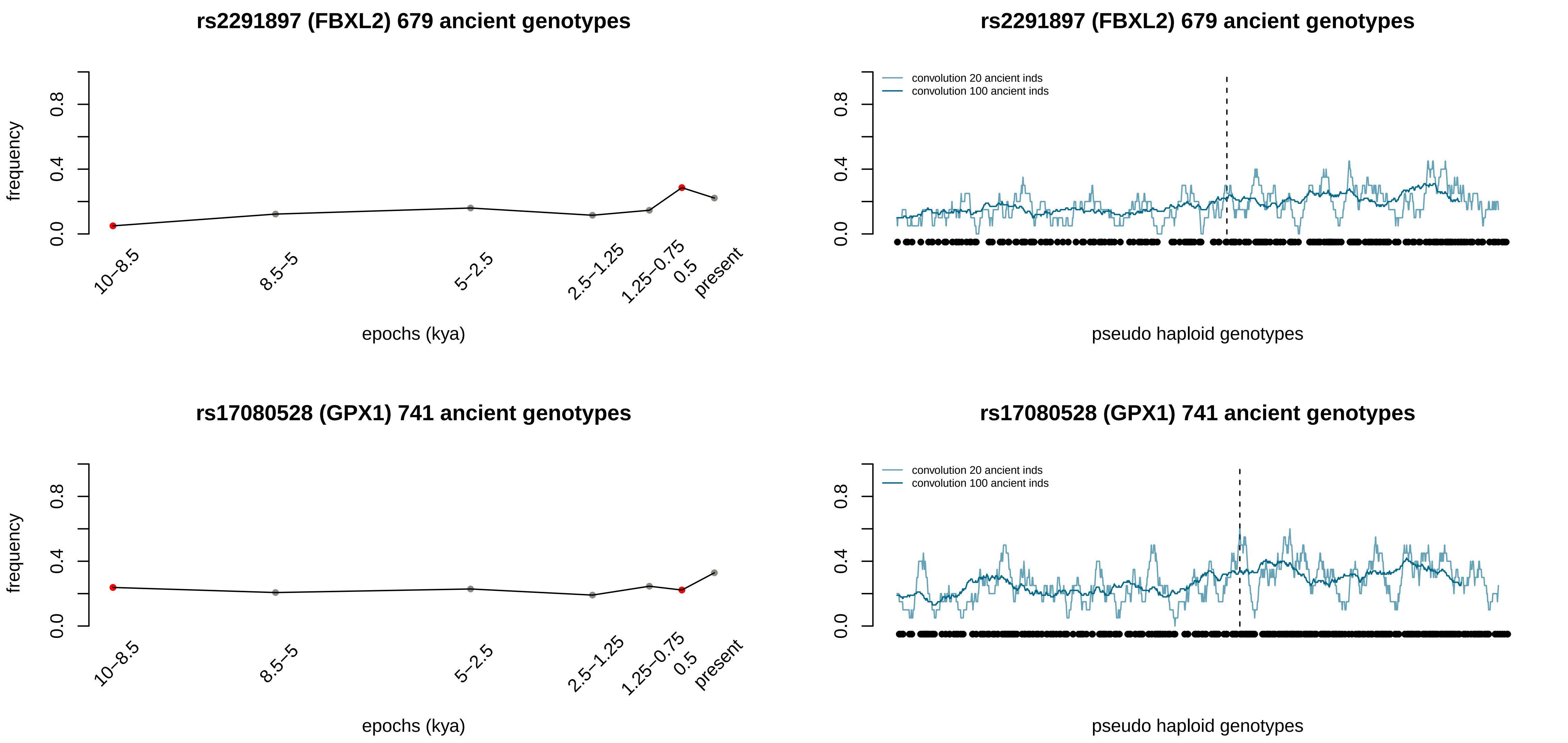
rs2291897 (FBXL2) 679 ancient genotypes
frequency
0.0
0.4
0.8
10–8.5
8.5–5
5–2.5
2.5–1.25
1.25–0.75
0.5
present
epochs (kya)
rs2291897 (FBXL2) 679 ancient genotypes
convolution 20 ancient inds
convolution 100 ancient inds
pseudo haploid genotypes
rs17080528 (GPX1) 741 ancient genotypes
frequency
epochs (kya)
rs17080528 (GPX1) 741 ancient genotypes
convolution 20 ancient inds
convolution 100 ancient inds
pseudo haploid genotypes

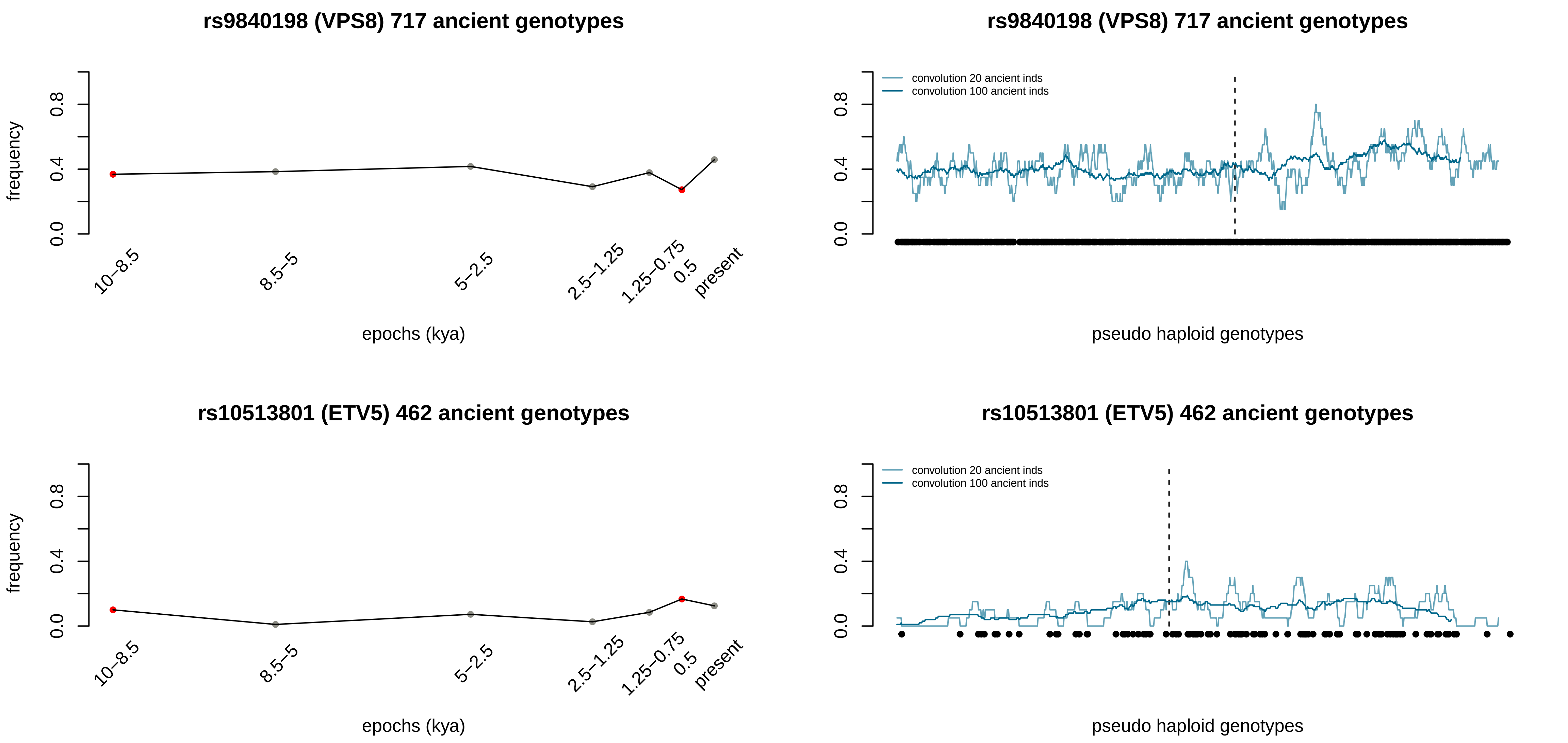
rs9840198 (VPS8) 717 ancient genotypes
frequency
0.0
0.4
0.8
10–8.5
8.5–5
5–2.5
2.5–1.25
1.25–0.75
0.5
present
epochs (kya)
rs9840198 (VPS8) 717 ancient genotypes
convolution 20 ancient inds
convolution 100 ancient inds
0.0
0.4
0.8
pseudo haploid genotypes
rs10513801 (ETV5) 462 ancient genotypes
frequency
0.0
0.4
0.8
10–8.5
8.5–5
5–2.5
2.5–1.25
1.25–0.75
0.5
present
epochs (kya)
rs10513801 (ETV5) 462 ancient genotypes
convolution 20 ancient inds
convolution 100 ancient inds
0.0
0.4
0.8
pseudo haploid genotypes

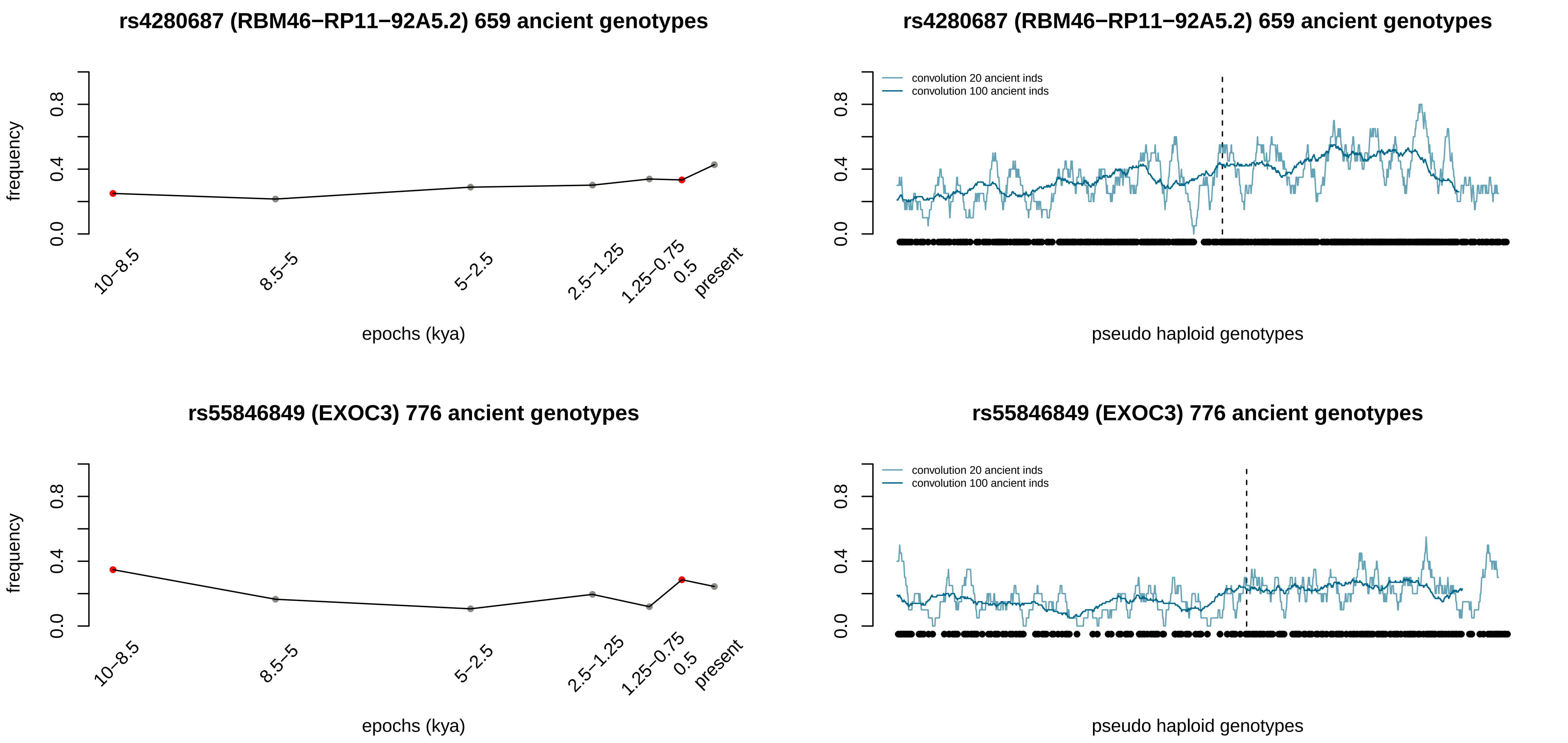

rs4280687 (RBM46−RP11−92A5.2) 659 ancient genotypes
frequency
0.0
0.4
0.8
10−8.5
8.5−5
5−2.5
2.5−1.25
1.25−0.75
0.5
present
epochs (kya)
rs4280687 (RBM46−RP11−92A5.2) 659 ancient genotypes
convolution 20 ancient inds
convolution 100 ancient inds
pseudo haploid genotypes
rs55846849 (EXOC3) 776 ancient genotypes
frequency
epochs (kya)
rs55846849 (EXOC3) 776 ancient genotypes
convolution 20 ancient inds
convolution 100 ancient inds
pseudo haploid genotypes

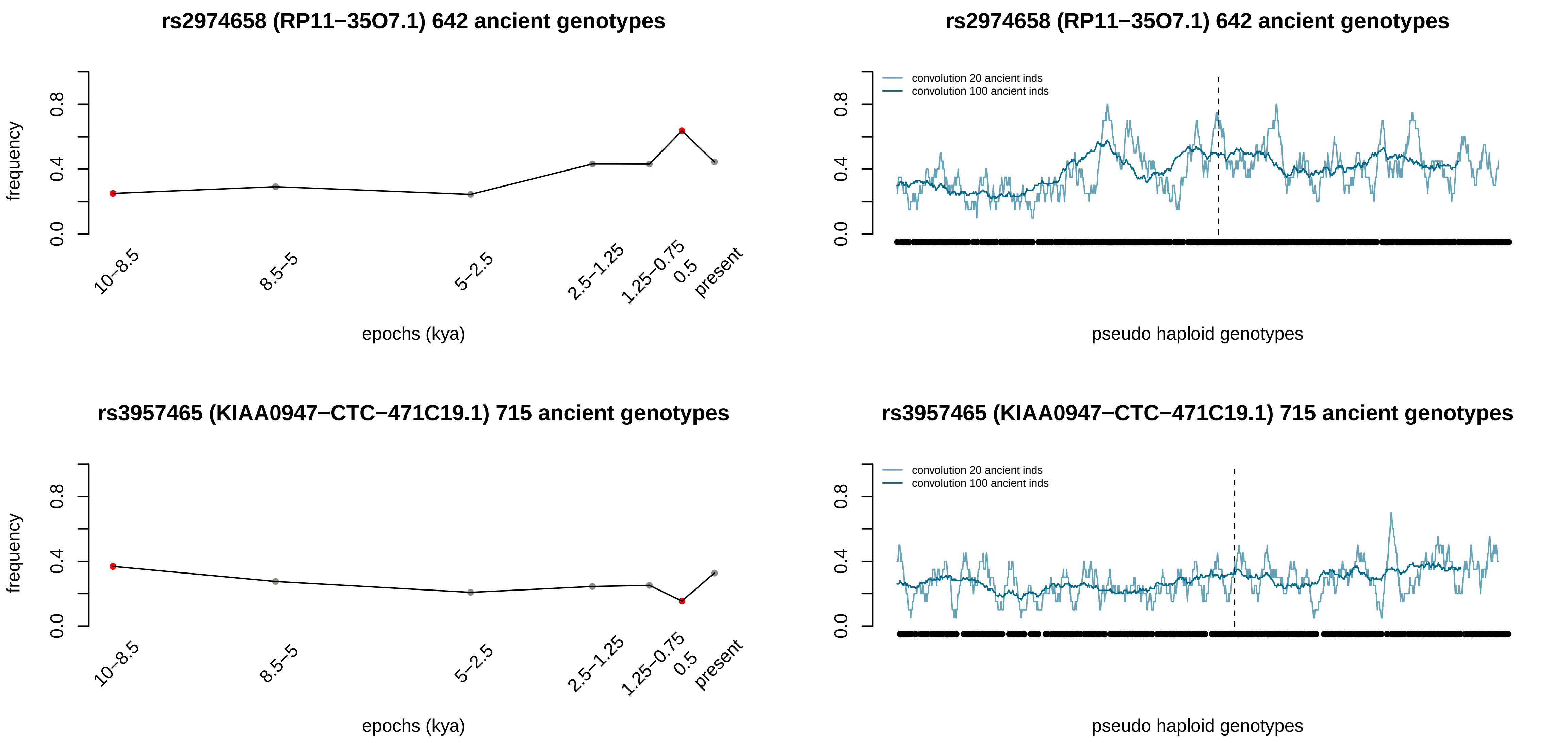
rs2974658 (RP11−35O7.1) 642 ancient genotypes
frequency
epochs (kya)
10−8.5
8.5−5
5−2.5
2.5−1.25
1.25−0.75
0.5
present
rs2974658 (RP11−35O7.1) 642 ancient genotypes
convolution 20 ancient inds
convolution 100 ancient inds
pseudo haploid genotypes
rs3957465 (KIAA0947−CTC−471C19.1) 715 ancient genotypes
frequency
epochs (kya)
rs3957465 (KIAA0947−CTC−471C19.1) 715 ancient genotypes
convolution 20 ancient inds
convolution 100 ancient inds
pseudo haploid genotypes

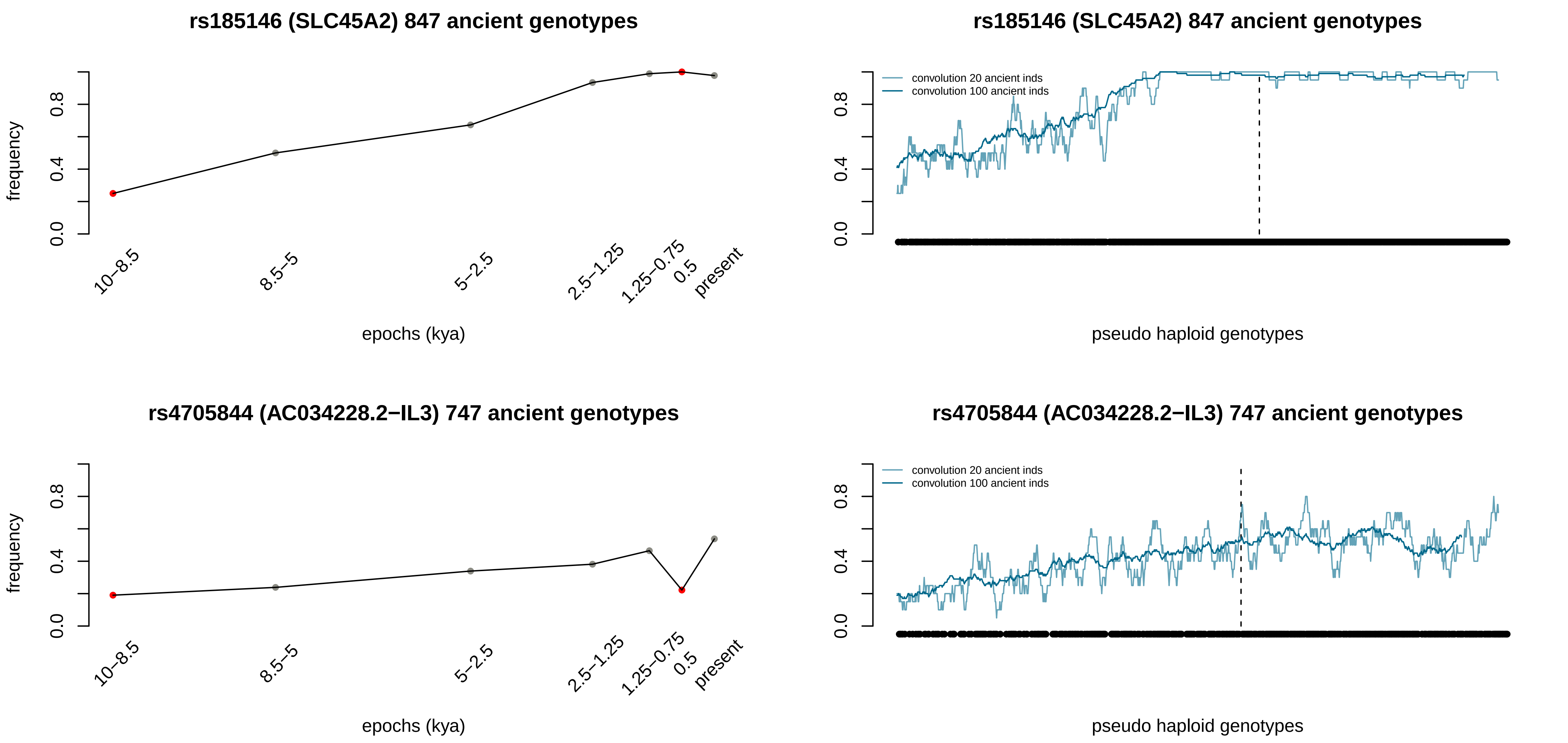

rs185146 (SLC45A2) 847 ancient genotypes
frequency
0.0
0.4
0.8
10–8.5
8.5–5
5–2.5
2.5–1.25
1.25–0.75
0.5
present
epochs (kya)
rs185146 (SLC45A2) 847 ancient genotypes
convolution 20 ancient inds
convolution 100 ancient inds
pseudo haploid genotypes
rs4705844 (AC034228.2−IL3) 747 ancient genotypes
frequency
epochs (kya)
rs4705844 (AC034228.2−IL3) 747 ancient genotypes
convolution 20 ancient inds
convolution 100 ancient inds
pseudo haploid genotypes

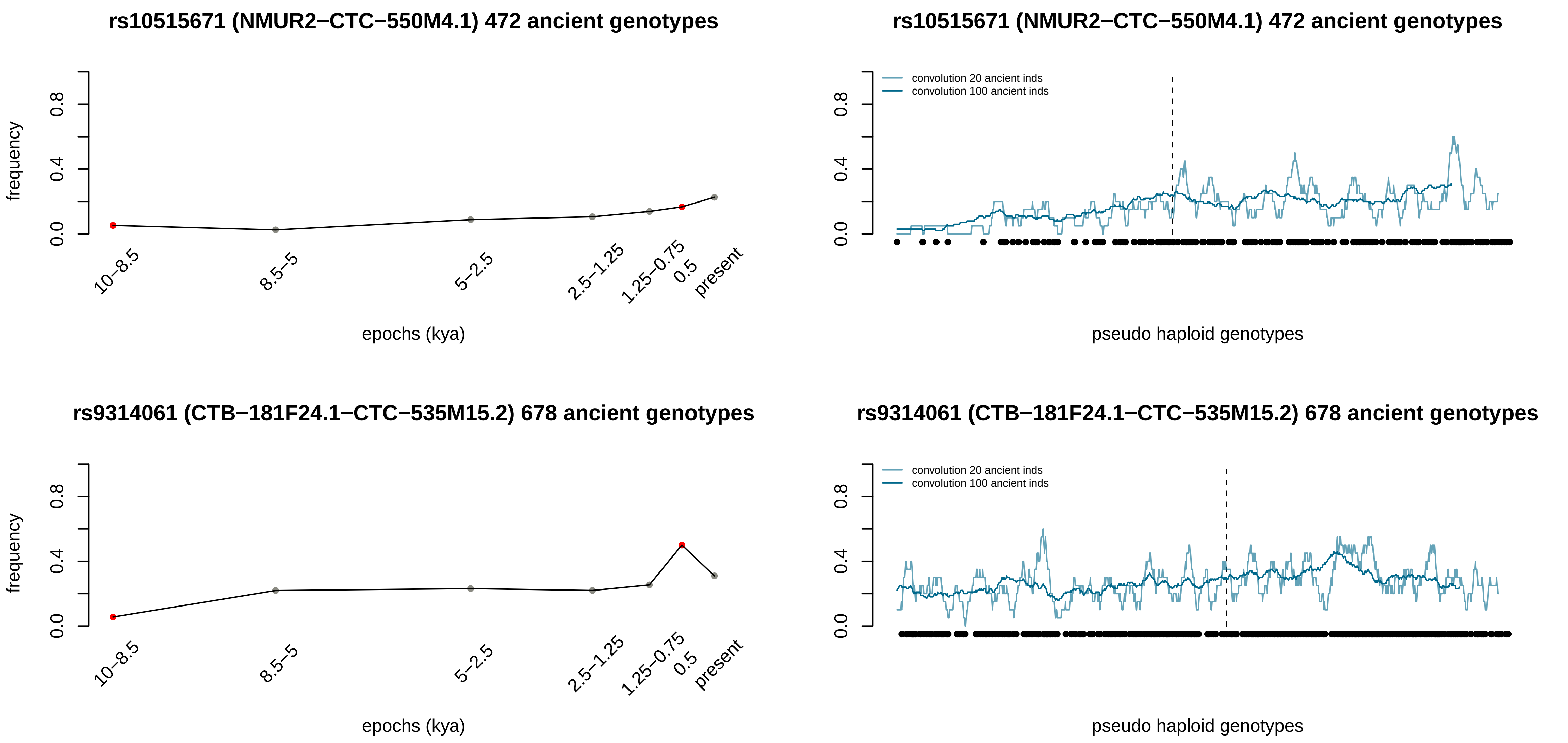
rs10515671 (NMUR2−CTC−550M4.1) 472 ancient genotypes
frequency
0.0
0.4
0.8
10−8.5
8.5−5
5−2.5
2.5−1.25
1.25−0.75
0.5
present
epochs (kya)
rs10515671 (NMUR2−CTC−550M4.1) 472 ancient genotypes
convolution 20 ancient inds
convolution 100 ancient inds
0.0
0.4
0.8
pseudo haploid genotypes
rs9314061 (CTB−181F24.1−CTC−535M15.2) 678 ancient genotypes
frequency
0.0
0.4
0.8
10−8.5
8.5−5
5−2.5
2.5−1.25
1.25−0.75
0.5
present
epochs (kya)
rs9314061 (CTB−181F24.1−CTC−535M15.2) 678 ancient genotypes
convolution 20 ancient inds
convolution 100 ancient inds
0.0
0.4
0.8
pseudo haploid genotypes

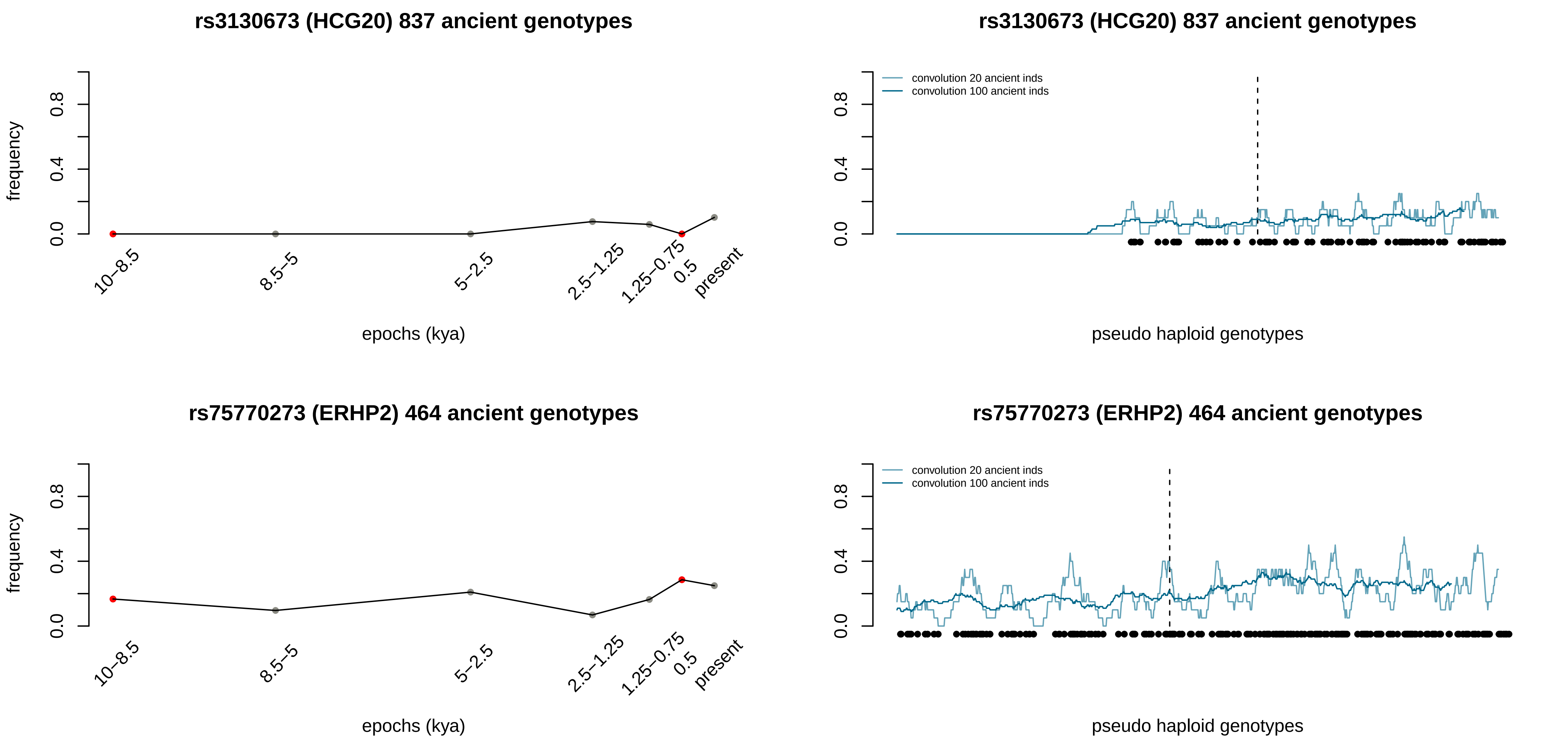

rs3130673 (HCG20) 837 ancient genotypes
frequency
epochs (kya)
10–8.5
8.5–5
5–2.5
2.5–1.25
1.25–0.75
0.5
present
rs3130673 (HCG20) 837 ancient genotypes
convolution 20 ancient inds
convolution 100 ancient inds
pseudo haploid genotypes
rs75770273 (ERHP2) 464 ancient genotypes
frequency
epochs (kya)
rs75770273 (ERHP2) 464 ancient genotypes
convolution 20 ancient inds
convolution 100 ancient inds
pseudo haploid genotypes

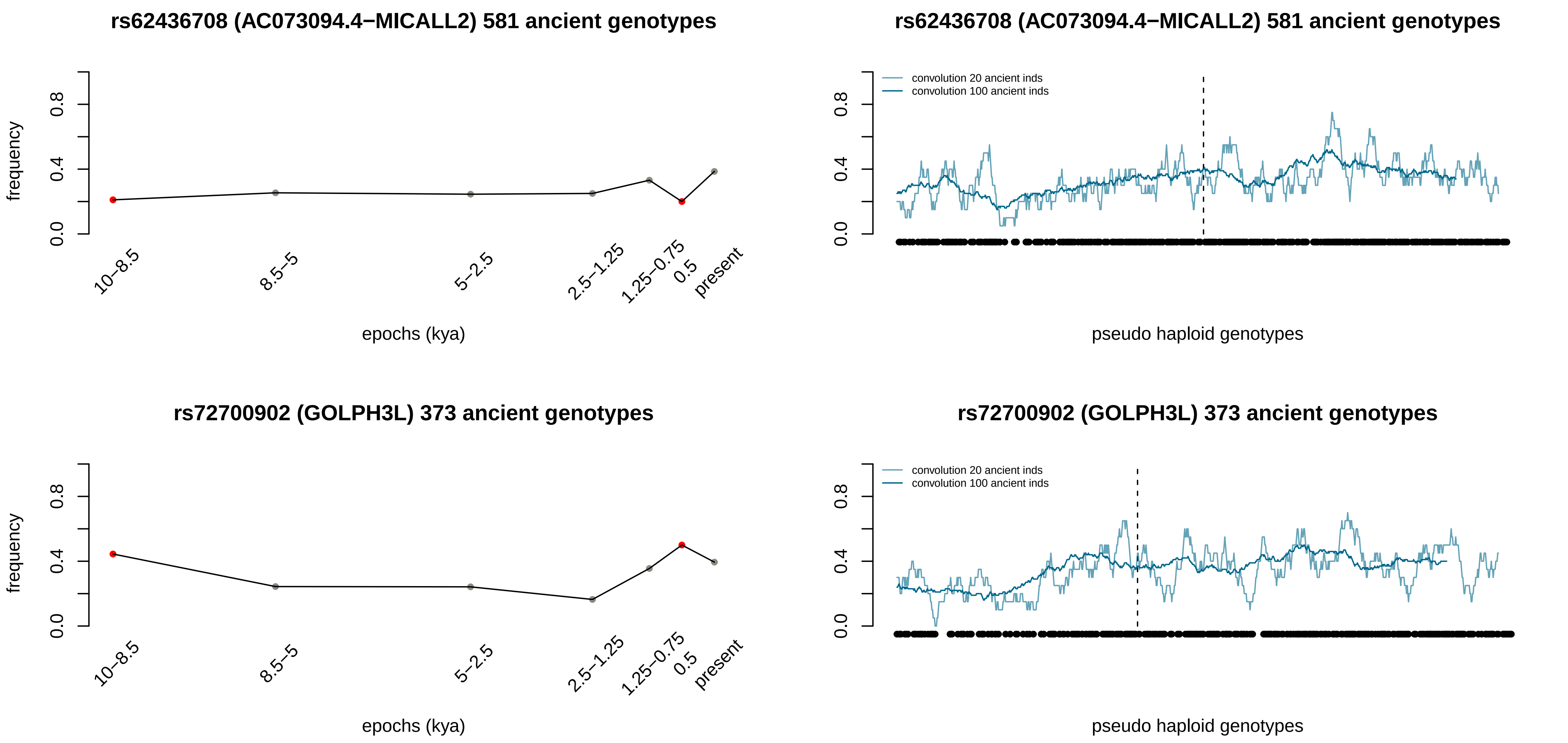

rs62436708 (AC073094.4−MICALL2) 581 ancient genotypes
frequency
0.0
0.4
0.8
10−8.5
8.5−5
5−2.5
2.5−1.25
1.25−0.75
0.5
present
epochs (kya)
rs62436708 (AC073094.4−MICALL2) 581 ancient genotypes
convolution 20 ancient inds
convolution 100 ancient inds
pseudo haploid genotypes
rs72700902 (GOLPH3L) 373 ancient genotypes
frequency
epochs (kya)
rs72700902 (GOLPH3L) 373 ancient genotypes
convolution 20 ancient inds
convolution 100 ancient inds
pseudo haploid genotypes

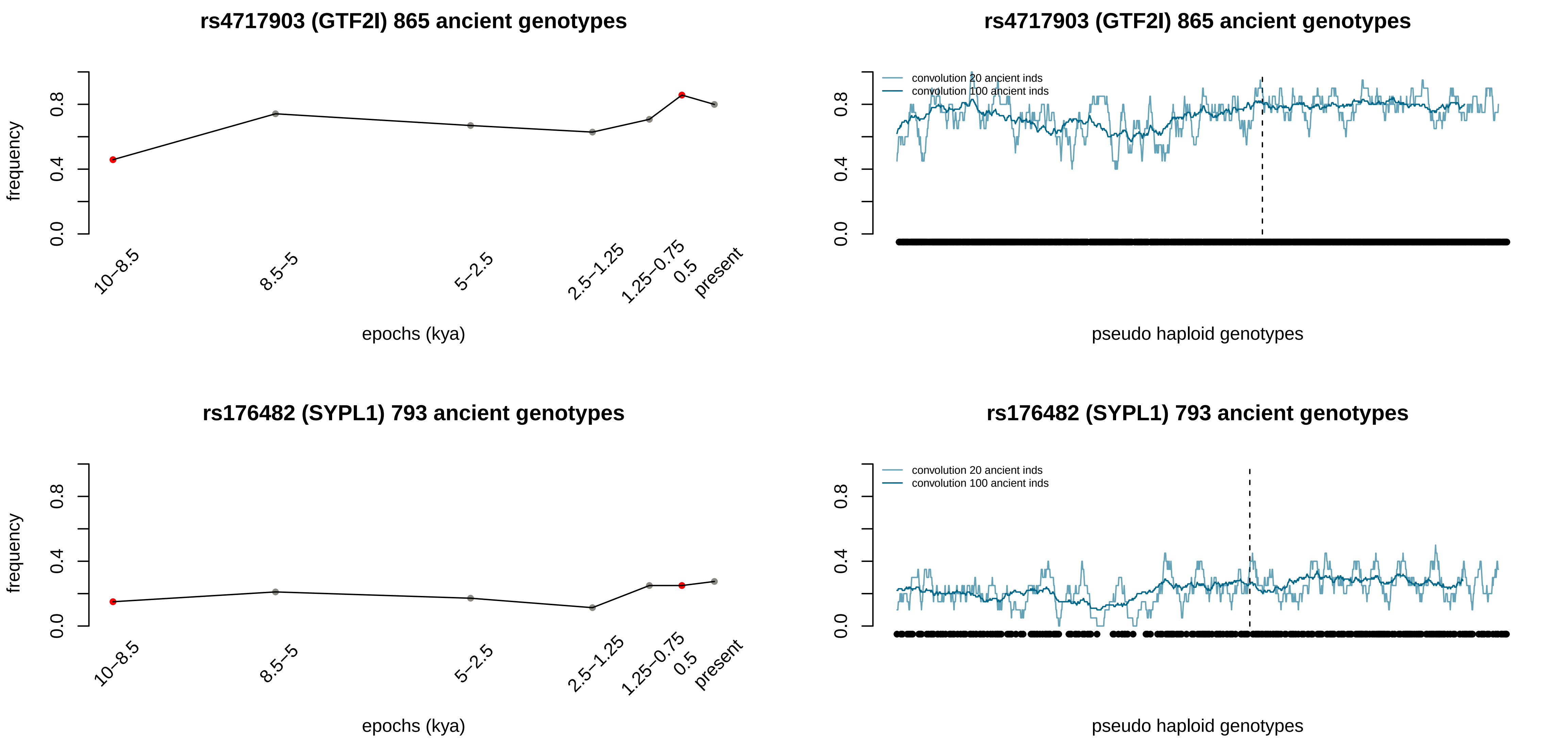
rs4717903 (GTF2I) 865 ancient genotypes
frequency
epochs (kya)
10–8.5
8.5–5
5–2.5
2.5–1.25
1.25–0.75
0.5
present
rs4717903 (GTF2I) 865 ancient genotypes
convolution 20 ancient inds
convolution 100 ancient inds
pseudo haploid genotypes
rs176482 (SYPL1) 793 ancient genotypes
frequency
epochs (kya)
rs176482 (SYPL1) 793 ancient genotypes
convolution 20 ancient inds
convolution 100 ancient inds
pseudo haploid genotypes

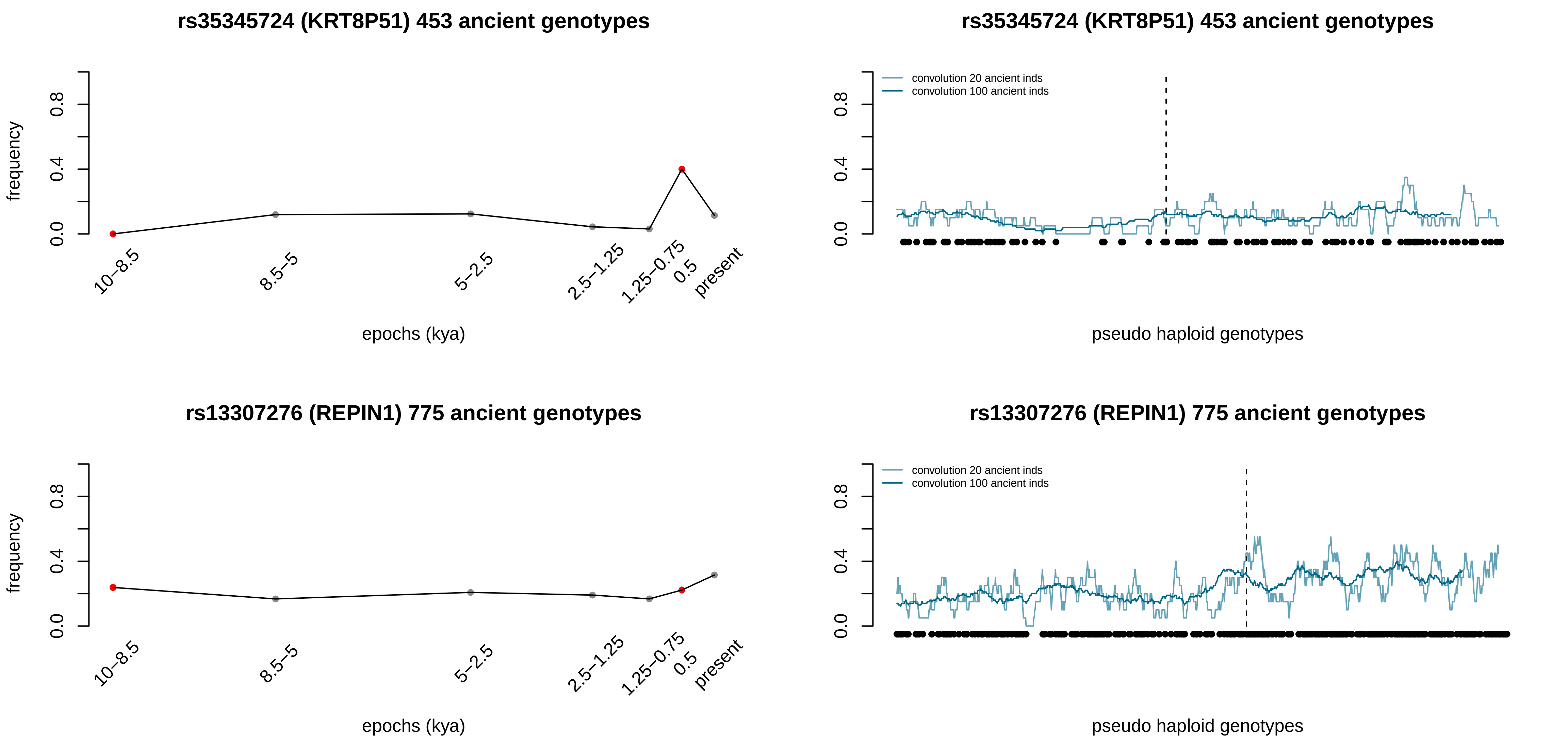
rs35345724 (KRT8P51) 453 ancient genotypes
frequency
0.0
0.4
0.8
10–8.5
8.5–5
5–2.5
2.5–1.25
1.25–0.75
0.5
present
epochs (kya)
rs35345724 (KRT8P51) 453 ancient genotypes
convolution 20 ancient inds
convolution 100 ancient inds
pseudo haploid genotypes
rs13307276 (REPIN1) 775 ancient genotypes
frequency
epochs (kya)
rs13307276 (REPIN1) 775 ancient genotypes
convolution 20 ancient inds
convolution 100 ancient inds
pseudo haploid genotypes

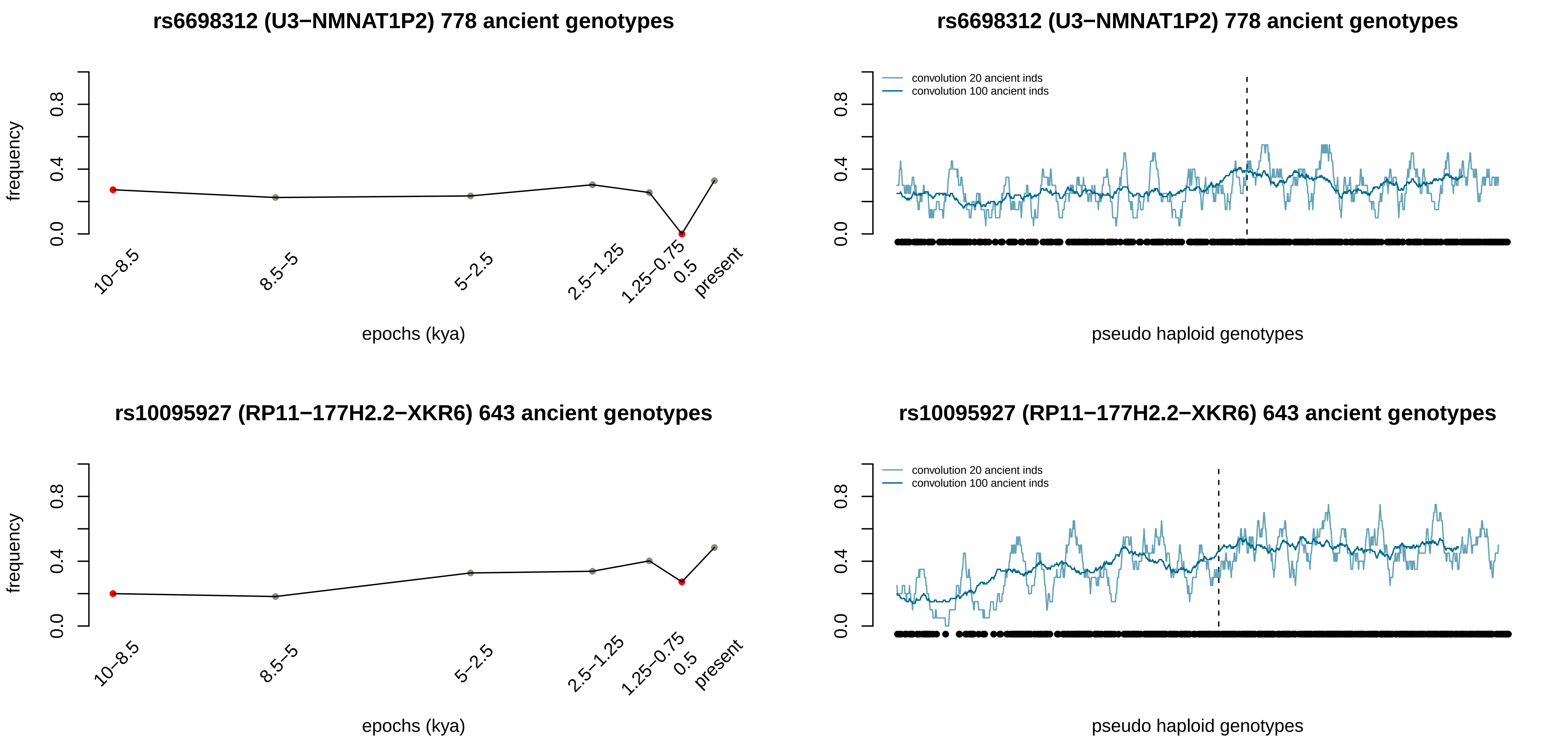
rs6698312 (U3−NMNAT1P2) 778 ancient genotypes
frequency
0.0
0.4
0.8
10−8.5
8.5−5
5−2.5
2.5−1.25
1.25−0.75
0.5
present
epochs (kya)
rs6698312 (U3−NMNAT1P2) 778 ancient genotypes
convolution 20 ancient inds
convolution 100 ancient inds
0.0
0.4
0.8
pseudo haploid genotypes
rs10095927 (RP11−177H2.2−XKR6) 643 ancient genotypes
frequency
0.0
0.4
0.8
10−8.5
8.5−5
5−2.5
2.5−1.25
1.25−0.75
0.5
present
epochs (kya)
rs10095927 (RP11−177H2.2−XKR6) 643 ancient genotypes
convolution 20 ancient inds
convolution 100 ancient inds
0.0
0.4
0.8
pseudo haploid genotypes

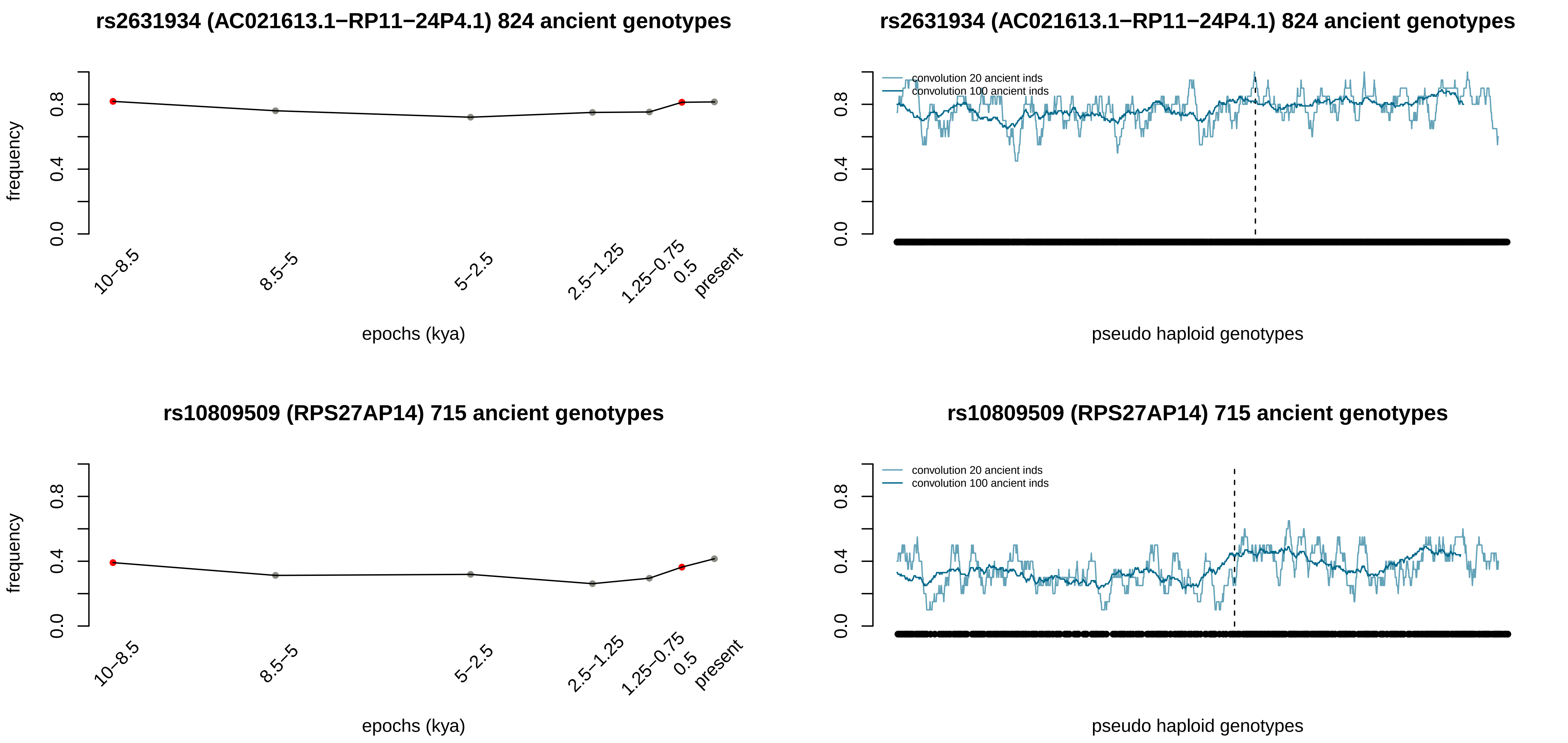

rs2631934 (AC021613.1−RP11−24P4.1) 824 ancient genotypes
frequency
0.0
0.4
0.8
10−8.5
8.5−5
5−2.5
2.5−1.25
1.25−0.75
0.5
present
epochs (kya)
rs2631934 (AC021613.1−RP11−24P4.1) 824 ancient genotypes
convolution 20 ancient inds
convolution 100 ancient inds
pseudo haploid genotypes
rs10809509 (RPS27AP14) 715 ancient genotypes
frequency
epochs (kya)
rs10809509 (RPS27AP14) 715 ancient genotypes
convolution 20 ancient inds
convolution 100 ancient inds
pseudo haploid genotypes

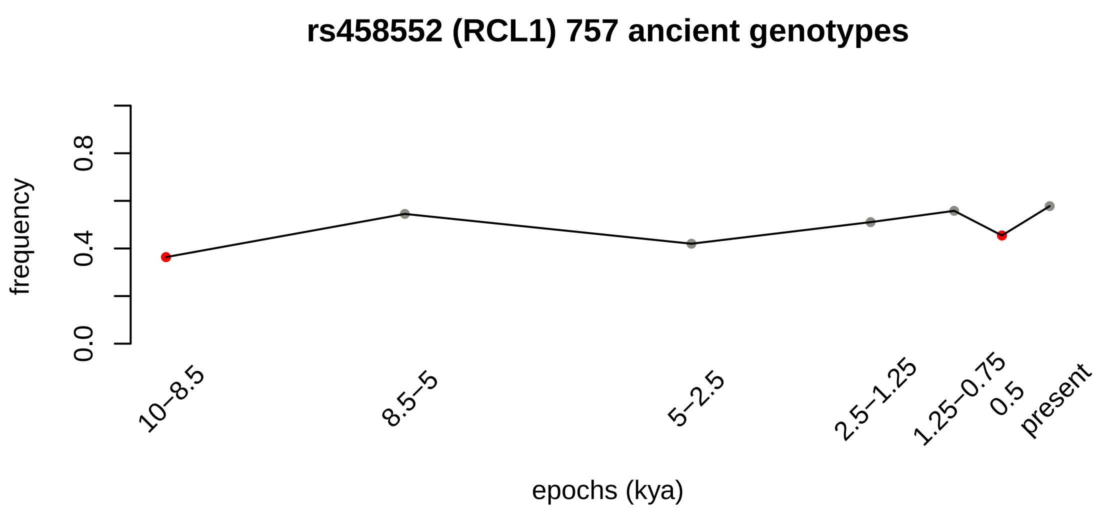
rs458552 (RCL1) 757 ancient genotypes
frequency
0.0
0.4
0.8
10–8.5
8.5–5
5–2.5
2.5–1.25
1.25–0.75
0.5
present
epochs (kya)

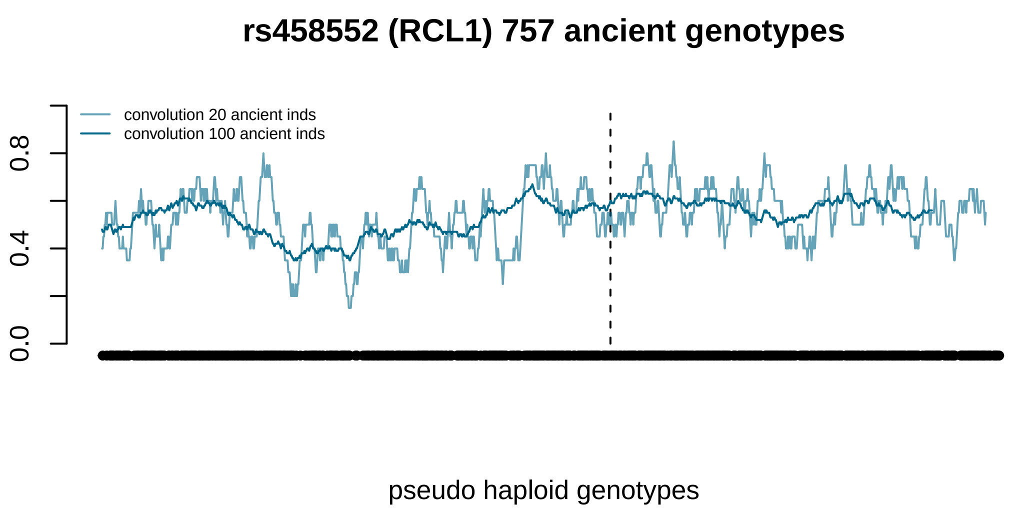
rs458552 (RCL1) 757 ancient genotypes
convolution 20 ancient inds
convolution 100 ancient inds
0.0
0.4
0.8
pseudo haploid genotypes

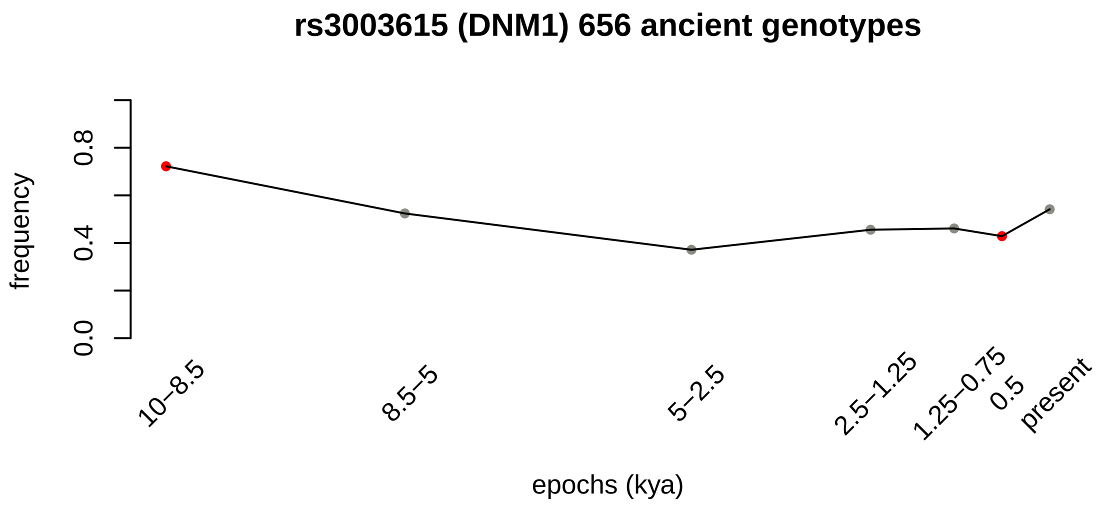
rs3003615 (DNM1) 656 ancient genotypes
frequency
0.0
0.4
0.8
10–8.5
8.5–5
5–2.5
2.5–1.25
1.25–0.75
0.5
present
epochs (kya)

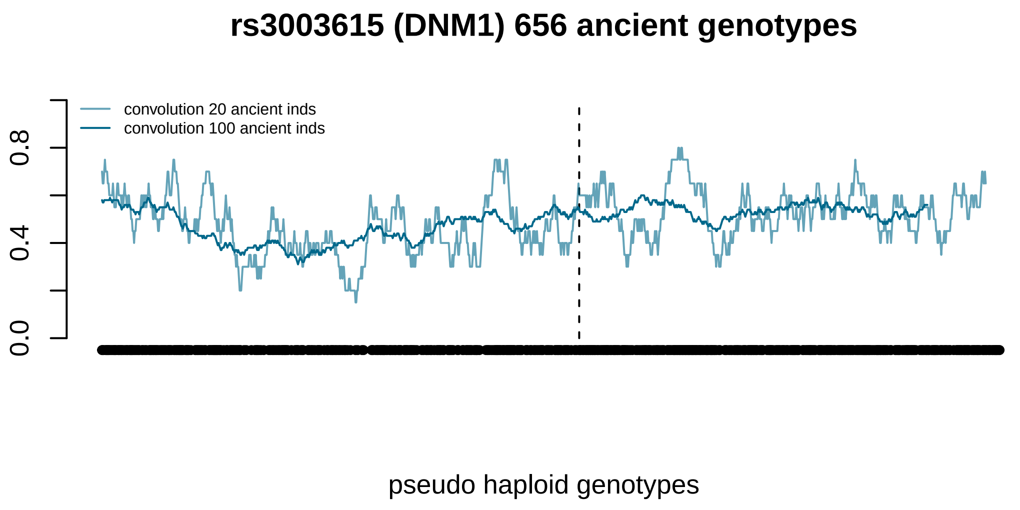
rs3003615 (DNM1) 656 ancient genotypes
convolution 20 ancient inds
convolution 100 ancient inds
0.0
0.4
0.8
pseudo haploid genotypes

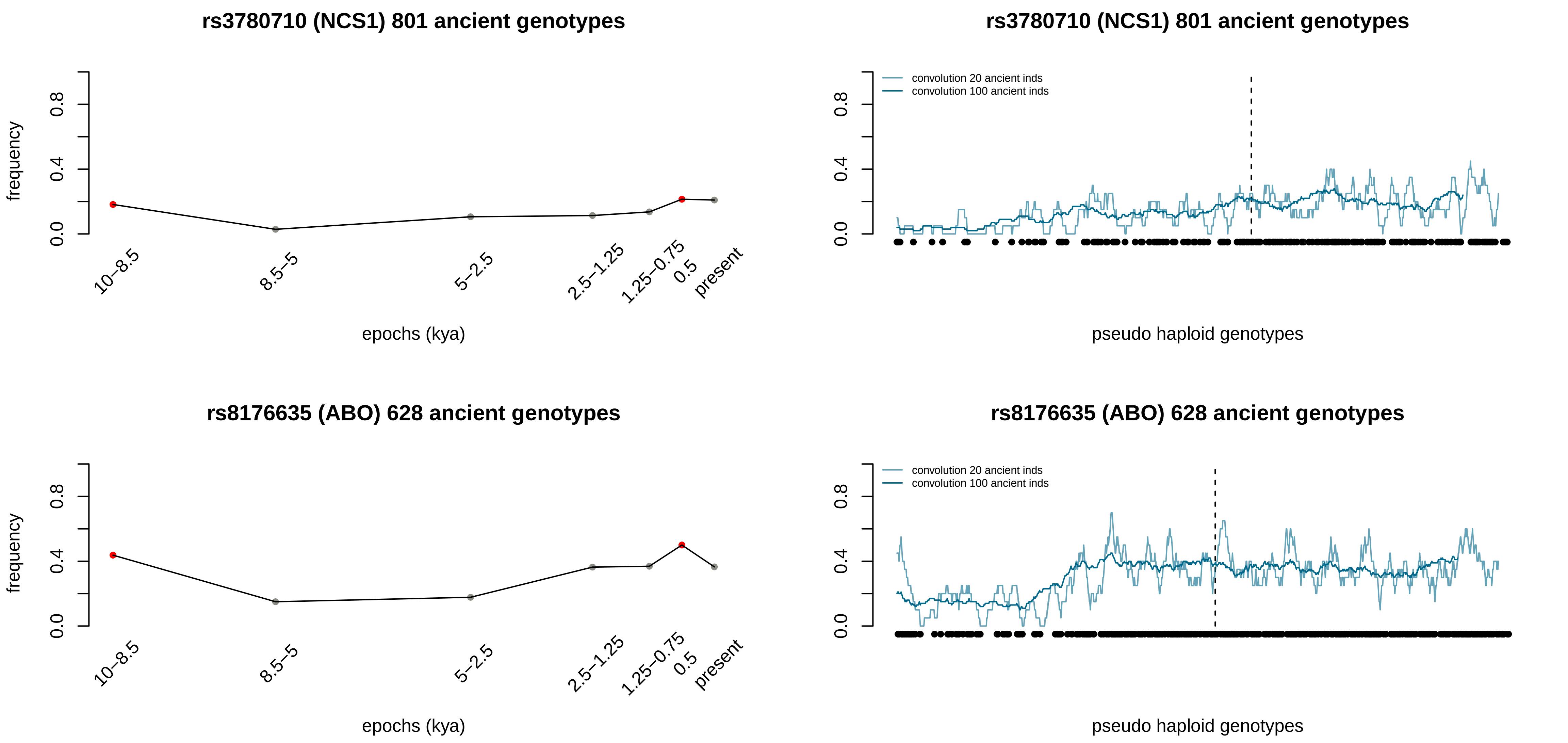

rs3780710 (NCS1) 801 ancient genotypes
frequency
0.0
0.4
0.8
10–8.5
8.5–5
5–2.5
2.5–1.25
1.25–0.75
0.5
present
epochs (kya)
rs3780710 (NCS1) 801 ancient genotypes
convolution 20 ancient inds
convolution 100 ancient inds
pseudo haploid genotypes
rs8176635 (ABO) 628 ancient genotypes
frequency
epochs (kya)
rs8176635 (ABO) 628 ancient genotypes
convolution 20 ancient inds
convolution 100 ancient inds
pseudo haploid genotypes

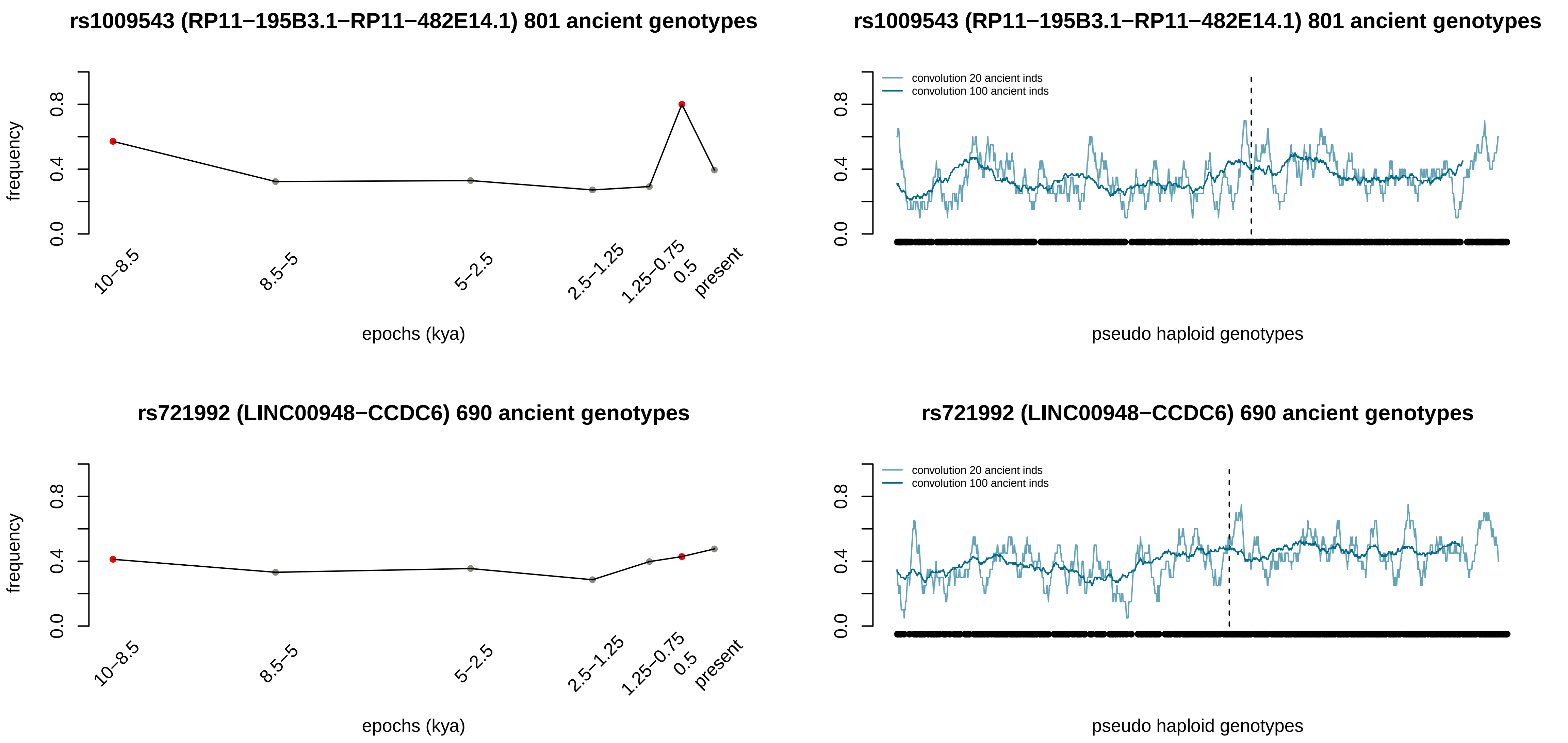

rs1009543 (RP11−195B3.1−RP11−482E14.1) 801 ancient genotypes
frequency
0.0
0.4
0.8
10−8.5
8.5−5
5−2.5
2.5−1.25
1.25−0.75
0.5
present
epochs (kya)
rs1009543 (RP11−195B3.1−RP11−482E14.1) 801 ancient genotypes
convolution 20 ancient inds
convolution 100 ancient inds
pseudo haploid genotypes
rs721992 (LINC00948−CCDC6) 690 ancient genotypes
frequency
epochs (kya)
rs721992 (LINC00948−CCDC6) 690 ancient genotypes
convolution 20 ancient inds
convolution 100 ancient inds
pseudo haploid genotypes

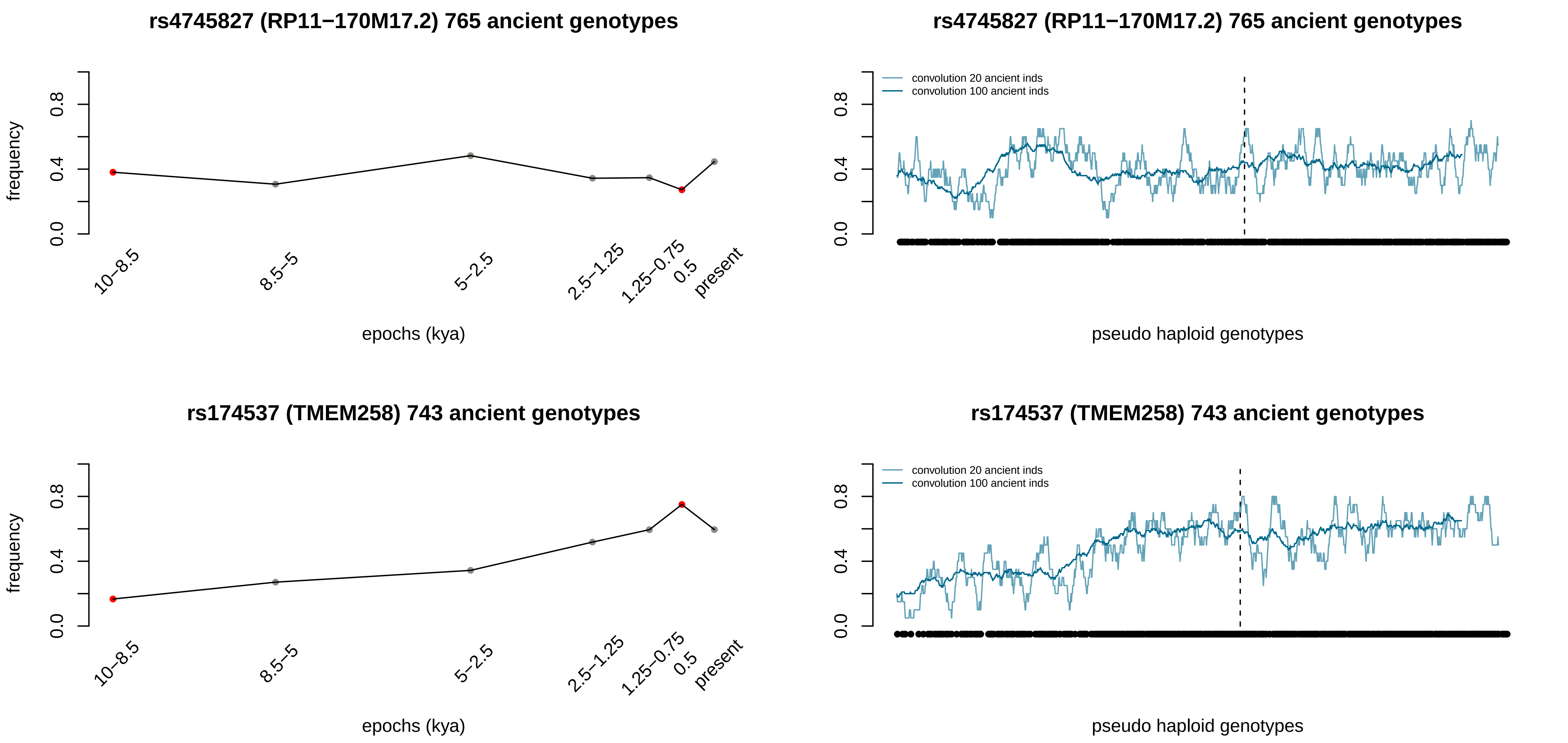
rs4745827 (RP11−170M17.2) 765 ancient genotypes
frequency
0.0
0.4
0.8
10–8.5
8.5–5
5–2.5
2.5–1.25
1.25–0.75
0.5
present
epochs (kya)
rs4745827 (RP11−170M17.2) 765 ancient genotypes
convolution 20 ancient inds
convolution 100 ancient inds
pseudo haploid genotypes
rs174537 (TMEM258) 743 ancient genotypes
frequency
epochs (kya)
rs174537 (TMEM258) 743 ancient genotypes
convolution 20 ancient inds
convolution 100 ancient inds
pseudo haploid genotypes

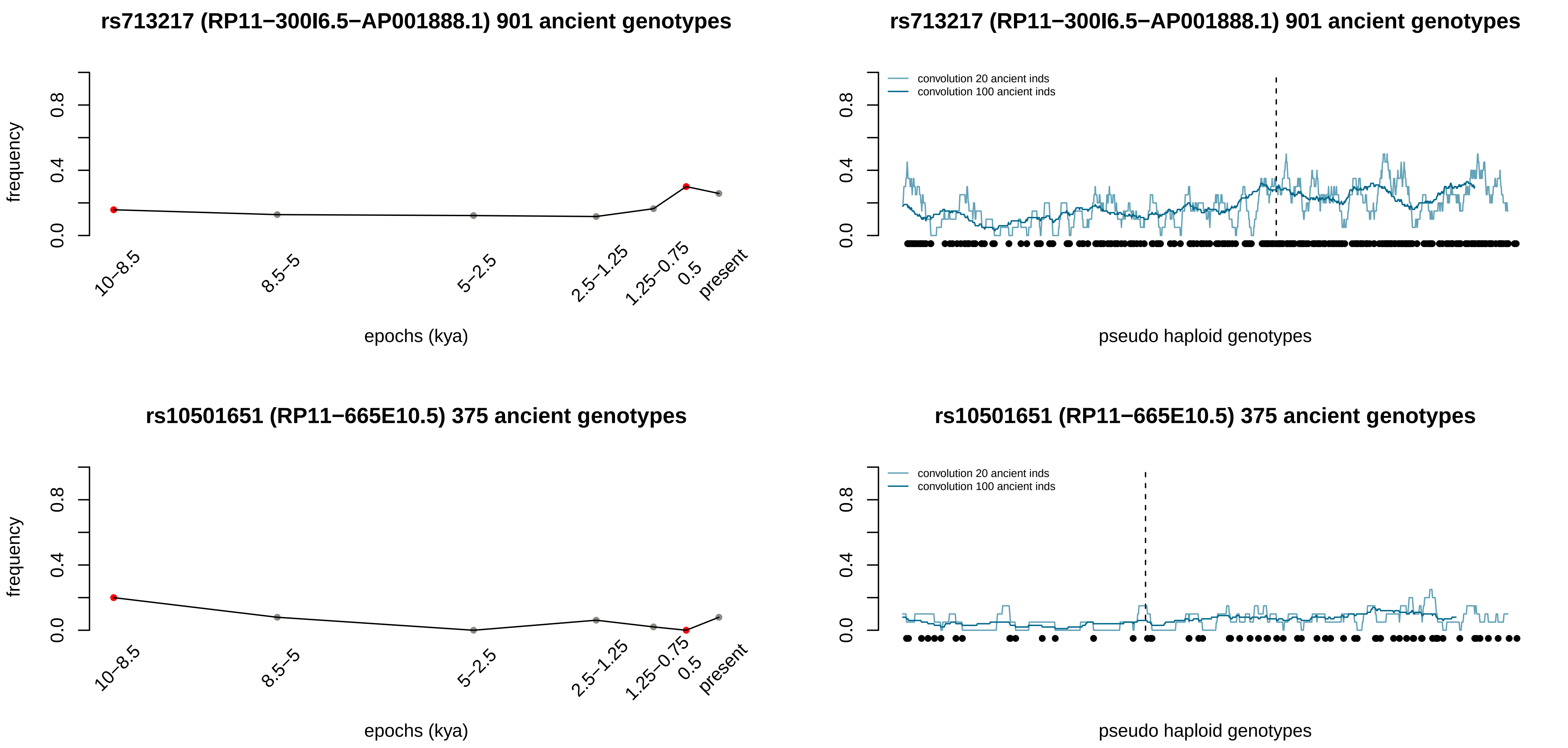
rs713217 (RP11−300I6.5−AP001888.1) 901 ancient genotypes
frequency
0.0
0.4
0.8
10−8.5
8.5−5
5−2.5
2.5−1.25
1.25−0.75
0.5
present
epochs (kya)
rs713217 (RP11−300I6.5−AP001888.1) 901 ancient genotypes
convolution 20 ancient inds
convolution 100 ancient inds
pseudo haploid genotypes
rs10501651 (RP11−665E10.5) 375 ancient genotypes
frequency
epochs (kya)
rs10501651 (RP11−665E10.5) 375 ancient genotypes
convolution 20 ancient inds
convolution 100 ancient inds
pseudo haploid genotypes

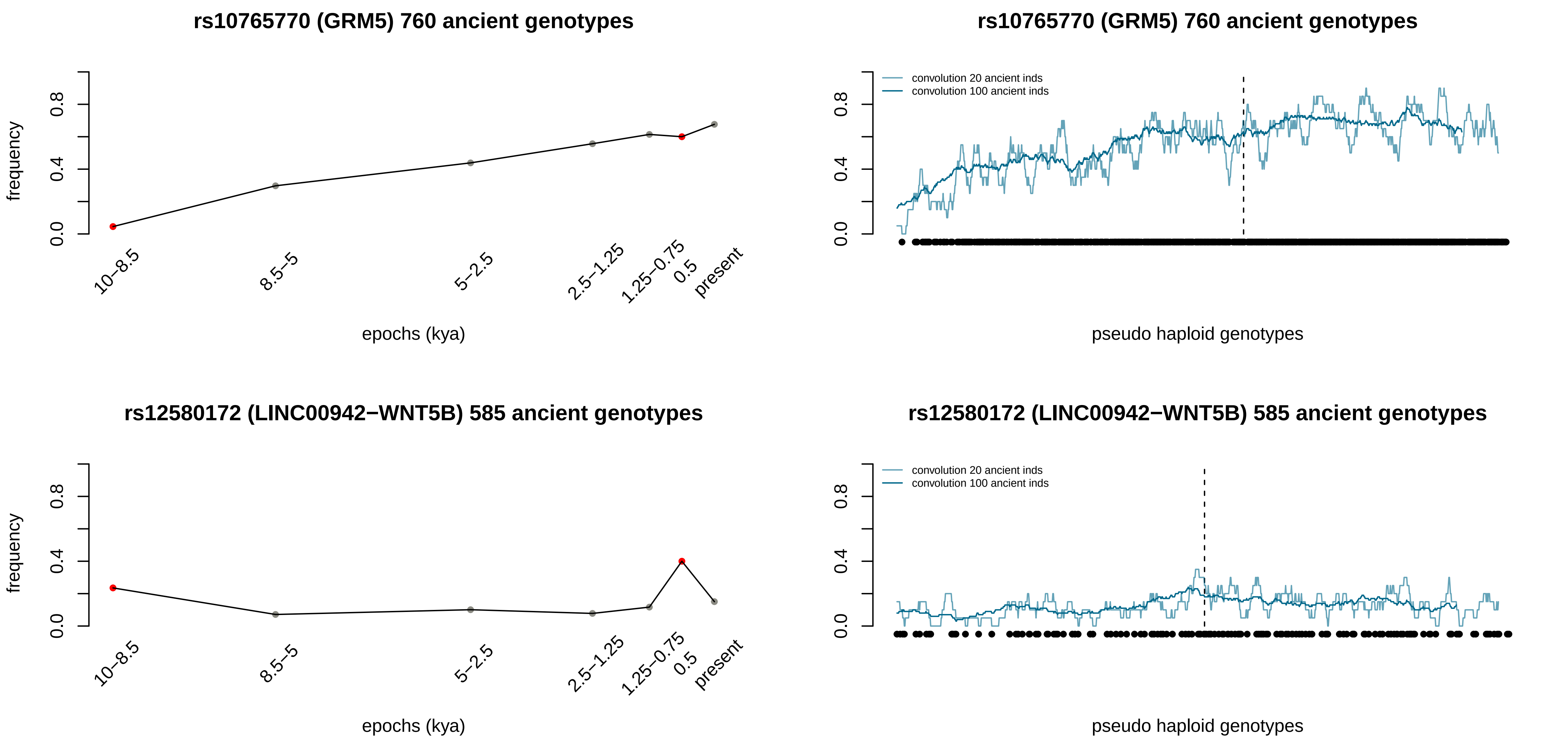
rs10765770 (GRM5) 760 ancient genotypes
frequency
0.0
0.4
0.8
10–8.5
8.5–5
5–2.5
2.5–1.25
1.25–0.75
0.5
present
epochs (kya)
rs10765770 (GRM5) 760 ancient genotypes
convolution 20 ancient inds
convolution 100 ancient inds
0.0
0.4
0.8
pseudo haploid genotypes
rs12580172 (LINC00942−WNT5B) 585 ancient genotypes
frequency
0.0
0.4
0.8
10–8.5
8.5–5
5–2.5
2.5–1.25
1.25–0.75
0.5
present
epochs (kya)
rs12580172 (LINC00942−WNT5B) 585 ancient genotypes
convolution 20 ancient inds
convolution 100 ancient inds
0.0
0.4
0.8
pseudo haploid genotypes

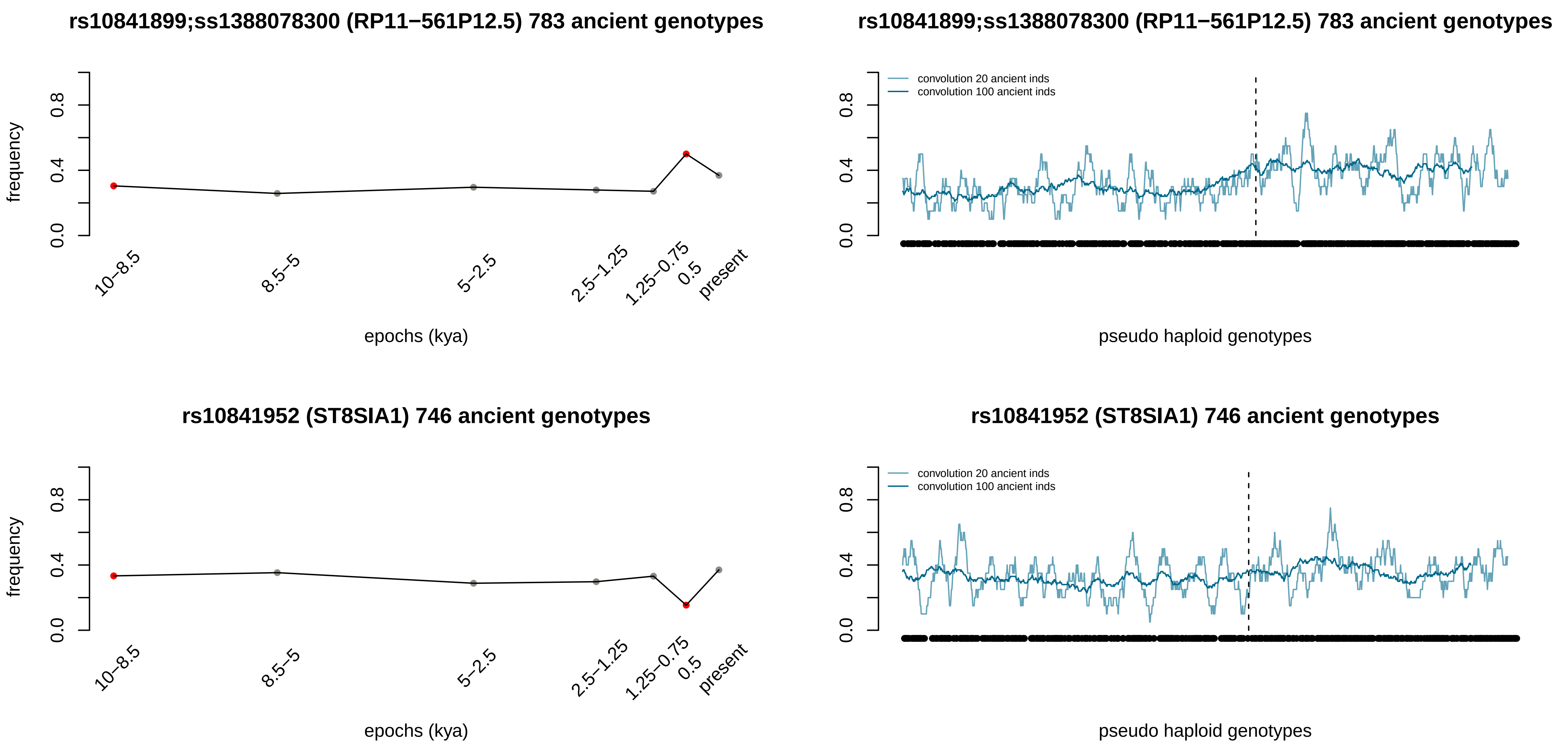
rs10841899;ss1388078300 (RP11−561P12.5) 783 ancient genotypes
frequency
0.0
0.4
0.8
10−8.5
8.5−5
5−2.5
2.5−1.25
1.25−0.75
0.5
present
epochs (kya)
rs10841899;ss1388078300 (RP11−561P12.5) 783 ancient genotypes
convolution 20 ancient inds
convolution 100 ancient inds
0.0
0.4
0.8
pseudo haploid genotypes
rs10841952 (ST8SIA1) 746 ancient genotypes
frequency
0.0
0.4
0.8
10−8.5
8.5−5
5−2.5
2.5−1.25
1.25−0.75
0.5
present
epochs (kya)
rs10841952 (ST8SIA1) 746 ancient genotypes
convolution 20 ancient inds
convolution 100 ancient inds
0.0
0.4
0.8
pseudo haploid genotypes

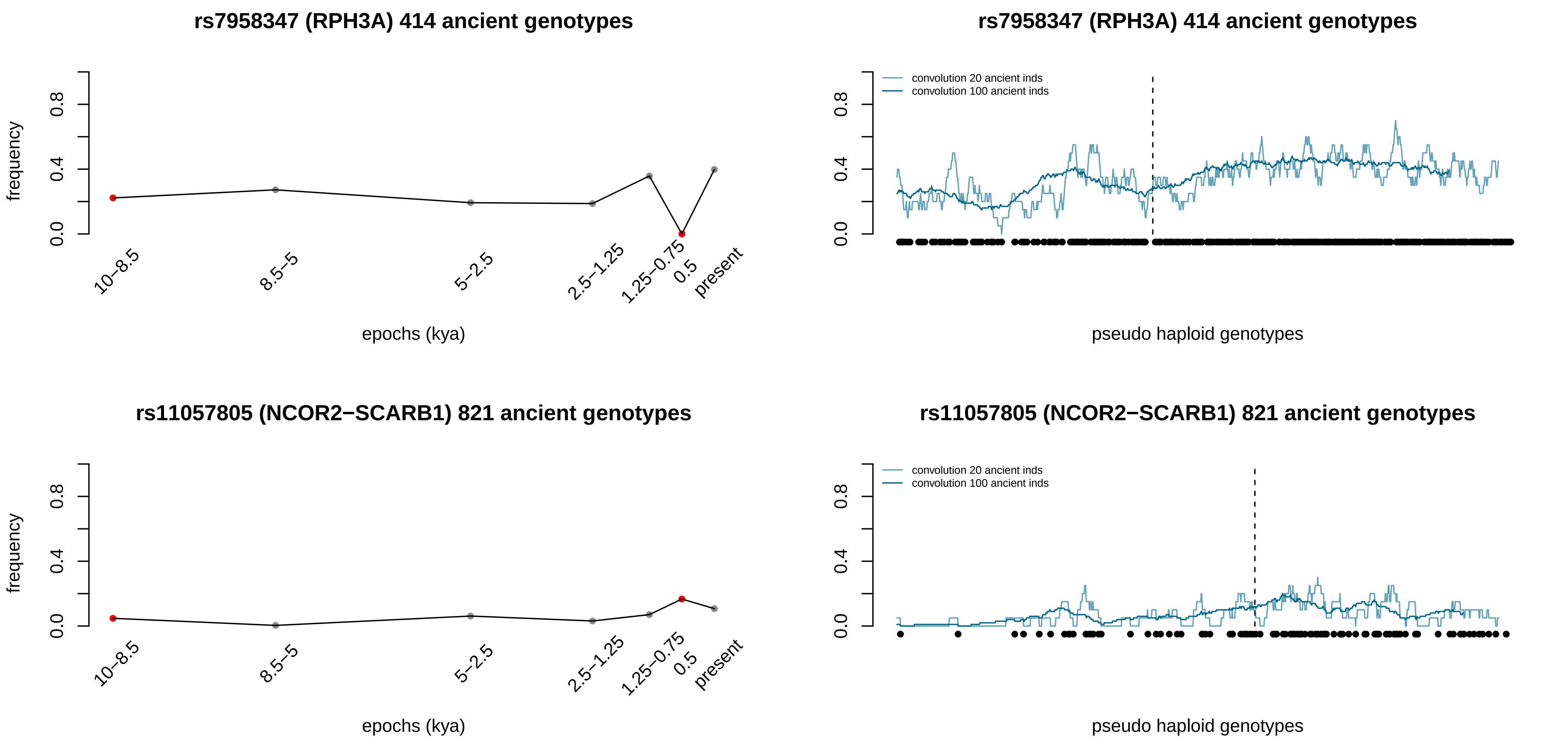
rs7958347 (RPH3A) 414 ancient genotypes
frequency
0.0
0.4
0.8
10–8.5
8.5–5
5–2.5
2.5–1.25
1.25–0.75
0.5
present
epochs (kya)
rs7958347 (RPH3A) 414 ancient genotypes
convolution 20 ancient inds
convolution 100 ancient inds
pseudo haploid genotypes
rs11057805 (NCOR2−SCARB1) 821 ancient genotypes
frequency
epochs (kya)
rs11057805 (NCOR2−SCARB1) 821 ancient genotypes
convolution 20 ancient inds
convolution 100 ancient inds
pseudo haploid genotypes

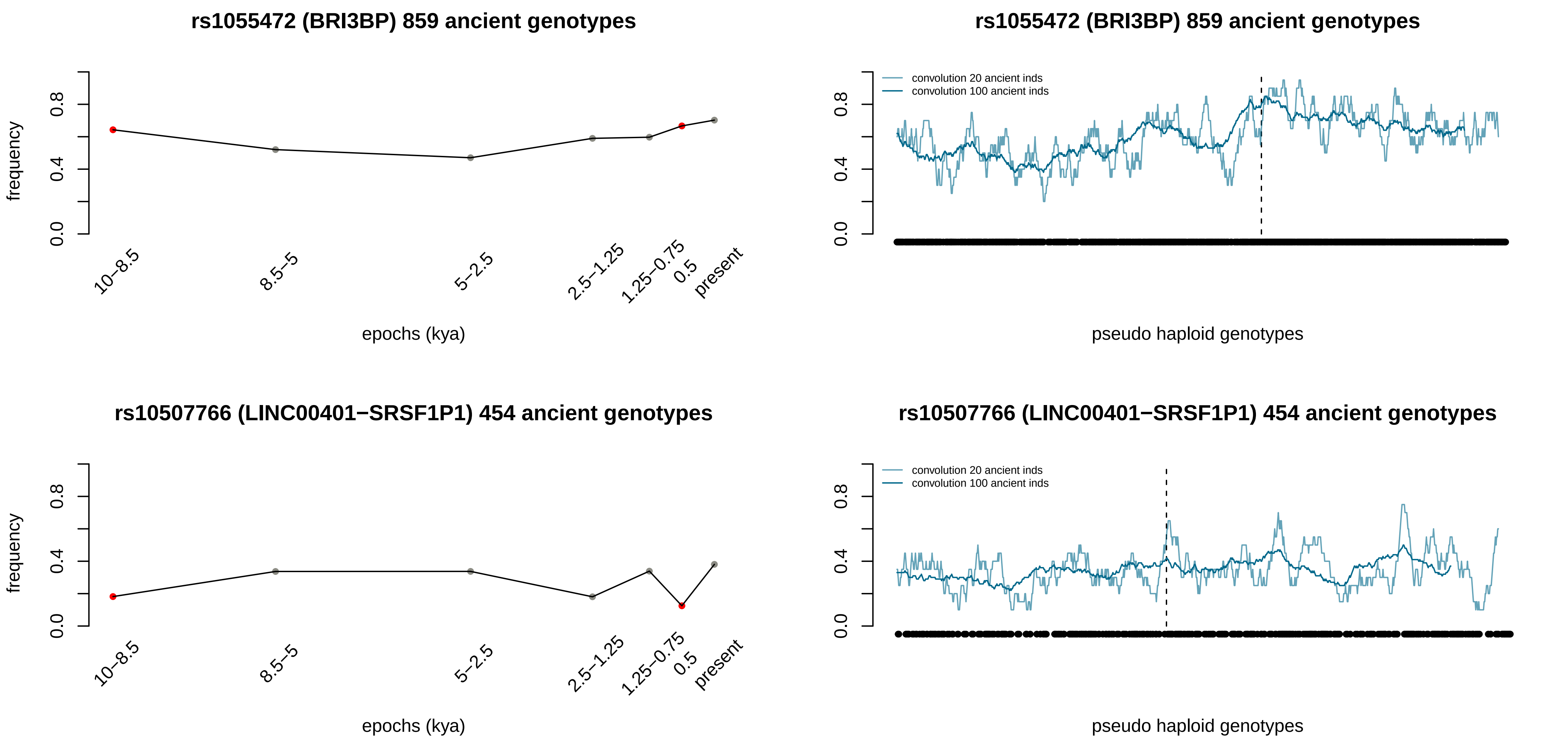
rs1055472 (BRI3BP) 859 ancient genotypes
frequency
0.0
0.4
0.8
10–8.5
8.5–5
5–2.5
2.5–1.25
1.25–0.75
0.5
present
epochs (kya)
rs1055472 (BRI3BP) 859 ancient genotypes
convolution 20 ancient inds
convolution 100 ancient inds
pseudo haploid genotypes
rs10507766 (LINC00401−SRSF1P1) 454 ancient genotypes
frequency
epochs (kya)
rs10507766 (LINC00401−SRSF1P1) 454 ancient genotypes
convolution 20 ancient inds
convolution 100 ancient inds
pseudo haploid genotypes

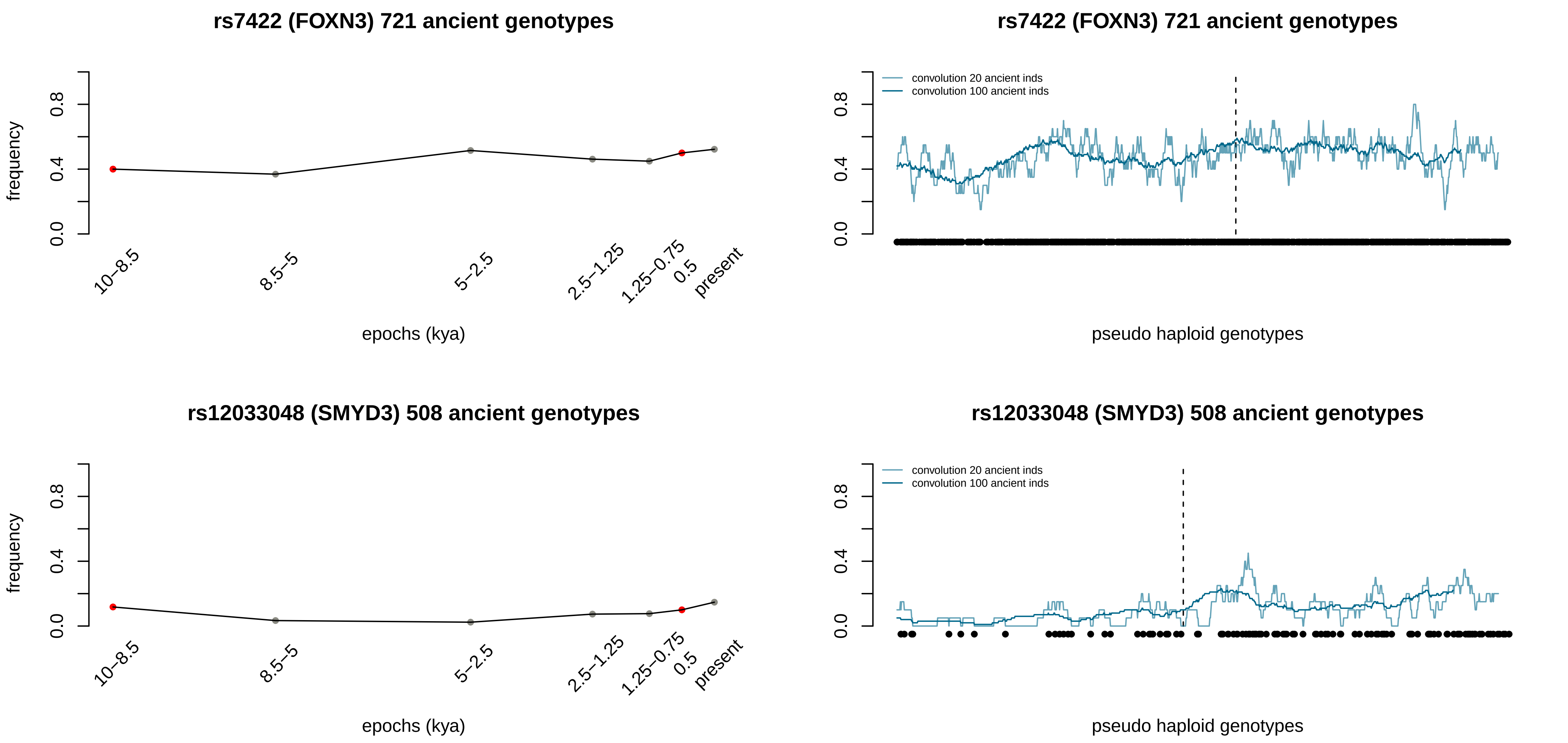
rs7422 (FOXN3) 721 ancient genotypes
frequency
0.0
0.4
0.8
10–8.5
8.5–5
5–2.5
2.5–1.25
1.25–0.75
0.5
present
epochs (kya)
rs7422 (FOXN3) 721 ancient genotypes
convolution 20 ancient inds
convolution 100 ancient inds
pseudo haploid genotypes
rs12033048 (SMYD3) 508 ancient genotypes
frequency
epochs (kya)
rs12033048 (SMYD3) 508 ancient genotypes
convolution 20 ancient inds
convolution 100 ancient inds
pseudo haploid genotypes

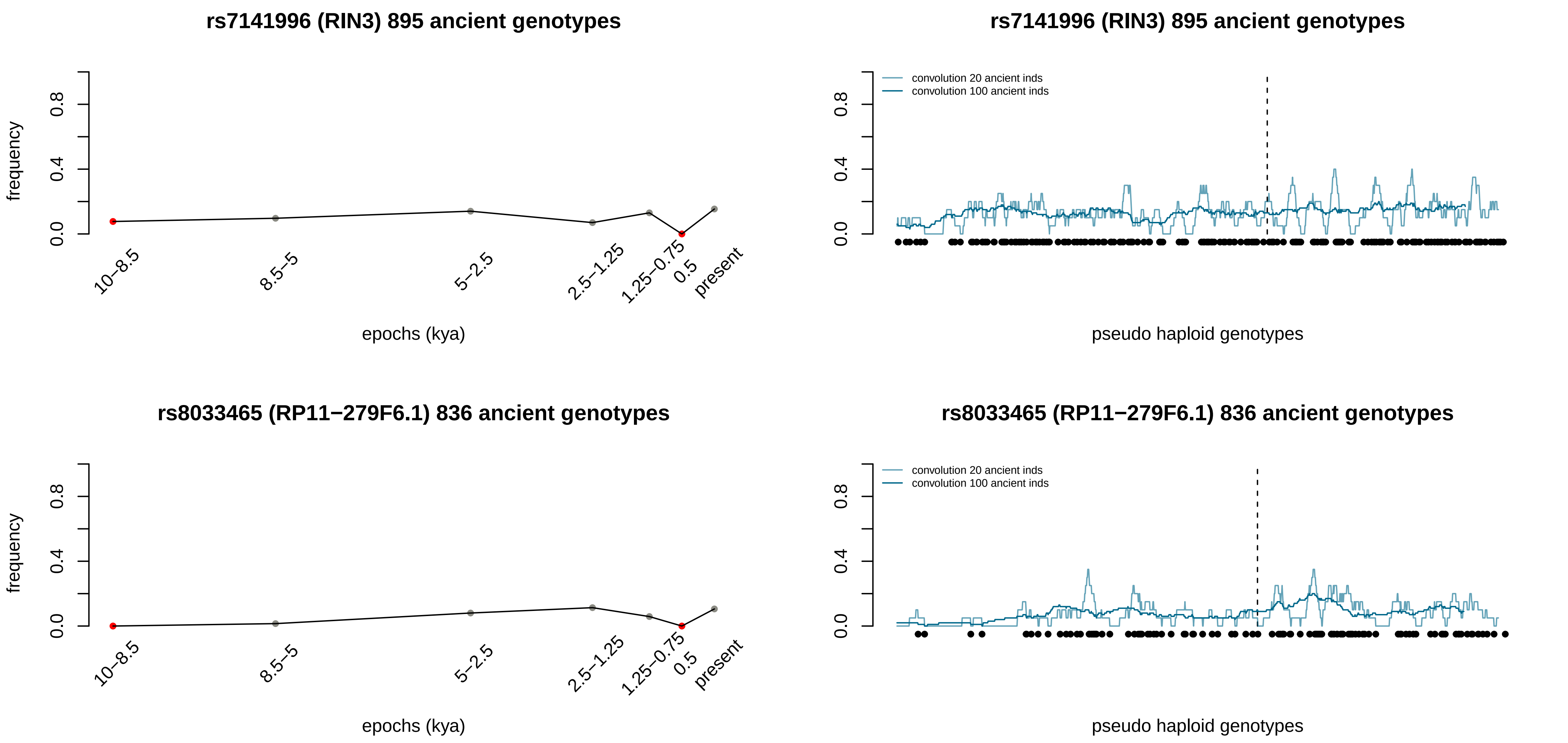
rs7141996 (RIN3) 895 ancient genotypes
frequency
0.0
0.4
0.8
10–8.5
8.5–5
5–2.5
2.5–1.25
1.25–0.75
0.5
present
epochs (kya)
rs7141996 (RIN3) 895 ancient genotypes
convolution 20 ancient inds
convolution 100 ancient inds
0.0
0.4
0.8
pseudo haploid genotypes
rs8033465 (RP11−279F6.1) 836 ancient genotypes
frequency
0.0
0.4
0.8
10–8.5
8.5–5
5–2.5
2.5–1.25
1.25–0.75
0.5
present
epochs (kya)
rs8033465 (RP11−279F6.1) 836 ancient genotypes
convolution 20 ancient inds
convolution 100 ancient inds
0.0
0.4
0.8
pseudo haploid genotypes

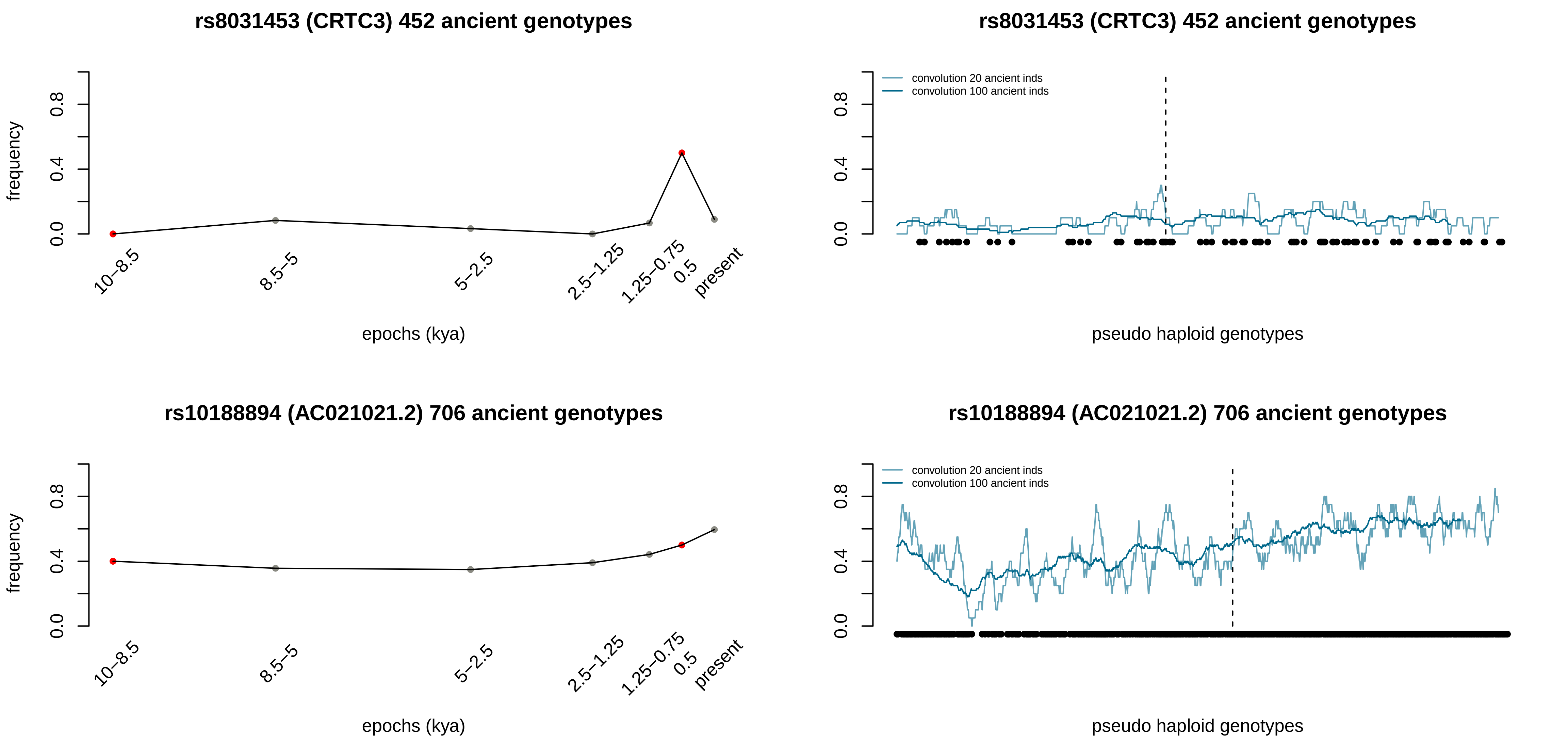
rs8031453 (CRTC3) 452 ancient genotypes
frequency
0.0
0.4
0.8
10–8.5
8.5–5
5–2.5
2.5–1.25
1.25–0.75
0.5
present
epochs (kya)
rs8031453 (CRTC3) 452 ancient genotypes
convolution 20 ancient inds
convolution 100 ancient inds
pseudo haploid genotypes
rs10188894 (AC021021.2) 706 ancient genotypes
frequency
epochs (kya)
rs10188894 (AC021021.2) 706 ancient genotypes
convolution 20 ancient inds
convolution 100 ancient inds
pseudo haploid genotypes

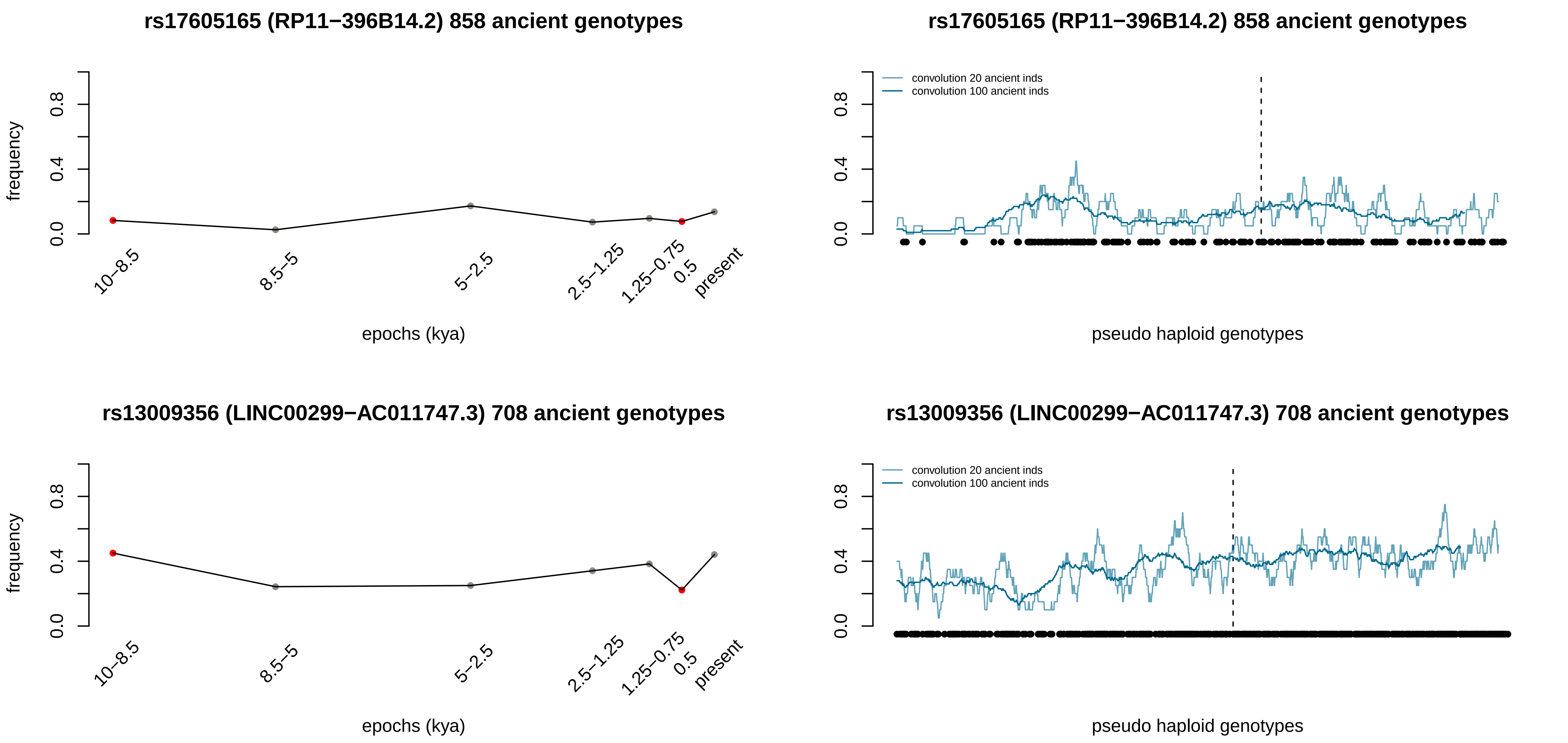
rs17605165 (RP11−396B14.2) 858 ancient genotypes
frequency
0.0
0.4
0.8
10−8.5
8.5−5
5−2.5
2.5−1.25
1.25−0.75
0.5
present
epochs (kya)
rs17605165 (RP11−396B14.2) 858 ancient genotypes
convolution 20 ancient inds
convolution 100 ancient inds
pseudo haploid genotypes
rs13009356 (LINC00299−AC011747.3) 708 ancient genotypes
frequency
epochs (kya)
rs13009356 (LINC00299−AC011747.3) 708 ancient genotypes
convolution 20 ancient inds
convolution 100 ancient inds
pseudo haploid genotypes

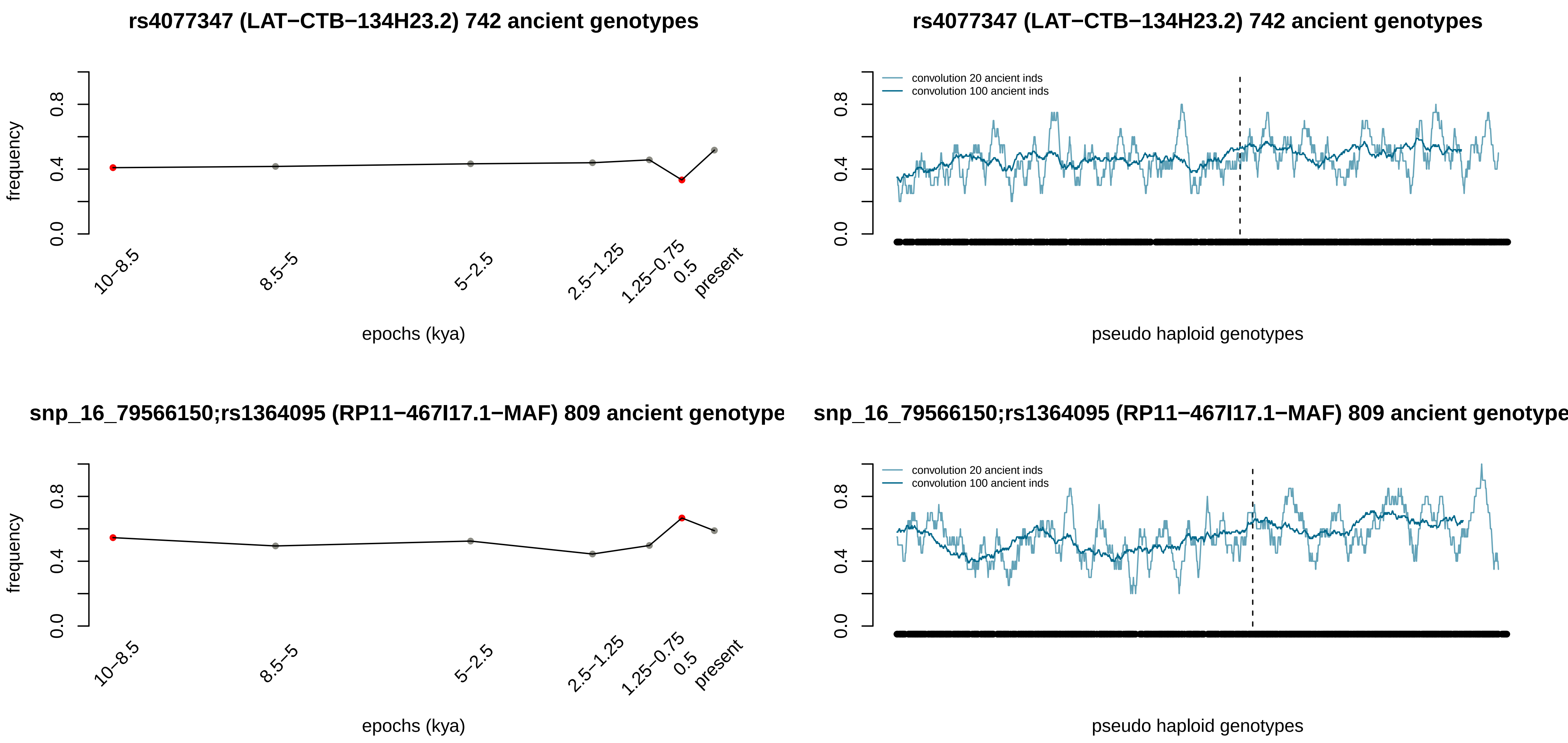
rs4077347 (LAT−CTB−134H23.2) 742 ancient genotypes
frequency
0.0
0.4
0.8
10−8.5
8.5−5
5−2.5
2.5−1.25
1.25−0.75
0.5
present
epochs (kya)
rs4077347 (LAT−CTB−134H23.2) 742 ancient genotypes
convolution 20 ancient inds
convolution 100 ancient inds
pseudo haploid genotypes
snp_16_79566150;rs1364095 (RP11−467I17.1−MAF) 809 ancient genotype
frequency
epochs (kya)
snp_16_79566150;rs1364095 (RP11−467I17.1−MAF) 809 ancient genotype
convolution 20 ancient inds
convolution 100 ancient inds
pseudo haploid genotypes

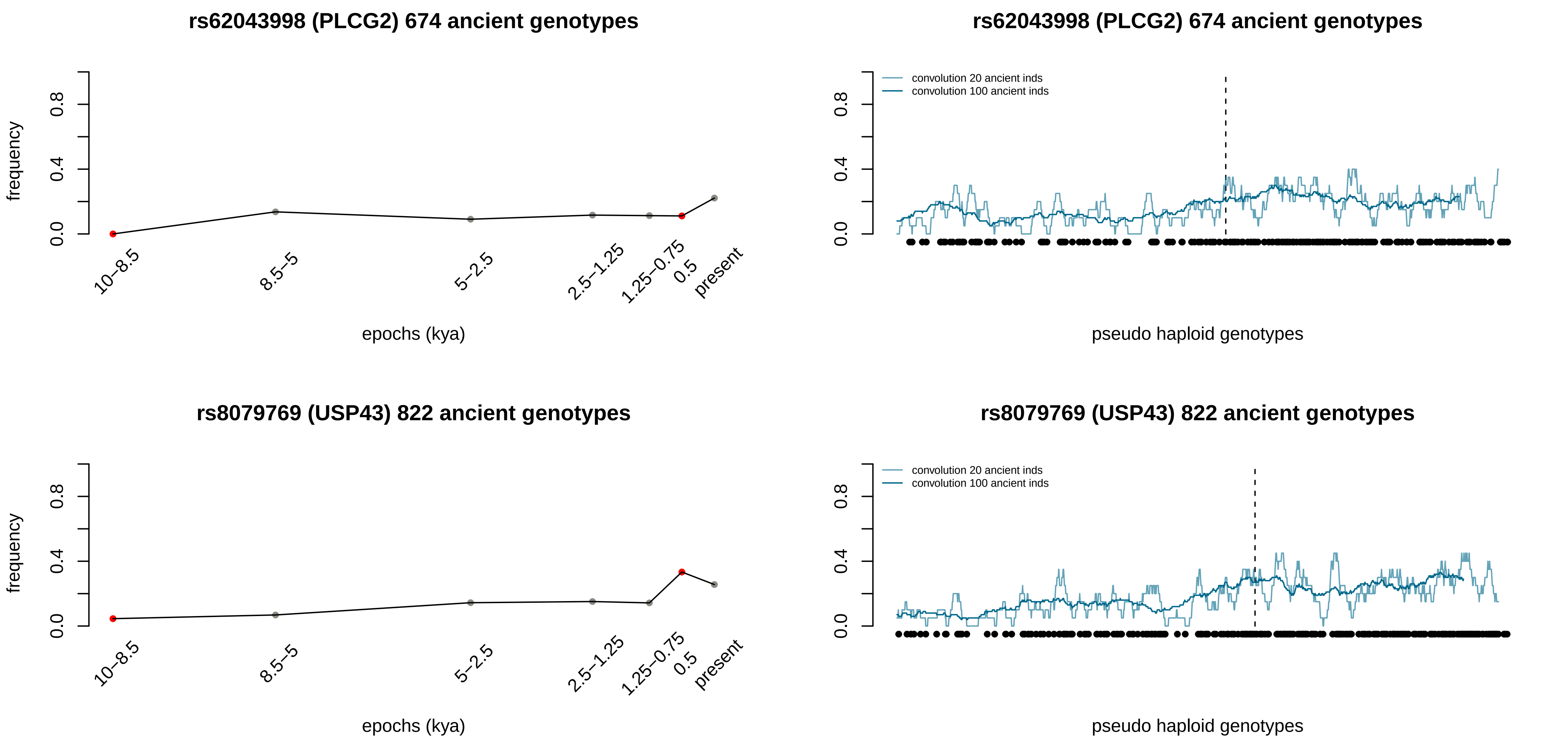
rs62043998 (PLCG2) 674 ancient genotypes
frequency
0.0
0.4
0.8
10–8.5
8.5–5
5–2.5
2.5–1.25
1.25–0.75
0.5
present
epochs (kya)
rs62043998 (PLCG2) 674 ancient genotypes
convolution 20 ancient inds
convolution 100 ancient inds
0.0
0.4
0.8
pseudo haploid genotypes
rs8079769 (USP43) 822 ancient genotypes
frequency
0.0
0.4
0.8
10–8.5
8.5–5
5–2.5
2.5–1.25
1.25–0.75
0.5
present
epochs (kya)
rs8079769 (USP43) 822 ancient genotypes
convolution 20 ancient inds
convolution 100 ancient inds
0.0
0.4
0.8
pseudo haploid genotypes

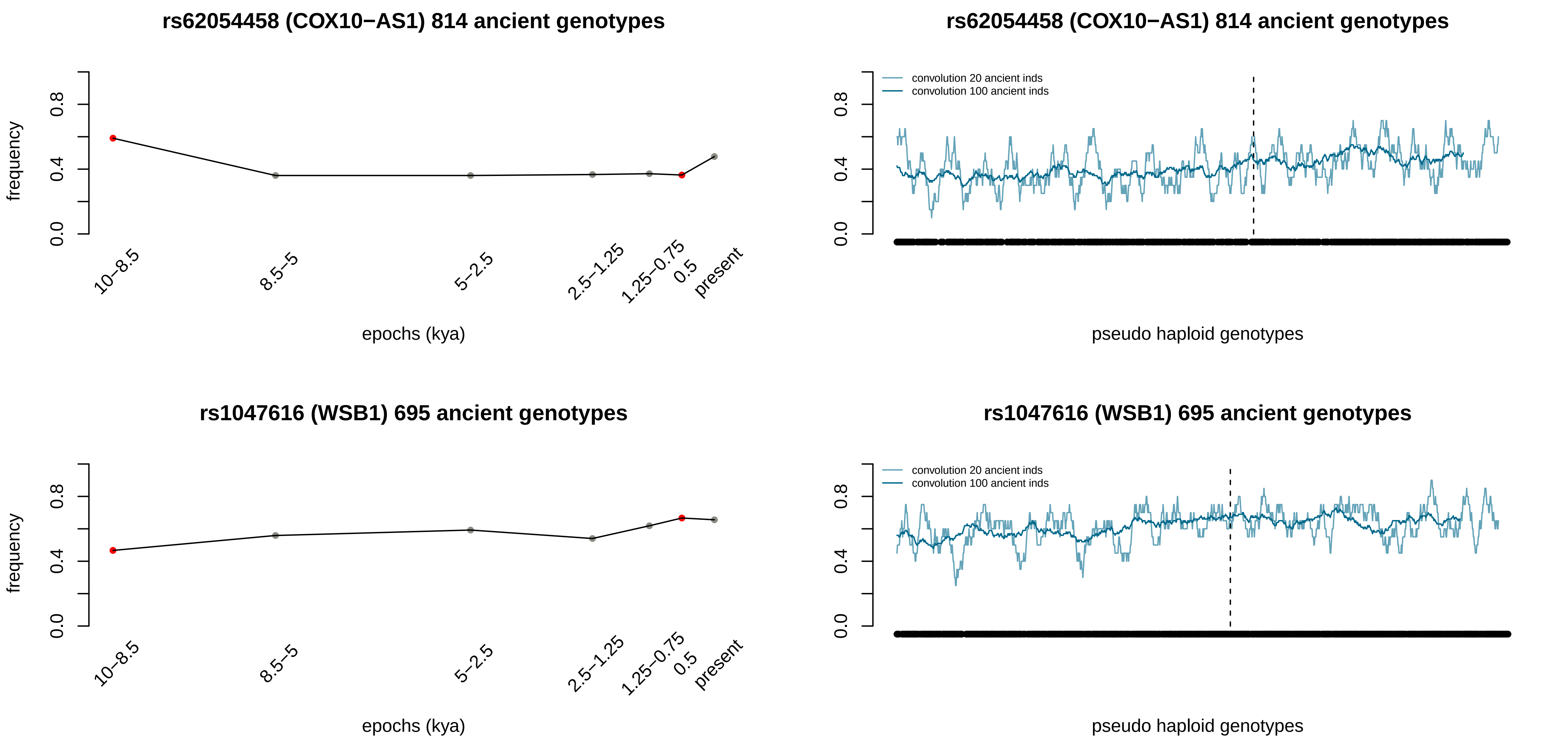

rs62054458 (COX10−AS1) 814 ancient genotypes
frequency
0.0
0.4
0.8
10–8.5
8.5–5
5–2.5
2.5–1.25
1.25–0.75
0.5
present
epochs (kya)
rs62054458 (COX10−AS1) 814 ancient genotypes
convolution 20 ancient inds
convolution 100 ancient inds
0.0
0.4
0.8
pseudo haploid genotypes
rs1047616 (WSB1) 695 ancient genotypes
frequency
0.0
0.4
0.8
10–8.5
8.5–5
5–2.5
2.5–1.25
1.25–0.75
0.5
present
epochs (kya)
rs1047616 (WSB1) 695 ancient genotypes
convolution 20 ancient inds
convolution 100 ancient inds
0.0
0.4
0.8
pseudo haploid genotypes

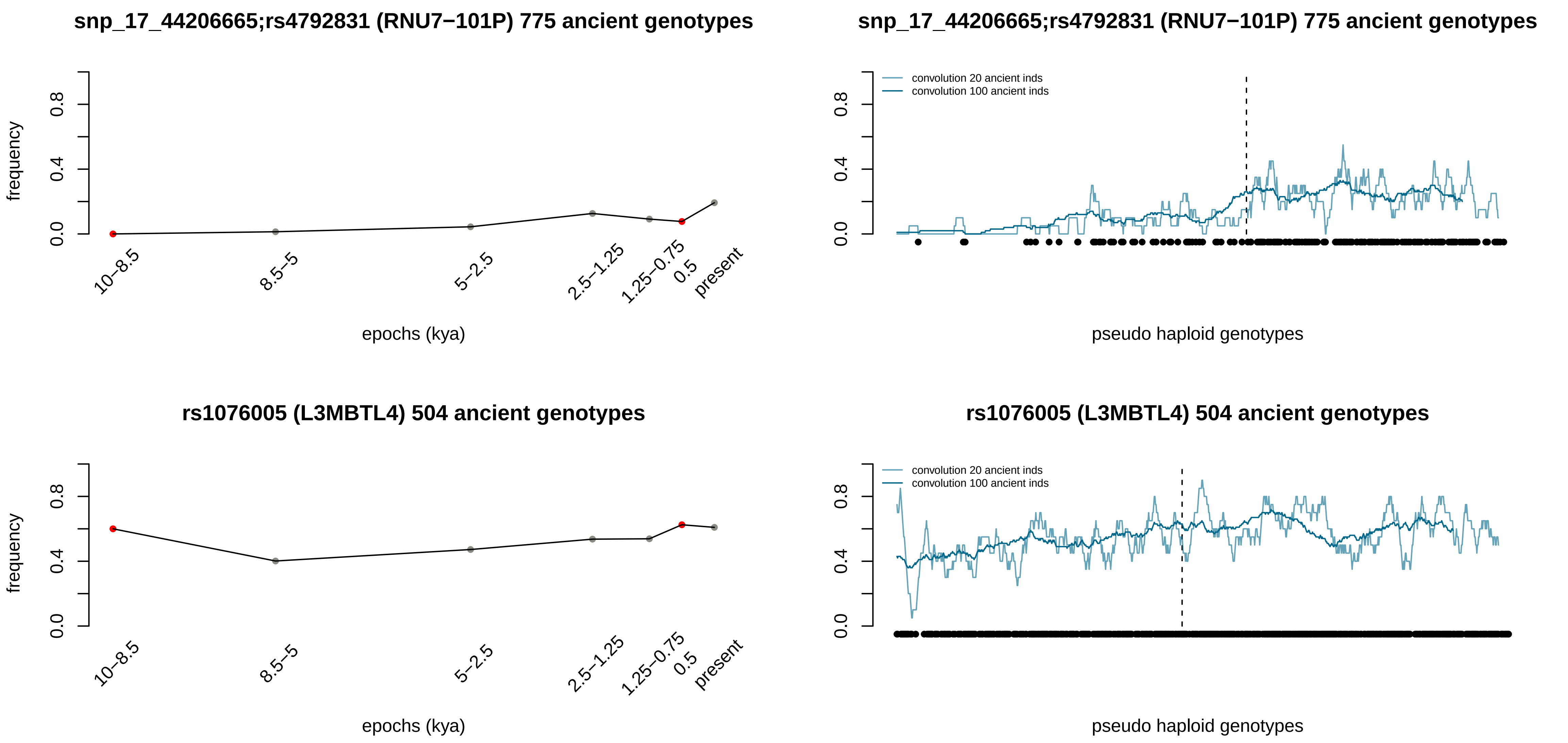
snp_17_44206665;rs4792831 (RNU7−101P) 775 ancient genotypes
frequency
0.0
0.4
0.8
10−8.5
8.5−5
5−2.5
2.5−1.25
1.25−0.75
0.5
present
epochs (kya)
snp_17_44206665;rs4792831 (RNU7−101P) 775 ancient genotypes
convolution 20 ancient inds
convolution 100 ancient inds
0.0
0.4
0.8
pseudo haploid genotypes
rs1076005 (L3MBTL4) 504 ancient genotypes
frequency
0.0
0.4
0.8
10−8.5
8.5−5
5−2.5
2.5−1.25
1.25−0.75
0.5
present
epochs (kya)
rs1076005 (L3MBTL4) 504 ancient genotypes
convolution 20 ancient inds
convolution 100 ancient inds
0.0
0.4
0.8
pseudo haploid genotypes

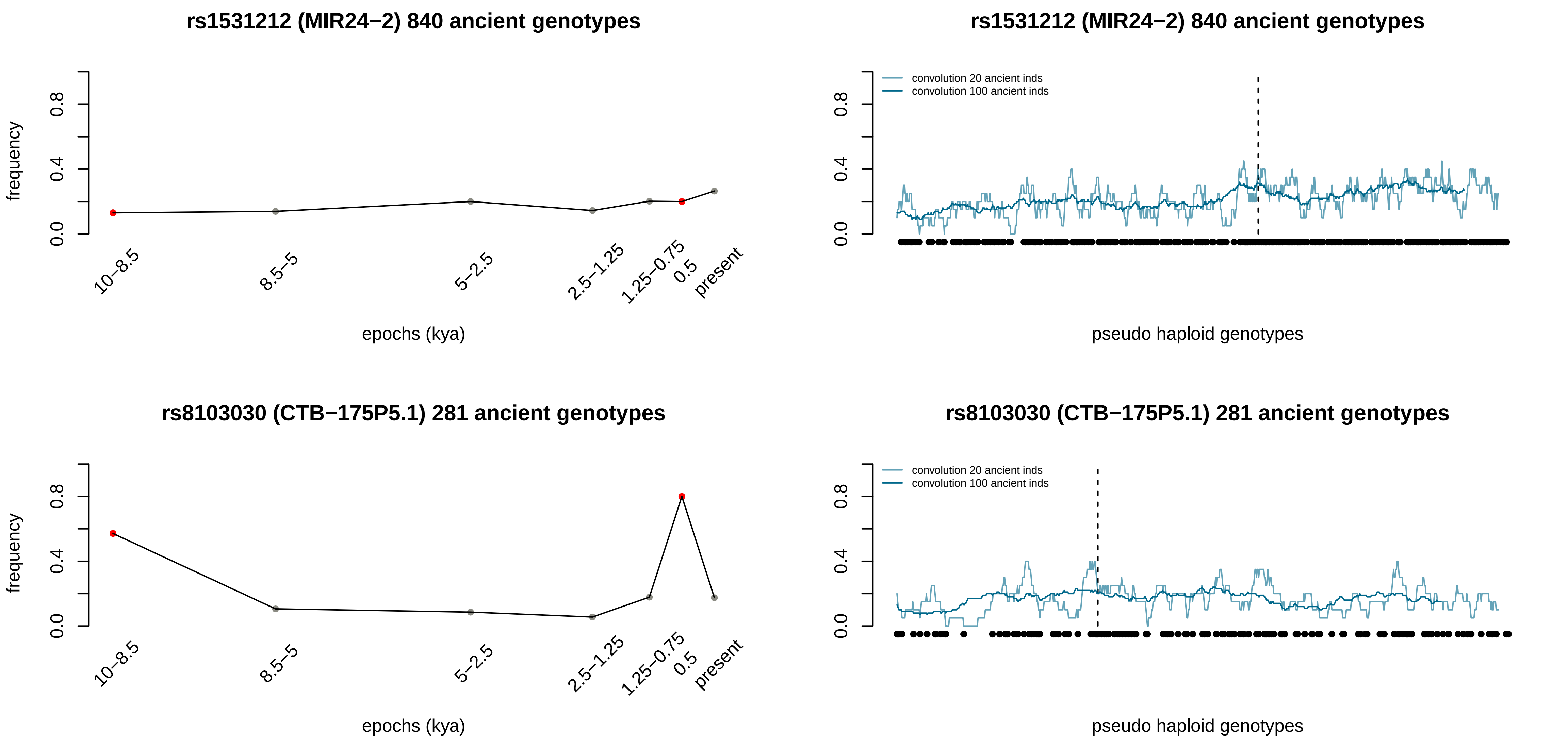
rs1531212 (MIR24−2) 840 ancient genotypes
frequency
0.0
0.4
0.8
10−8.5
8.5−5
5−2.5
2.5−1.25
1.25−0.75
0.5
present
epochs (kya)
rs1531212 (MIR24−2) 840 ancient genotypes
convolution 20 ancient inds
convolution 100 ancient inds
pseudo haploid genotypes
rs8103030 (CTB−175P5.1) 281 ancient genotypes
frequency
epochs (kya)
rs8103030 (CTB−175P5.1) 281 ancient genotypes
convolution 20 ancient inds
convolution 100 ancient inds
pseudo haploid genotypes

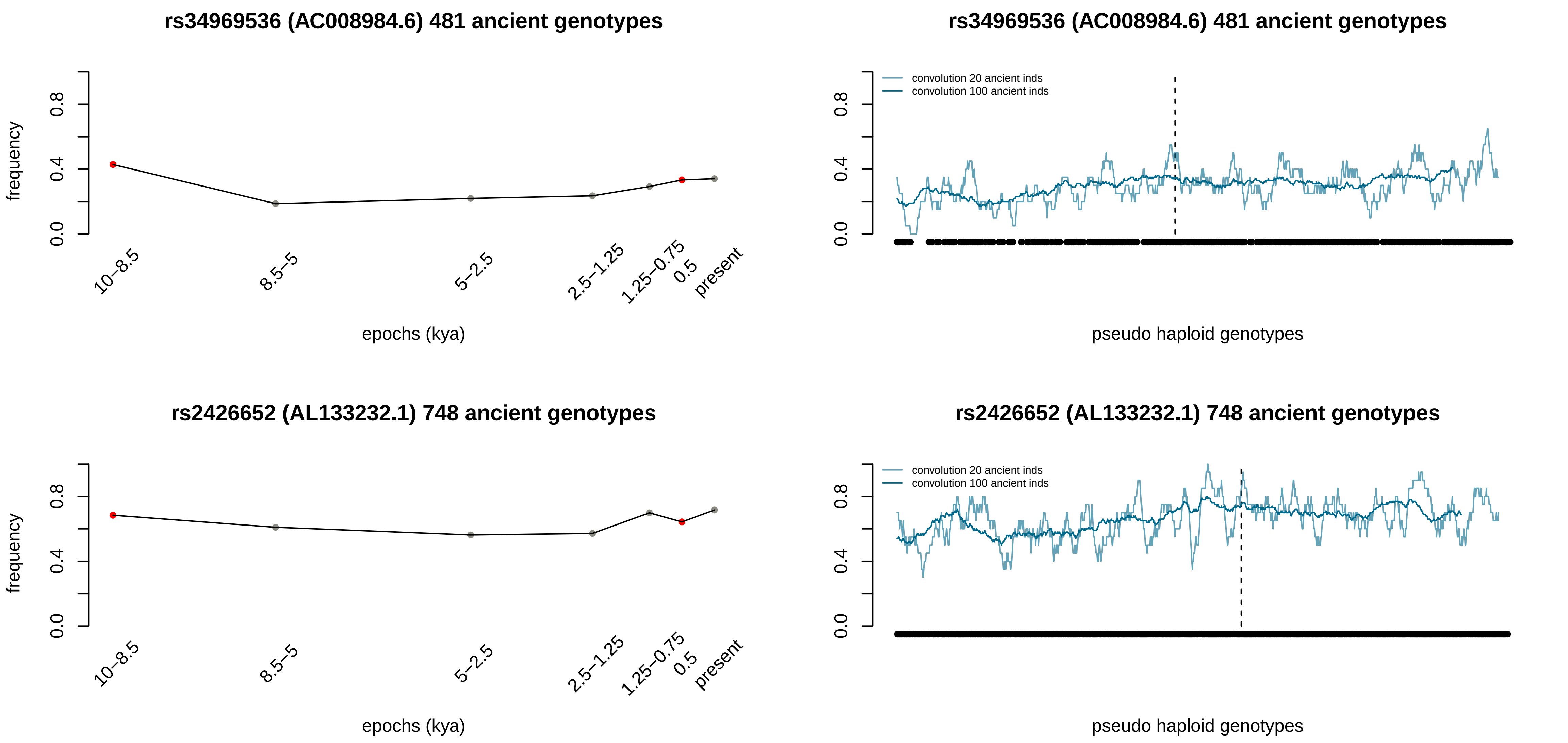
rs34969536 (AC008984.6) 481 ancient genotypes
frequency
0.0
0.4
0.8
10–8.5
8.5–5
5–2.5
2.5–1.25
1.25–0.75
0.5
present
epochs (kya)
rs34969536 (AC008984.6) 481 ancient genotypes
convolution 20 ancient inds
convolution 100 ancient inds
pseudo haploid genotypes
rs2426652 (AL133232.1) 748 ancient genotypes
frequency
epochs (kya)
rs2426652 (AL133232.1) 748 ancient genotypes
convolution 20 ancient inds
convolution 100 ancient inds
pseudo haploid genotypes

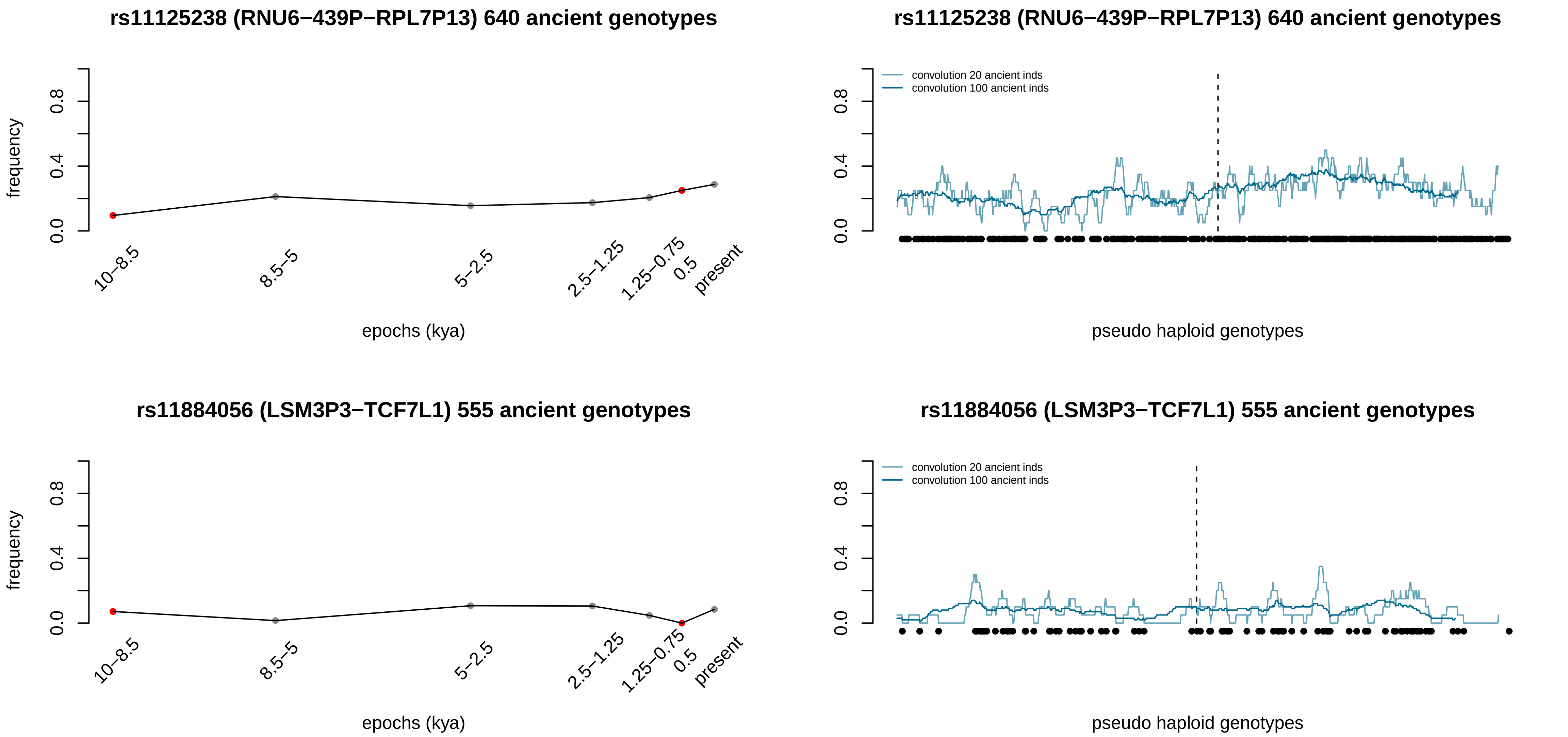
rs11125238 (RNU6−439P−RPL7P13) 640 ancient genotypes
frequency
0.0
0.4
0.8
10−8.5
8.5−5
5−2.5
2.5−1.25
1.25−0.75
0.5
present
epochs (kya)
rs11125238 (RNU6−439P−RPL7P13) 640 ancient genotypes
convolution 20 ancient inds
convolution 100 ancient inds
0.0
0.4
0.8
pseudo haploid genotypes
rs11884056 (LSM3P3−TCF7L1) 555 ancient genotypes
frequency
0.0
0.4
0.8
10−8.5
8.5−5
5−2.5
2.5−1.25
1.25−0.75
0.5
present
epochs (kya)
rs11884056 (LSM3P3−TCF7L1) 555 ancient genotypes
convolution 20 ancient inds
convolution 100 ancient inds
0.0
0.4
0.8
pseudo haploid genotypes

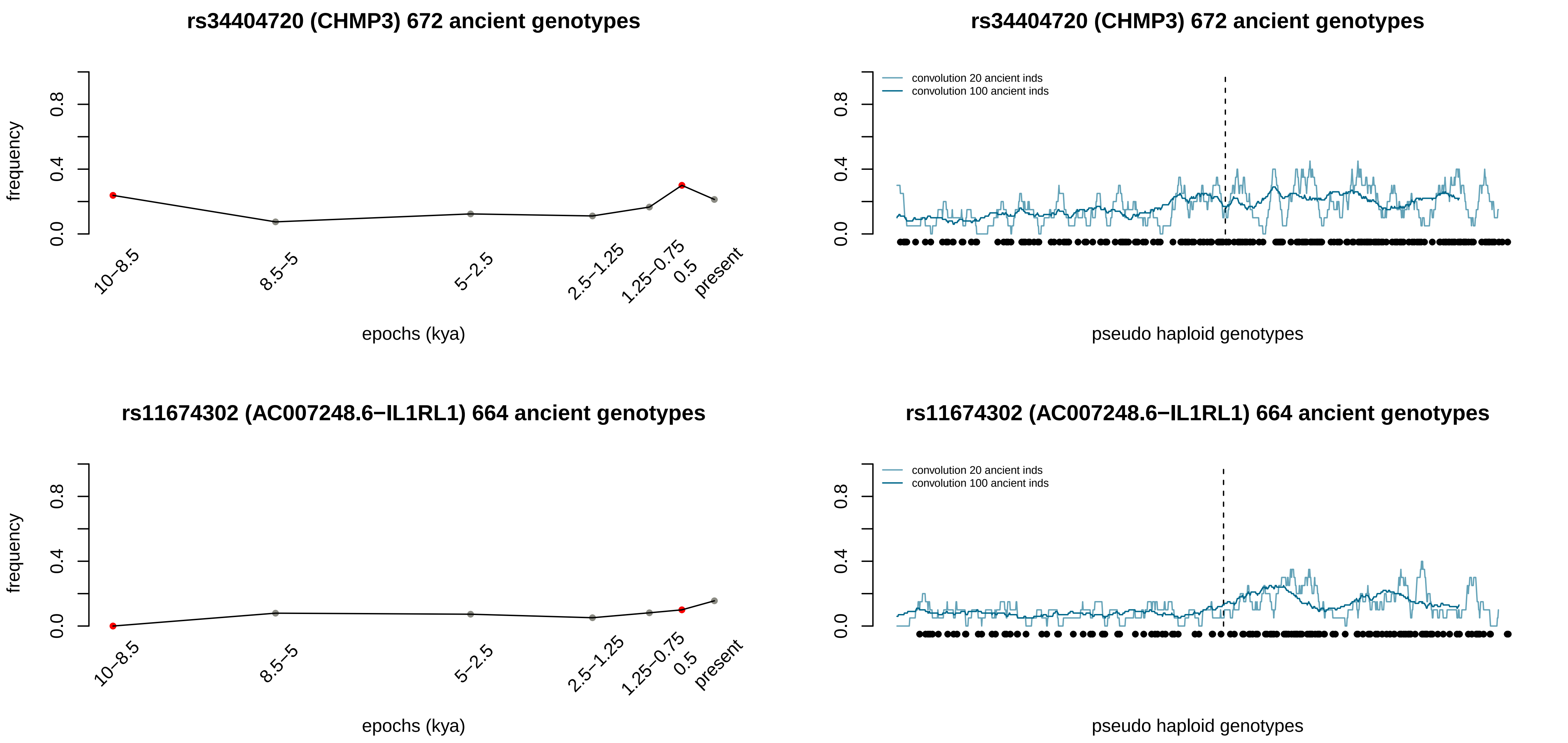
rs34404720 (CHMP3) 672 ancient genotypes
frequency
0.0
0.4
0.8
10–8.5
8.5–5
5–2.5
2.5–1.25
1.25–0.75
0.5
present
epochs (kya)
rs34404720 (CHMP3) 672 ancient genotypes
convolution 20 ancient inds
convolution 100 ancient inds
0.0
0.4
0.8
pseudo haploid genotypes
rs11674302 (AC007248.6−IL1RL1) 664 ancient genotypes
frequency
0.0
0.4
0.8
10–8.5
8.5–5
5–2.5
2.5–1.25
1.25–0.75
0.5
present
epochs (kya)
rs11674302 (AC007248.6−IL1RL1) 664 ancient genotypes
convolution 20 ancient inds
convolution 100 ancient inds
0.0
0.4
0.8
pseudo haploid genotypes

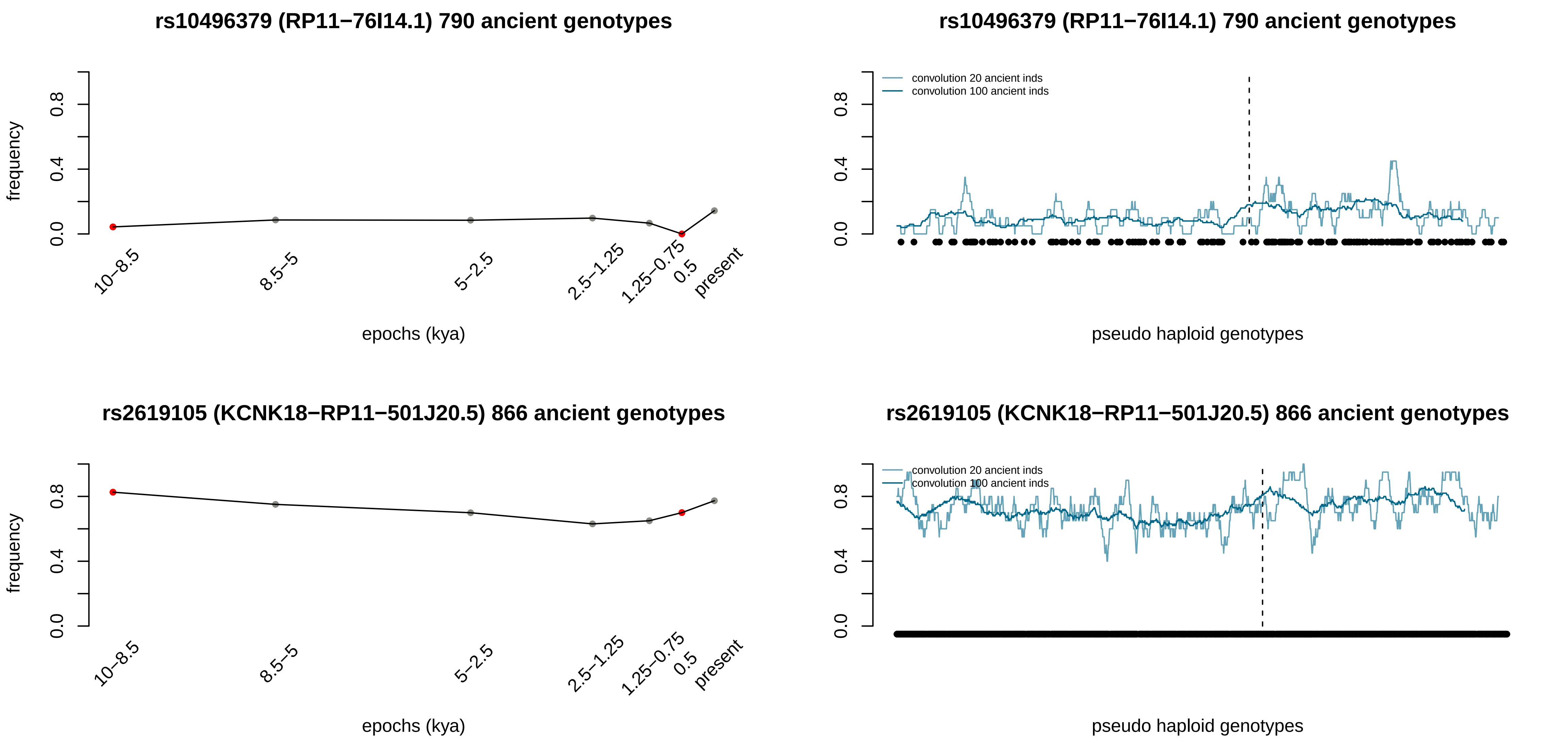
rs10496379 (RP11−76I14.1) 790 ancient genotypes
frequency
0.0
0.4
0.8
10−8.5
8.5−5
5−2.5
2.5−1.25
1.25−0.75
0.5
present
epochs (kya)
rs10496379 (RP11−76I14.1) 790 ancient genotypes
convolution 20 ancient inds
convolution 100 ancient inds
0.0
0.4
0.8
pseudo haploid genotypes
rs2619105 (KCNK18−RP11−501J20.5) 866 ancient genotypes
frequency
0.0
0.4
0.8
10−8.5
8.5−5
5−2.5
2.5−1.25
1.25−0.75
0.5
present
epochs (kya)
rs2619105 (KCNK18−RP11−501J20.5) 866 ancient genotypes
convolution 20 ancient inds
convolution 100 ancient inds
0.0
0.4
0.8
pseudo haploid genotypes

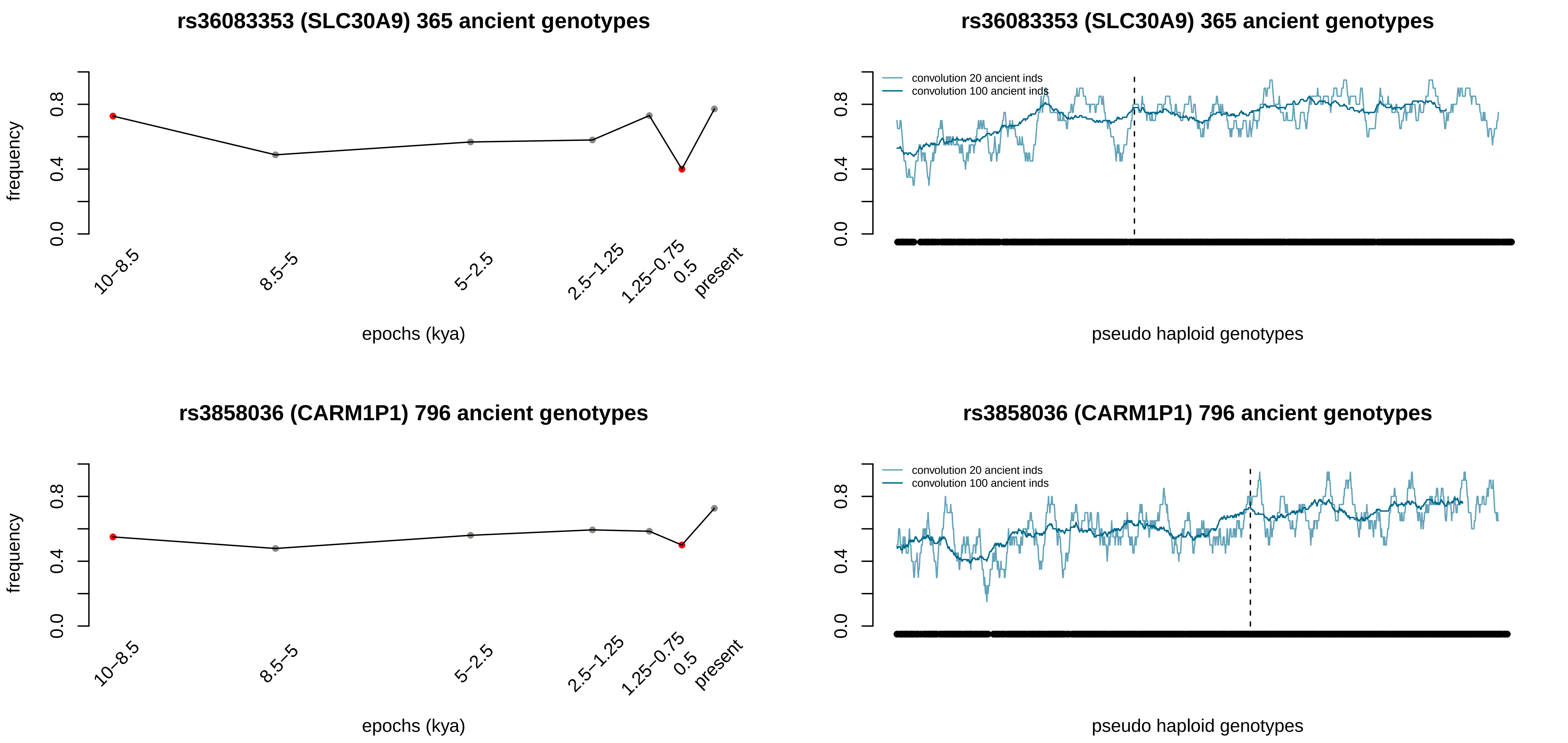
rs36083353 (SLC30A9) 365 ancient genotypes
frequency
0.0
0.4
0.8
10–8.5
8.5–5
5–2.5
2.5–1.25
1.25–0.75
0.5
present
epochs (kya)
rs36083353 (SLC30A9) 365 ancient genotypes
convolution 20 ancient inds
convolution 100 ancient inds
0.0
0.4
0.8
pseudo haploid genotypes
rs3858036 (CARM1P1) 796 ancient genotypes
frequency
0.0
0.4
0.8
10–8.5
8.5–5
5–2.5
2.5–1.25
1.25–0.75
0.5
present
epochs (kya)
rs3858036 (CARM1P1) 796 ancient genotypes
convolution 20 ancient inds
convolution 100 ancient inds
0.0
0.4
0.8
pseudo haploid genotypes

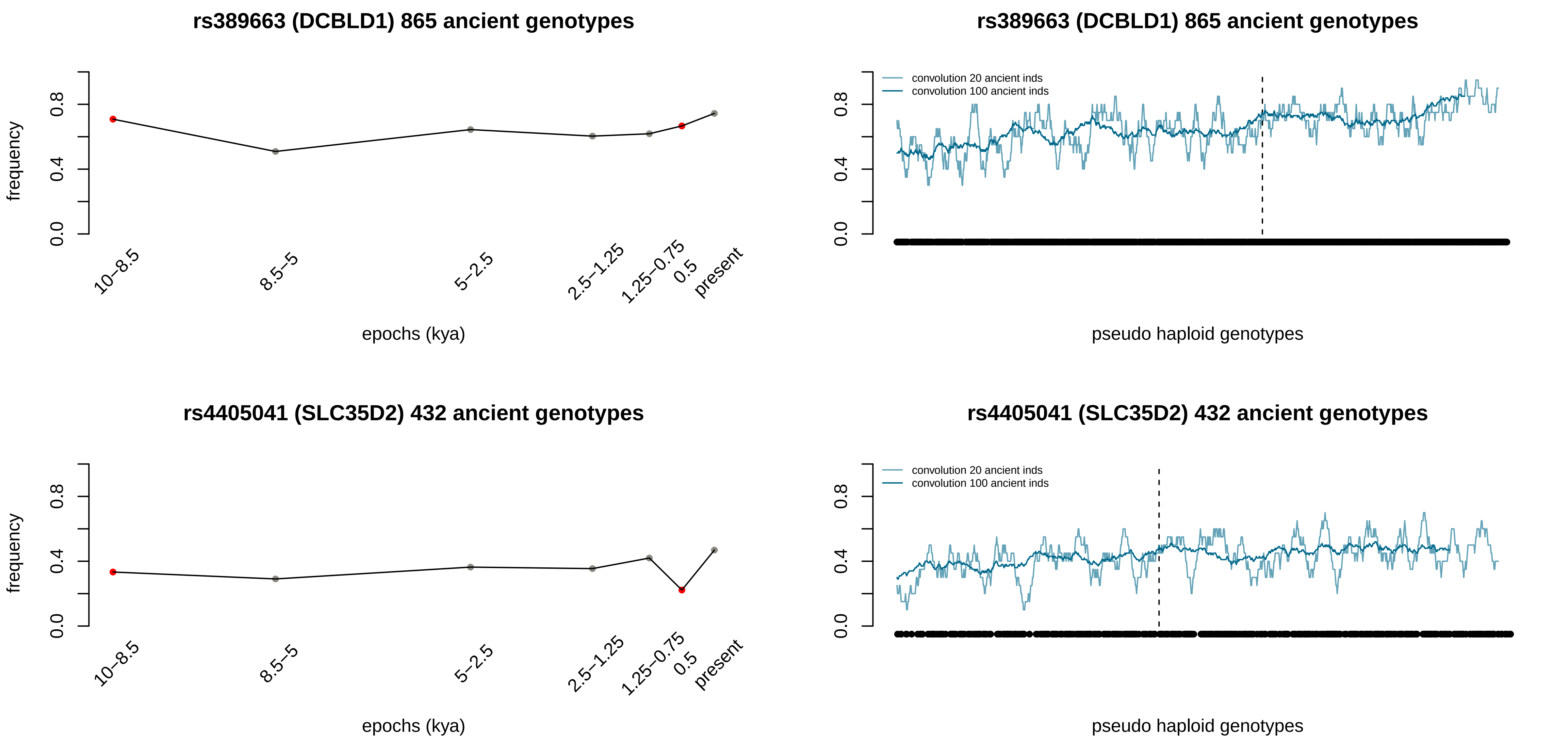

rs389663 (DCBLD1) 865 ancient genotypes
frequency
0.0
0.4
0.8
10–8.5
8.5–5
5–2.5
2.5–1.25
1.25–0.75
0.5
present
epochs (kya)
rs389663 (DCBLD1) 865 ancient genotypes
convolution 20 ancient inds
convolution 100 ancient inds
pseudo haploid genotypes
rs4405041 (SLC35D2) 432 ancient genotypes
frequency
epochs (kya)
rs4405041 (SLC35D2) 432 ancient genotypes
convolution 20 ancient inds
convolution 100 ancient inds
pseudo haploid genotypes

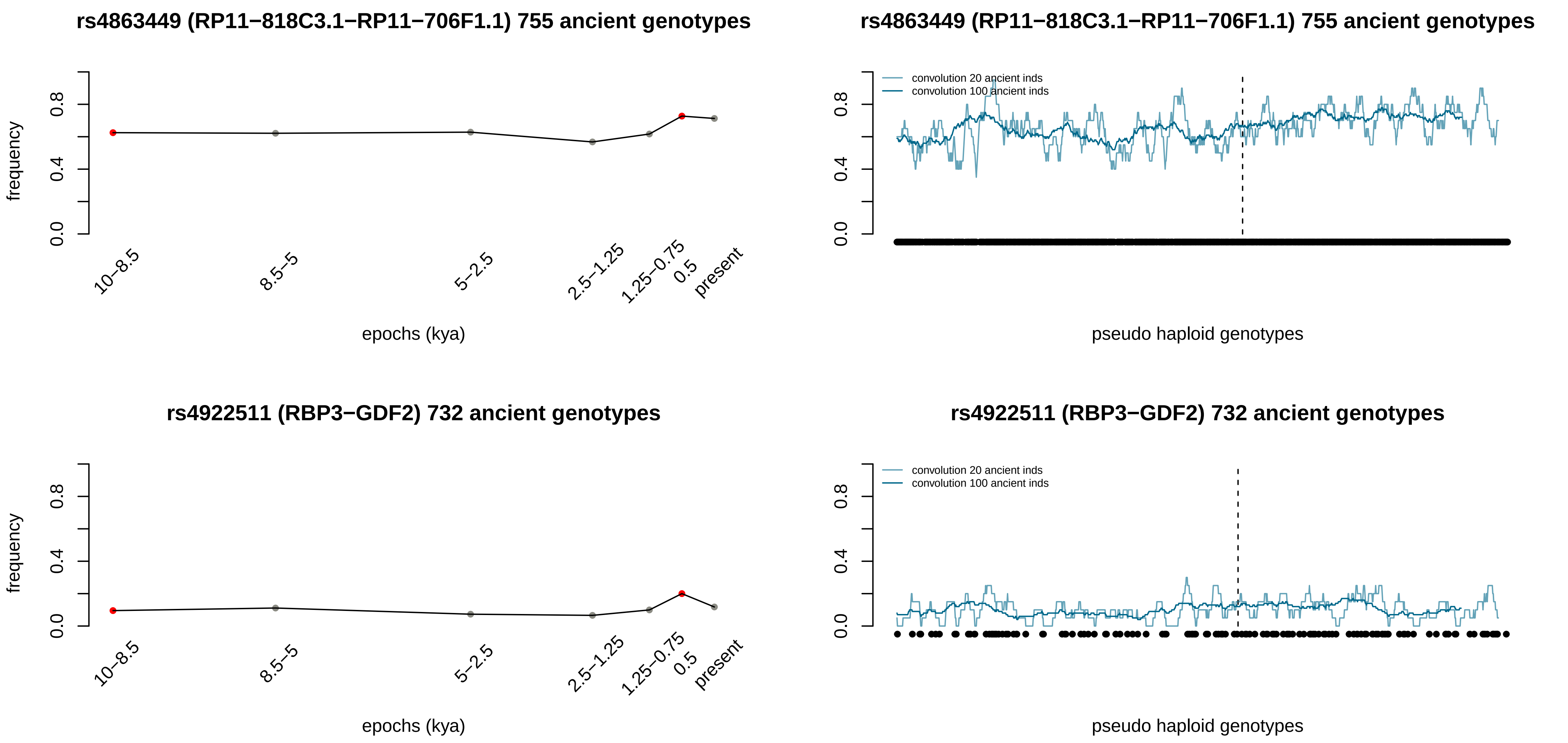

rs4863449 (RP11−818C3.1−RP11−706F1.1) 755 ancient genotypes
frequency
0.0
0.4
0.8
10−8.5
8.5−5
5−2.5
2.5−1.25
1.25−0.75
0.5
present
epochs (kya)
rs4863449 (RP11−818C3.1−RP11−706F1.1) 755 ancient genotypes
convolution 20 ancient inds
convolution 100 ancient inds
pseudo haploid genotypes
rs4922511 (RBP3−GDF2) 732 ancient genotypes
frequency
epochs (kya)
rs4922511 (RBP3−GDF2) 732 ancient genotypes
convolution 20 ancient inds
convolution 100 ancient inds
pseudo haploid genotypes

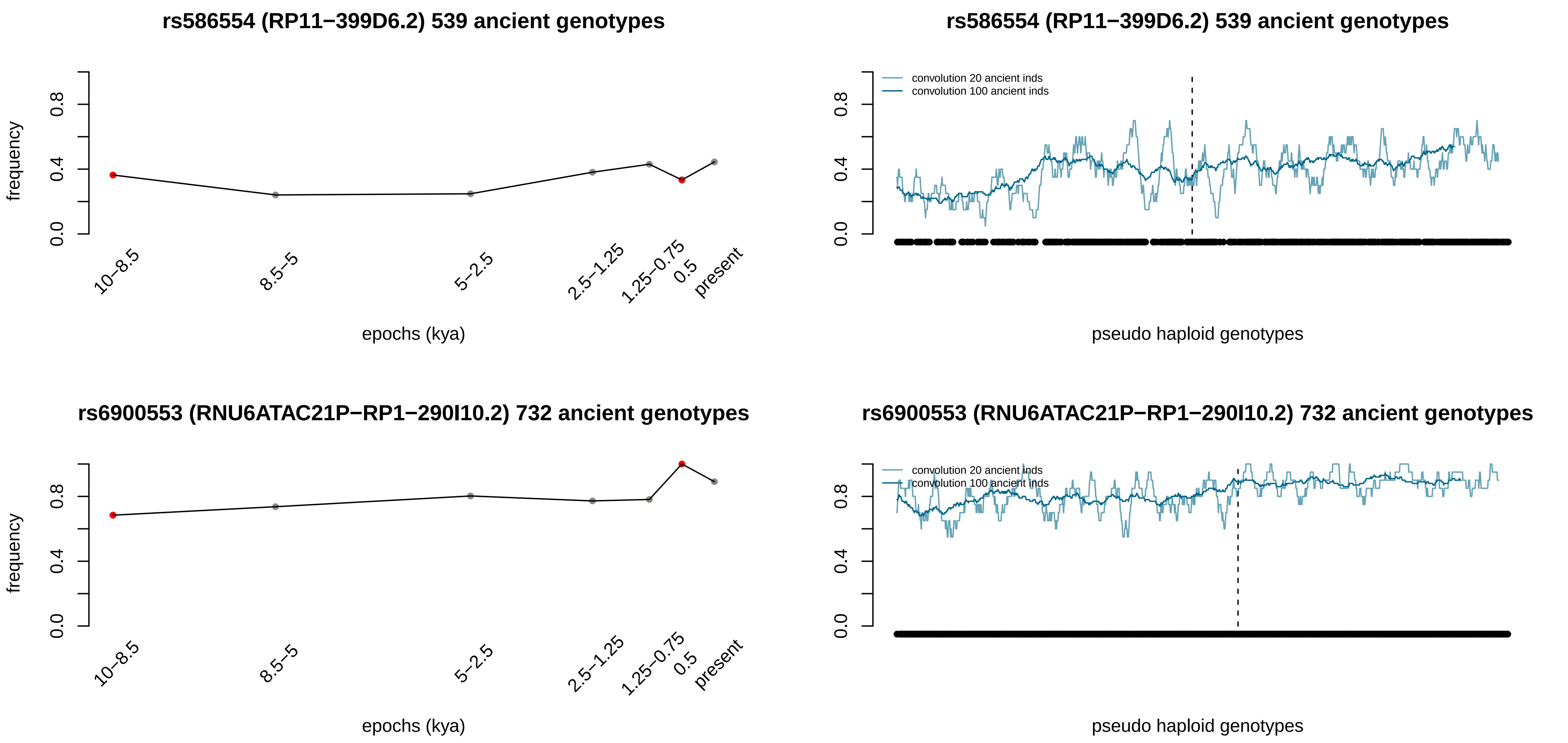
rs586554 (RP11−399D6.2) 539 ancient genotypes
frequency
0.0
0.4
0.8
10−8.5
8.5−5
5−2.5
2.5−1.25
1.25−0.75
0.5
present
epochs (kya)
rs586554 (RP11−399D6.2) 539 ancient genotypes
convolution 20 ancient inds
convolution 100 ancient inds
0.0
0.4
0.8
pseudo haploid genotypes
rs6900553 (RNU6ATAC21P−RP1−290I10.2) 732 ancient genotypes
frequency
0.0
0.4
0.8
10−8.5
8.5−5
5−2.5
2.5−1.25
1.25−0.75
0.5
present
epochs (kya)
rs6900553 (RNU6ATAC21P−RP1−290I10.2) 732 ancient genotypes
convolution 20 ancient inds
convolution 100 ancient inds
0.0
0.4
0.8
pseudo haploid genotypes

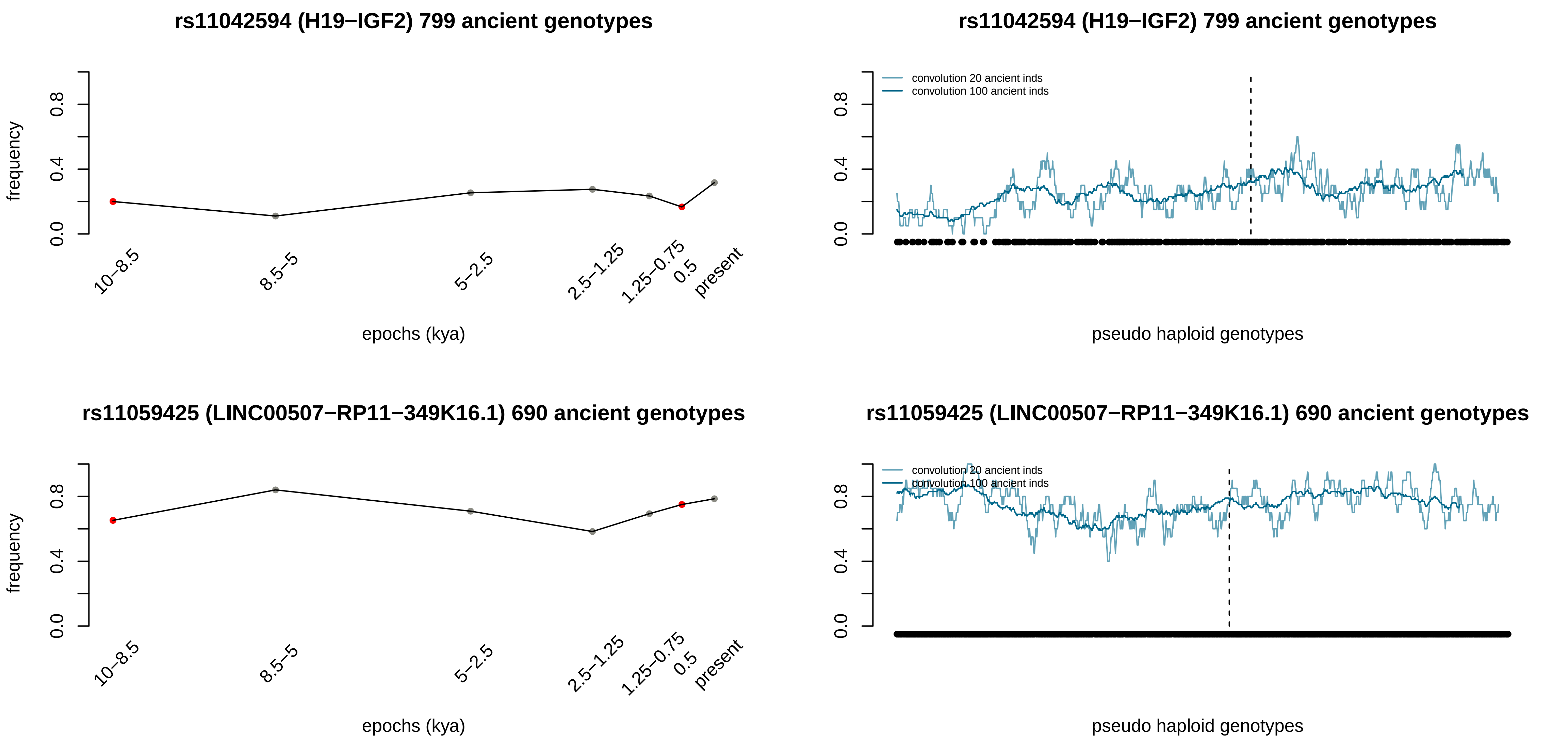

rs11042594 (H19−IGF2) 799 ancient genotypes
frequency
0.0
0.4
0.8
10−8.5
8.5−5
5−2.5
2.5−1.25
1.25−0.75
0.5
present
epochs (kya)
rs11042594 (H19−IGF2) 799 ancient genotypes
convolution 20 ancient inds
convolution 100 ancient inds
pseudo haploid genotypes
rs11059425 (LINC00507−RP11−349K16.1) 690 ancient genotypes
frequency
epochs (kya)
rs11059425 (LINC00507−RP11−349K16.1) 690 ancient genotypes
convolution 20 ancient inds
convolution 100 ancient inds
pseudo haploid genotypes

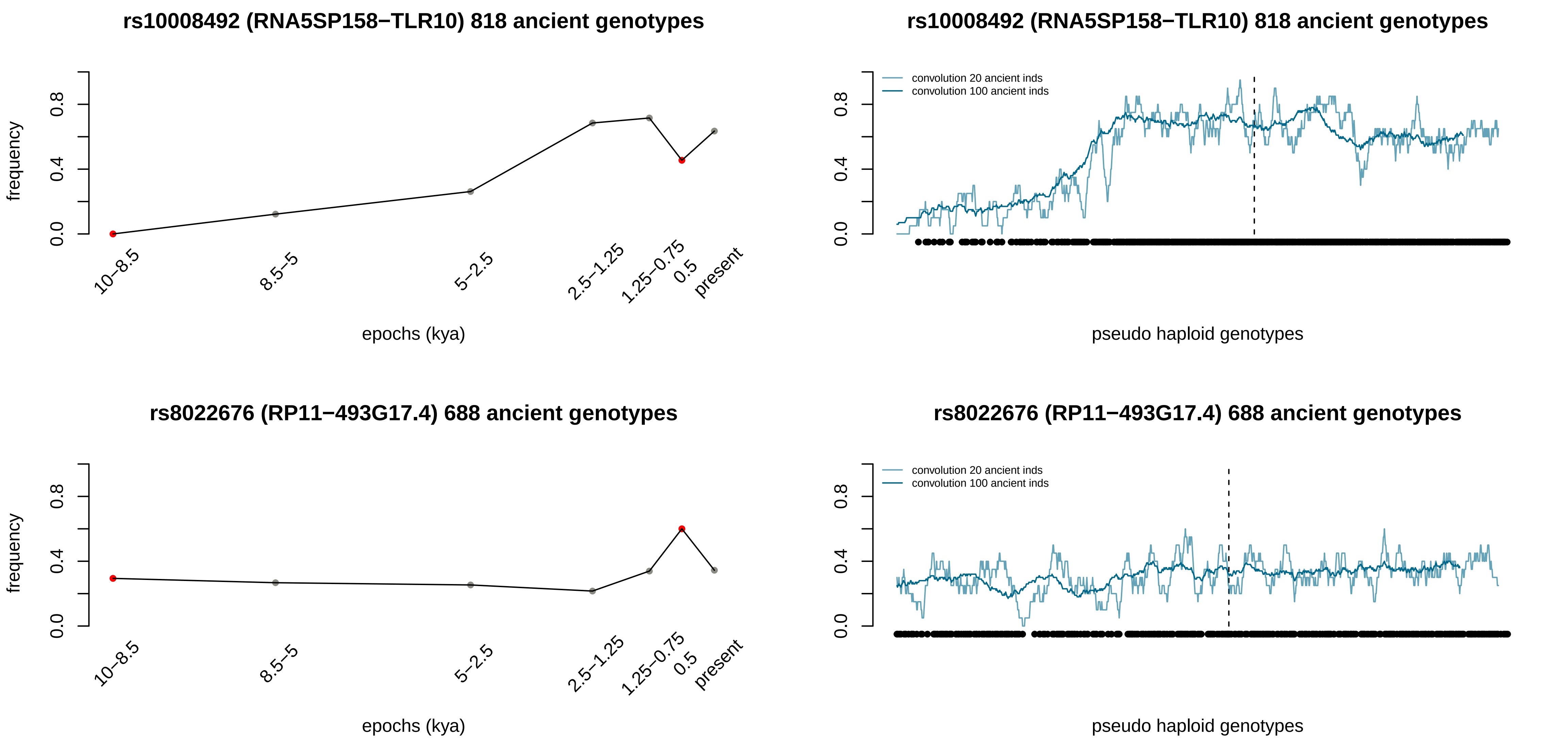

rs10008492 (RNA5SP158−TLR10) 818 ancient genotypes
frequency
0.0
0.4
0.8
10−8.5
8.5−5
5−2.5
2.5−1.25
1.25−0.75
0.5
present
epochs (kya)
rs10008492 (RNA5SP158−TLR10) 818 ancient genotypes
convolution 20 ancient inds
convolution 100 ancient inds
pseudo haploid genotypes
rs8022676 (RP11−493G17.4) 688 ancient genotypes
frequency
epochs (kya)
rs8022676 (RP11−493G17.4) 688 ancient genotypes
convolution 20 ancient inds
convolution 100 ancient inds
pseudo haploid genotypes

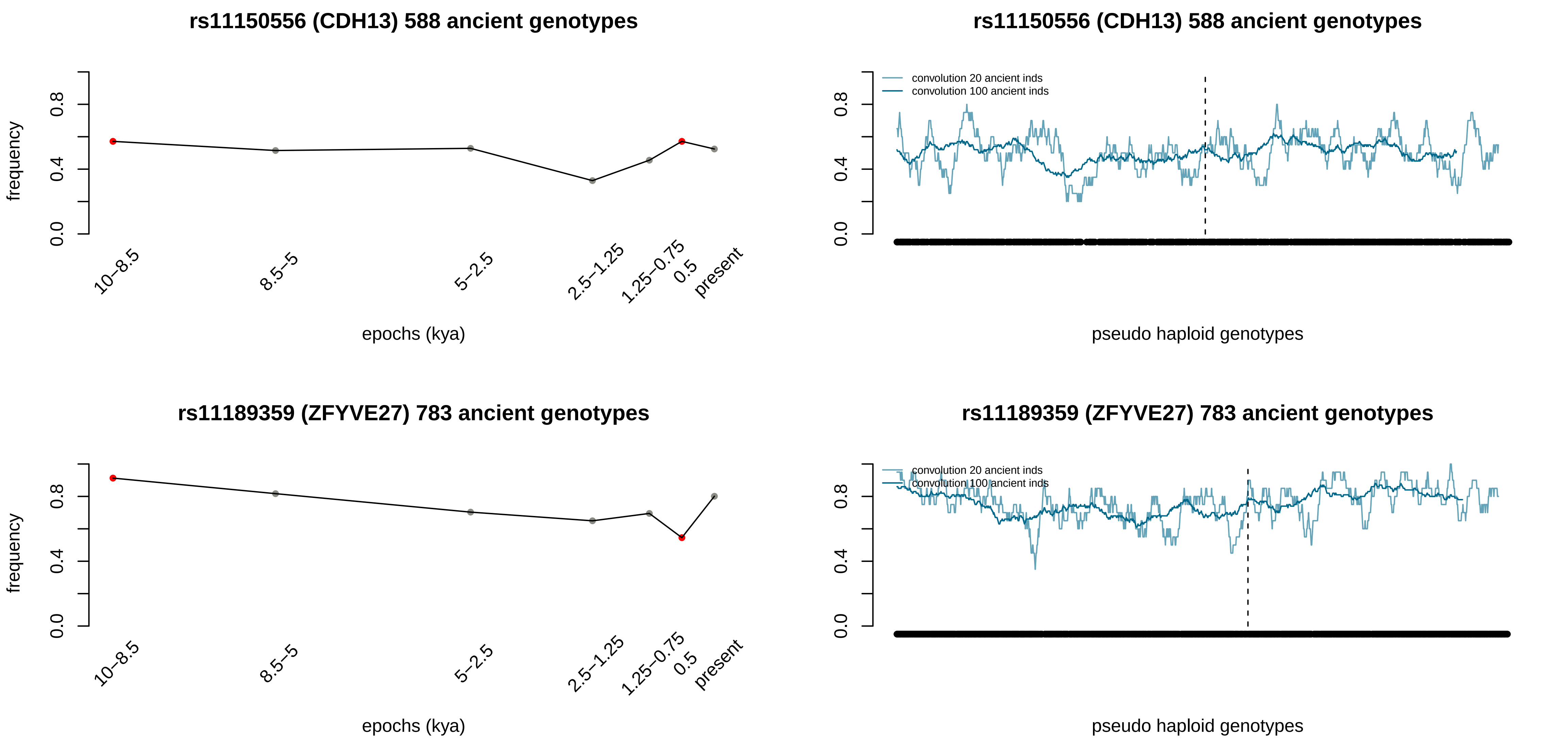
rs11150556 (CDH13) 588 ancient genotypes
frequency
0.0
0.4
0.8
10–8.5
8.5–5
5–2.5
2.5–1.25
1.25–0.75
0.5
present
epochs (kya)
rs11150556 (CDH13) 588 ancient genotypes
convolution 20 ancient inds
convolution 100 ancient inds
0.0
0.4
0.8
pseudo haploid genotypes
rs11189359 (ZFYVE27) 783 ancient genotypes
frequency
0.0
0.4
0.8
10–8.5
8.5–5
5–2.5
2.5–1.25
1.25–0.75
0.5
present
epochs (kya)
rs11189359 (ZFYVE27) 783 ancient genotypes
convolution 20 ancient inds
convolution 100 ancient inds
0.0
0.4
0.8
pseudo haploid genotypes

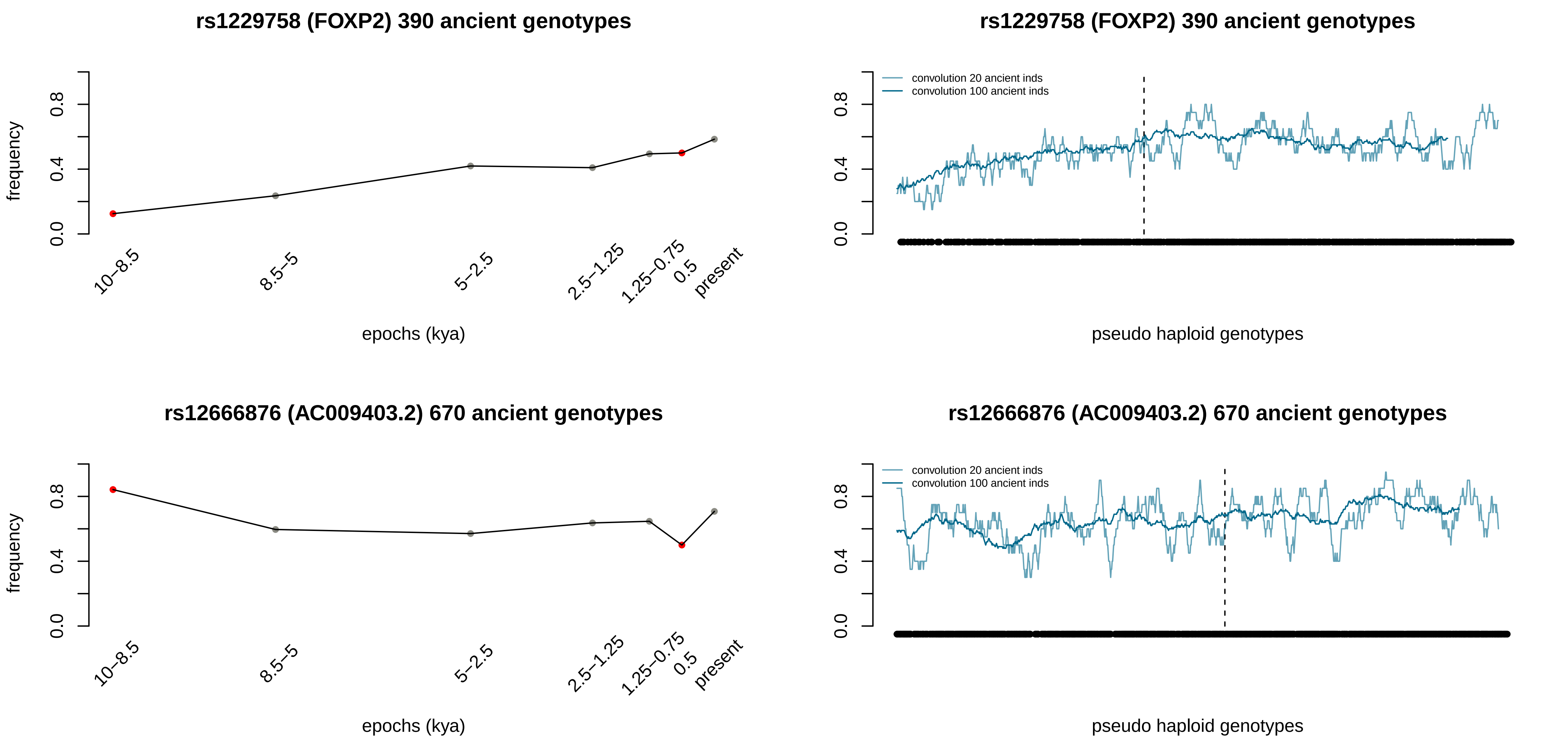
rs1229758 (FOXP2) 390 ancient genotypes
frequency
0.0
0.4
0.8
10–8.5
8.5–5
5–2.5
2.5–1.25
1.25–0.75
0.5
present
epochs (kya)
rs1229758 (FOXP2) 390 ancient genotypes
convolution 20 ancient inds
convolution 100 ancient inds
pseudo haploid genotypes
rs12666876 (AC009403.2) 670 ancient genotypes
frequency
epochs (kya)
rs12666876 (AC009403.2) 670 ancient genotypes
convolution 20 ancient inds
convolution 100 ancient inds
pseudo haploid genotypes

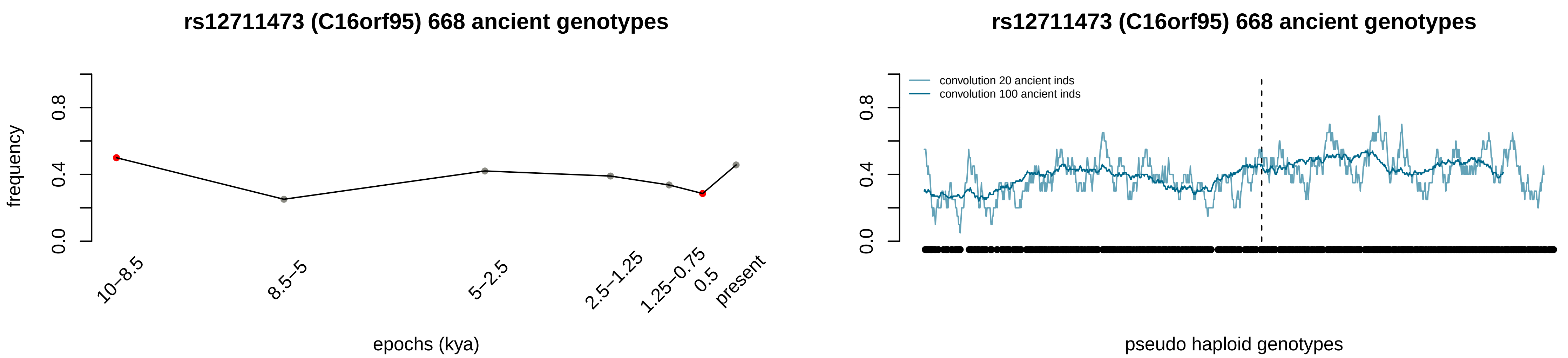

rs12711473 (C16orf95) 668 ancient genotypes
frequency
0.0
0.4
0.8
10–8.5
8.5–5
5–2.5
2.5–1.25
1.25–0.75
0.5
present
epochs (kya)
rs12711473 (C16orf95) 668 ancient genotypes
convolution 20 ancient inds
convolution 100 ancient inds
0.0
0.4
0.8
pseudo haploid genotypes
