## Supplementary material for "Deep estimation of the intensity and timing of selection from ancient genomes": 1D images for the 89 positively selected variants in southern Europe

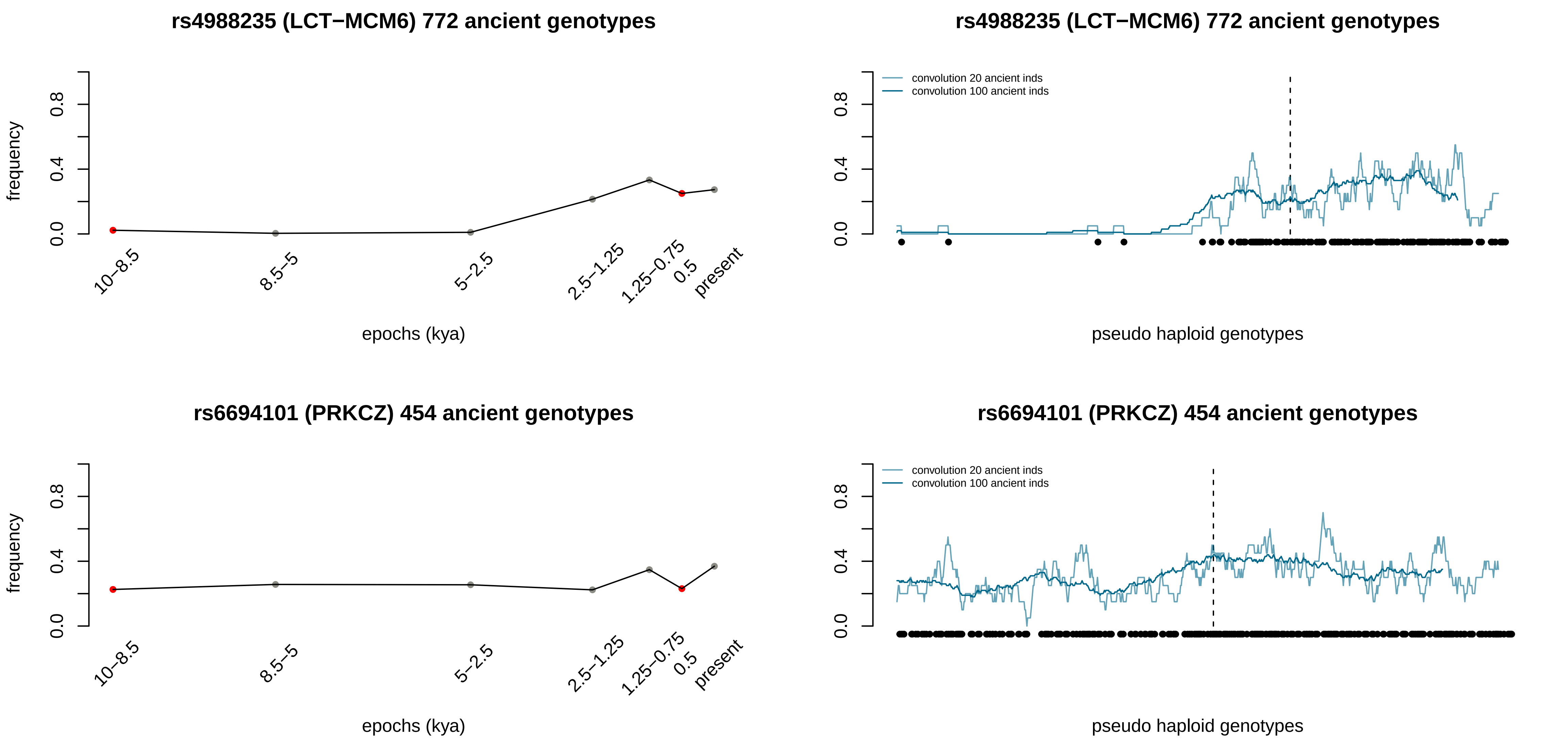

rs4988235 (LCT−MCM6) 772 ancient genotypes
frequency
0.0
0.4
0.8
10–8.5
8.5–5
5–2.5
2.5–1.25
1.25–0.75
0.5
present
epochs (kya)
rs4988235 (LCT−MCM6) 772 ancient genotypes
convolution 20 ancient inds
convolution 100 ancient inds
pseudo haploid genotypes
rs6694101 (PRKCZ) 454 ancient genotypes
frequency
epochs (kya)
rs6694101 (PRKCZ) 454 ancient genotypes
convolution 20 ancient inds
convolution 100 ancient inds
pseudo haploid genotypes

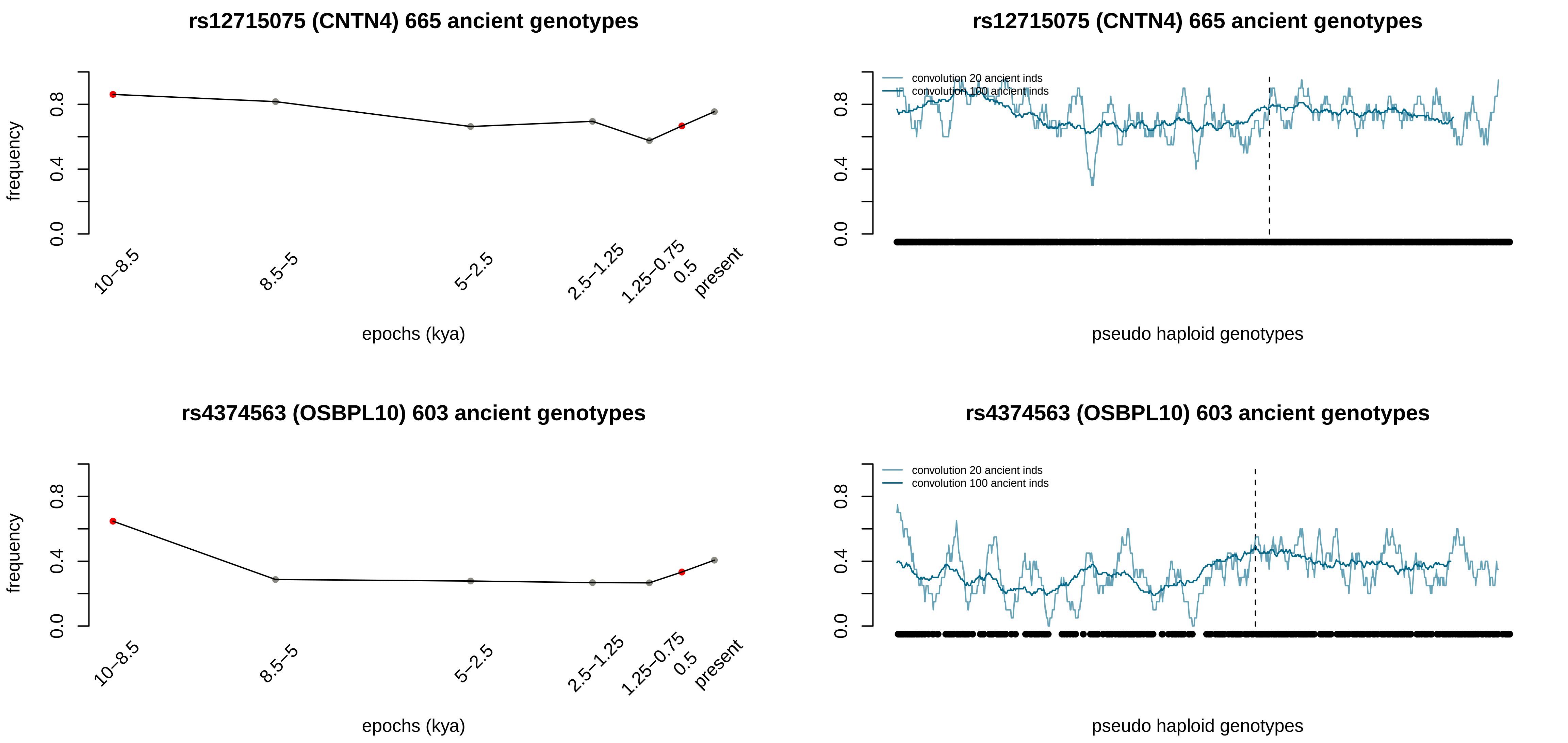

rs12715075 (CNTN4) 665 ancient genotypes
frequency
0.0
0.4
0.8
10–8.5
8.5–5
5–2.5
2.5–1.25
1.25–0.75
0.5
present
epochs (kya)
rs12715075 (CNTN4) 665 ancient genotypes
convolution 20 ancient inds
convolution 100 ancient inds
pseudo haploid genotypes
rs4374563 (OSBPL10) 603 ancient genotypes
frequency
epochs (kya)
rs4374563 (OSBPL10) 603 ancient genotypes
convolution 20 ancient inds
convolution 100 ancient inds
pseudo haploid genotypes

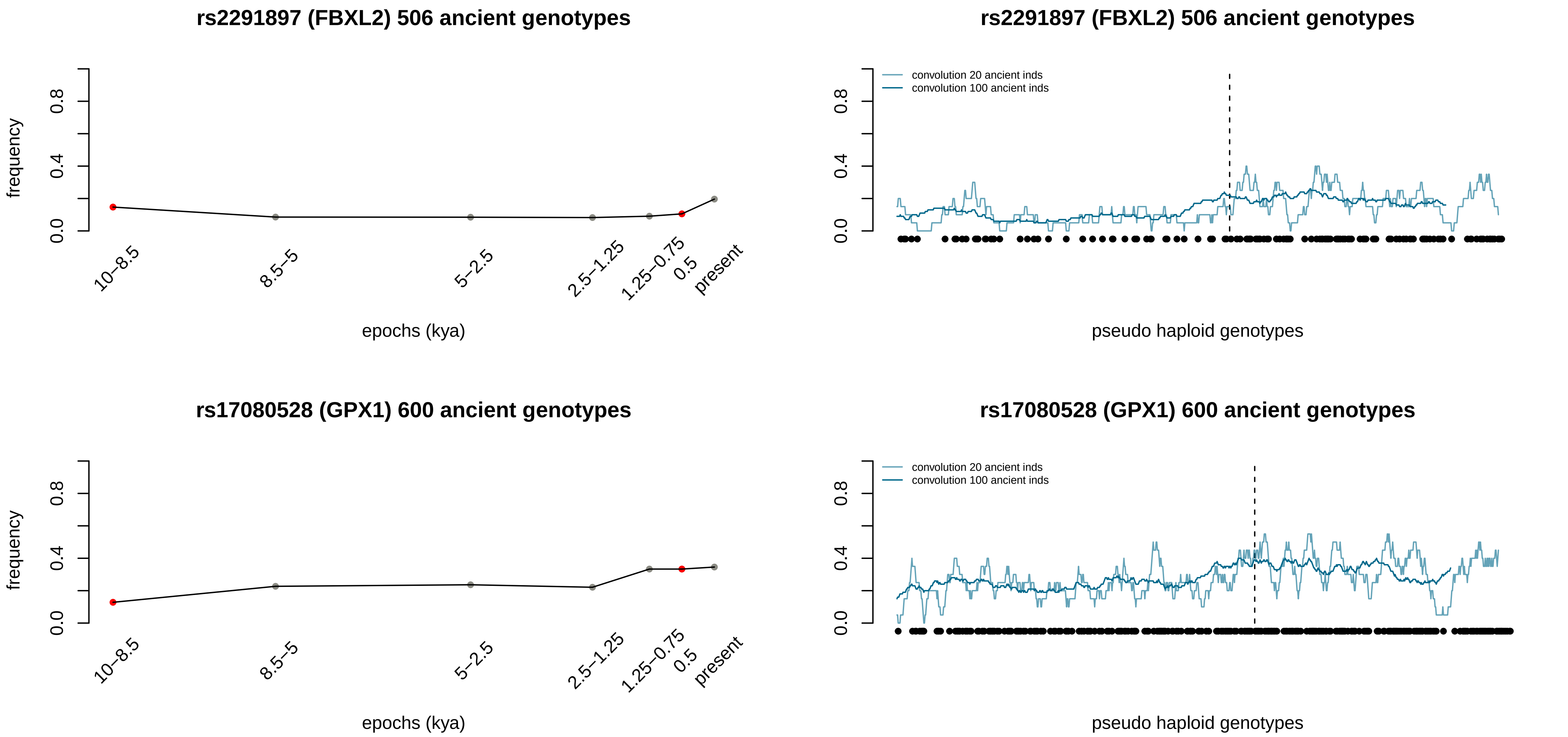
rs2291897 (FBXL2) 506 ancient genotypes
frequency
0.0
0.4
0.8
10–8.5
8.5–5
5–2.5
2.5–1.25
1.25–0.75
0.5
present
epochs (kya)
rs2291897 (FBXL2) 506 ancient genotypes
convolution 20 ancient inds
convolution 100 ancient inds
pseudo haploid genotypes
rs17080528 (GPX1) 600 ancient genotypes
frequency
epochs (kya)
rs17080528 (GPX1) 600 ancient genotypes
convolution 20 ancient inds
convolution 100 ancient inds
pseudo haploid genotypes

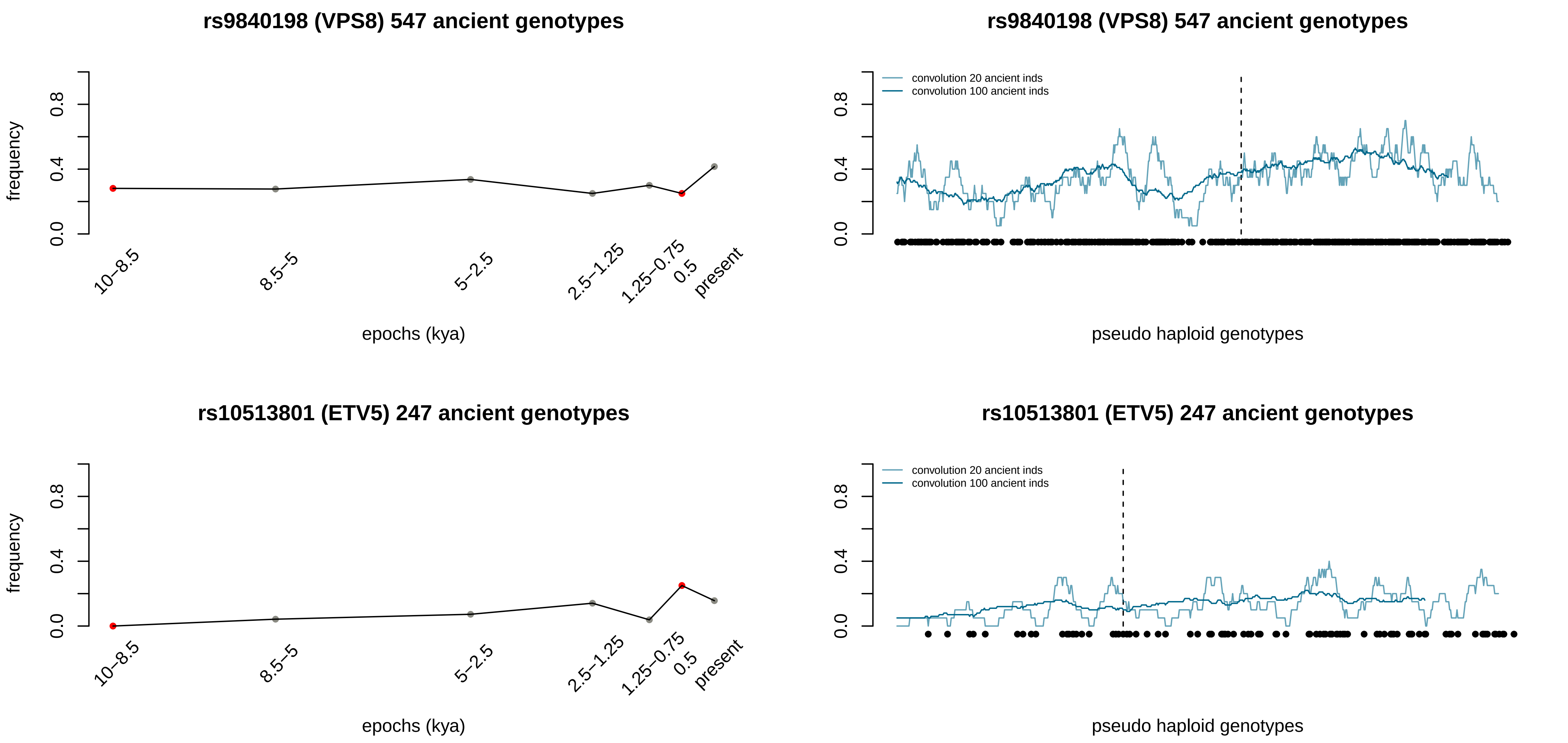
rs9840198 (VPS8) 547 ancient genotypes
frequency
0.0
0.4
0.8
10–8.5
8.5–5
5–2.5
2.5–1.25
1.25–0.75
0.5
present
epochs (kya)
rs9840198 (VPS8) 547 ancient genotypes
convolution 20 ancient inds
convolution 100 ancient inds
pseudo haploid genotypes
rs10513801 (ETV5) 247 ancient genotypes
frequency
epochs (kya)
rs10513801 (ETV5) 247 ancient genotypes
convolution 20 ancient inds
convolution 100 ancient inds
pseudo haploid genotypes

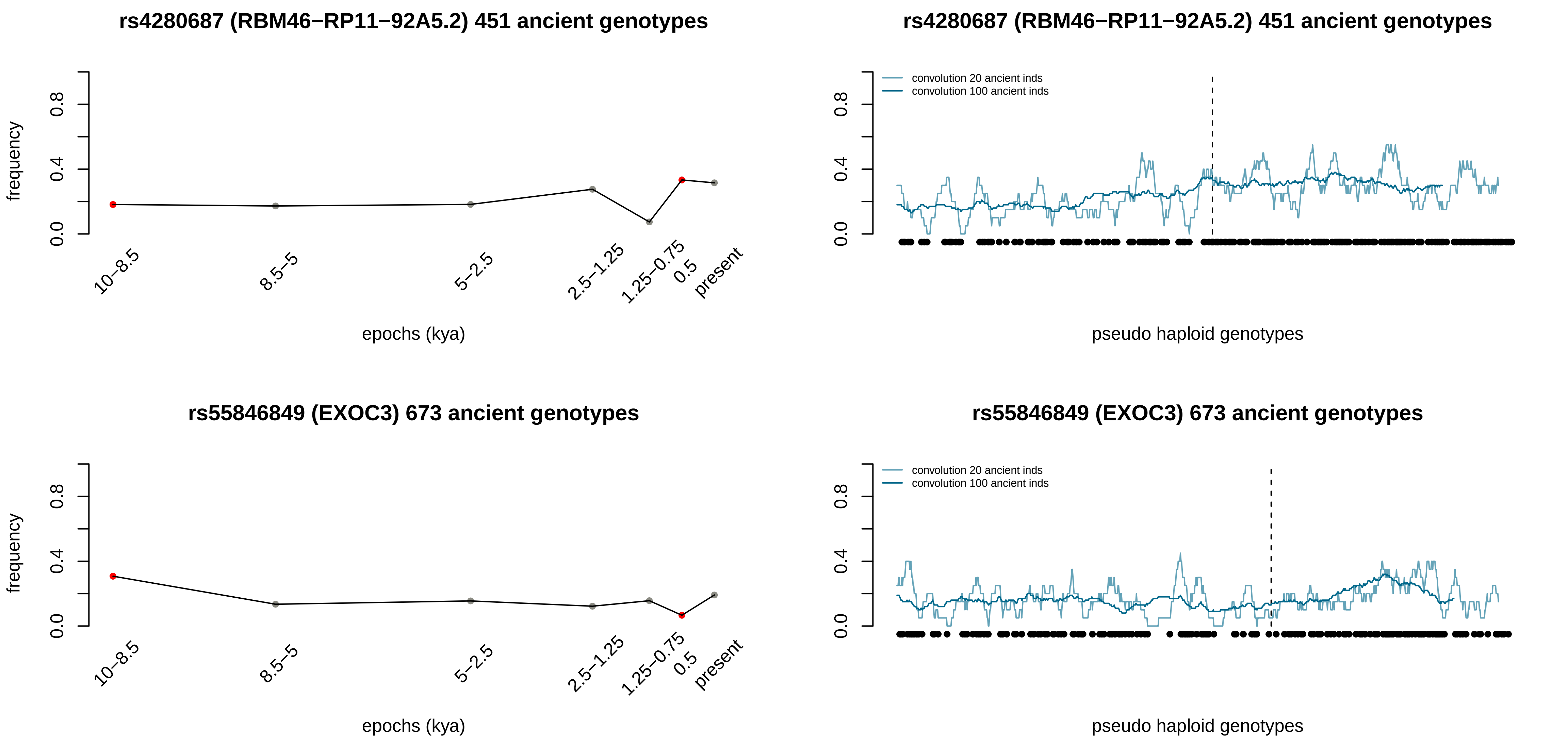

rs4280687 (RBM46−RP11−92A5.2) 451 ancient genotypes
frequency
0.0
0.4
0.8
10−8.5
8.5−5
5−2.5
2.5−1.25
1.25−0.75
0.5
present
epochs (kya)
rs4280687 (RBM46−RP11−92A5.2) 451 ancient genotypes
convolution 20 ancient inds
convolution 100 ancient inds
pseudo haploid genotypes
rs55846849 (EXOC3) 673 ancient genotypes
frequency
epochs (kya)
rs55846849 (EXOC3) 673 ancient genotypes
convolution 20 ancient inds
convolution 100 ancient inds
pseudo haploid genotypes

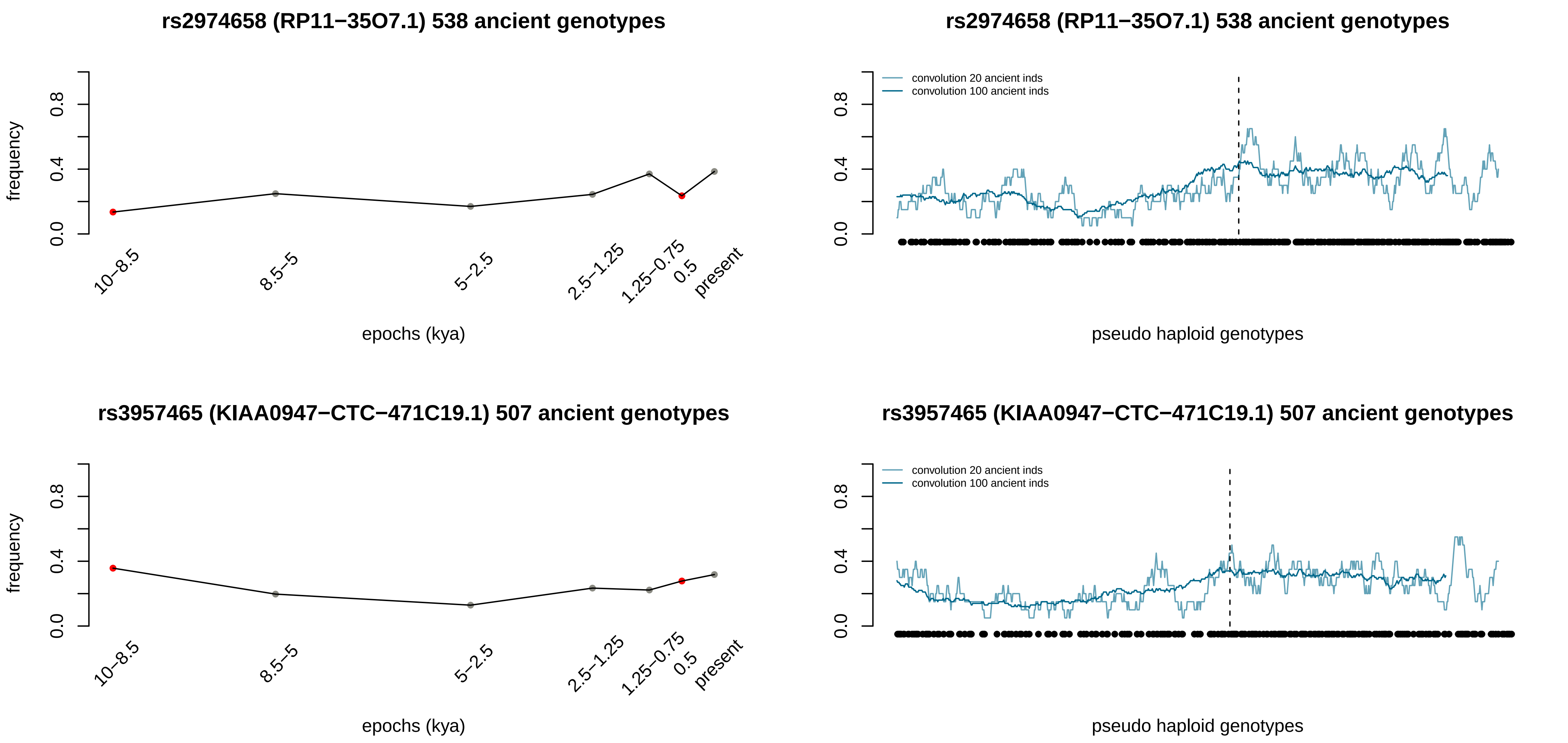

rs2974658 (RP11−35O7.1) 538 ancient genotypes
frequency
0.0
0.4
0.8
10−8.5
8.5−5
5−2.5
2.5−1.25
1.25−0.75
0.5
present
epochs (kya)
rs2974658 (RP11−35O7.1) 538 ancient genotypes
convolution 20 ancient inds
convolution 100 ancient inds
pseudo haploid genotypes
rs3957465 (KIAA0947−CTC−471C19.1) 507 ancient genotypes
frequency
epochs (kya)
rs3957465 (KIAA0947−CTC−471C19.1) 507 ancient genotypes
convolution 20 ancient inds
convolution 100 ancient inds
pseudo haploid genotypes

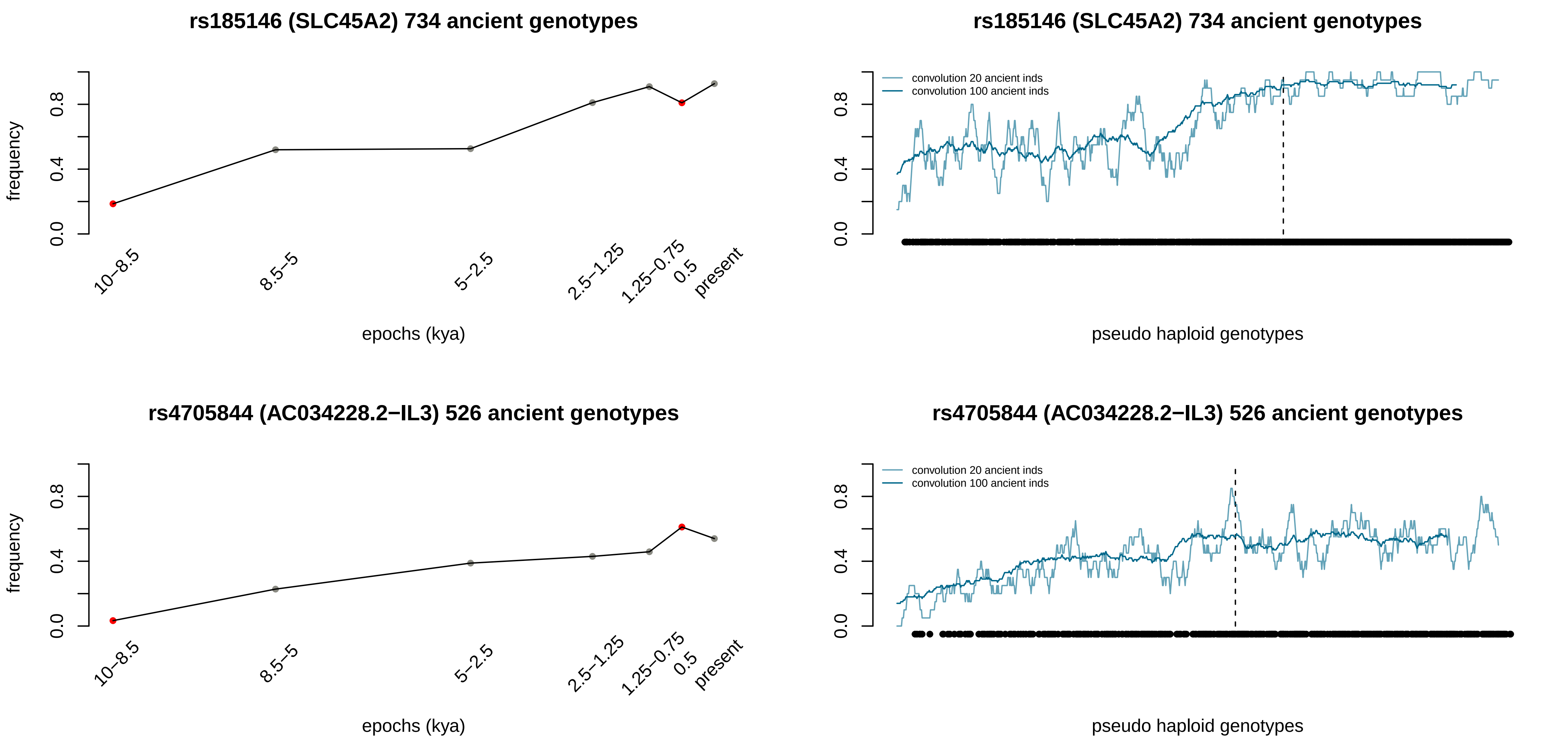
rs185146 (SLC45A2) 734 ancient genotypes
frequency
epochs (kya)
10–8.5
8.5–5
5–2.5
2.5–1.25
1.25–0.75
0.5
present
convolution 20 ancient inds
convolution 100 ancient inds
pseudo haploid genotypes
rs4705844 (AC034228.2−IL3) 526 ancient genotypes

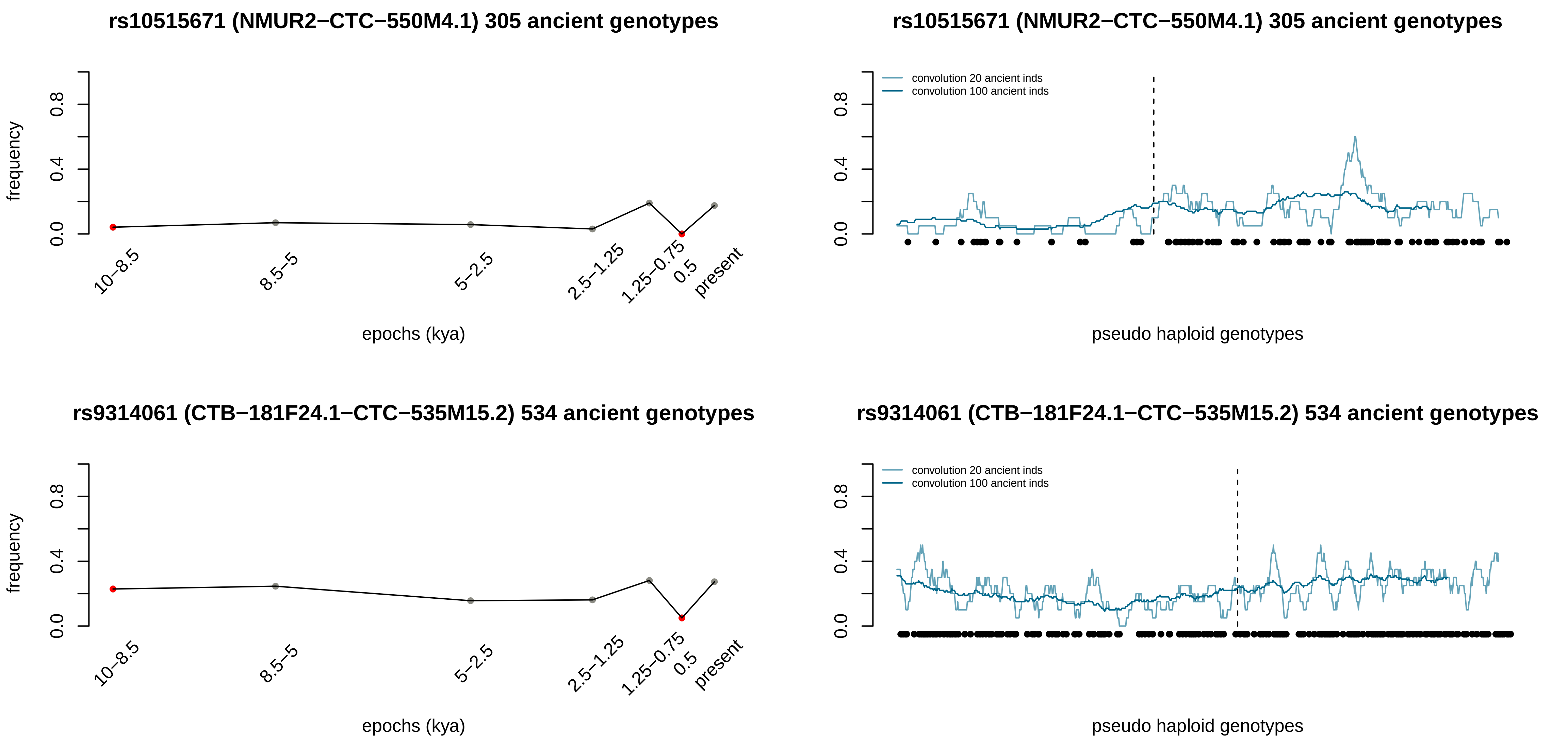

rs10515671 (NMUR2−CTC−550M4.1) 305 ancient genotypes
frequency
0.0
0.4
0.8
10−8.5
8.5−5
5−2.5
2.5−1.25
1.25−0.75
0.5
present
epochs (kya)
rs10515671 (NMUR2−CTC−550M4.1) 305 ancient genotypes
convolution 20 ancient inds
convolution 100 ancient inds
0.0
0.4
0.8
pseudo haploid genotypes
rs9314061 (CTB−181F24.1−CTC−535M15.2) 534 ancient genotypes
frequency
0.0
0.4
0.8
10−8.5
8.5−5
5−2.5
2.5−1.25
1.25−0.75
0.5
present
epochs (kya)
rs9314061 (CTB−181F24.1−CTC−535M15.2) 534 ancient genotypes
convolution 20 ancient inds
convolution 100 ancient inds
0.0
0.4
0.8
pseudo haploid genotypes

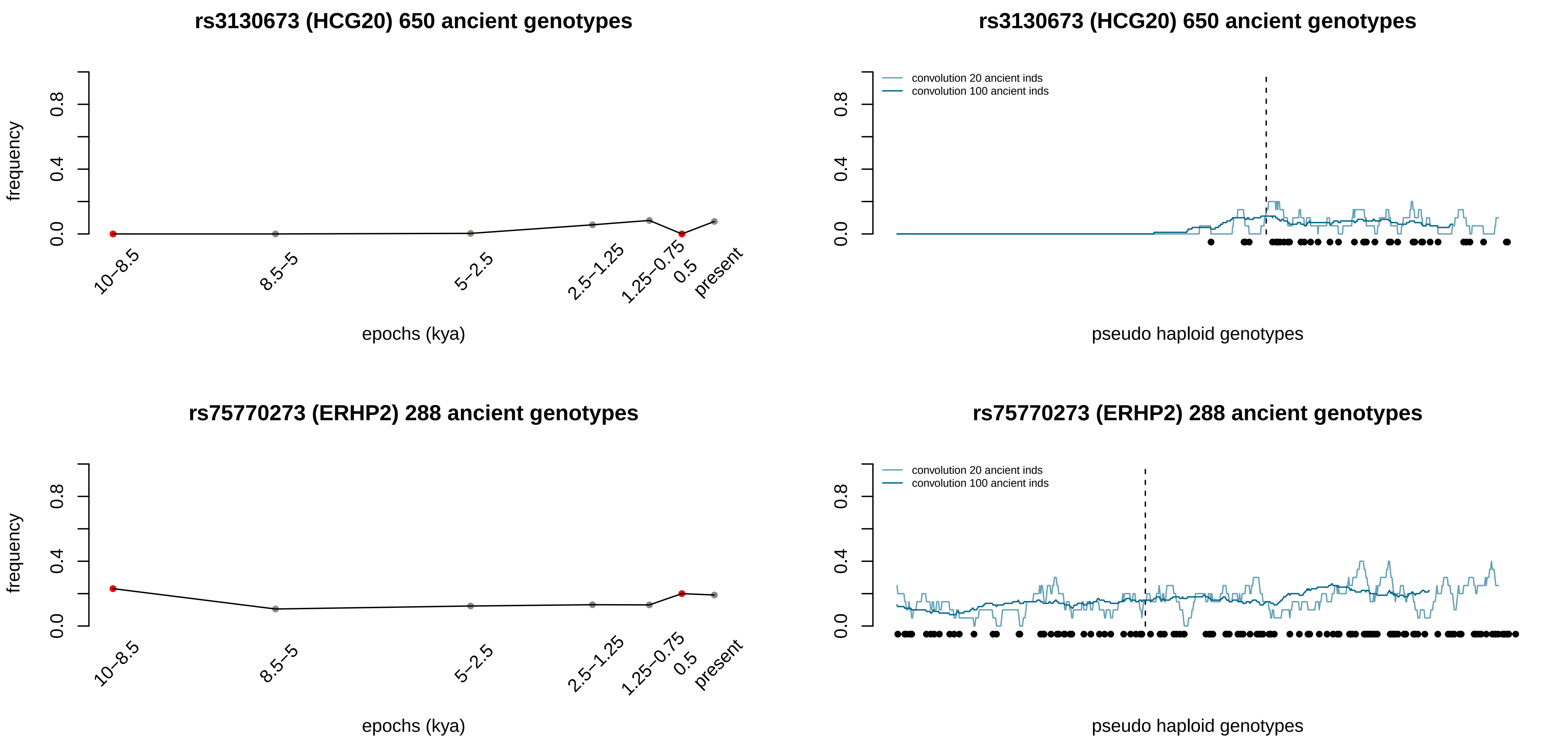
rs3130673 (HCG20) 650 ancient genotypes
frequency
epochs (kya)
10–8.5
8.5–5
5–2.5
2.5–1.25
1.25–0.75
0.5
present
0.0
0.4
0.8
convolution 20 ancient inds
convolution 100 ancient inds
pseudo haploid genotypes
rs75770273 (ERHP2) 288 ancient genotypes

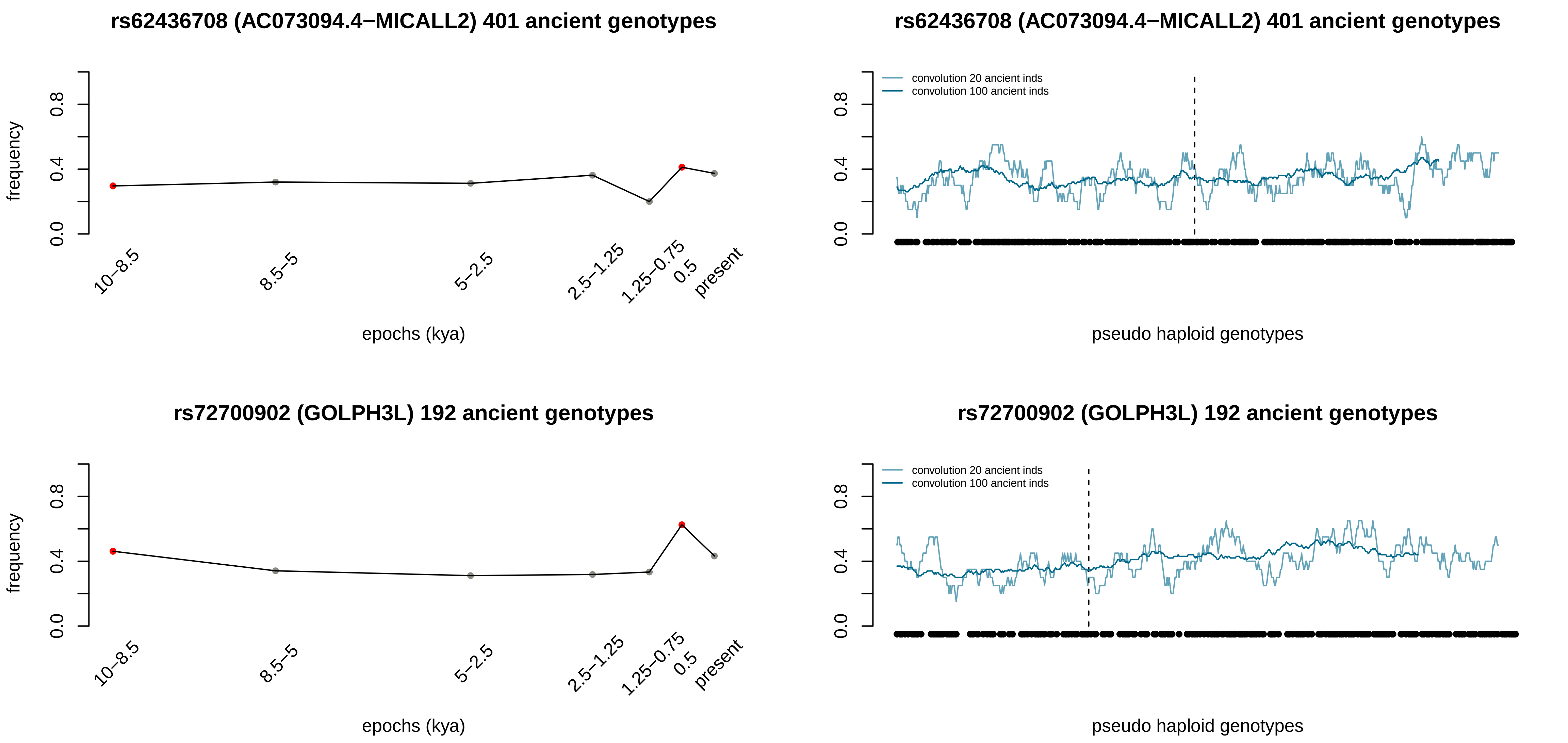
rs62436708 (AC073094.4−MICALL2) 401 ancient genotypes
frequency
0.0
0.4
0.8
10−8.5
8.5−5
5−2.5
2.5−1.25
1.25−0.75
0.5
present
epochs (kya)
rs62436708 (AC073094.4−MICALL2) 401 ancient genotypes
convolution 20 ancient inds
convolution 100 ancient inds
0.0
0.4
0.8
pseudo haploid genotypes
rs72700902 (GOLPH3L) 192 ancient genotypes
frequency
0.0
0.4
0.8
10−8.5
8.5−5
5−2.5
2.5−1.25
1.25−0.75
0.5
present
epochs (kya)
rs72700902 (GOLPH3L) 192 ancient genotypes
convolution 20 ancient inds
convolution 100 ancient inds
0.0
0.4
0.8
pseudo haploid genotypes

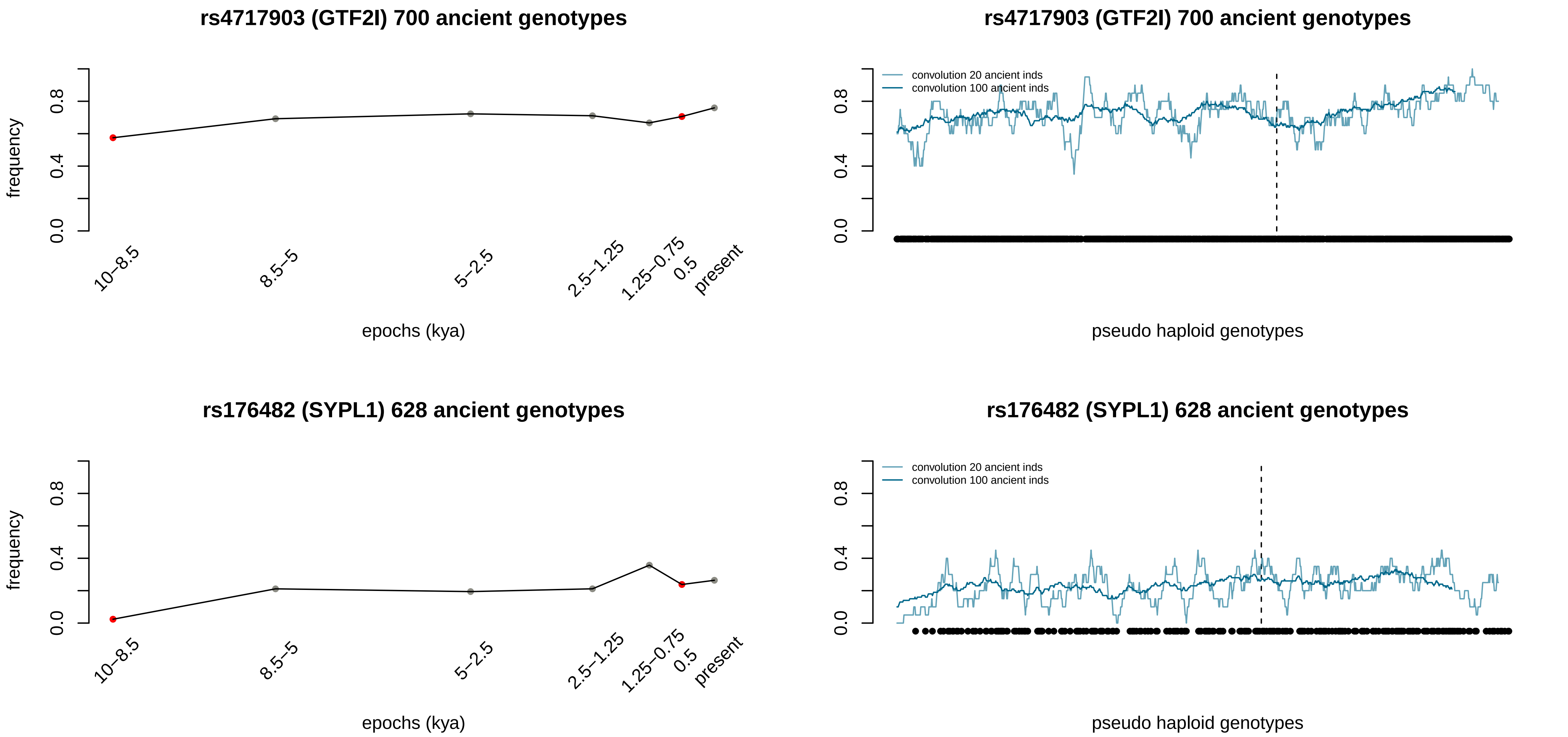
rs4717903 (GTF2I) 700 ancient genotypes
frequency
0.0
0.4
0.8
10–8.5
8.5–5
5–2.5
2.5–1.25
1.25–0.75
0.5
present
epochs (kya)
rs4717903 (GTF2I) 700 ancient genotypes
convolution 20 ancient inds
convolution 100 ancient inds
pseudo haploid genotypes
rs176482 (SYPL1) 628 ancient genotypes
frequency
epochs (kya)
rs176482 (SYPL1) 628 ancient genotypes
convolution 20 ancient inds
convolution 100 ancient inds
pseudo haploid genotypes

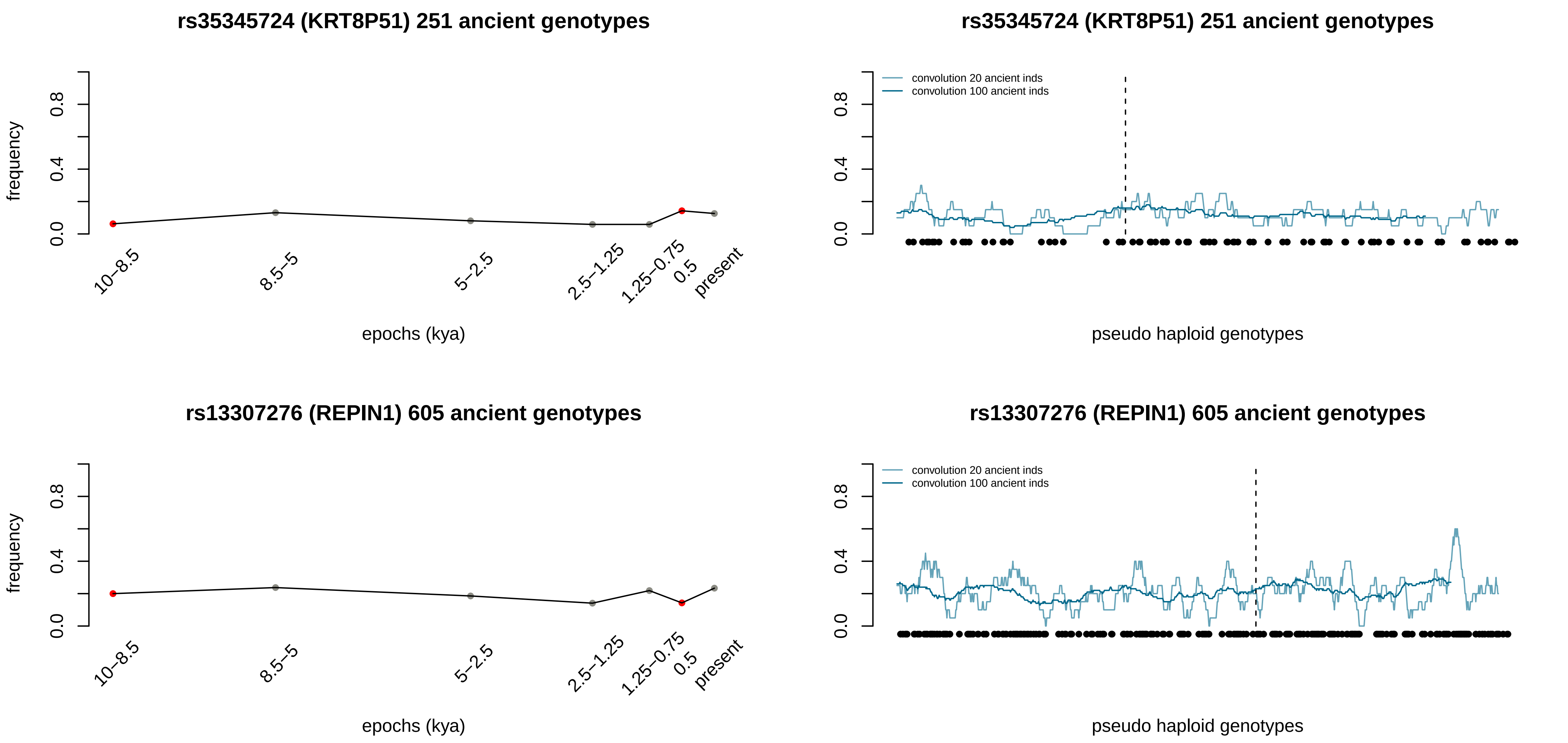
rs35345724 (KRT8P51) 251 ancient genotypes
frequency
0.0
0.4
0.8
10–8.5
8.5–5
5–2.5
2.5–1.25
1.25–0.75
0.5
present
epochs (kya)
rs35345724 (KRT8P51) 251 ancient genotypes
convolution 20 ancient inds
convolution 100 ancient inds
pseudo haploid genotypes
rs13307276 (REPIN1) 605 ancient genotypes
frequency
epochs (kya)
rs13307276 (REPIN1) 605 ancient genotypes
convolution 20 ancient inds
convolution 100 ancient inds
pseudo haploid genotypes

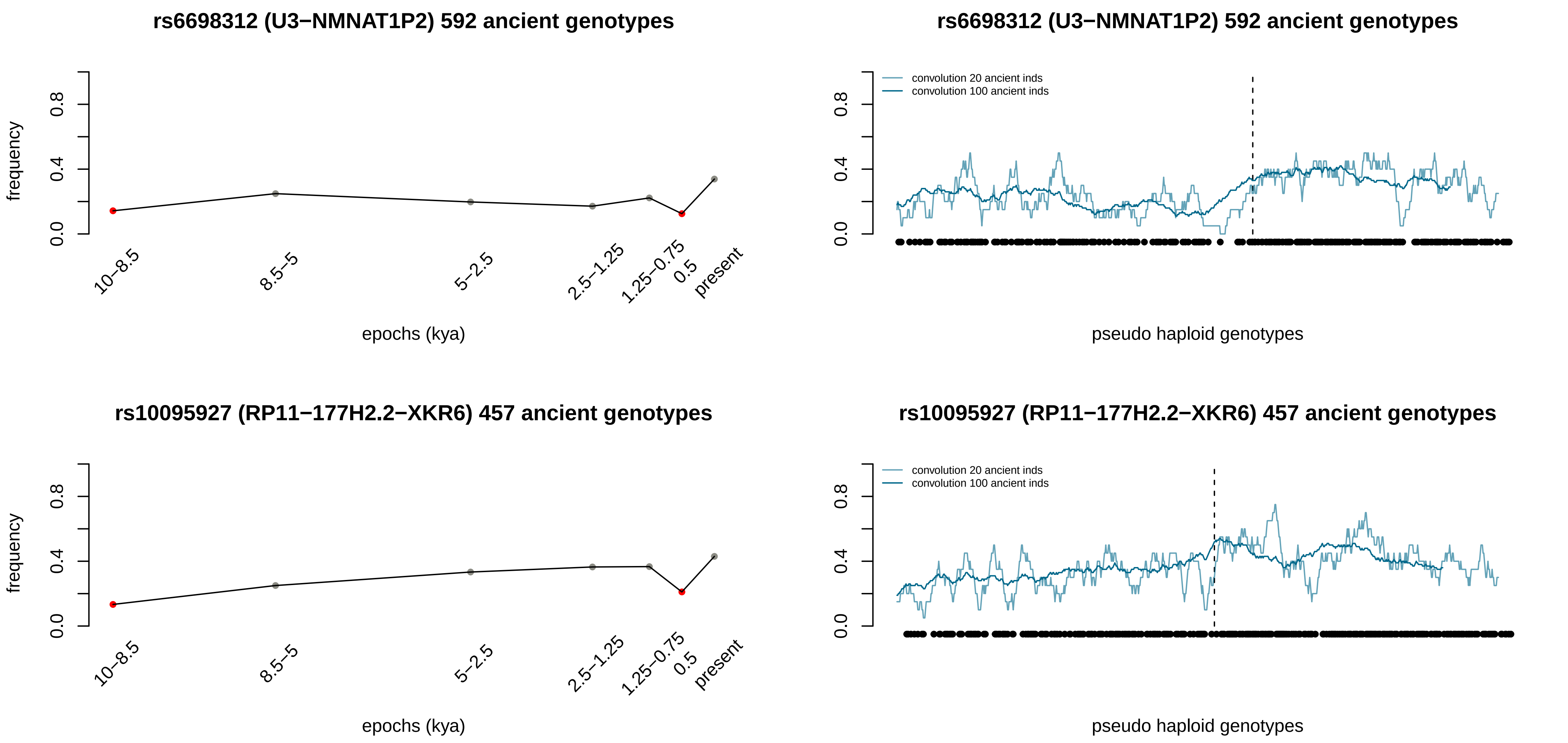
rs6698312 (U3−NMNAT1P2) 592 ancient genotypes
frequency
0.0
0.4
0.8
10−8.5
8.5−5
5−2.5
2.5−1.25
1.25−0.75
0.5
present
epochs (kya)
rs6698312 (U3−NMNAT1P2) 592 ancient genotypes
convolution 20 ancient inds
convolution 100 ancient inds
0.0
0.4
0.8
pseudo haploid genotypes
rs10095927 (RP11−177H2.2−XKR6) 457 ancient genotypes
frequency
0.0
0.4
0.8
10−8.5
8.5−5
5−2.5
2.5−1.25
1.25−0.75
0.5
present
epochs (kya)
rs10095927 (RP11−177H2.2−XKR6) 457 ancient genotypes
convolution 20 ancient inds
convolution 100 ancient inds
0.0
0.4
0.8
pseudo haploid genotypes

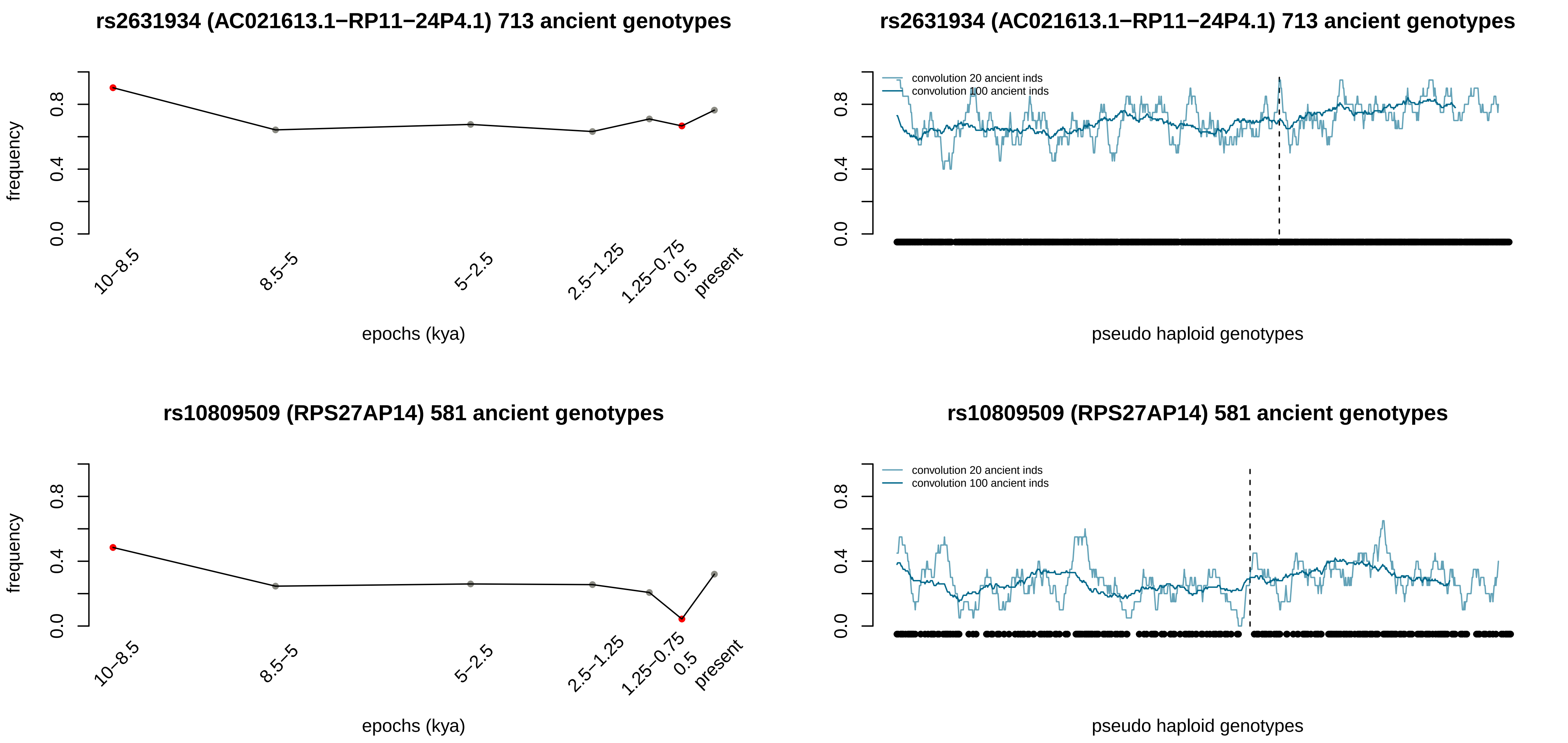
rs2631934 (AC021613.1−RP11−24P4.1) 713 ancient genotypes
frequency
0.0
0.4
0.8
10−8.5
8.5−5
5−2.5
2.5−1.25
1.25−0.75
0.5
present
epochs (kya)
rs2631934 (AC021613.1−RP11−24P4.1) 713 ancient genotypes
convolution 20 ancient inds
convolution 100 ancient inds
0.0
0.4
0.8
pseudo haploid genotypes
rs10809509 (RPS27AP14) 581 ancient genotypes
frequency
0.0
0.4
0.8
10−8.5
8.5−5
5−2.5
2.5−1.25
1.25−0.75
0.5
present
epochs (kya)
rs10809509 (RPS27AP14) 581 ancient genotypes
convolution 20 ancient inds
convolution 100 ancient inds
0.0
0.4
0.8
pseudo haploid genotypes

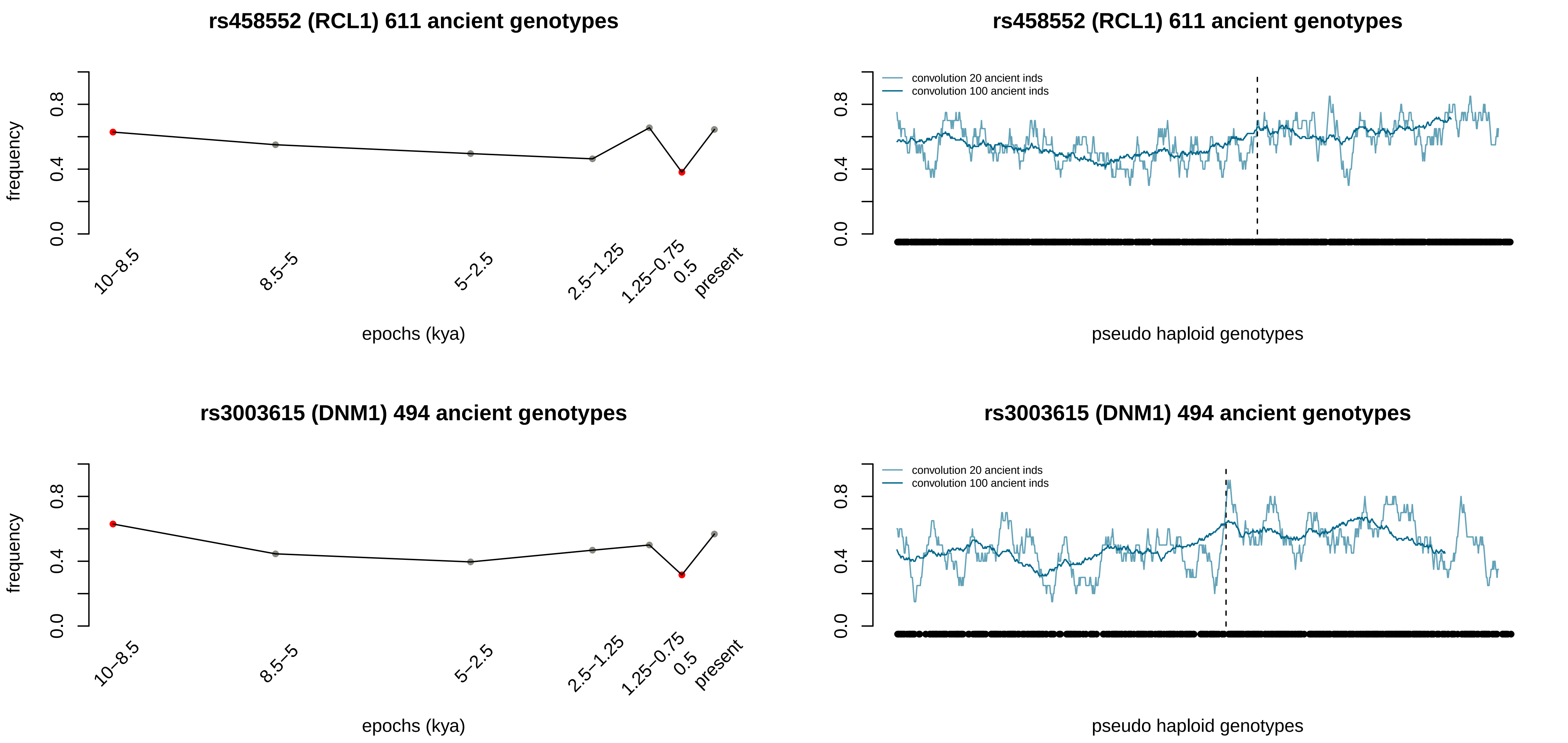
rs458552 (RCL1) 611 ancient genotypes
frequency
0.0
0.4
0.8
10–8.5
8.5–5
5–2.5
2.5–1.25
1.25–0.75
0.5
present
epochs (kya)
rs458552 (RCL1) 611 ancient genotypes
convolution 20 ancient inds
convolution 100 ancient inds
pseudo haploid genotypes
rs3003615 (DNM1) 494 ancient genotypes
frequency
epochs (kya)
rs3003615 (DNM1) 494 ancient genotypes
convolution 20 ancient inds
convolution 100 ancient inds
pseudo haploid genotypes

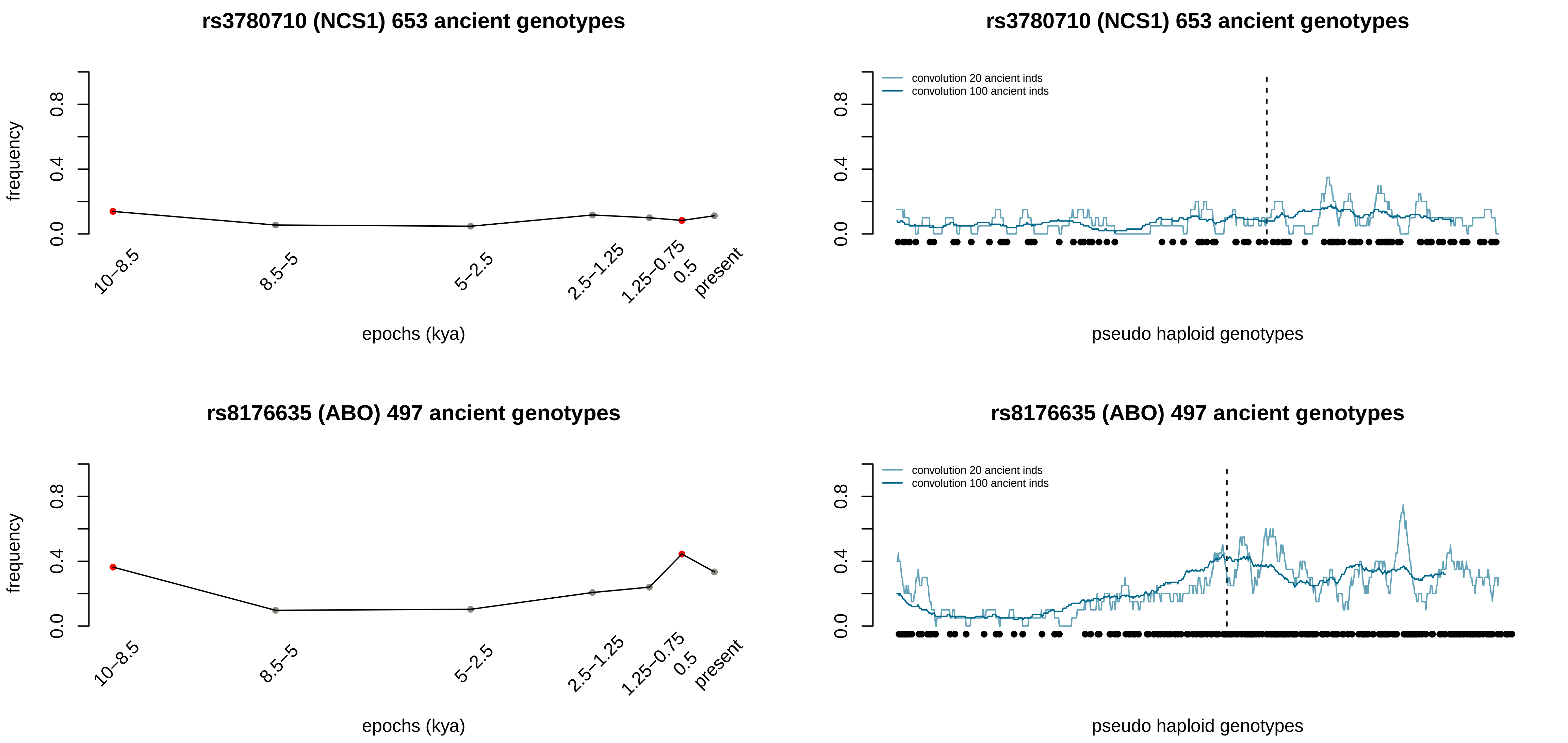
rs3780710 (NCS1) 653 ancient genotypes
frequency
0.0
0.4
0.8
10–8.5
8.5–5
5–2.5
2.5–1.25
1.25–0.75
0.5
present
epochs (kya)
rs3780710 (NCS1) 653 ancient genotypes
convolution 20 ancient inds
convolution 100 ancient inds
pseudo haploid genotypes
rs8176635 (ABO) 497 ancient genotypes
frequency
epochs (kya)
rs8176635 (ABO) 497 ancient genotypes
convolution 20 ancient inds
convolution 100 ancient inds
pseudo haploid genotypes

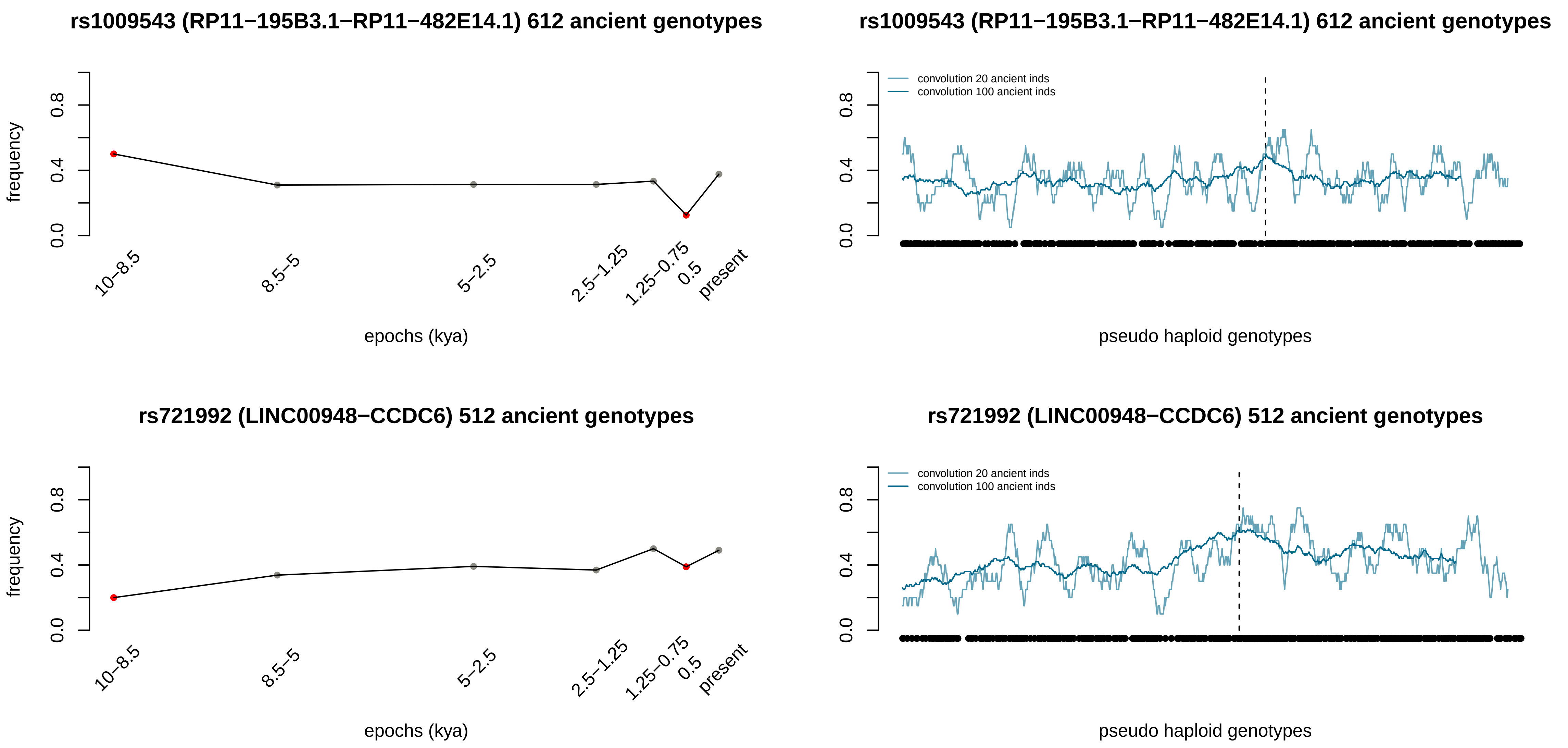
rs1009543 (RP11−195B3.1−RP11−482E14.1) 612 ancient genotypes
frequency
0.0
0.4
0.8
10−8.5
8.5−5
5−2.5
2.5−1.25
1.25−0.75
0.5
present
epochs (kya)
rs1009543 (RP11−195B3.1−RP11−482E14.1) 612 ancient genotypes
convolution 20 ancient inds
convolution 100 ancient inds
0.0
0.4
0.8
pseudo haploid genotypes
rs721992 (LINC00948−CCDC6) 512 ancient genotypes
frequency
0.0
0.4
0.8
10−8.5
8.5−5
5−2.5
2.5−1.25
1.25−0.75
0.5
present
epochs (kya)
rs721992 (LINC00948−CCDC6) 512 ancient genotypes
convolution 20 ancient inds
convolution 100 ancient inds
0.0
0.4
0.8
pseudo haploid genotypes

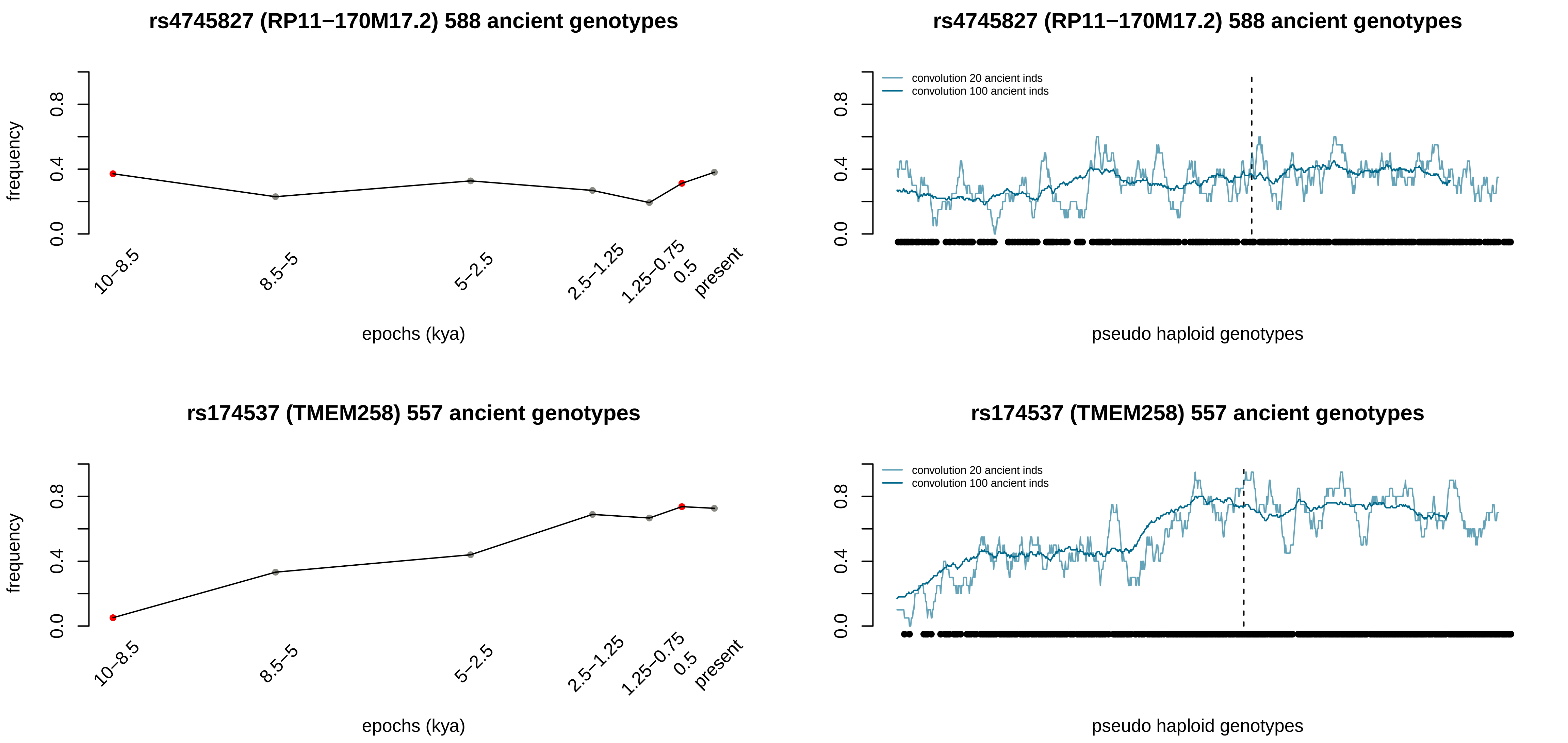

rs4745827 (RP11−170M17.2) 588 ancient genotypes
frequency
0.0
0.4
0.8
10−8.5
8.5−5
5−2.5
2.5−1.25
1.25−0.75
0.5
present
epochs (kya)
rs4745827 (RP11−170M17.2) 588 ancient genotypes
convolution 20 ancient inds
convolution 100 ancient inds
pseudo haploid genotypes
rs174537 (TMEM258) 557 ancient genotypes
frequency
epochs (kya)
rs174537 (TMEM258) 557 ancient genotypes
convolution 20 ancient inds
convolution 100 ancient inds
pseudo haploid genotypes

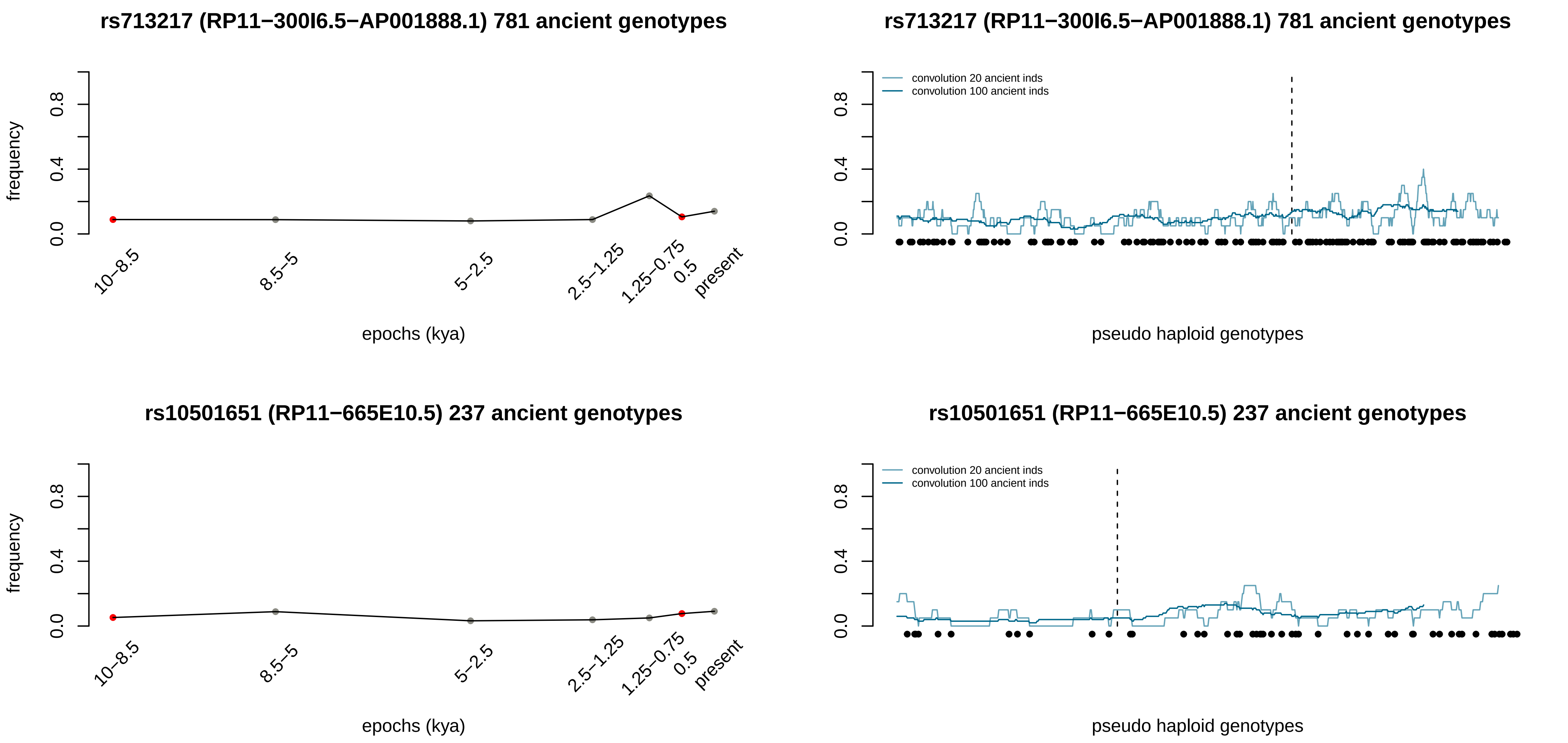
rs713217 (RP11−300I6.5−AP001888.1) 781 ancient genotypes
frequency
0.0
0.4
0.8
10−8.5
8.5−5
5−2.5
2.5−1.25
1.25−0.75
0.5
present
epochs (kya)
rs713217 (RP11−300I6.5−AP001888.1) 781 ancient genotypes
convolution 20 ancient inds
convolution 100 ancient inds
pseudo haploid genotypes
rs10501651 (RP11−665E10.5) 237 ancient genotypes
frequency
epochs (kya)
rs10501651 (RP11−665E10.5) 237 ancient genotypes
convolution 20 ancient inds
convolution 100 ancient inds
pseudo haploid genotypes

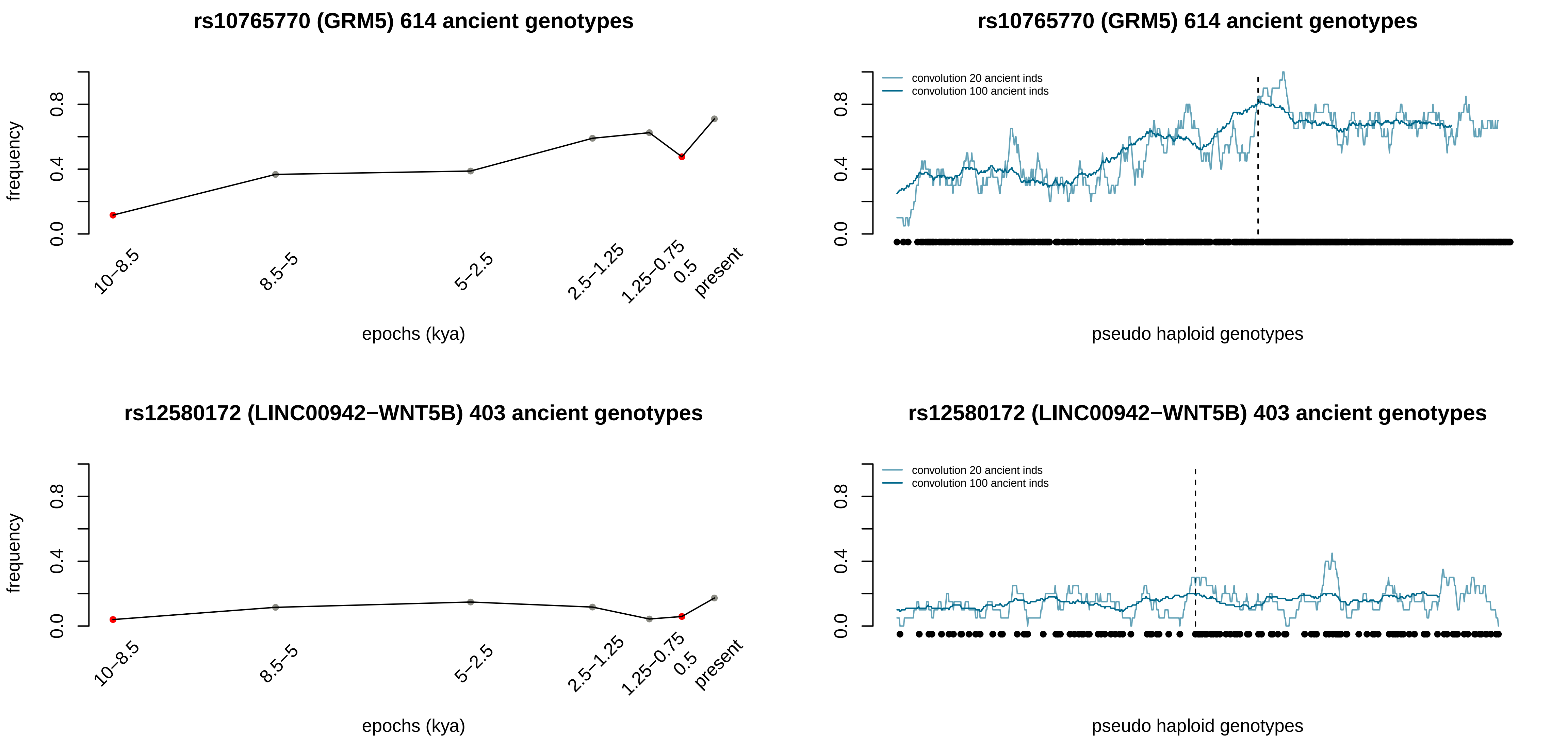
rs10765770 (GRM5) 614 ancient genotypes
frequency
0.0
0.4
0.8
10–8.5
8.5–5
5–2.5
2.5–1.25
1.25–0.75
0.5
present
epochs (kya)
rs10765770 (GRM5) 614 ancient genotypes
convolution 20 ancient inds
convolution 100 ancient inds
0.0
0.4
0.8
pseudo haploid genotypes
rs12580172 (LINC00942−WNT5B) 403 ancient genotypes
frequency
0.0
0.4
0.8
10–8.5
8.5–5
5–2.5
2.5–1.25
1.25–0.75
0.5
present
epochs (kya)
rs12580172 (LINC00942−WNT5B) 403 ancient genotypes
convolution 20 ancient inds
convolution 100 ancient inds
0.0
0.4
0.8
pseudo haploid genotypes

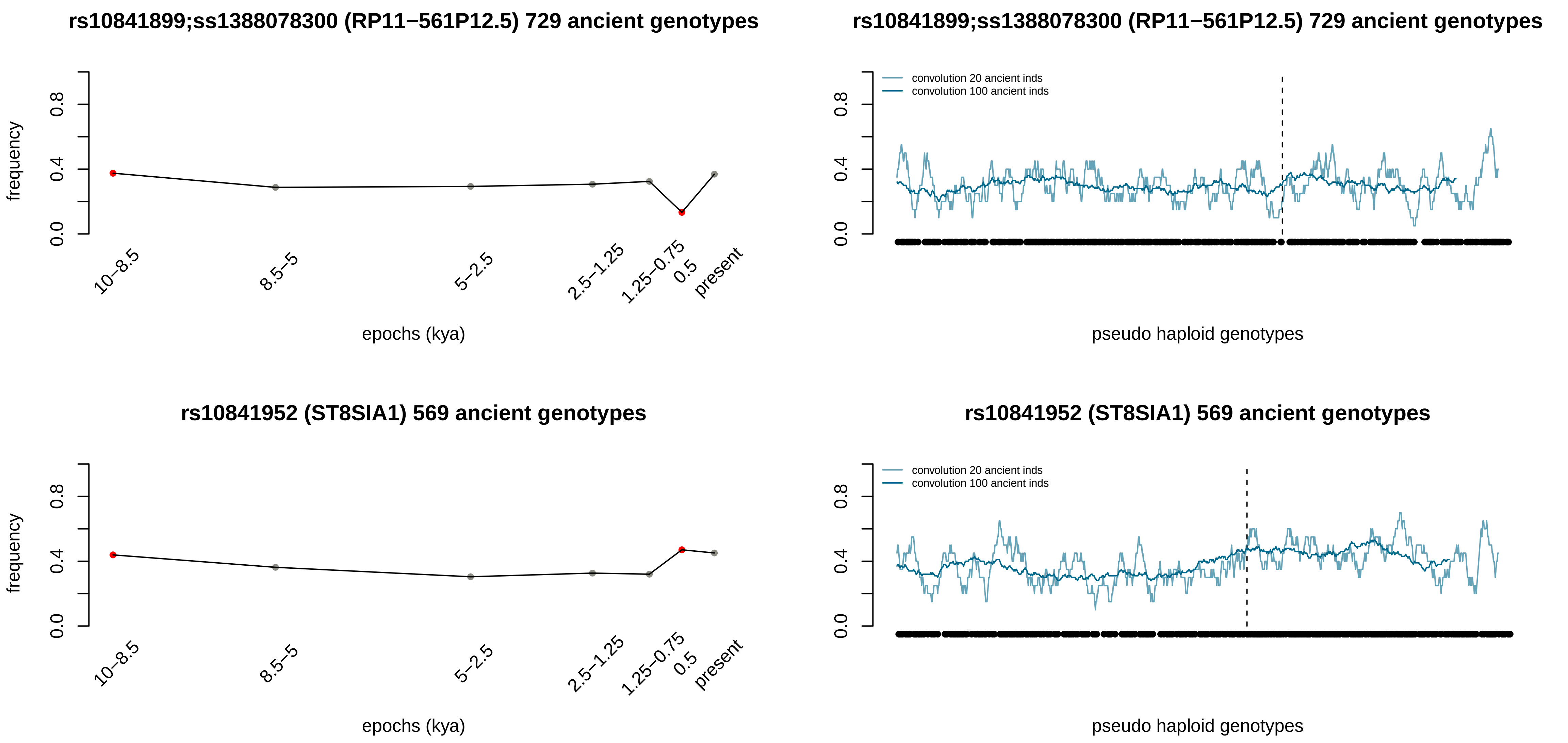
rs10841899;ss1388078300 (RP11−561P12.5) 729 ancient genotypes
frequency
0.0
0.4
0.8
10−8.5
8.5−5
5−2.5
2.5−1.25
1.25−0.75
0.5
present
epochs (kya)
rs10841899;ss1388078300 (RP11−561P12.5) 729 ancient genotypes
convolution 20 ancient inds
convolution 100 ancient inds
0.0
0.4
0.8
pseudo haploid genotypes
rs10841952 (ST8SIA1) 569 ancient genotypes
frequency
0.0
0.4
0.8
10−8.5
8.5−5
5−2.5
2.5−1.25
1.25−0.75
0.5
present
epochs (kya)
rs10841952 (ST8SIA1) 569 ancient genotypes
convolution 20 ancient inds
convolution 100 ancient inds
0.0
0.4
0.8
pseudo haploid genotypes

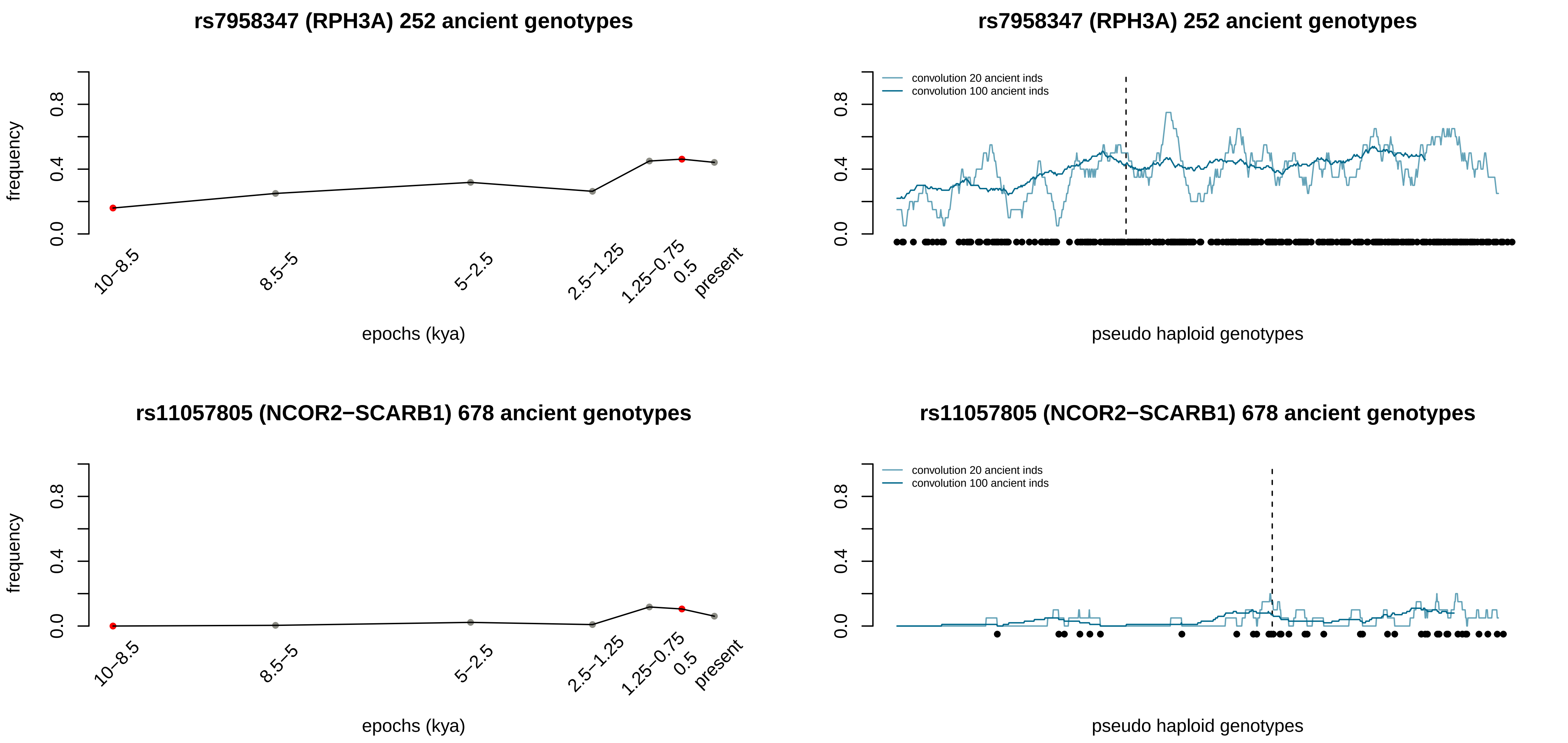
rs7958347 (RPH3A) 252 ancient genotypes
frequency
0.0
0.4
0.8
10−8.5
8.5−5
5−2.5
2.5−1.25
1.25−0.75
0.5
present
epochs (kya)
rs7958347 (RPH3A) 252 ancient genotypes
convolution 20 ancient inds
convolution 100 ancient inds
pseudo haploid genotypes
rs11057805 (NCOR2−SCARB1) 678 ancient genotypes
frequency
epochs (kya)
rs11057805 (NCOR2−SCARB1) 678 ancient genotypes
convolution 20 ancient inds
convolution 100 ancient inds
pseudo haploid genotypes

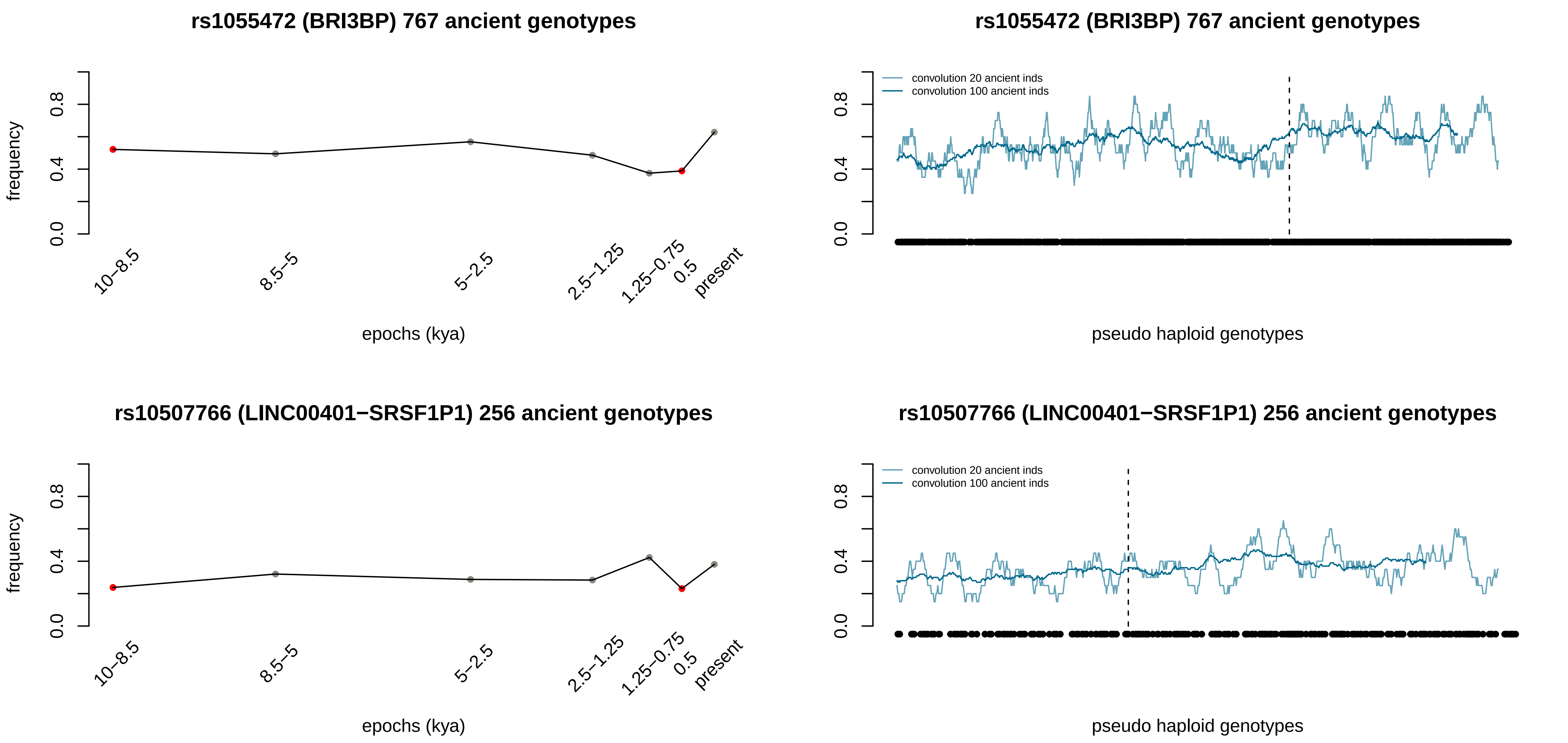

rs1055472 (BRI3BP) 767 ancient genotypes
frequency
0.0
0.4
0.8
10–8.5
8.5–5
5–2.5
2.5–1.25
1.25–0.75
0.5
present
epochs (kya)
rs1055472 (BRI3BP) 767 ancient genotypes
convolution 20 ancient inds
convolution 100 ancient inds
pseudo haploid genotypes
rs10507766 (LINC00401−SRSF1P1) 256 ancient genotypes
frequency
epochs (kya)
rs10507766 (LINC00401−SRSF1P1) 256 ancient genotypes
convolution 20 ancient inds
convolution 100 ancient inds
pseudo haploid genotypes

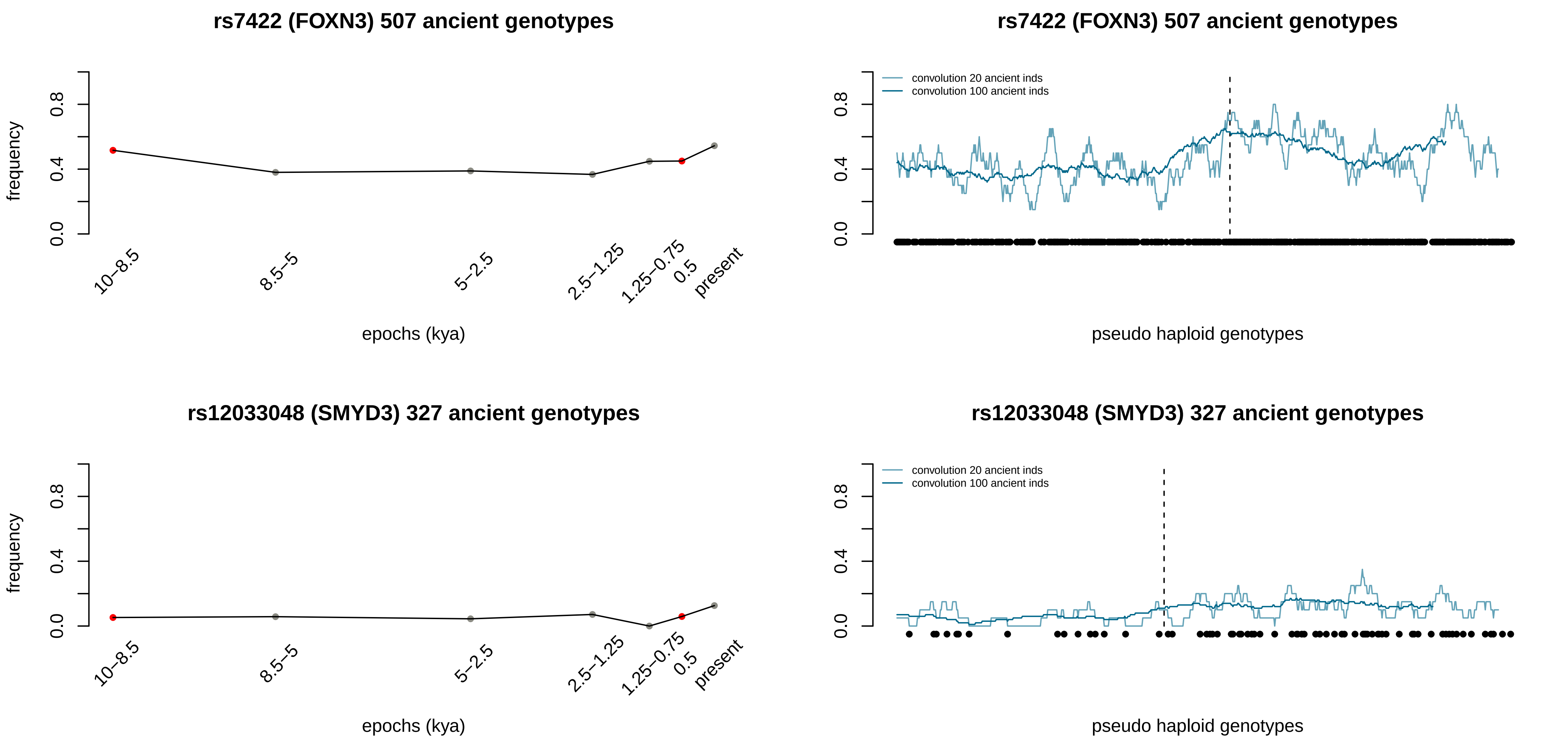
rs7422 (FOXN3) 507 ancient genotypes
frequency
0.0
0.4
0.8
10–8.5
8.5–5
5–2.5
2.5–1.25
1.25–0.75
0.5
present
epochs (kya)
rs7422 (FOXN3) 507 ancient genotypes
convolution 20 ancient inds
convolution 100 ancient inds
0.0
0.4
0.8
pseudo haploid genotypes
rs12033048 (SMYD3) 327 ancient genotypes
frequency
0.0
0.4
0.8
10–8.5
8.5–5
5–2.5
2.5–1.25
1.25–0.75
0.5
present
epochs (kya)
rs12033048 (SMYD3) 327 ancient genotypes
convolution 20 ancient inds
convolution 100 ancient inds
0.0
0.4
0.8
pseudo haploid genotypes

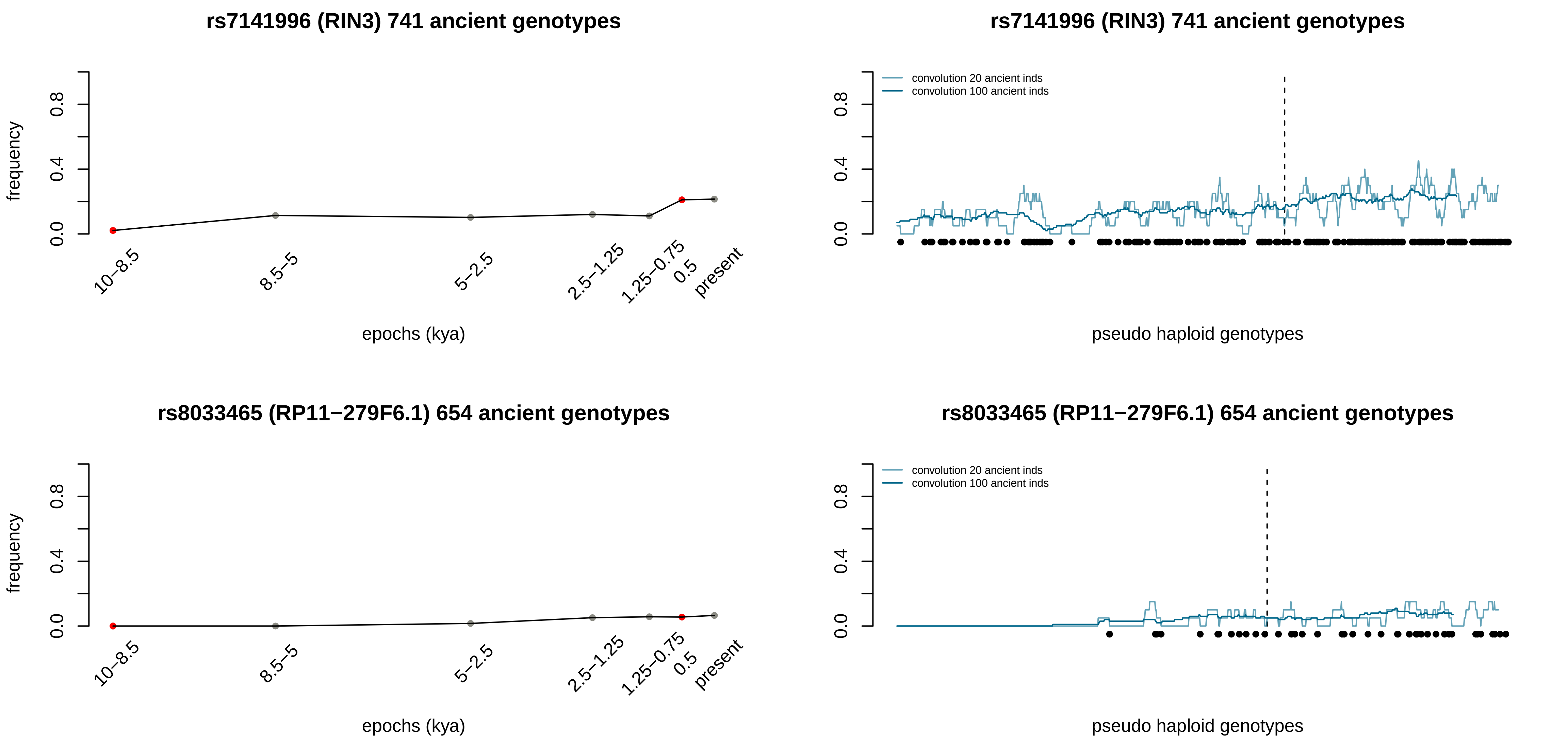
rs7141996 (RIN3) 741 ancient genotypes
frequency
0.0
0.4
0.8
10–8.5
8.5–5
5–2.5
2.5–1.25
1.25–0.75
0.5
present
epochs (kya)
rs7141996 (RIN3) 741 ancient genotypes
convolution 20 ancient inds
convolution 100 ancient inds
pseudo haploid genotypes
rs8033465 (RP11−279F6.1) 654 ancient genotypes
frequency
epochs (kya)
rs8033465 (RP11−279F6.1) 654 ancient genotypes
convolution 20 ancient inds
convolution 100 ancient inds
pseudo haploid genotypes

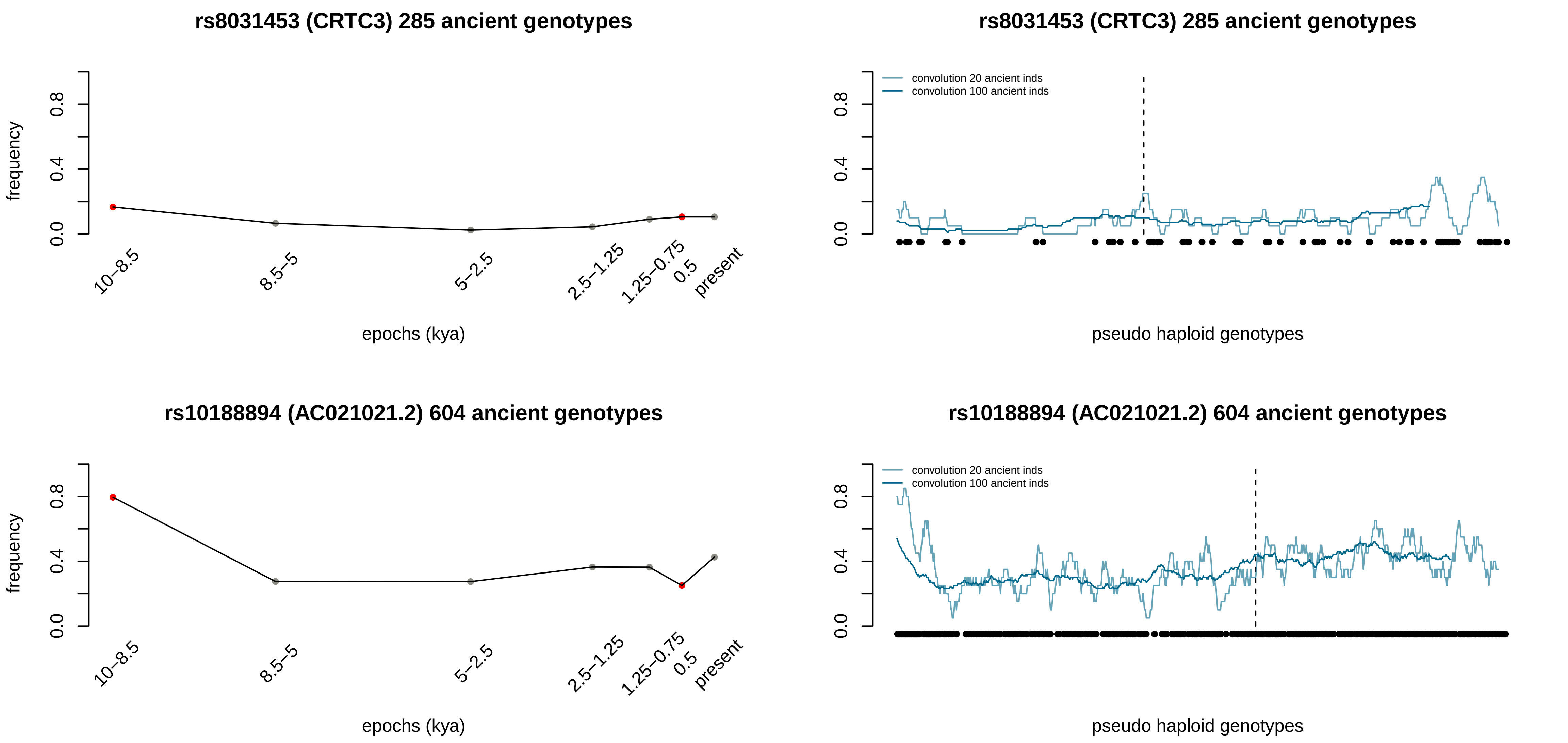
rs8031453 (CRTC3) 285 ancient genotypes
frequency
0.0
0.4
0.8
10–8.5
8.5–5
5–2.5
2.5–1.25
1.25–0.75
0.5
present
epochs (kya)
rs8031453 (CRTC3) 285 ancient genotypes
convolution 20 ancient inds
convolution 100 ancient inds
pseudo haploid genotypes
rs10188894 (AC021021.2) 604 ancient genotypes
frequency
epochs (kya)
rs10188894 (AC021021.2) 604 ancient genotypes
convolution 20 ancient inds
convolution 100 ancient inds
pseudo haploid genotypes

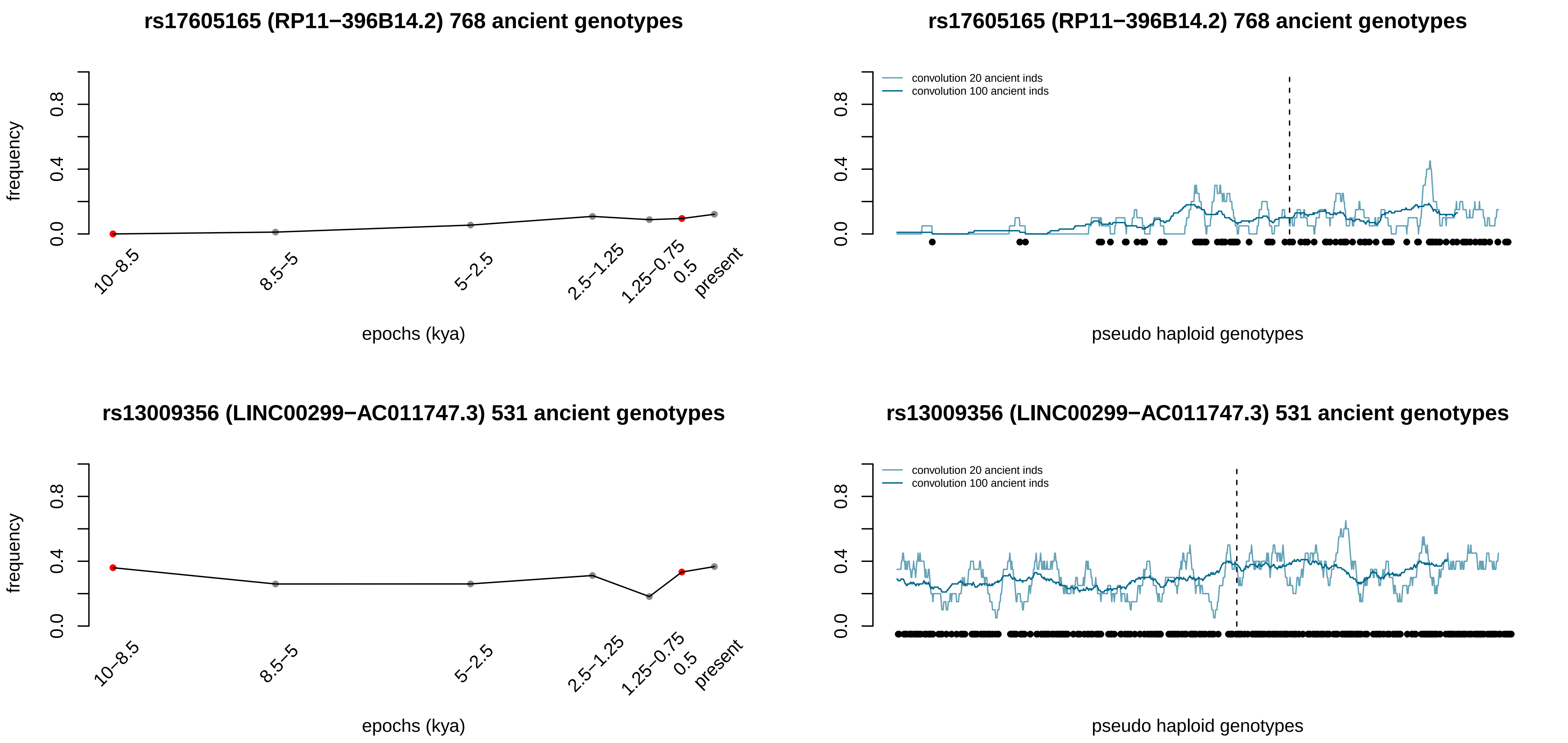

rs17605165 (RP11−396B14.2) 768 ancient genotypes
frequency
0.0
0.4
0.8
10−8.5
8.5−5
5−2.5
2.5−1.25
1.25−0.75
0.5
present
epochs (kya)
rs17605165 (RP11−396B14.2) 768 ancient genotypes
convolution 20 ancient inds
convolution 100 ancient inds
pseudo haploid genotypes
rs13009356 (LINC00299−AC011747.3) 531 ancient genotypes
frequency
epochs (kya)
rs13009356 (LINC00299−AC011747.3) 531 ancient genotypes
convolution 20 ancient inds
convolution 100 ancient inds
pseudo haploid genotypes

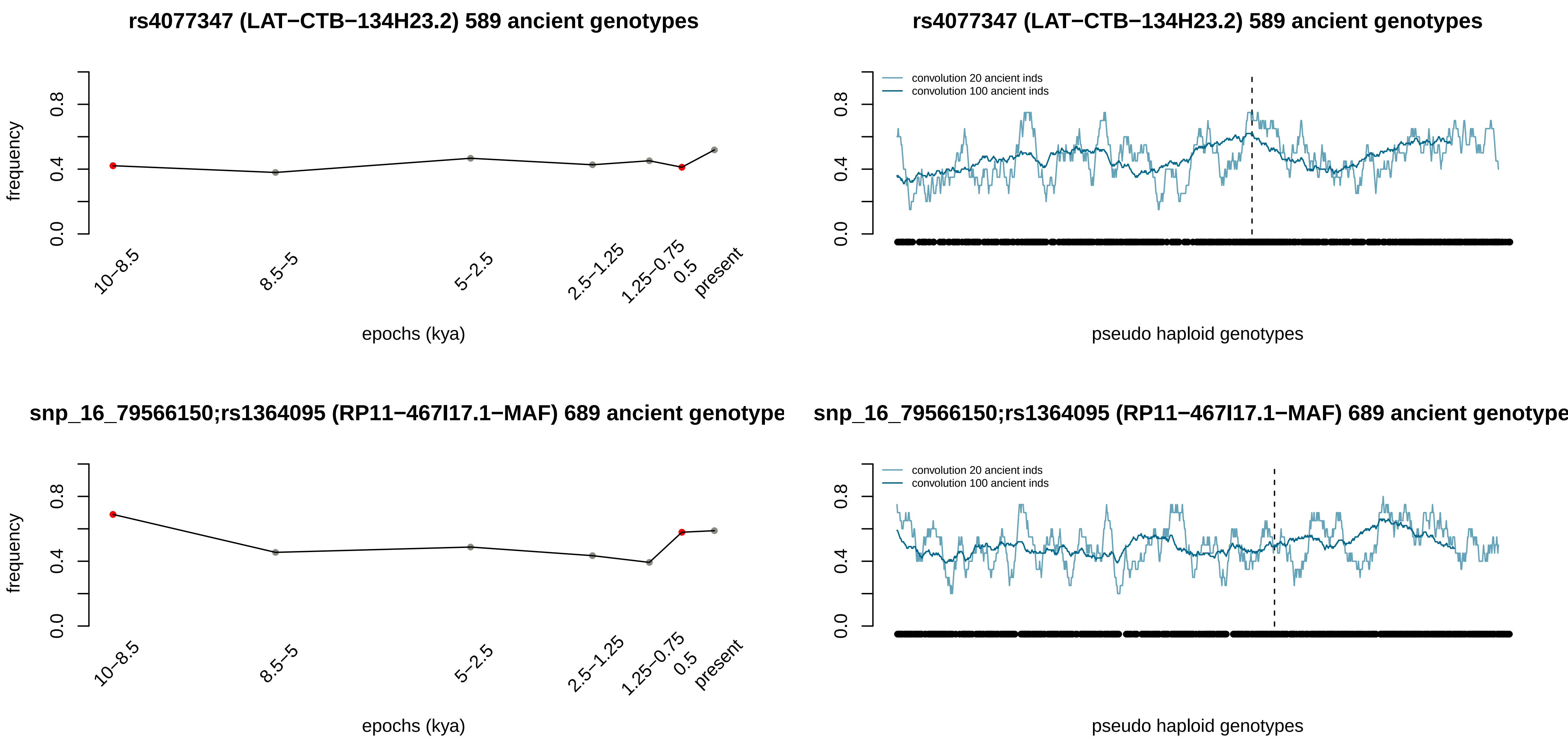
rs4077347 (LAT−CTB−134H23.2) 589 ancient genotypes
frequency
0.0
0.4
0.8
10−8.5
8.5−5
5−2.5
2.5−1.25
1.25−0.75
0.5
present
epochs (kya)
rs4077347 (LAT−CTB−134H23.2) 589 ancient genotypes
convolution 20 ancient inds
convolution 100 ancient inds
0.0
0.4
0.8
pseudo haploid genotypes
snp_16_79566150;rs1364095 (RP11−467I17.1−MAF) 689 ancient genotype
frequency
0.0
0.4
0.8
10−8.5
8.5−5
5−2.5
2.5−1.25
1.25−0.75
0.5
present
epochs (kya)
snp_16_79566150;rs1364095 (RP11−467I17.1−MAF) 689 ancient genotype
convolution 20 ancient inds
convolution 100 ancient inds
0.0
0.4
0.8
pseudo haploid genotypes

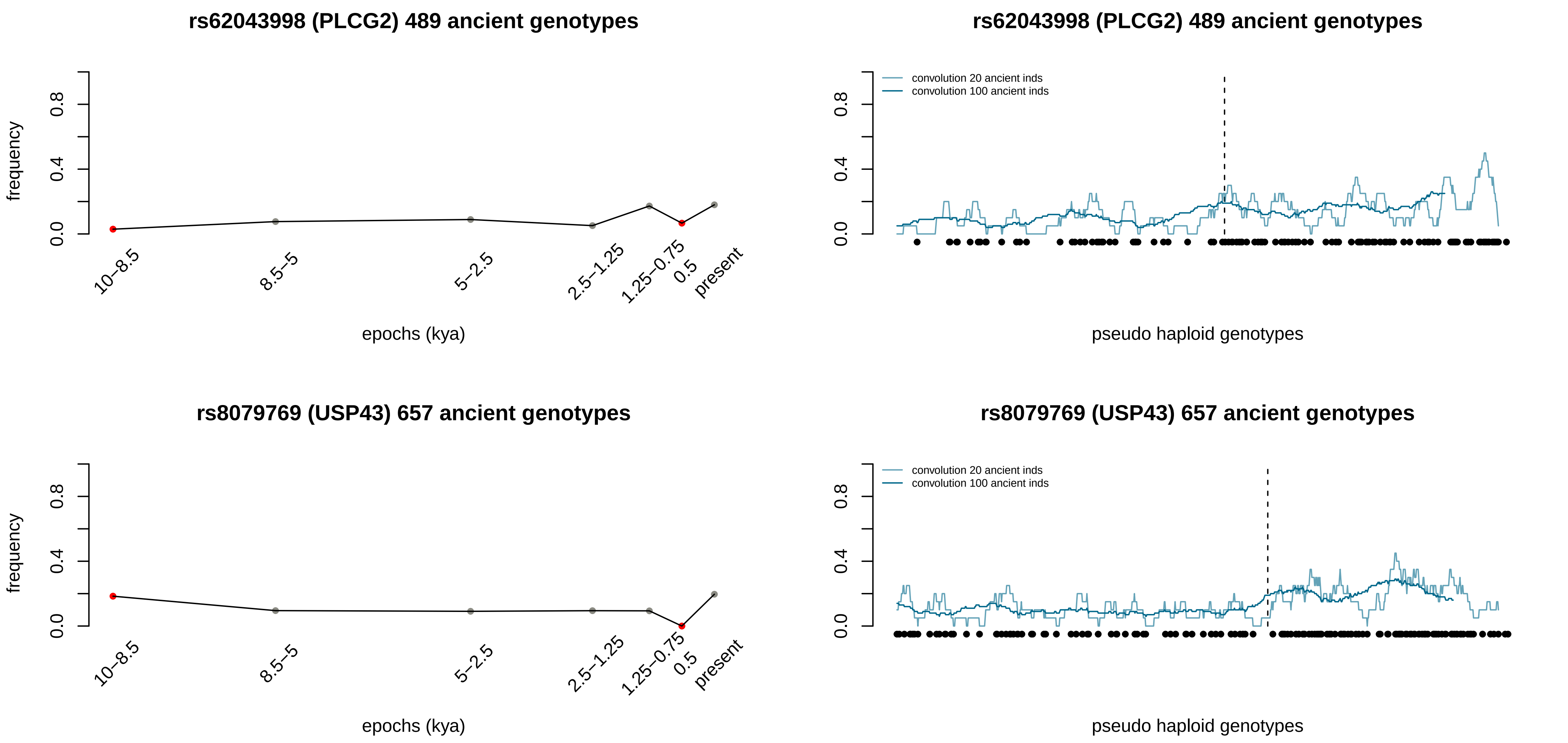

rs62043998 (PLCG2) 489 ancient genotypes
frequency
0.0
0.4
0.8
10–8.5
8.5–5
5–2.5
2.5–1.25
1.25–0.75
0.5
present
epochs (kya)
rs62043998 (PLCG2) 489 ancient genotypes
convolution 20 ancient inds
convolution 100 ancient inds
pseudo haploid genotypes
rs8079769 (USP43) 657 ancient genotypes
frequency
epochs (kya)
rs8079769 (USP43) 657 ancient genotypes
convolution 20 ancient inds
convolution 100 ancient inds
pseudo haploid genotypes

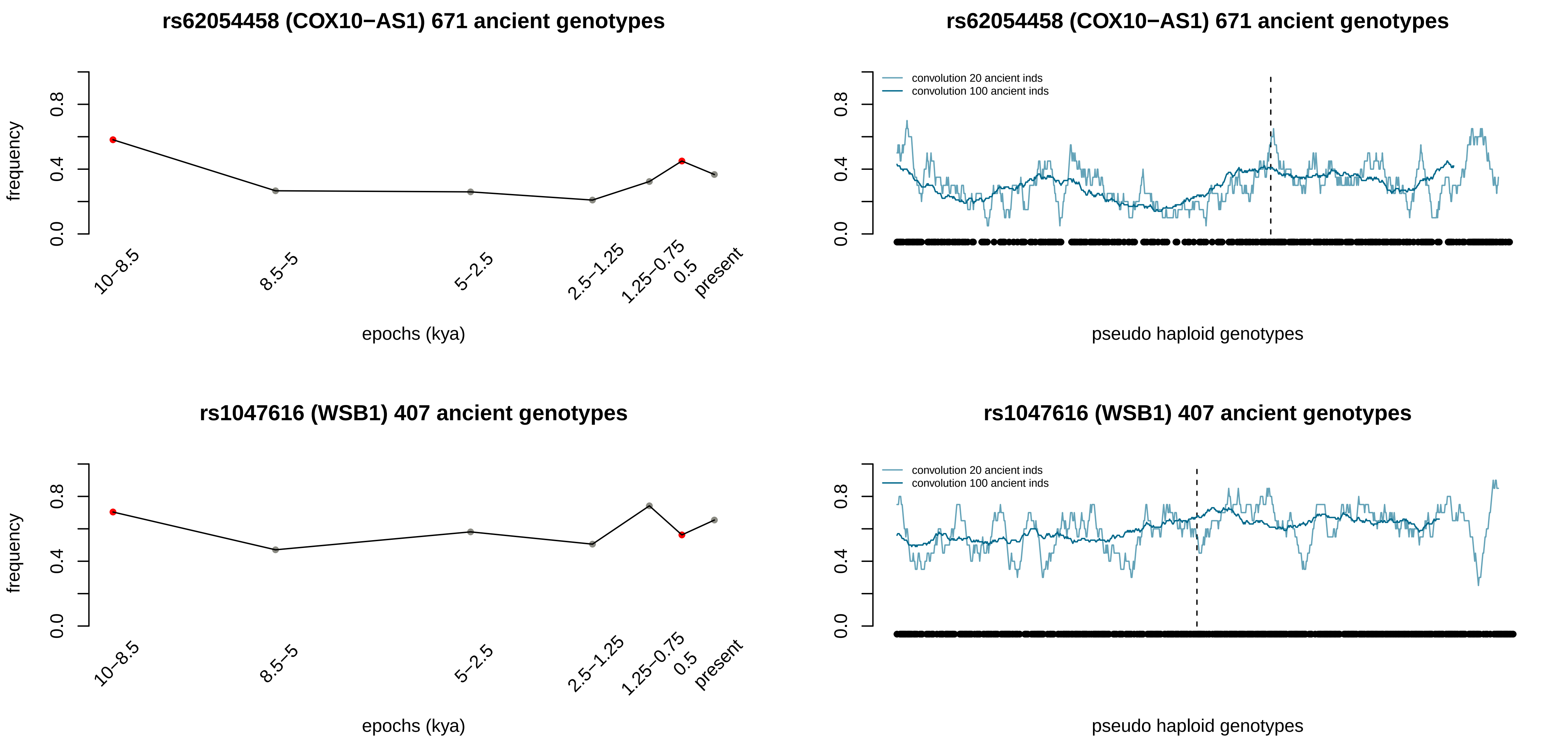
rs62054458 (COX10−AS1) 671 ancient genotypes
frequency
0.0
0.4
0.8
10−8.5
8.5−5
5−2.5
2.5−1.25
1.25−0.75
0.5
present
epochs (kya)
rs62054458 (COX10−AS1) 671 ancient genotypes
convolution 20 ancient inds
convolution 100 ancient inds
pseudo haploid genotypes
rs1047616 (WSB1) 407 ancient genotypes
frequency
epochs (kya)
rs1047616 (WSB1) 407 ancient genotypes
convolution 20 ancient inds
convolution 100 ancient inds
pseudo haploid genotypes

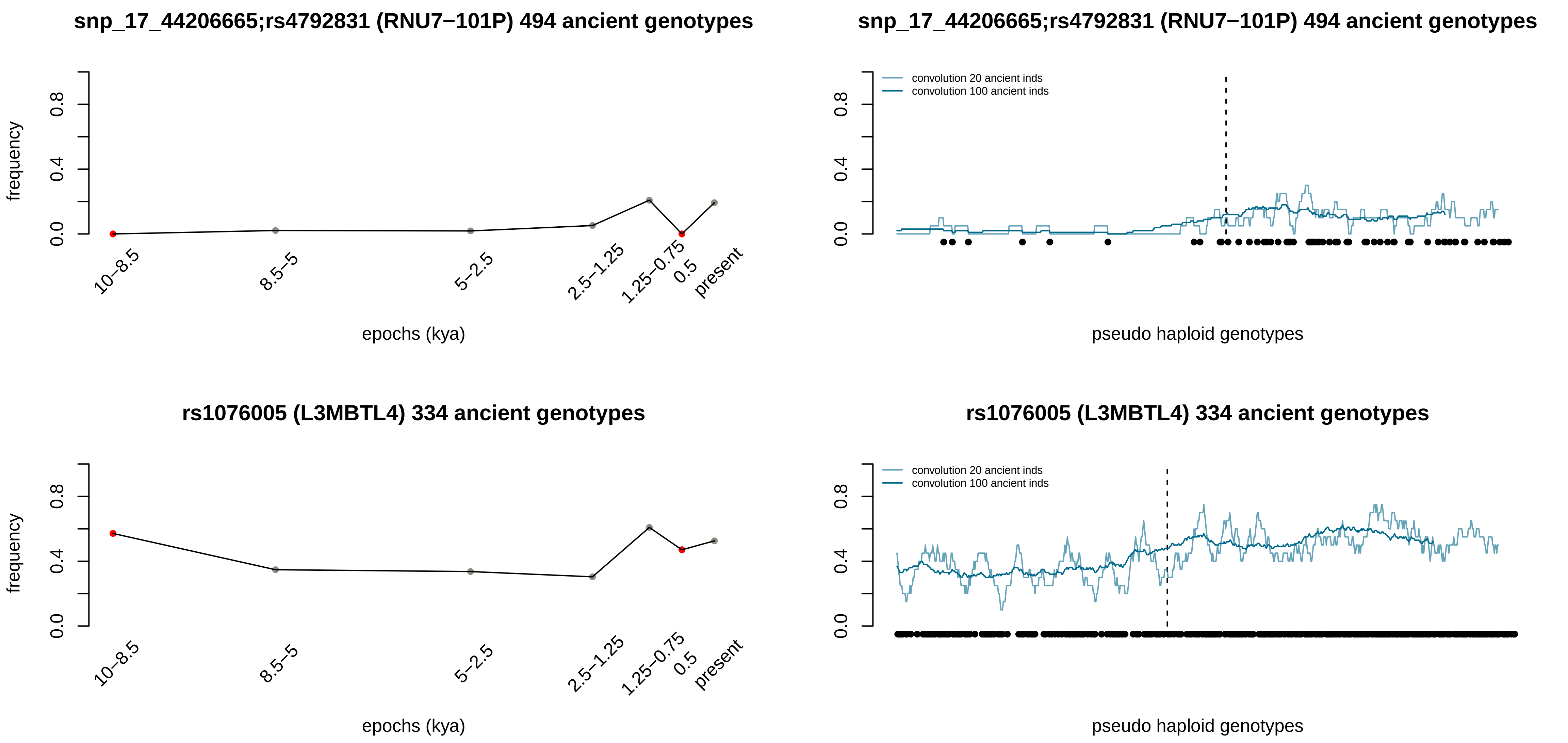

snp_17_44206665;rs4792831 (RNU7−101P) 494 ancient genotypes
frequency
epochs (kya)
10–8.5
8.5–5
5–2.5
2.5–1.25
1.25–0.75
0.5
present
snp_17_44206665;rs4792831 (RNU7−101P) 494 ancient genotypes
convolution 20 ancient inds
convolution 100 ancient inds
pseudo haploid genotypes
rs1076005 (L3MBTL4) 334 ancient genotypes
frequency
epochs (kya)
rs1076005 (L3MBTL4) 334 ancient genotypes
convolution 20 ancient inds
convolution 100 ancient inds
pseudo haploid genotypes

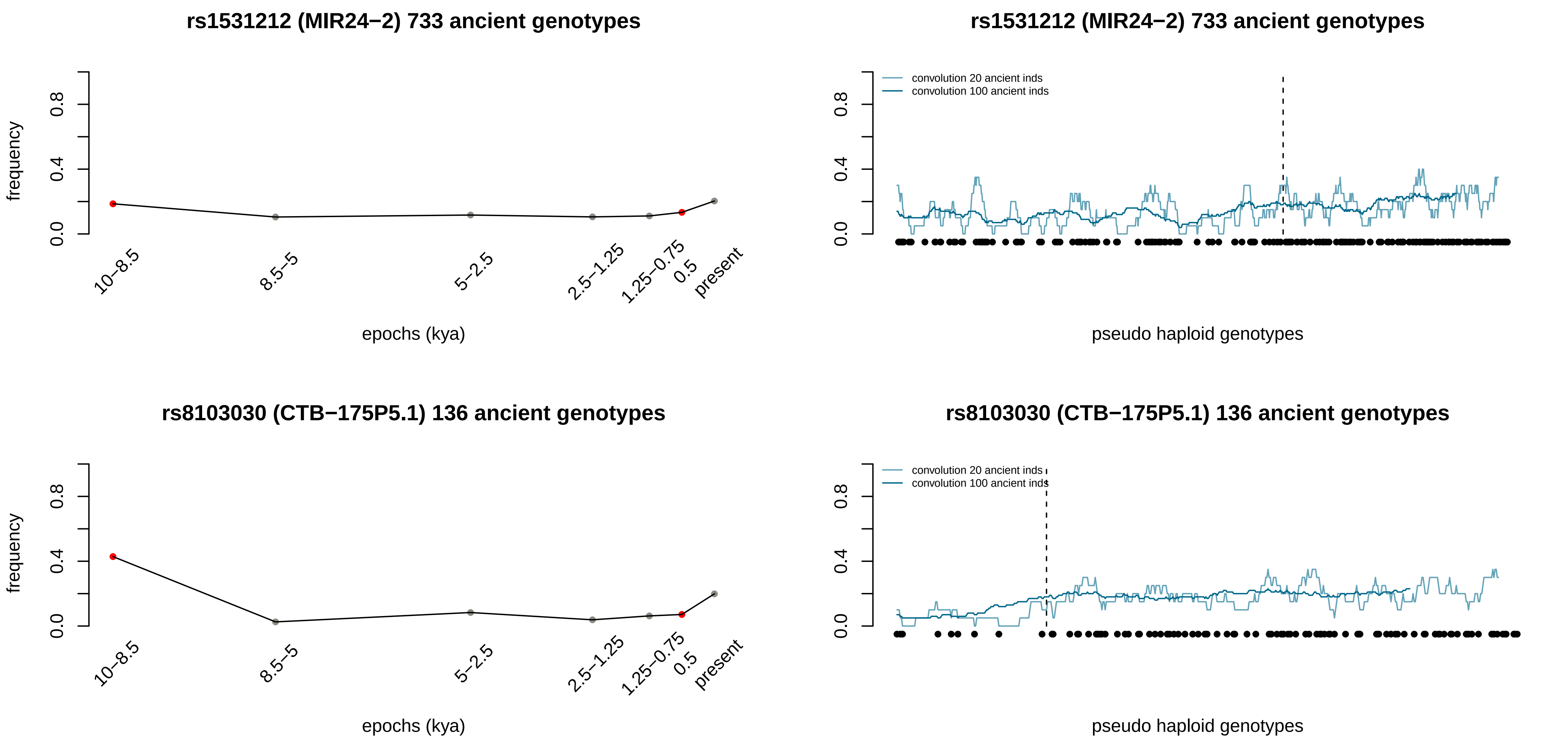
rs1531212 (MIR24−2) 733 ancient genotypes
frequency
0.0
0.4
0.8
10−8.5
8.5−5
5−2.5
2.5−1.25
1.25−0.75
0.5
present
epochs (kya)
rs1531212 (MIR24−2) 733 ancient genotypes
convolution 20 ancient inds
convolution 100 ancient inds
pseudo haploid genotypes
rs8103030 (CTB−175P5.1) 136 ancient genotypes
frequency
epochs (kya)
rs8103030 (CTB−175P5.1) 136 ancient genotypes
convolution 20 ancient inds
convolution 100 ancient inds
pseudo haploid genotypes

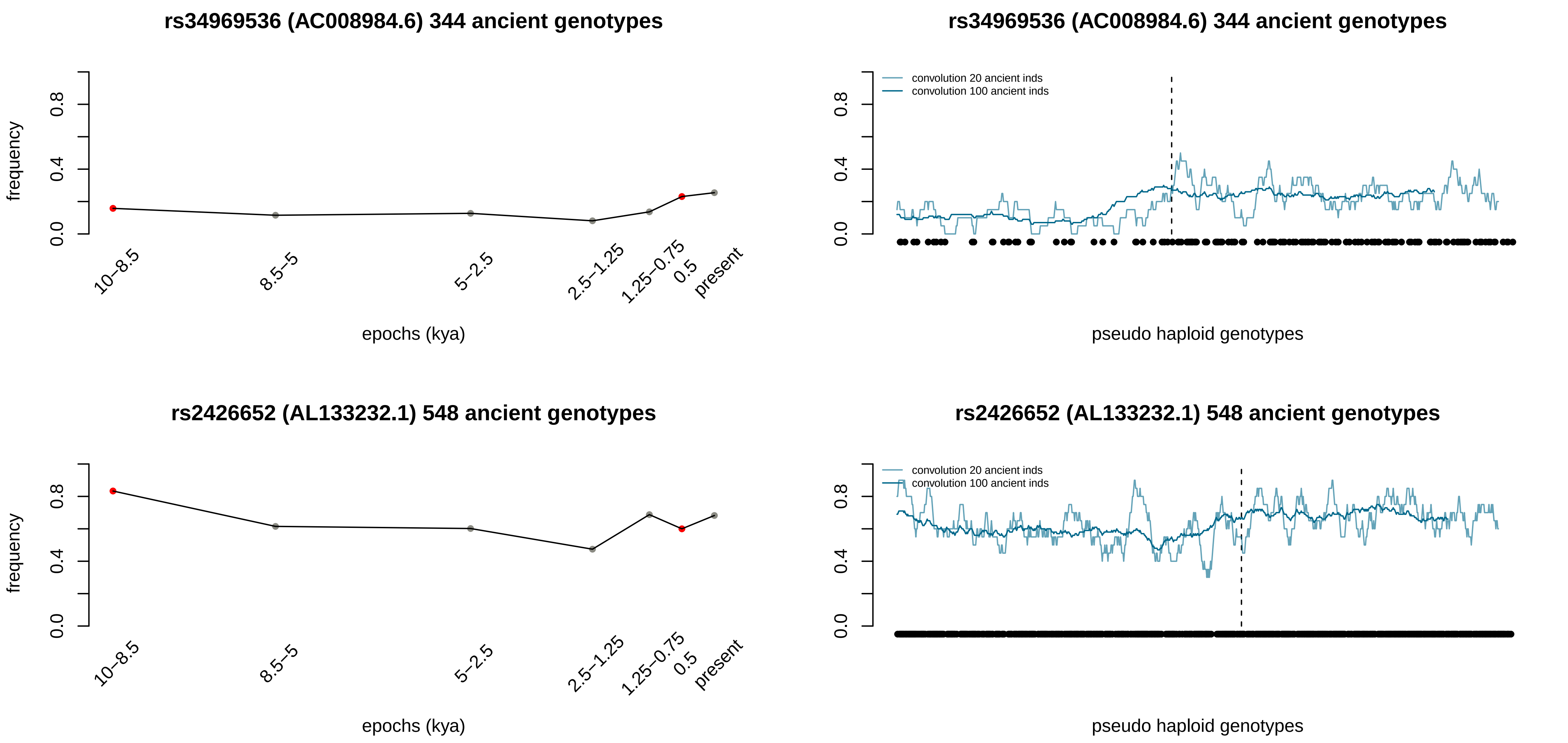
rs34969536 (AC008984.6) 344 ancient genotypes
frequency
epochs (kya)
10–8.5
8.5–5
5–2.5
2.5–1.25
1.25–0.75
0.5
present
rs34969536 (AC008984.6) 344 ancient genotypes
convolution 20 ancient inds
convolution 100 ancient inds
pseudo haploid genotypes
rs2426652 (AL133232.1) 548 ancient genotypes
frequency
epochs (kya)
rs2426652 (AL133232.1) 548 ancient genotypes
convolution 20 ancient inds
convolution 100 ancient inds
pseudo haploid genotypes

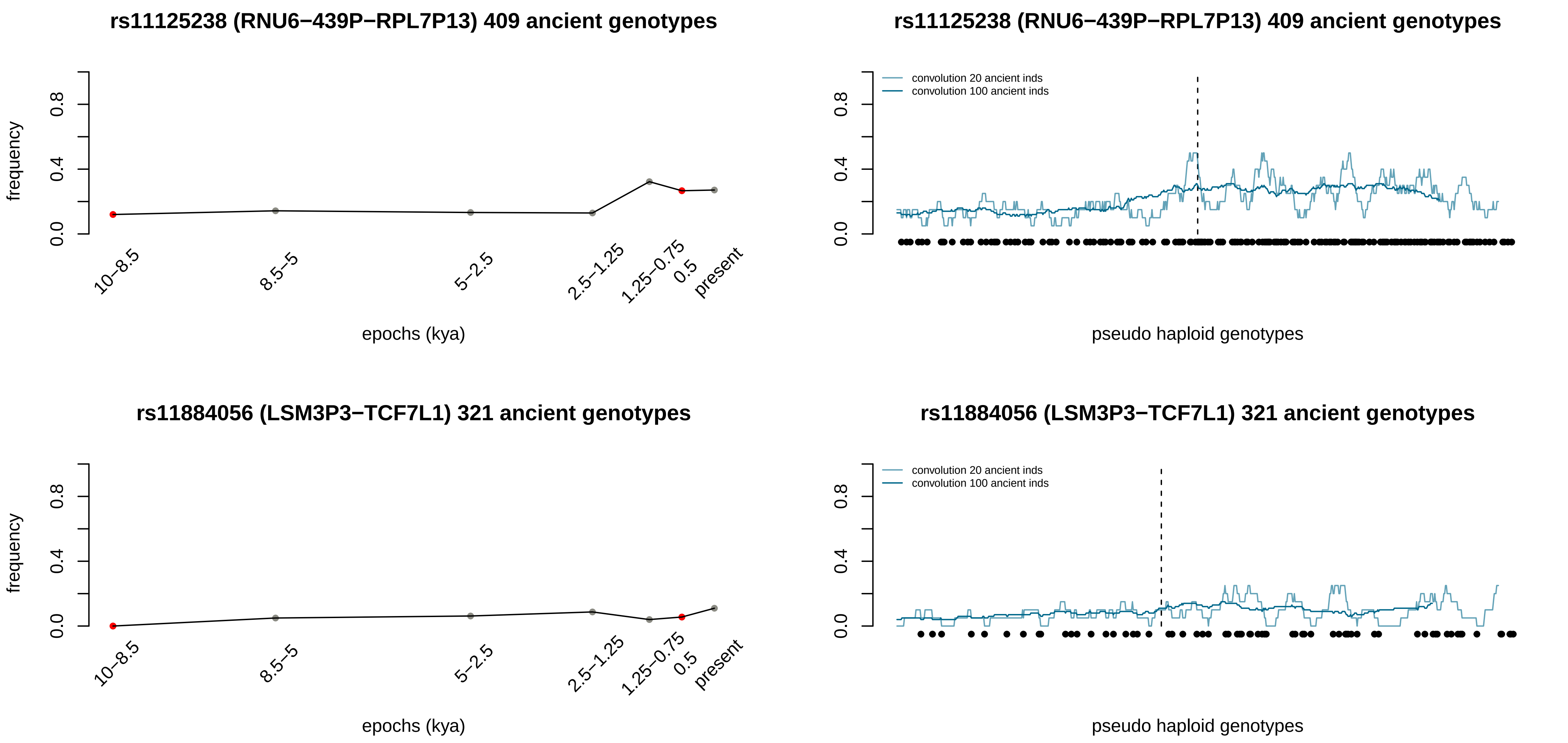

rs11125238 (RNU6−439P−RPL7P13) 409 ancient genotypes
frequency
0.0
0.4
0.8
10−8.5
8.5−5
5−2.5
2.5−1.25
1.25−0.75
0.5
present
epochs (kya)
rs11125238 (RNU6−439P−RPL7P13) 409 ancient genotypes
convolution 20 ancient inds
convolution 100 ancient inds
pseudo haploid genotypes
rs11884056 (LSM3P3−TCF7L1) 321 ancient genotypes
frequency
epochs (kya)
rs11884056 (LSM3P3−TCF7L1) 321 ancient genotypes
convolution 20 ancient inds
convolution 100 ancient inds
pseudo haploid genotypes

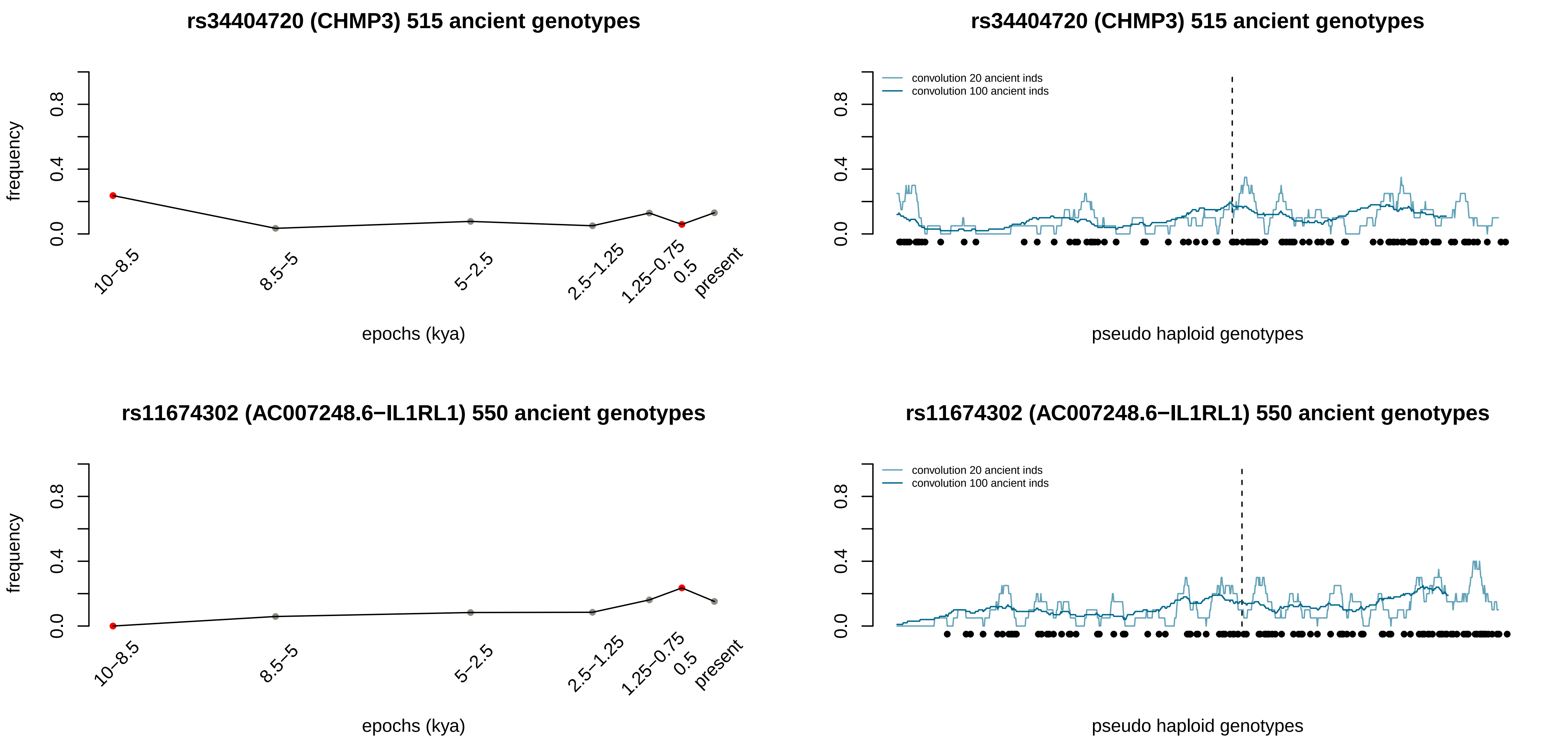

rs34404720 (CHMP3) 515 ancient genotypes
frequency
0.0
0.4
0.8
10–8.5
8.5–5
5–2.5
2.5–1.25
1.25–0.75
0.5
present
epochs (kya)
rs34404720 (CHMP3) 515 ancient genotypes
convolution 20 ancient inds
convolution 100 ancient inds
pseudo haploid genotypes
rs11674302 (AC007248.6−IL1RL1) 550 ancient genotypes
frequency
epochs (kya)
rs11674302 (AC007248.6−IL1RL1) 550 ancient genotypes
convolution 20 ancient inds
convolution 100 ancient inds
pseudo haploid genotypes

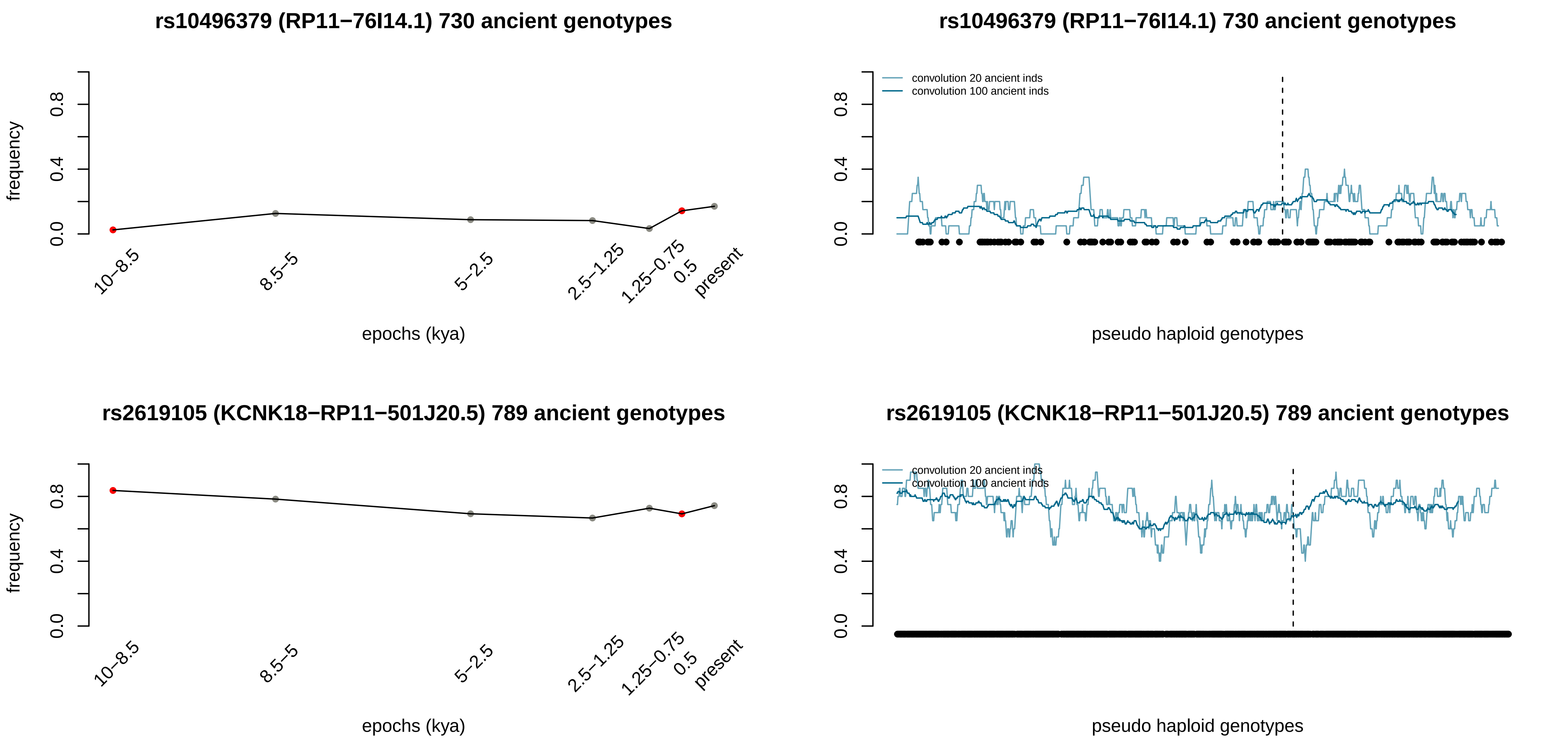

rs10496379 (RP11−76I14.1) 730 ancient genotypes
frequency
0.0
0.4
0.8
10−8.5
8.5−5
5−2.5
2.5−1.25
1.25−0.75
0.5
present
epochs (kya)
rs10496379 (RP11−76I14.1) 730 ancient genotypes
convolution 20 ancient inds
convolution 100 ancient inds
pseudo haploid genotypes
rs2619105 (KCNK18−RP11−501J20.5) 789 ancient genotypes
frequency
epochs (kya)
rs2619105 (KCNK18−RP11−501J20.5) 789 ancient genotypes
convolution 20 ancient inds
convolution 100 ancient inds
pseudo haploid genotypes

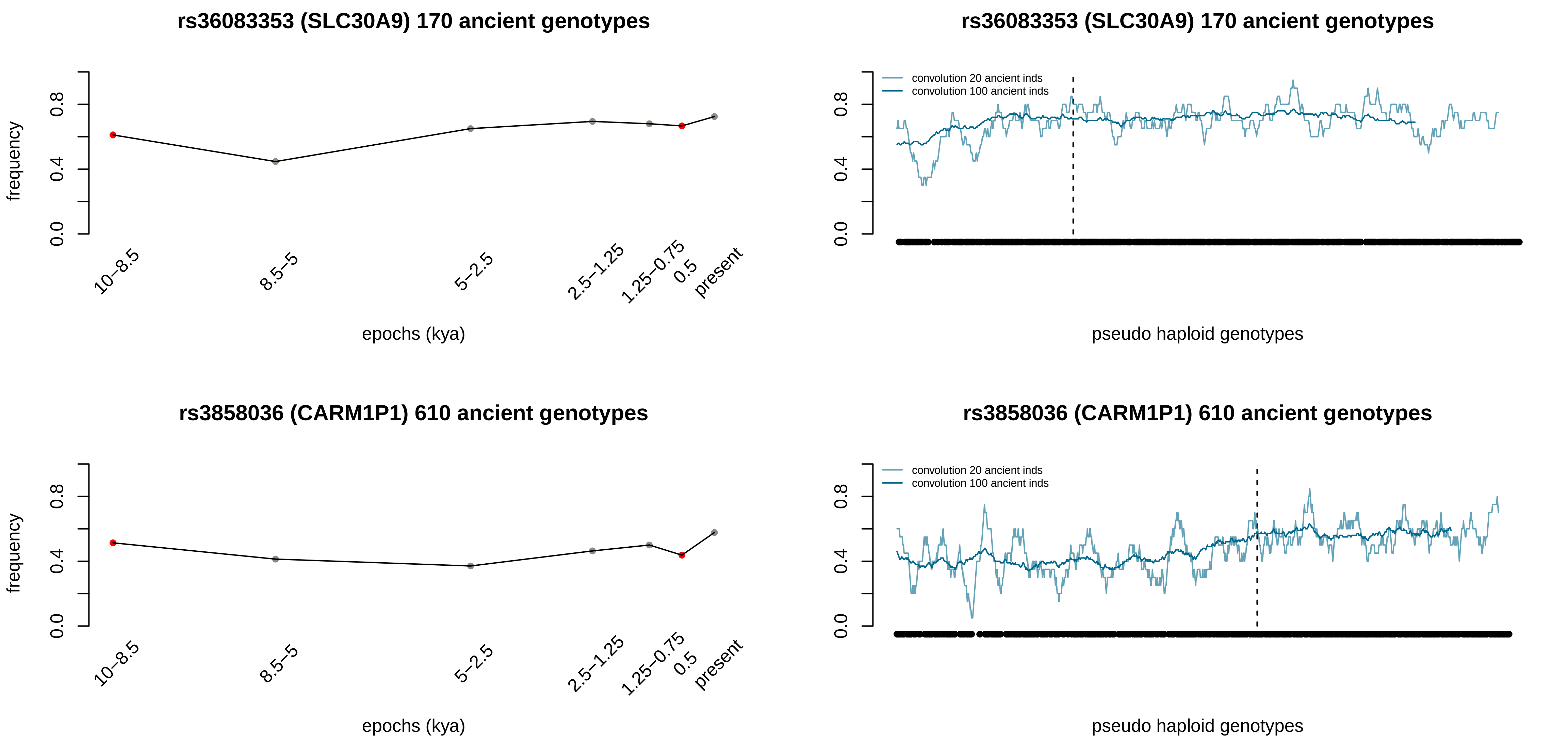
rs36083353 (SLC30A9) 170 ancient genotypes
frequency
0.0
0.4
0.8
10–8.5
8.5–5
5–2.5
2.5–1.25
1.25–0.75
0.5
present
epochs (kya)
rs36083353 (SLC30A9) 170 ancient genotypes
convolution 20 ancient inds
convolution 100 ancient inds
0.0
0.4
0.8
pseudo haploid genotypes
rs3858036 (CARM1P1) 610 ancient genotypes
frequency
0.0
0.4
0.8
10–8.5
8.5–5
5–2.5
2.5–1.25
1.25–0.75
0.5
present
epochs (kya)
rs3858036 (CARM1P1) 610 ancient genotypes
convolution 20 ancient inds
convolution 100 ancient inds
0.0
0.4
0.8
pseudo haploid genotypes

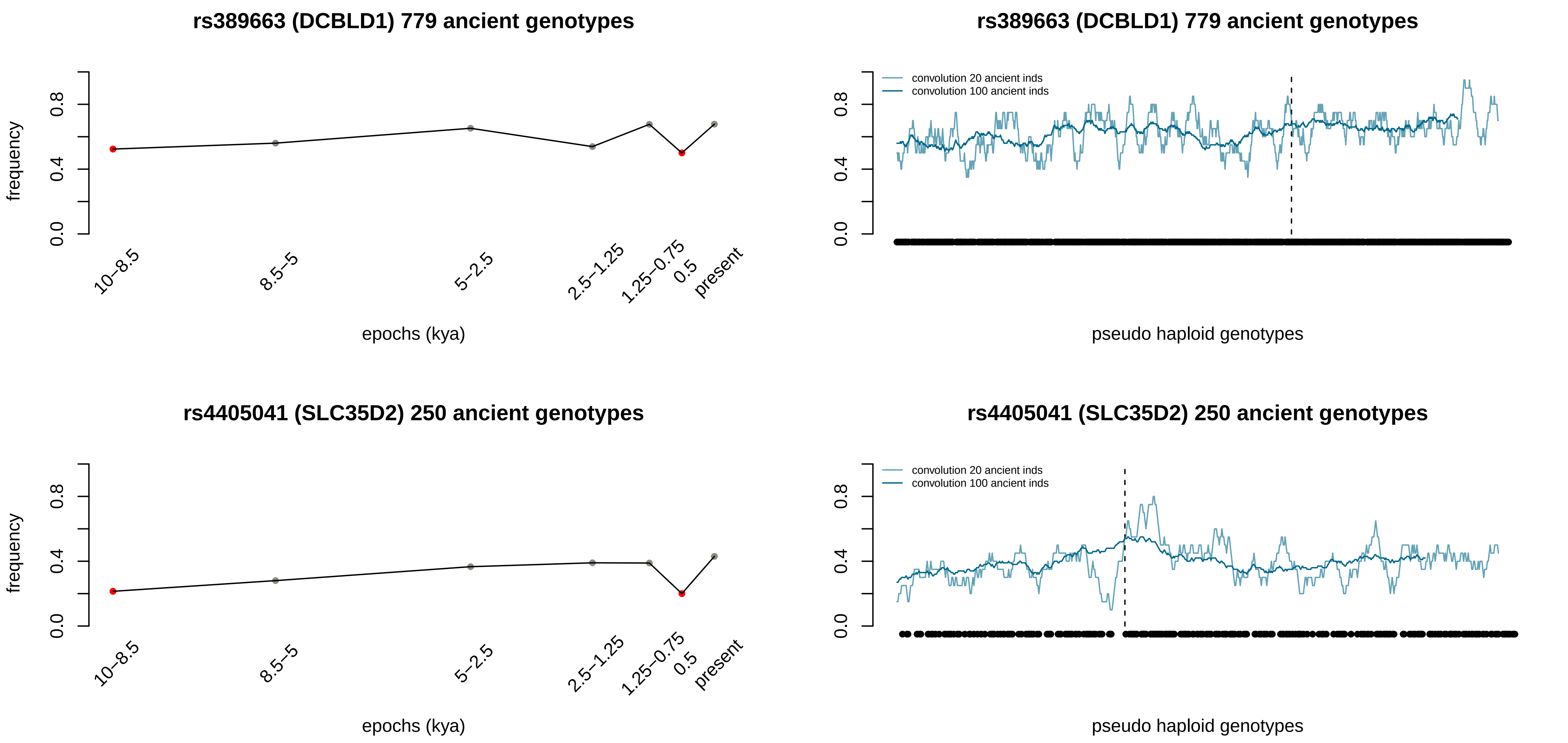
rs389663 (DCBLD1) 779 ancient genotypes
frequency
0.0
0.4
0.8
10–8.5
8.5–5
5–2.5
2.5–1.25
1.25–0.75
0.5
present
epochs (kya)
rs389663 (DCBLD1) 779 ancient genotypes
convolution 20 ancient inds
convolution 100 ancient inds
pseudo haploid genotypes
rs4405041 (SLC35D2) 250 ancient genotypes
frequency
epochs (kya)
rs4405041 (SLC35D2) 250 ancient genotypes
convolution 20 ancient inds
convolution 100 ancient inds
pseudo haploid genotypes

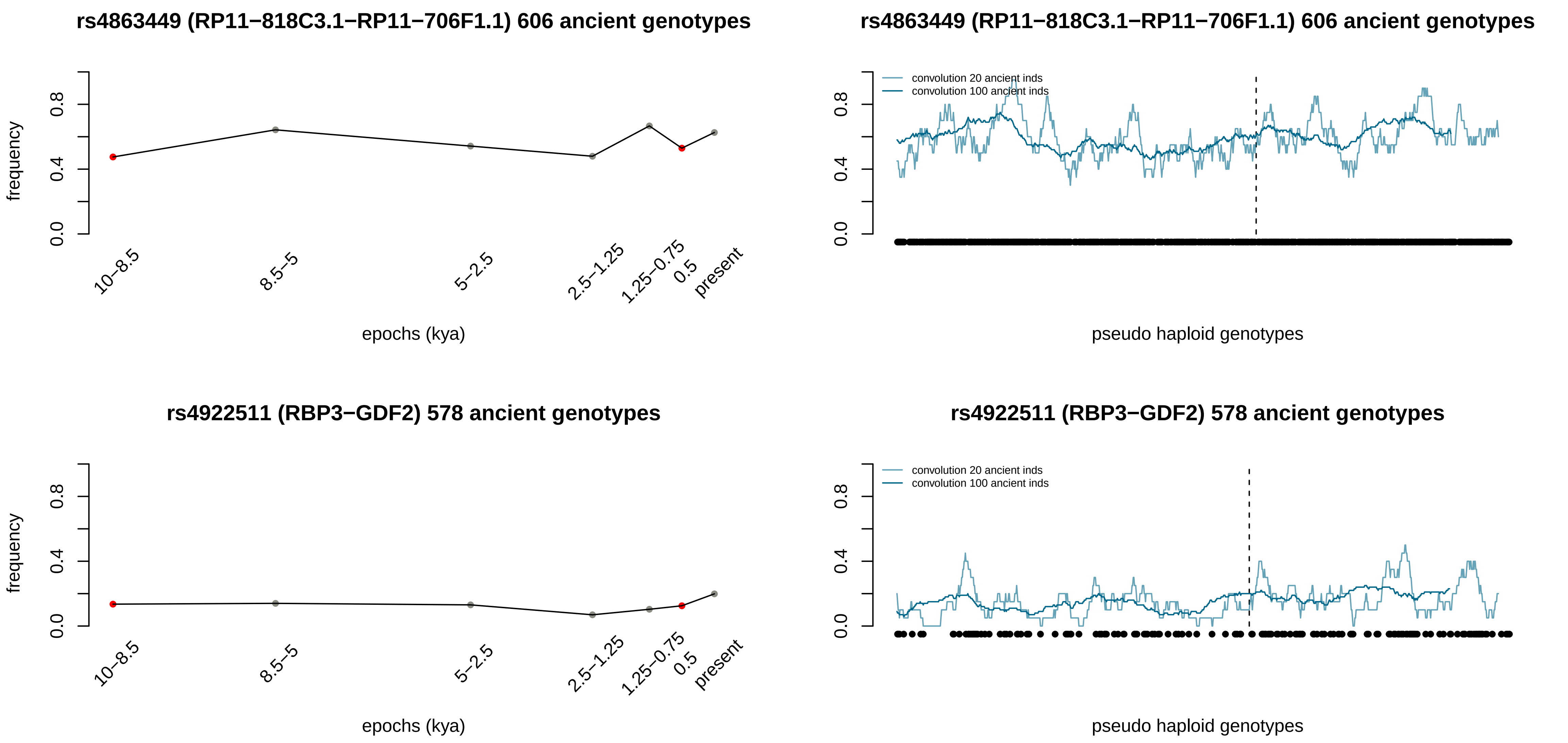
rs4863449 (RP11−818C3.1−RP11−706F1.1) 606 ancient genotypes
frequency
0.0
0.4
0.8
10−8.5
8.5−5
5−2.5
2.5−1.25
1.25−0.75
0.5
present
epochs (kya)
rs4863449 (RP11−818C3.1−RP11−706F1.1) 606 ancient genotypes
convolution 20 ancient inds
convolution 100 ancient inds
pseudo haploid genotypes
rs4922511 (RBP3−GDF2) 578 ancient genotypes
frequency
epochs (kya)
rs4922511 (RBP3−GDF2) 578 ancient genotypes
convolution 20 ancient inds
convolution 100 ancient inds
pseudo haploid genotypes

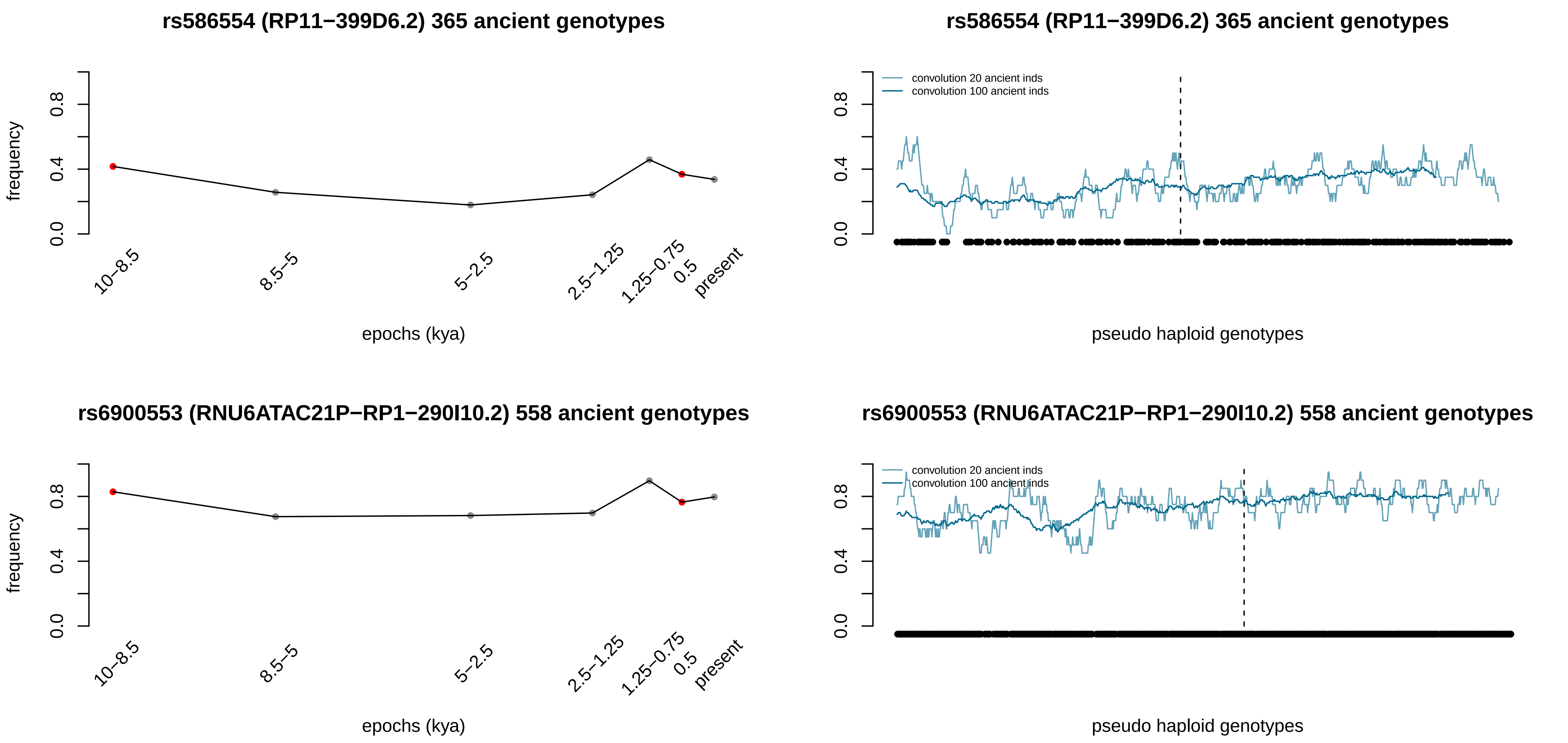

rs586554 (RP11−399D6.2) 365 ancient genotypes
frequency
0.0
0.4
0.8
10−8.5
8.5−5
5−2.5
2.5−1.25
1.25−0.75
0.5
present
epochs (kya)
rs586554 (RP11−399D6.2) 365 ancient genotypes
convolution 20 ancient inds
convolution 100 ancient inds
pseudo haploid genotypes
rs6900553 (RNU6ATAC21P−RP1−290I10.2) 558 ancient genotypes
epochs (kya)
rs6900553 (RNU6ATAC21P−RP1−290I10.2) 558 ancient genotypes
pseudo haploid genotypes

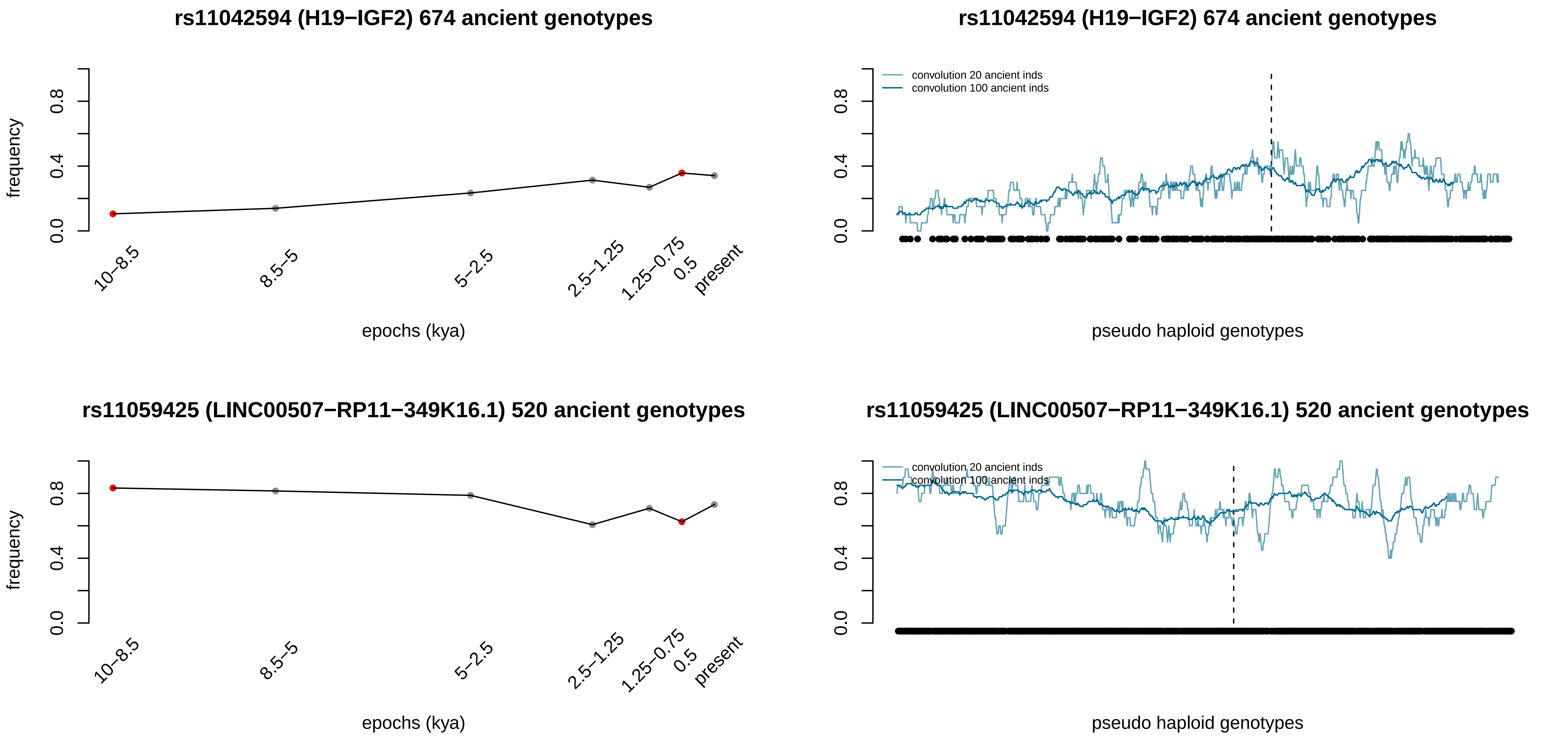

rs11042594 (H19−IGF2) 674 ancient genotypes
frequency
0.0
0.4
0.8
10−8.5
8.5−5
5−2.5
2.5−1.25
1.25−0.75
0.5
present
epochs (kya)
rs11042594 (H19−IGF2) 674 ancient genotypes
convolution 20 ancient inds
convolution 100 ancient inds
pseudo haploid genotypes
rs11059425 (LINC00507−RP11−349K16.1) 520 ancient genotypes
frequency
epochs (kya)
rs11059425 (LINC00507−RP11−349K16.1) 520 ancient genotypes
convolution 20 ancient inds
convolution 100 ancient inds
pseudo haploid genotypes

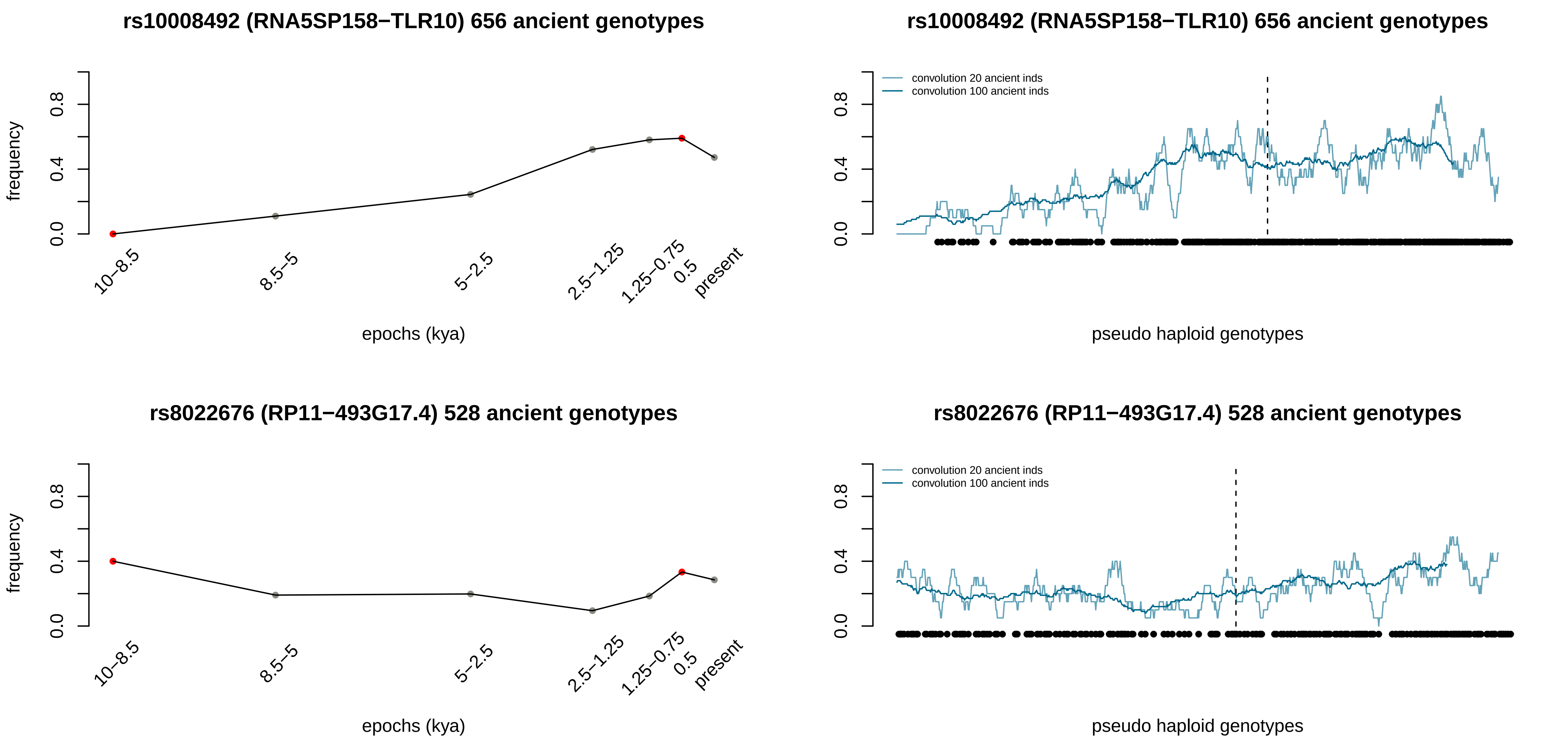
rs10008492 (RNA5SP158−TLR10) 656 ancient genotypes
frequency
0.0
0.4
0.8
10−8.5
8.5−5
5−2.5
2.5−1.25
1.25−0.75
0.5
present
epochs (kya)
rs10008492 (RNA5SP158−TLR10) 656 ancient genotypes
convolution 20 ancient inds
convolution 100 ancient inds
0.0
0.4
0.8
pseudo haploid genotypes
rs8022676 (RP11−493G17.4) 528 ancient genotypes
frequency
0.0
0.4
0.8
10−8.5
8.5−5
5−2.5
2.5−1.25
1.25−0.75
0.5
present
epochs (kya)
rs8022676 (RP11−493G17.4) 528 ancient genotypes
convolution 20 ancient inds
convolution 100 ancient inds
0.0
0.4
0.8
pseudo haploid genotypes

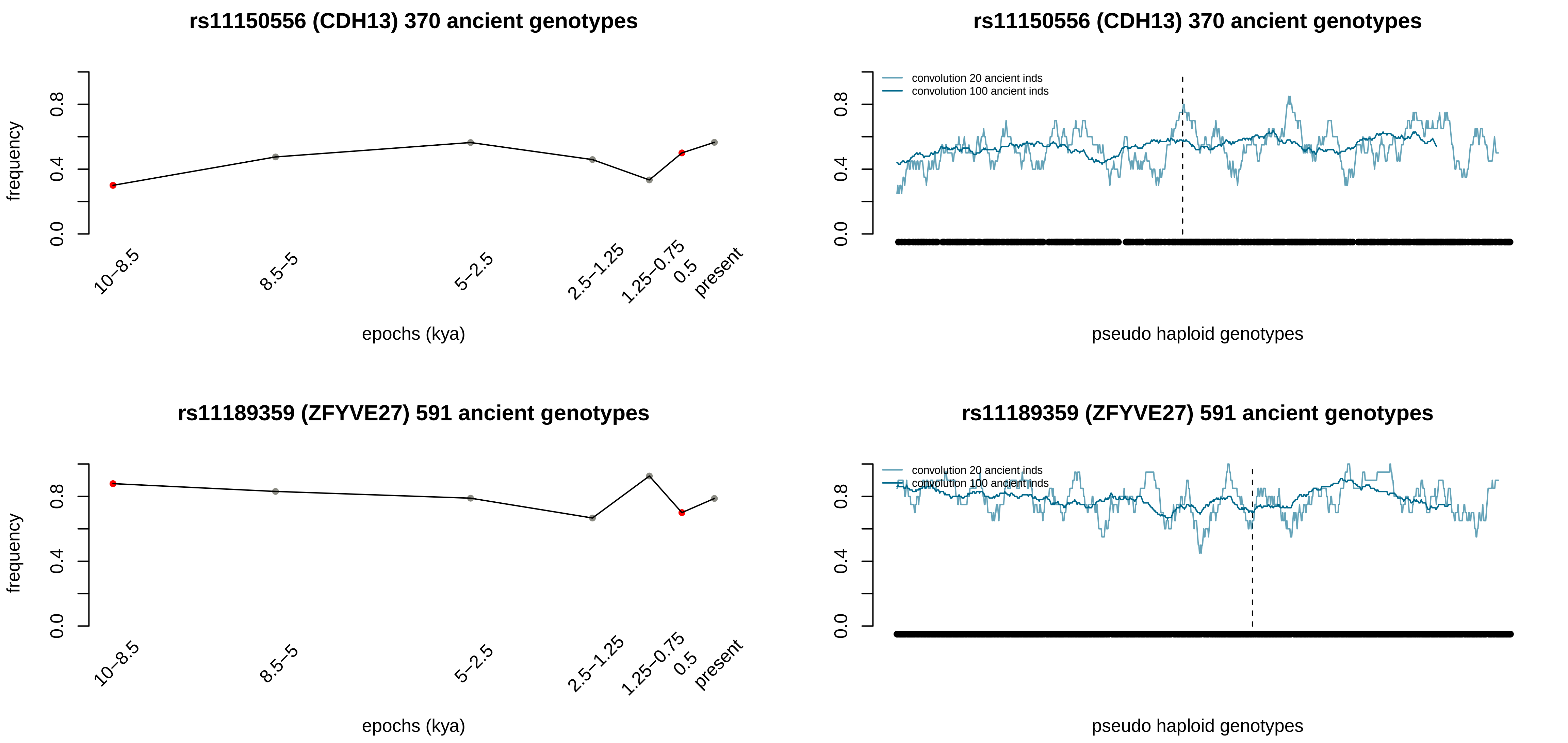
rs11150556 (CDH13) 370 ancient genotypes
frequency
0.0
0.4
0.8
10–8.5
8.5–5
5–2.5
2.5–1.25
1.25–0.75
0.5
present
epochs (kya)
rs11150556 (CDH13) 370 ancient genotypes
convolution 20 ancient inds
convolution 100 ancient inds
pseudo haploid genotypes
rs11189359 (ZFYVE27) 591 ancient genotypes
frequency
epochs (kya)
rs11189359 (ZFYVE27) 591 ancient genotypes
convolution 20 ancient inds
convolution 100 ancient inds
pseudo haploid genotypes

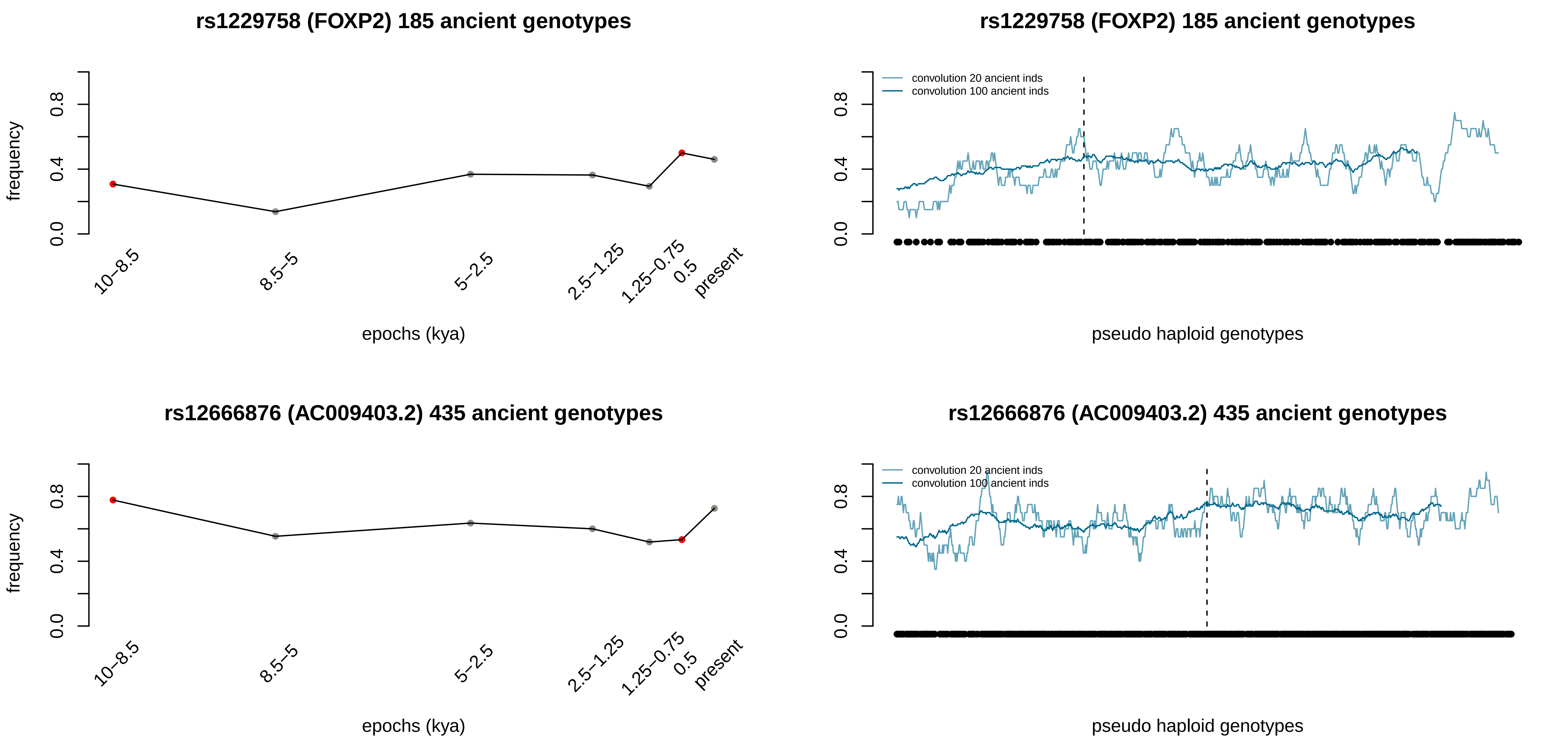

rs1229758 (FOXP2) 185 ancient genotypes
frequency
0.0
0.4
0.8
10–8.5
8.5–5
5–2.5
2.5–1.25
1.25–0.75
0.5
present
epochs (kya)
rs1229758 (FOXP2) 185 ancient genotypes
convolution 20 ancient inds
convolution 100 ancient inds
0.0
0.4
0.8
pseudo haploid genotypes
rs12666876 (AC009403.2) 435 ancient genotypes
frequency
0.0
0.4
0.8
10–8.5
8.5–5
5–2.5
2.5–1.25
1.25–0.75
0.5
present
epochs (kya)
rs12666876 (AC009403.2) 435 ancient genotypes
convolution 20 ancient inds
convolution 100 ancient inds
0.0
0.4
0.8
pseudo haploid genotypes

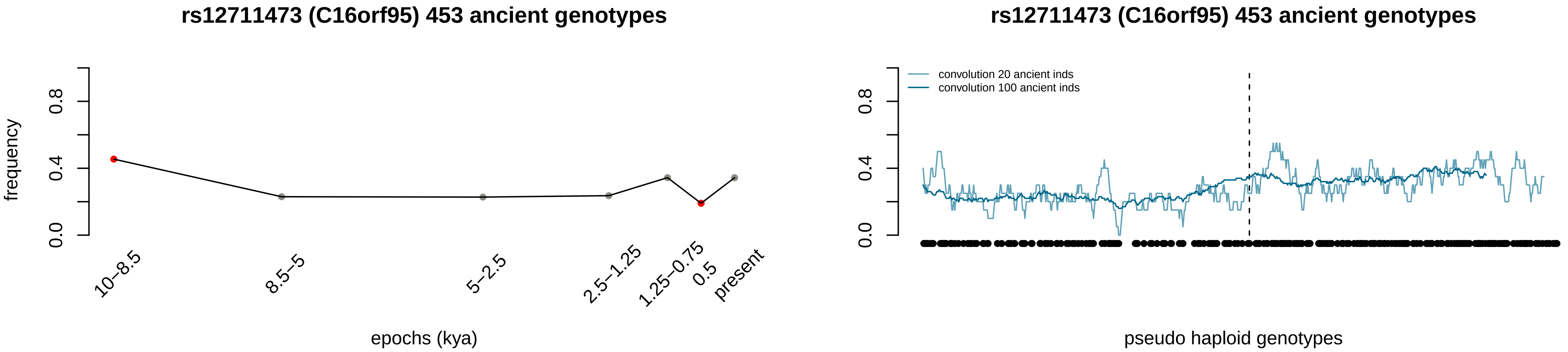

rs12711473 (C16orf95) 453 ancient genotypes
frequency
0.0
0.4
0.8
10–8.5
8.5–5
5–2.5
2.5–1.25
1.25–0.75
0.5
present
epochs (kya)
rs12711473 (C16orf95) 453 ancient genotypes
convolution 20 ancient inds
convolution 100 ancient inds
0.0
0.4
0.8
pseudo haploid genotypes
