## Supplementary figures and images for "Deep estimation of the intensity and timing of selection from ancient genomes"

### 1D images for the positively selected variants associated to Black Death pandemic (ERAP2)

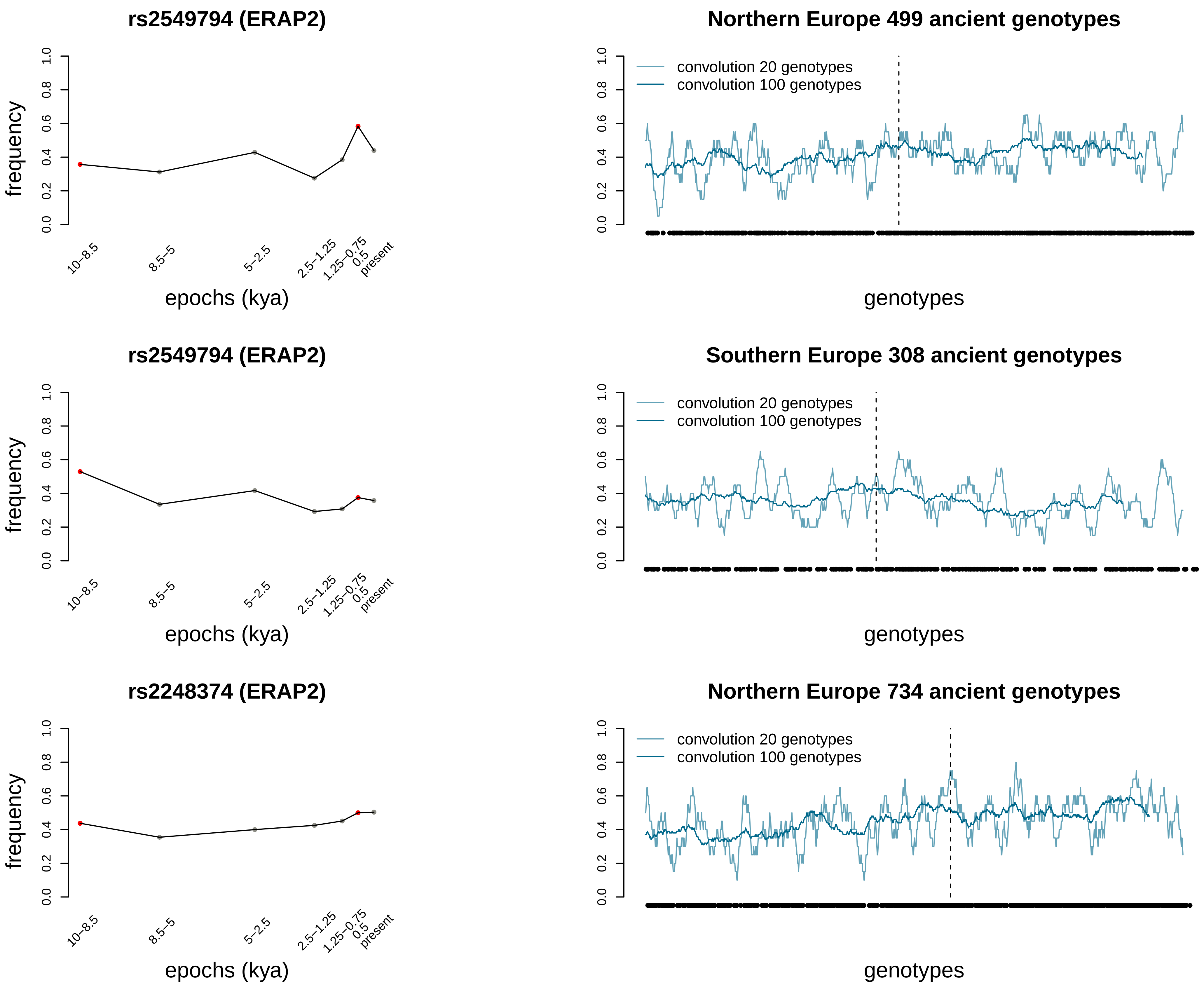

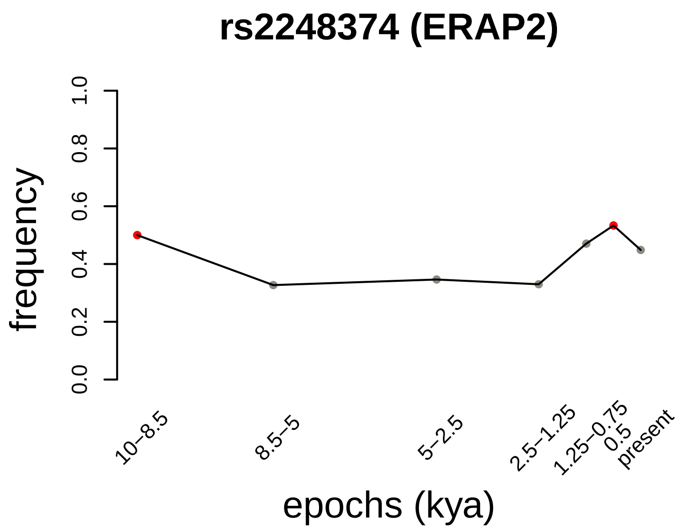

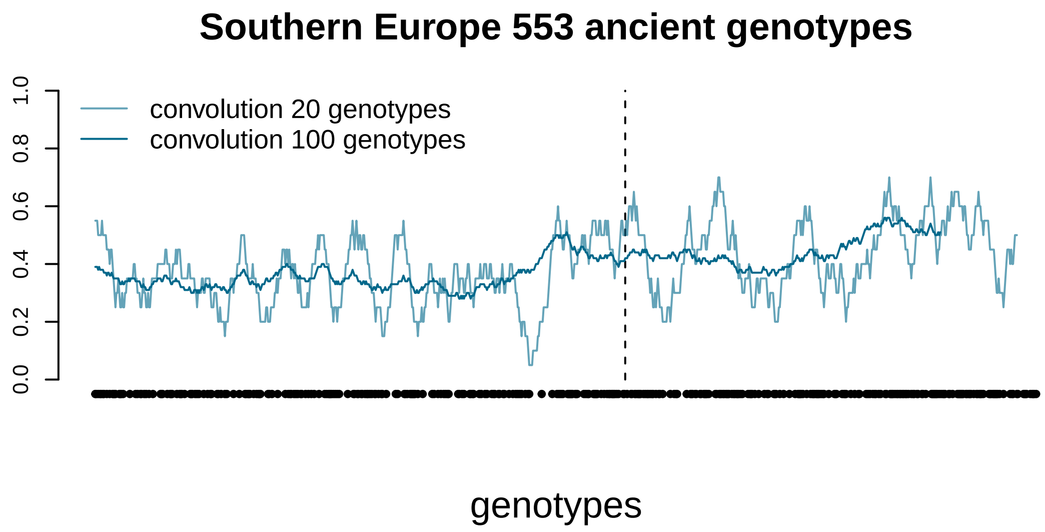
