## Supplementary material for "Deep estimation of the intensity and timing of selection from ancient genomes": 1D images for the 50 negatively selected variants

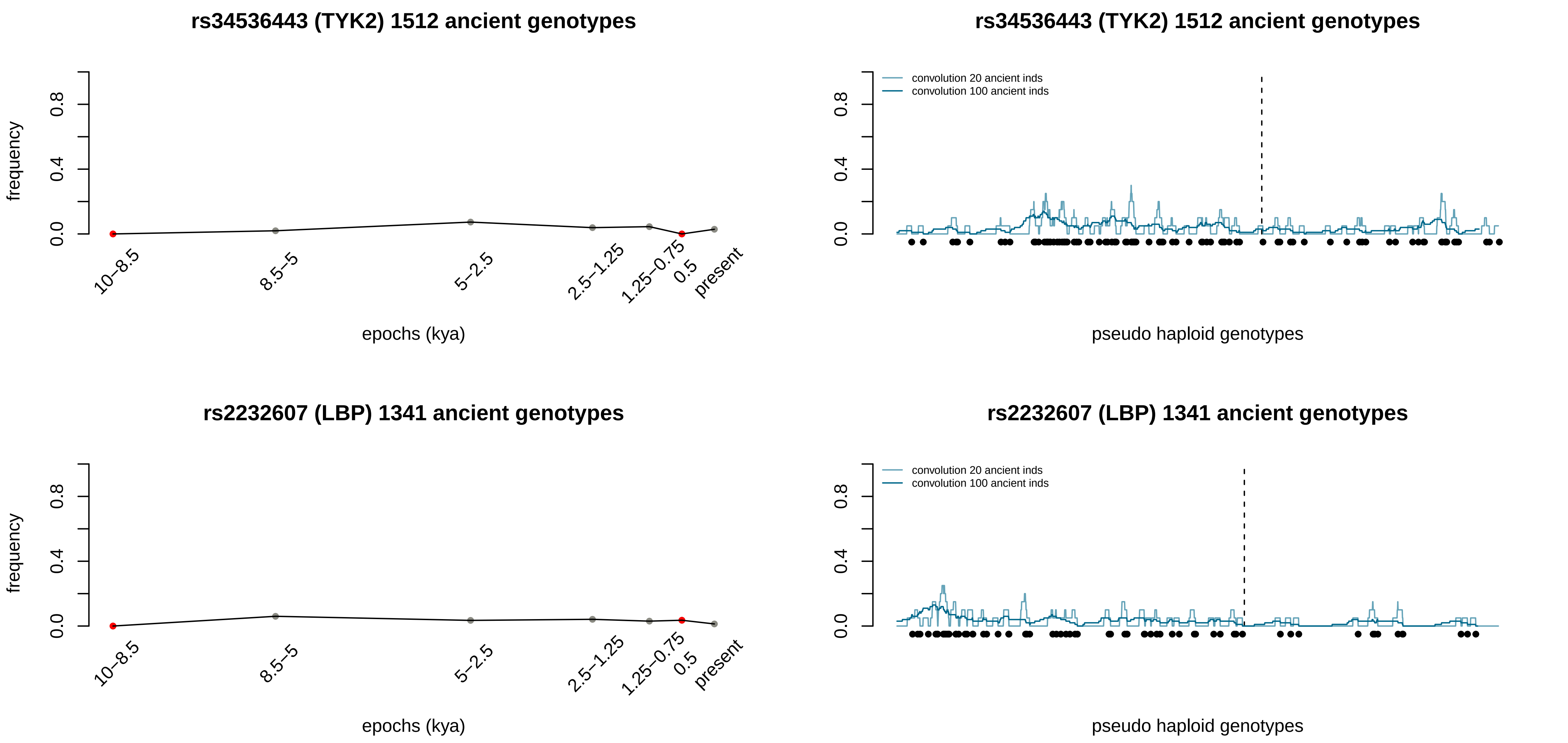

rs34536443 (TYK2) 1512 ancient genotypes
frequency
0.0
0.4
0.8
10–8.5
8.5–5
5–2.5
2.5–1.25
1.25–0.75
0.5
present
epochs (kya)
rs34536443 (TYK2) 1512 ancient genotypes
convolution 20 ancient inds
convolution 100 ancient inds
pseudo haploid genotypes
rs2232607 (LBP) 1341 ancient genotypes
frequency
epochs (kya)
rs2232607 (LBP) 1341 ancient genotypes
convolution 20 ancient inds
convolution 100 ancient inds
pseudo haploid genotypes

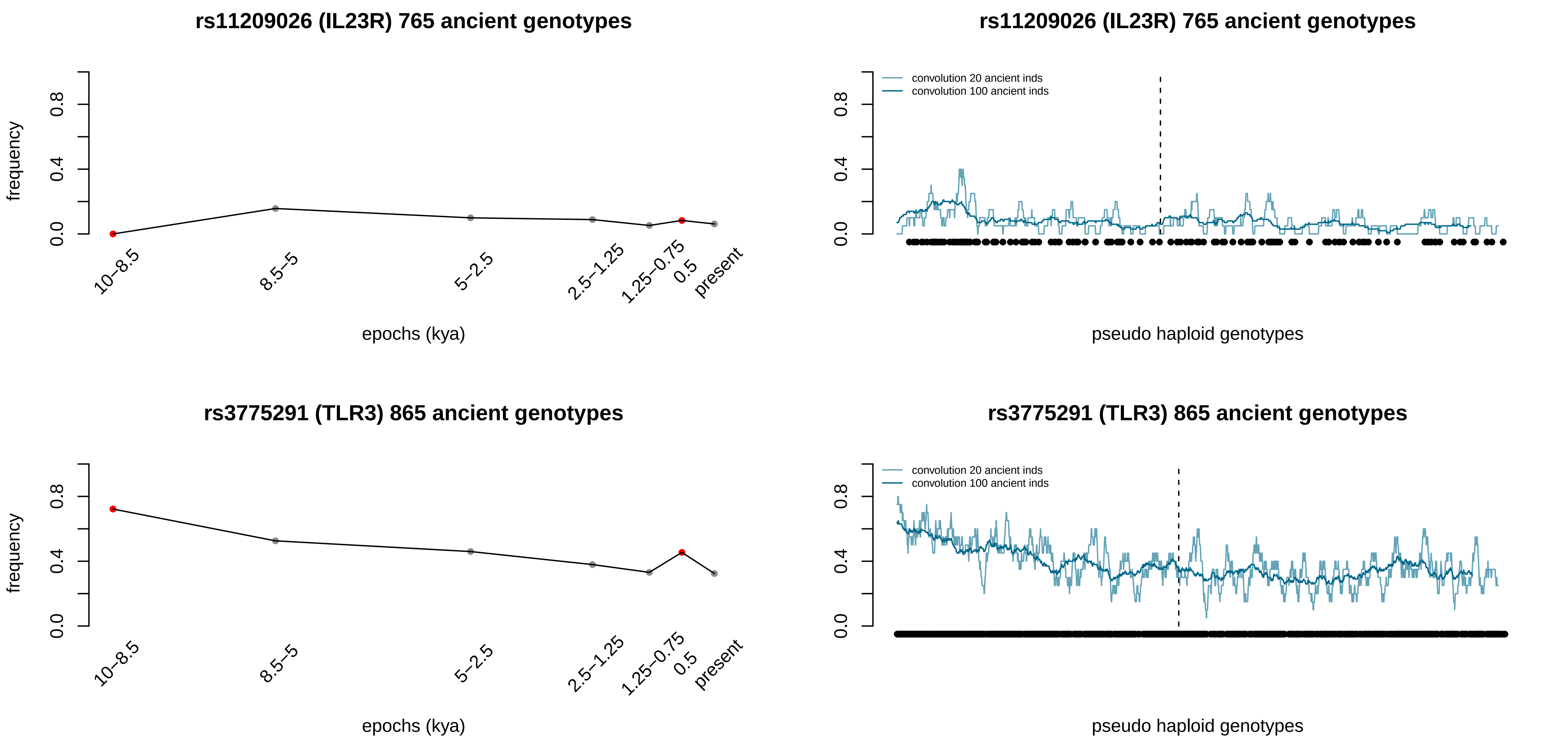

rs11209026 (IL23R) 765 ancient genotypes
frequency
0.0
0.4
0.8
10–8.5
8.5–5
5–2.5
2.5–1.25
1.25–0.75
0.5
present
epochs (kya)
rs11209026 (IL23R) 765 ancient genotypes
convolution 20 ancient inds
convolution 100 ancient inds
pseudo haploid genotypes
rs3775291 (TLR3) 865 ancient genotypes
frequency
epochs (kya)
rs3775291 (TLR3) 865 ancient genotypes
convolution 20 ancient inds
convolution 100 ancient inds
pseudo haploid genotypes

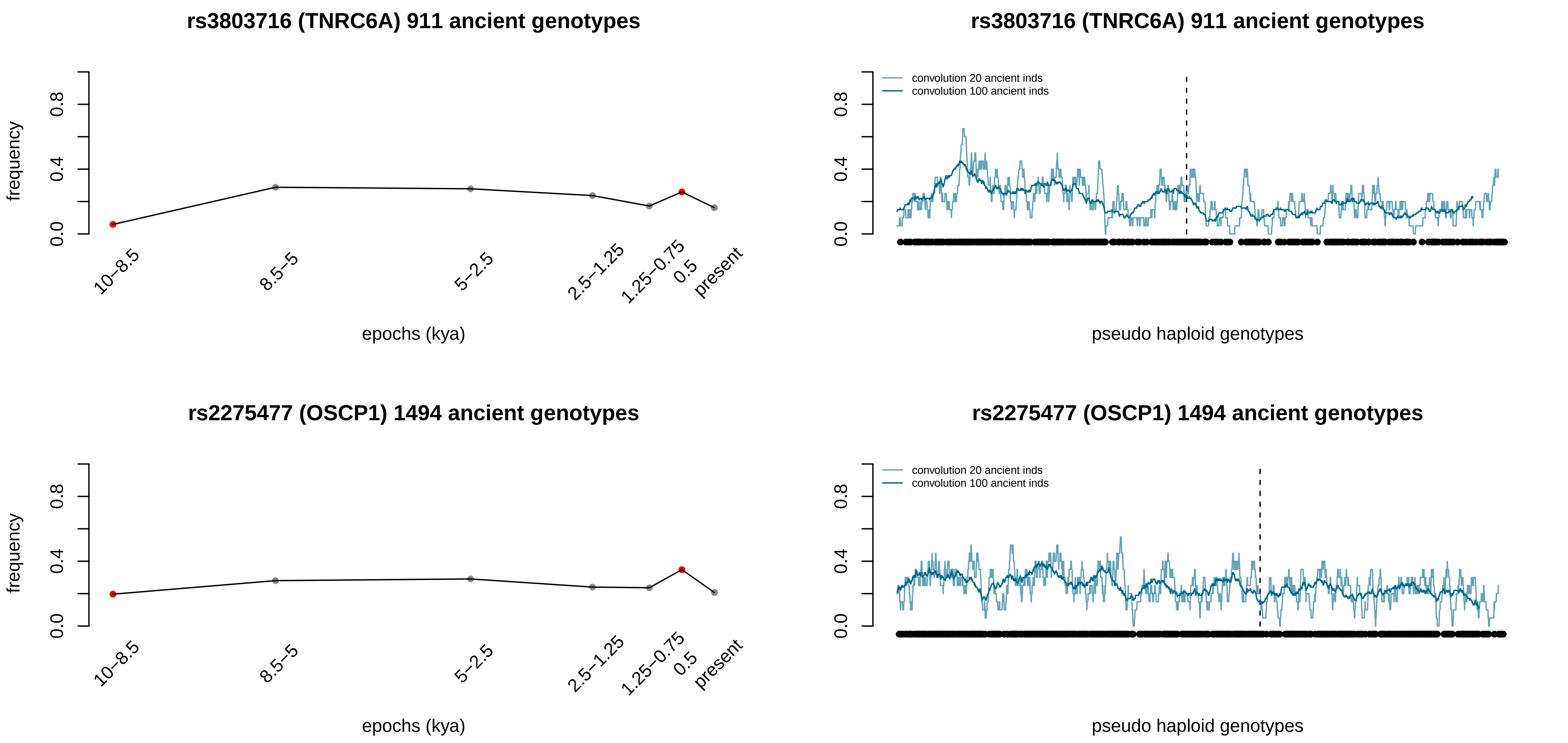
rs3803716 (TNRC6A) 911 ancient genotypes
frequency
0.0
0.4
0.8
10–8.5
8.5–5
5–2.5
2.5–1.25
1.25–0.75
0.5
present
epochs (kya)
rs3803716 (TNRC6A) 911 ancient genotypes
convolution 20 ancient inds
convolution 100 ancient inds
pseudo haploid genotypes
rs2275477 (OSCP1) 1494 ancient genotypes
frequency
epochs (kya)
rs2275477 (OSCP1) 1494 ancient genotypes
convolution 20 ancient inds
convolution 100 ancient inds
pseudo haploid genotypes

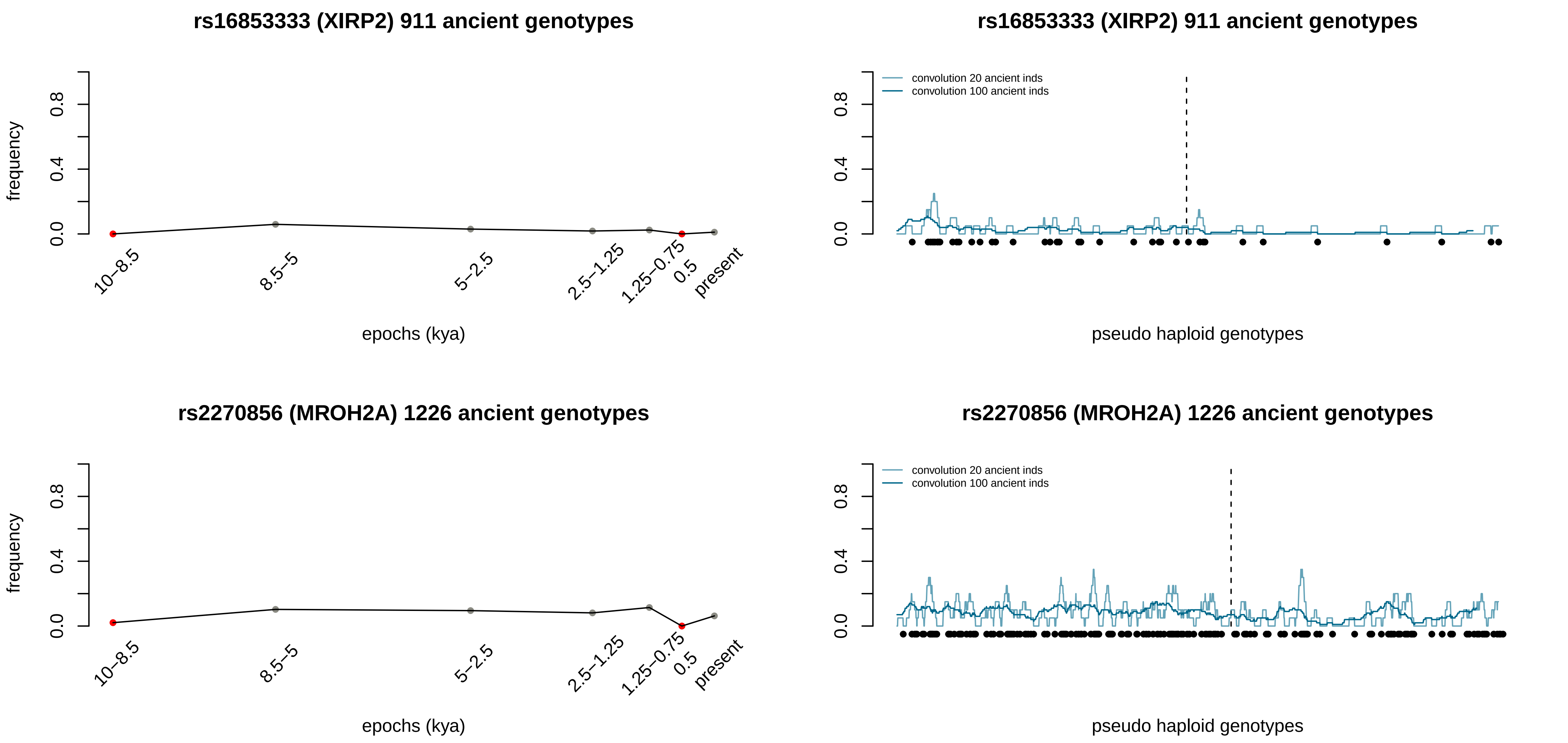

rs16853333 (XIRP2) 911 ancient genotypes
frequency
0.0
0.4
0.8
10–8.5
8.5–5
5–2.5
2.5–1.25
1.25–0.75
0.5
present
epochs (kya)
rs16853333 (XIRP2) 911 ancient genotypes
convolution 20 ancient inds
convolution 100 ancient inds
pseudo haploid genotypes
rs2270856 (MROH2A) 1226 ancient genotypes
frequency
epochs (kya)
rs2270856 (MROH2A) 1226 ancient genotypes
convolution 20 ancient inds
convolution 100 ancient inds
pseudo haploid genotypes

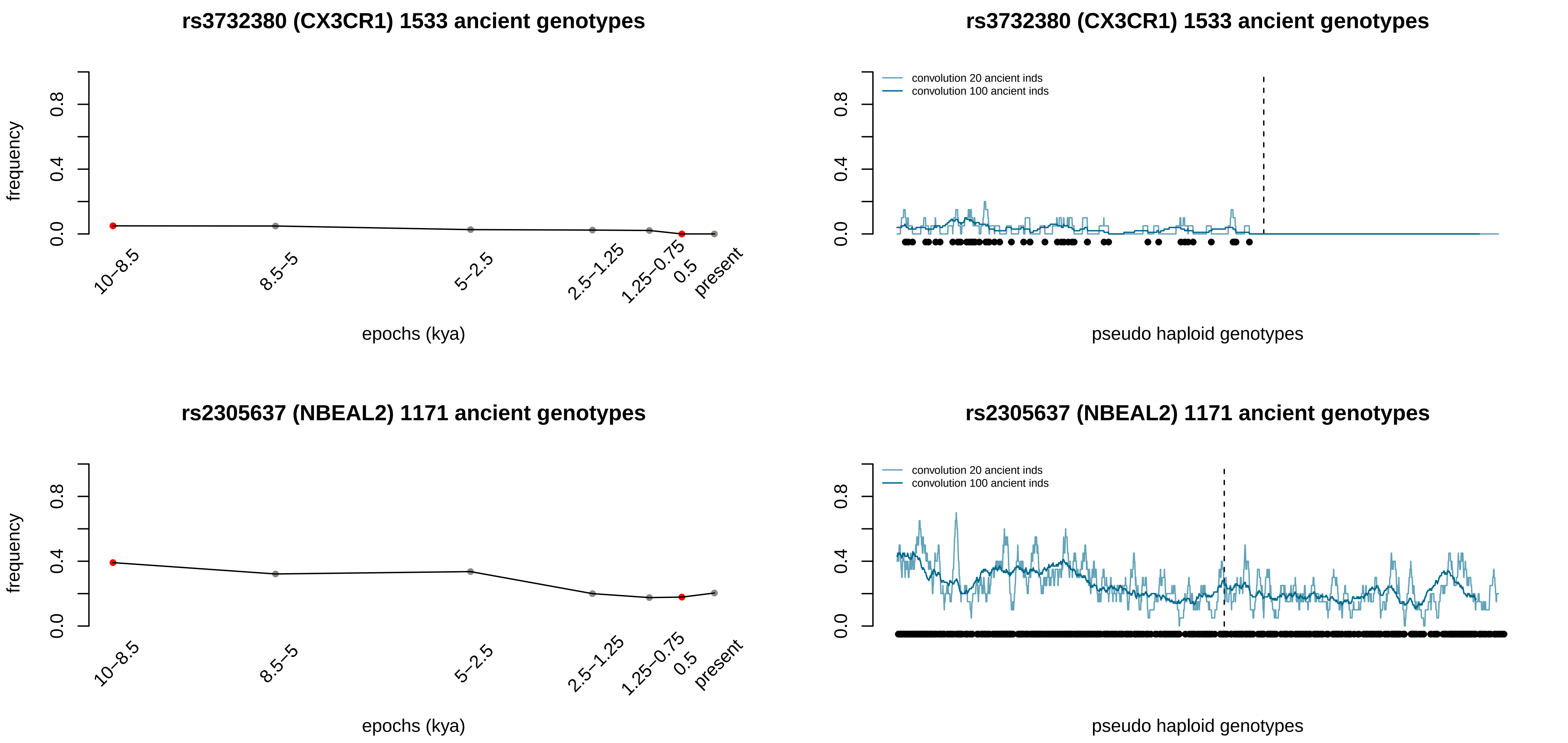

rs3732380 (CX3CR1) 1533 ancient genotypes
frequency
0.0
0.4
0.8
10–8.5
8.5–5
5–2.5
2.5–1.25
1.25–0.75
0.5
present
epochs (kya)
rs3732380 (CX3CR1) 1533 ancient genotypes
convolution 20 ancient inds
convolution 100 ancient inds
pseudo haploid genotypes
rs2305637 (NBEAL2) 1171 ancient genotypes
frequency
epochs (kya)
rs2305637 (NBEAL2) 1171 ancient genotypes
convolution 20 ancient inds
convolution 100 ancient inds
pseudo haploid genotypes

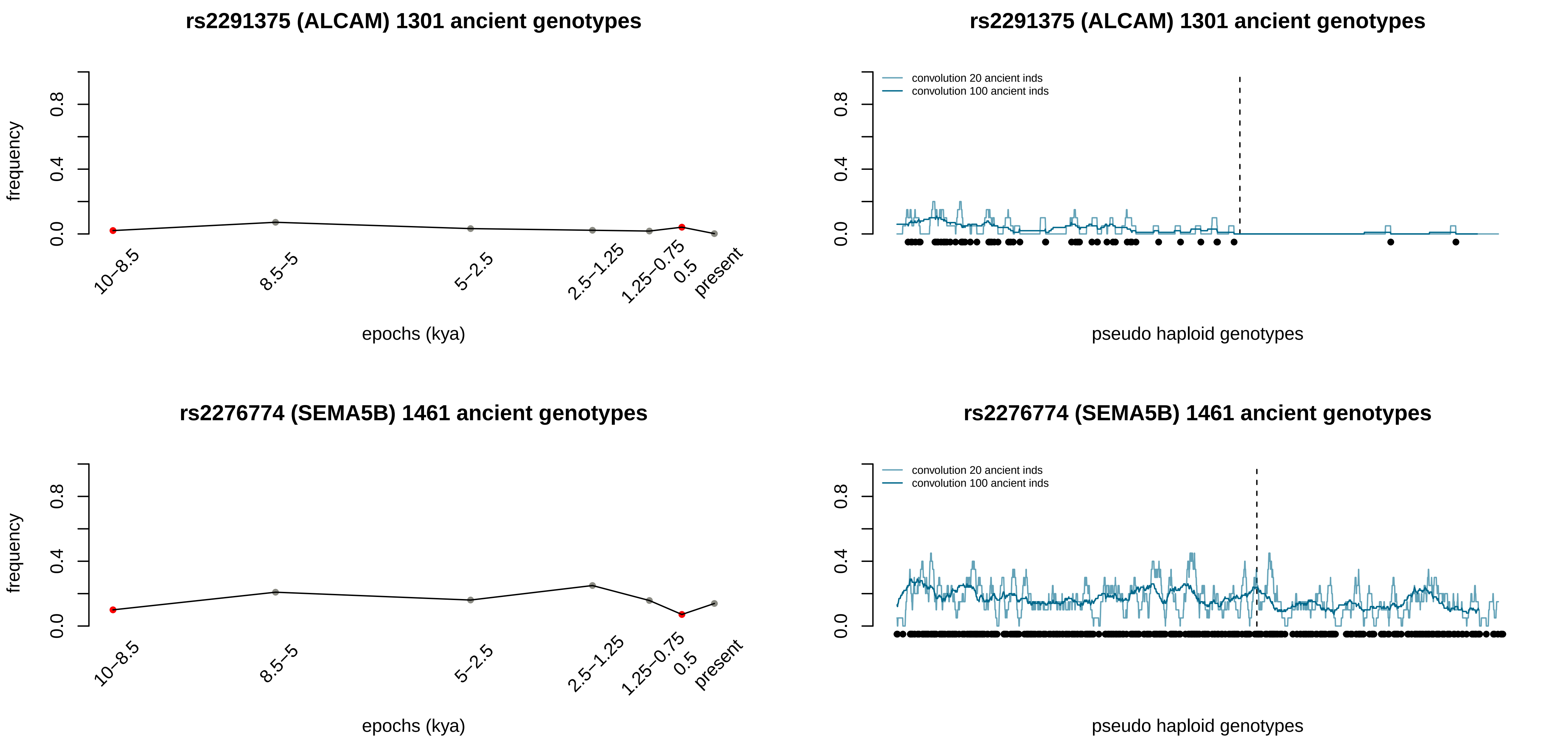
rs2291375 (ALCAM) 1301 ancient genotypes
frequency
0.0
0.4
0.8
10–8.5
8.5–5
5–2.5
2.5–1.25
1.25–0.75
0.5
present
epochs (kya)
rs2291375 (ALCAM) 1301 ancient genotypes
convolution 20 ancient inds
convolution 100 ancient inds
pseudo haploid genotypes
rs2276774 (SEMA5B) 1461 ancient genotypes
frequency
epochs (kya)
rs2276774 (SEMA5B) 1461 ancient genotypes
convolution 20 ancient inds
convolution 100 ancient inds
pseudo haploid genotypes

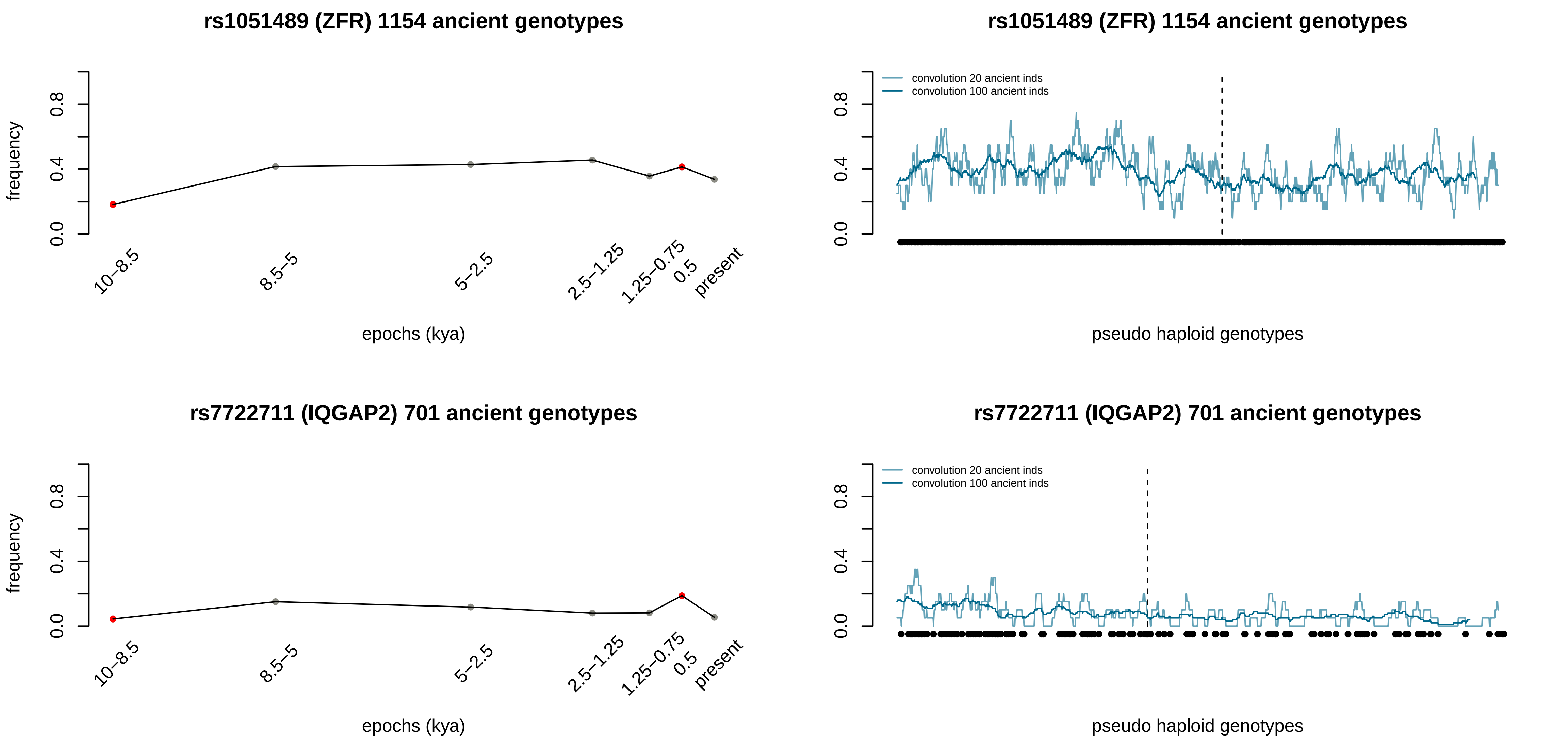

rs1051489 (ZFR) 1154 ancient genotypes
frequency
0.0
0.4
0.8
10–8.5
8.5–5
5–2.5
2.5–1.25
1.25–0.75
0.5
present
epochs (kya)
rs1051489 (ZFR) 1154 ancient genotypes
convolution 20 ancient inds
convolution 100 ancient inds
pseudo haploid genotypes
rs7722711 (IQGAP2) 701 ancient genotypes
frequency
epochs (kya)
rs7722711 (IQGAP2) 701 ancient genotypes
convolution 20 ancient inds
convolution 100 ancient inds
pseudo haploid genotypes

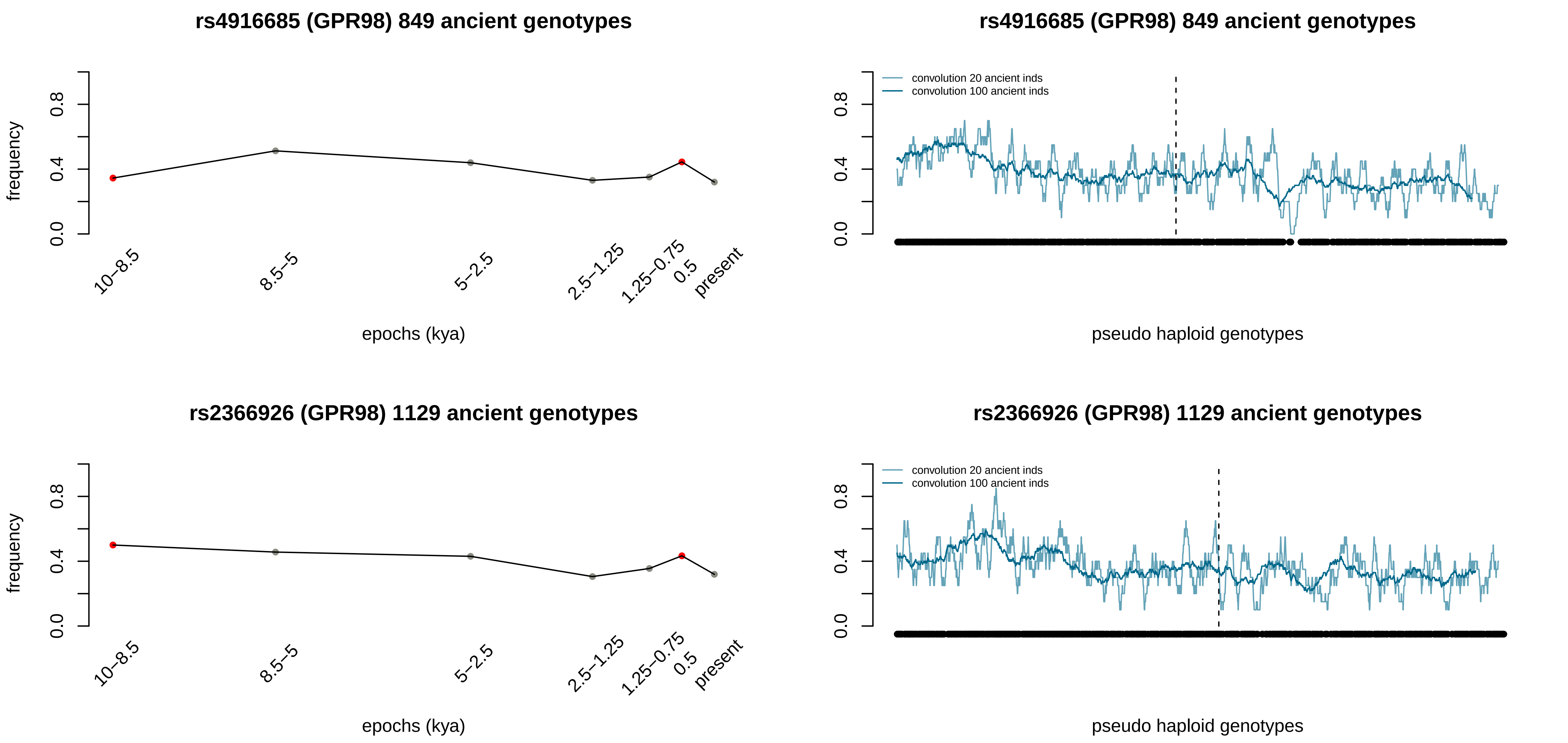
rs4916685 (GPR98) 849 ancient genotypes
frequency
0.0
0.4
0.8
10–8.5
8.5–5
5–2.5
2.5–1.25
1.25–0.75
0.5
present
epochs (kya)
rs4916685 (GPR98) 849 ancient genotypes
convolution 20 ancient inds
convolution 100 ancient inds
pseudo haploid genotypes
rs2366926 (GPR98) 1129 ancient genotypes
frequency
epochs (kya)
rs2366926 (GPR98) 1129 ancient genotypes
convolution 20 ancient inds
convolution 100 ancient inds
pseudo haploid genotypes

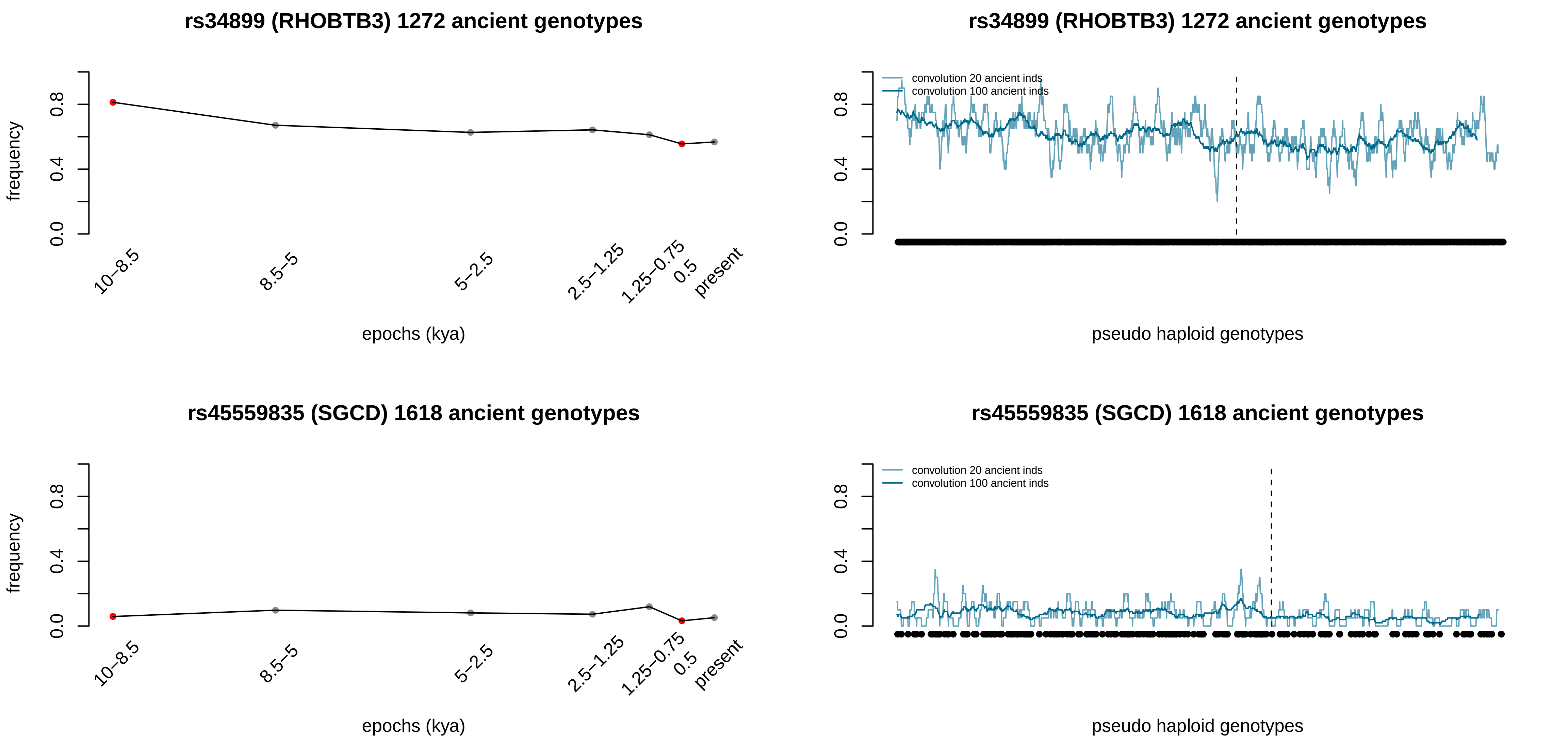

rs34899 (RHOBTB3) 1272 ancient genotypes
frequency
0.0
0.4
0.8
10–8.5
8.5–5
5–2.5
2.5–1.25
1.25–0.75
0.5
present
epochs (kya)
rs34899 (RHOBTB3) 1272 ancient genotypes
convolution 20 ancient inds
convolution 100 ancient inds
pseudo haploid genotypes
rs45559835 (SGCD) 1618 ancient genotypes
frequency
epochs (kya)
rs45559835 (SGCD) 1618 ancient genotypes
convolution 20 ancient inds
convolution 100 ancient inds
pseudo haploid genotypes

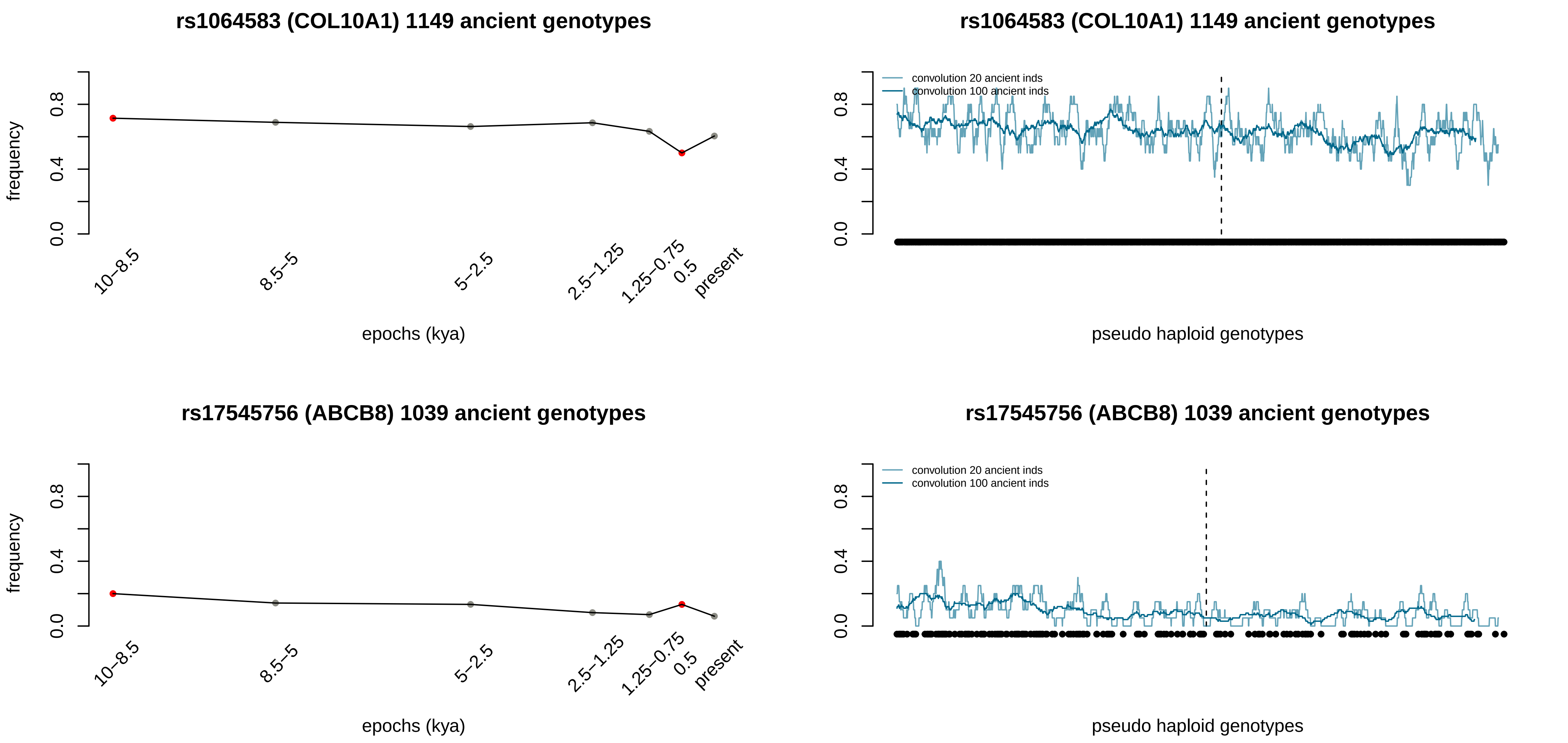
rs1064583 (COL10A1) 1149 ancient genotypes
frequency
0.0
0.4
0.8
10–8.5
8.5–5
5–2.5
2.5–1.25
1.25–0.75
0.5
present
epochs (kya)
rs1064583 (COL10A1) 1149 ancient genotypes
convolution 20 ancient inds
convolution 100 ancient inds
pseudo haploid genotypes
rs17545756 (ABCB8) 1039 ancient genotypes
frequency
epochs (kya)
rs17545756 (ABCB8) 1039 ancient genotypes
convolution 20 ancient inds
convolution 100 ancient inds
pseudo haploid genotypes

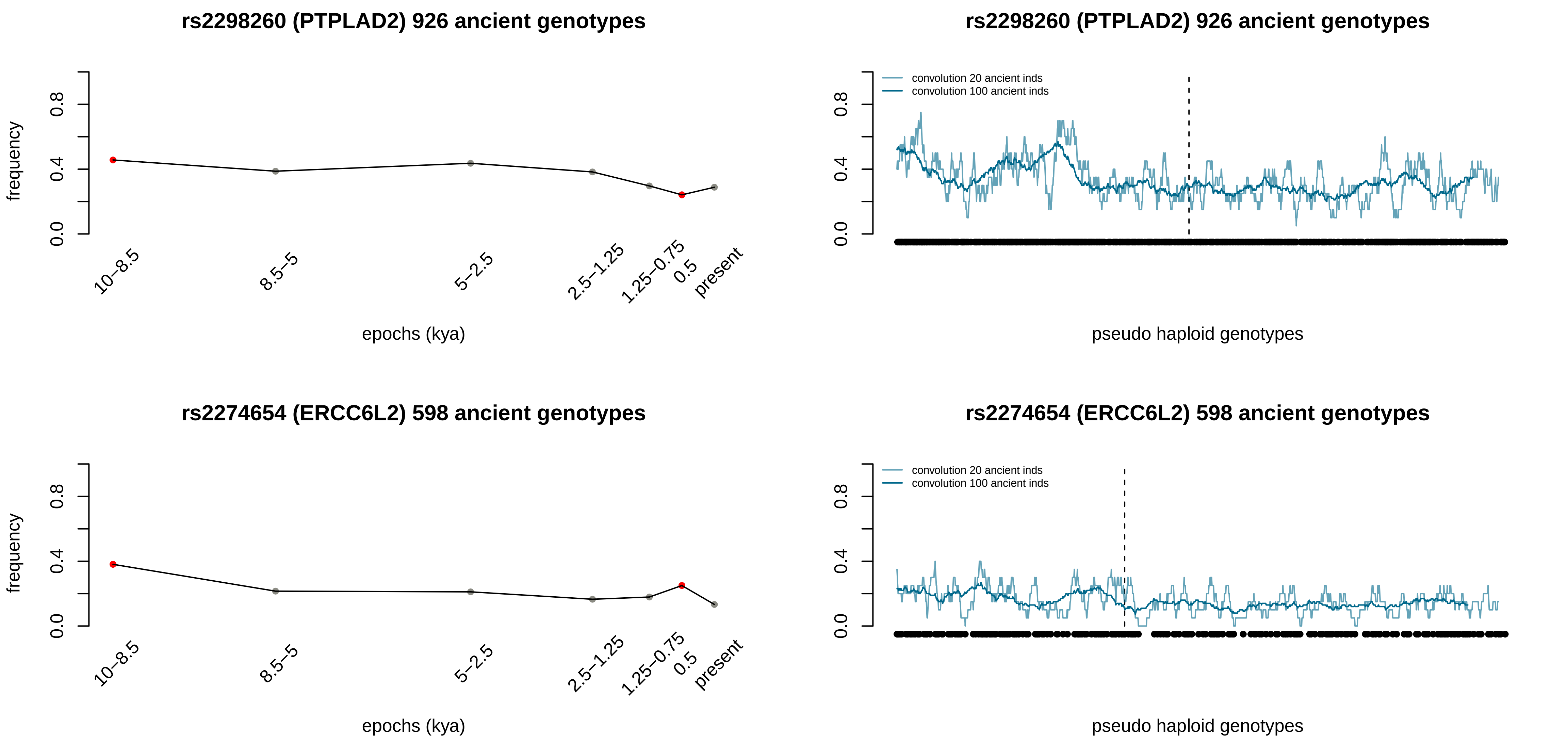
rs2298260 (PTPLAD2) 926 ancient genotypes
frequency
0.0
0.4
0.8
10–8.5
8.5–5
5–2.5
2.5–1.25
1.25–0.75
0.5
present
epochs (kya)
rs2298260 (PTPLAD2) 926 ancient genotypes
convolution 20 ancient inds
convolution 100 ancient inds
pseudo haploid genotypes
rs2274654 (ERCC6L2) 598 ancient genotypes
frequency
epochs (kya)
rs2274654 (ERCC6L2) 598 ancient genotypes
convolution 20 ancient inds
convolution 100 ancient inds
pseudo haploid genotypes

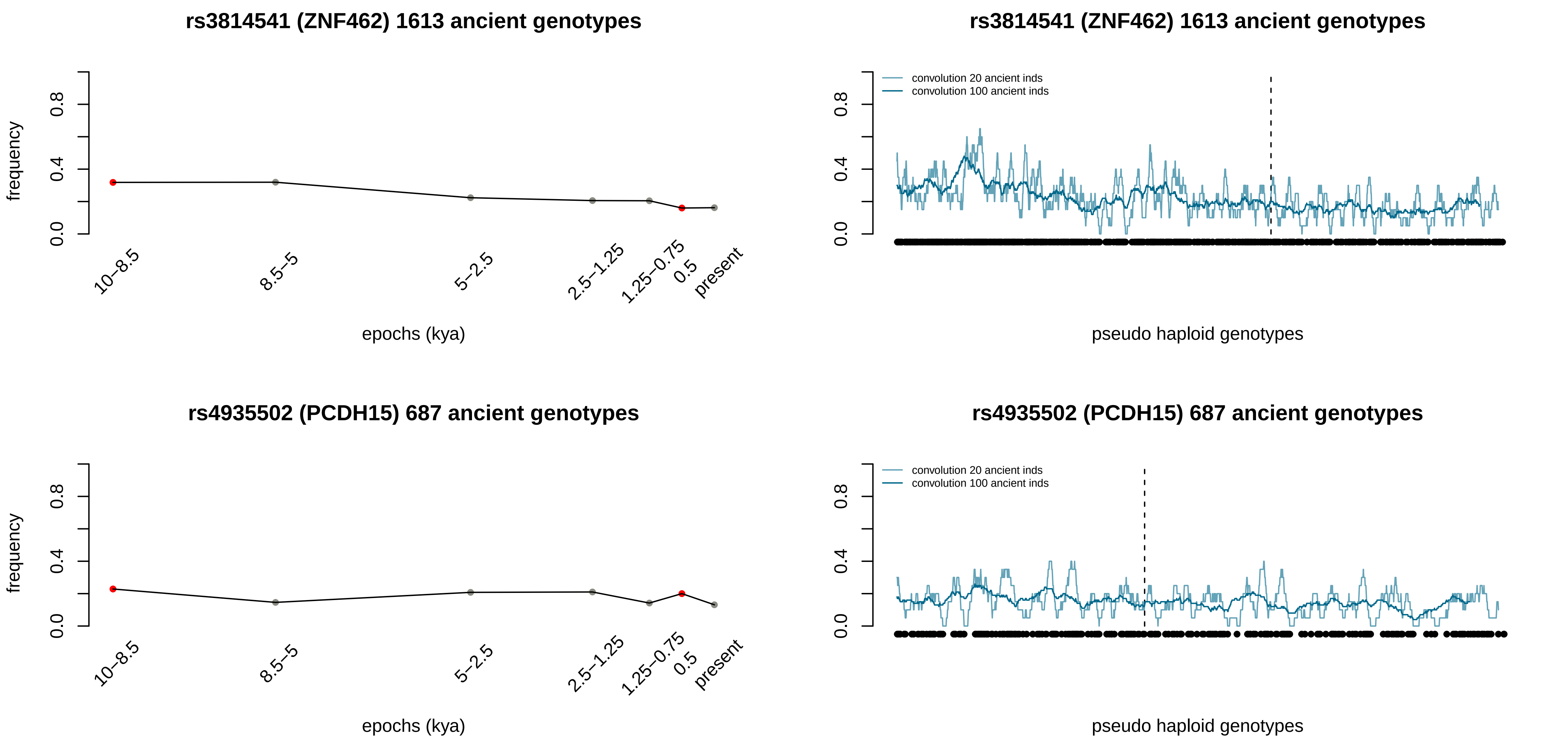
rs3814541 (ZNF462) 1613 ancient genotypes
frequency
0.0
0.4
0.8
10–8.5
8.5–5
5–2.5
2.5–1.25
1.25–0.75
0.5
present
epochs (kya)
rs3814541 (ZNF462) 1613 ancient genotypes
convolution 20 ancient inds
convolution 100 ancient inds
0.0
0.4
0.8
pseudo haploid genotypes
rs4935502 (PCDH15) 687 ancient genotypes
frequency
0.0
0.4
0.8
10–8.5
8.5–5
5–2.5
2.5–1.25
1.25–0.75
0.5
present
epochs (kya)
rs4935502 (PCDH15) 687 ancient genotypes
convolution 20 ancient inds
convolution 100 ancient inds
0.0
0.4
0.8
pseudo haploid genotypes

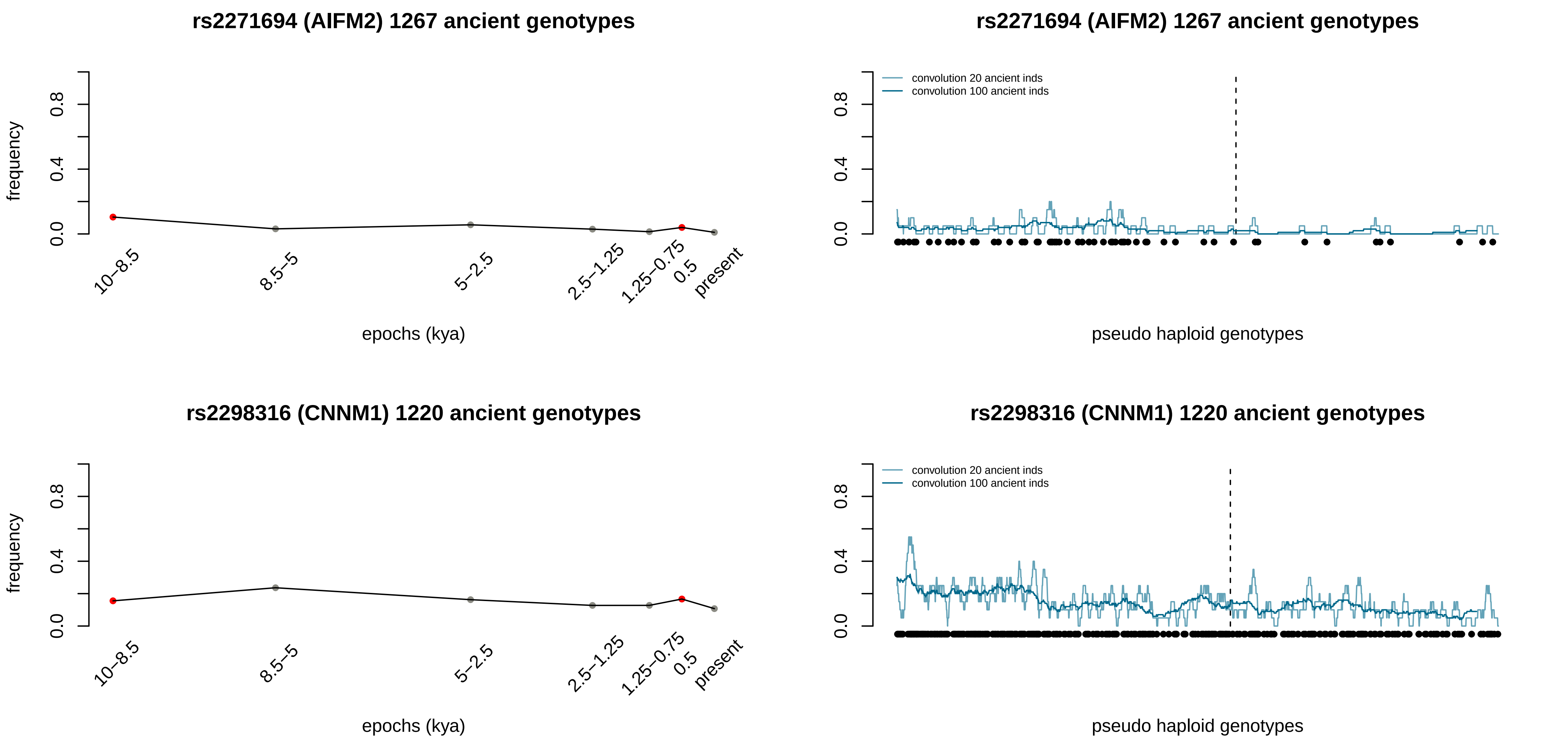
rs2271694 (AIFM2) 1267 ancient genotypes
frequency
0.0
0.4
0.8
10–8.5
8.5–5
5–2.5
2.5–1.25
1.25–0.75
0.5
present
epochs (kya)
rs2271694 (AIFM2) 1267 ancient genotypes
convolution 20 ancient inds
convolution 100 ancient inds
pseudo haploid genotypes
rs2298316 (CNNM1) 1220 ancient genotypes
frequency
epochs (kya)
rs2298316 (CNNM1) 1220 ancient genotypes
convolution 20 ancient inds
convolution 100 ancient inds
pseudo haploid genotypes

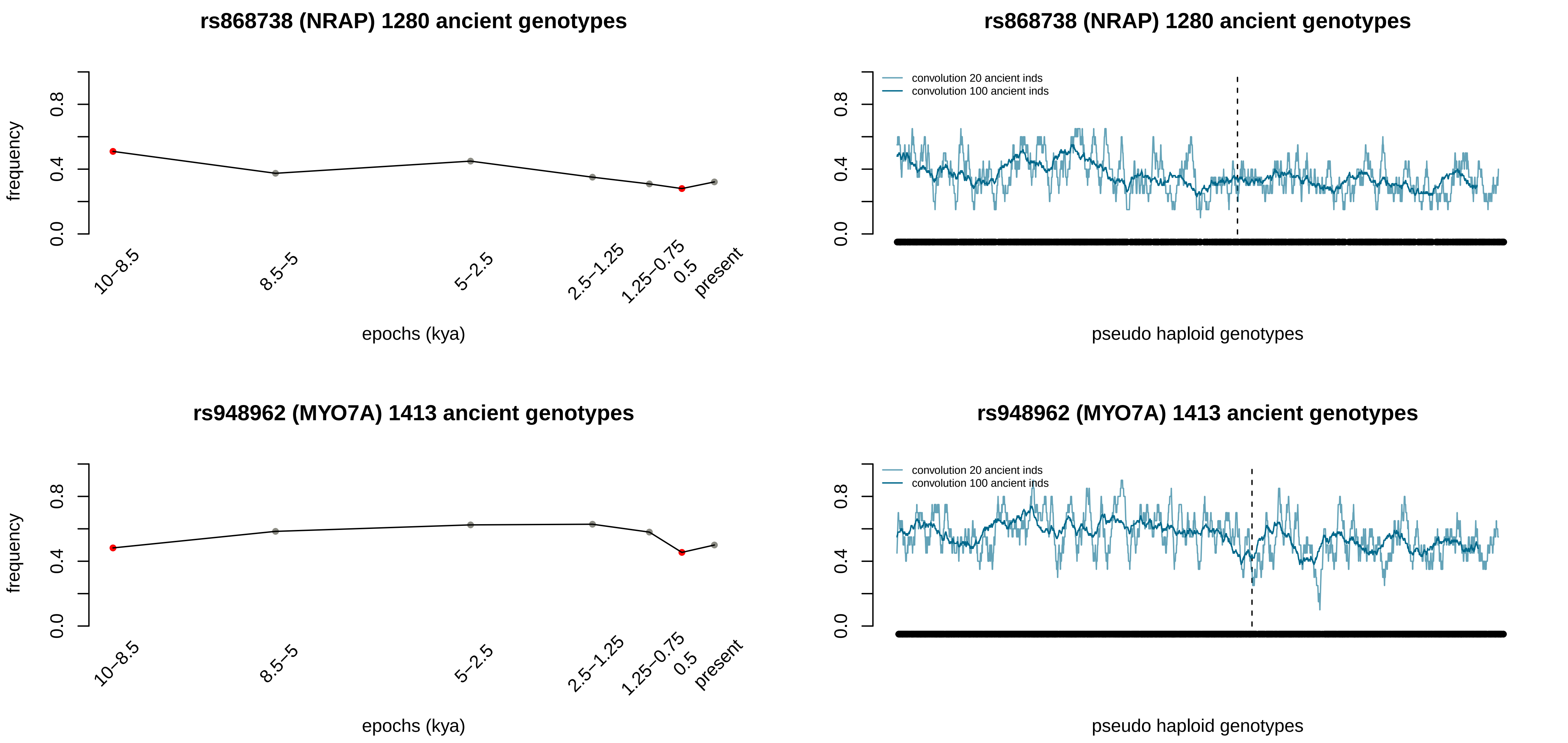

rs868738 (NRAP) 1280 ancient genotypes
frequency
0.0
0.4
0.8
10–8.5
8.5–5
5–2.5
2.5–1.25
1.25–0.75
0.5
present
epochs (kya)
rs868738 (NRAP) 1280 ancient genotypes
convolution 20 ancient inds
convolution 100 ancient inds
pseudo haploid genotypes
rs948962 (MYO7A) 1413 ancient genotypes
frequency
epochs (kya)
rs948962 (MYO7A) 1413 ancient genotypes
convolution 20 ancient inds
convolution 100 ancient inds
pseudo haploid genotypes

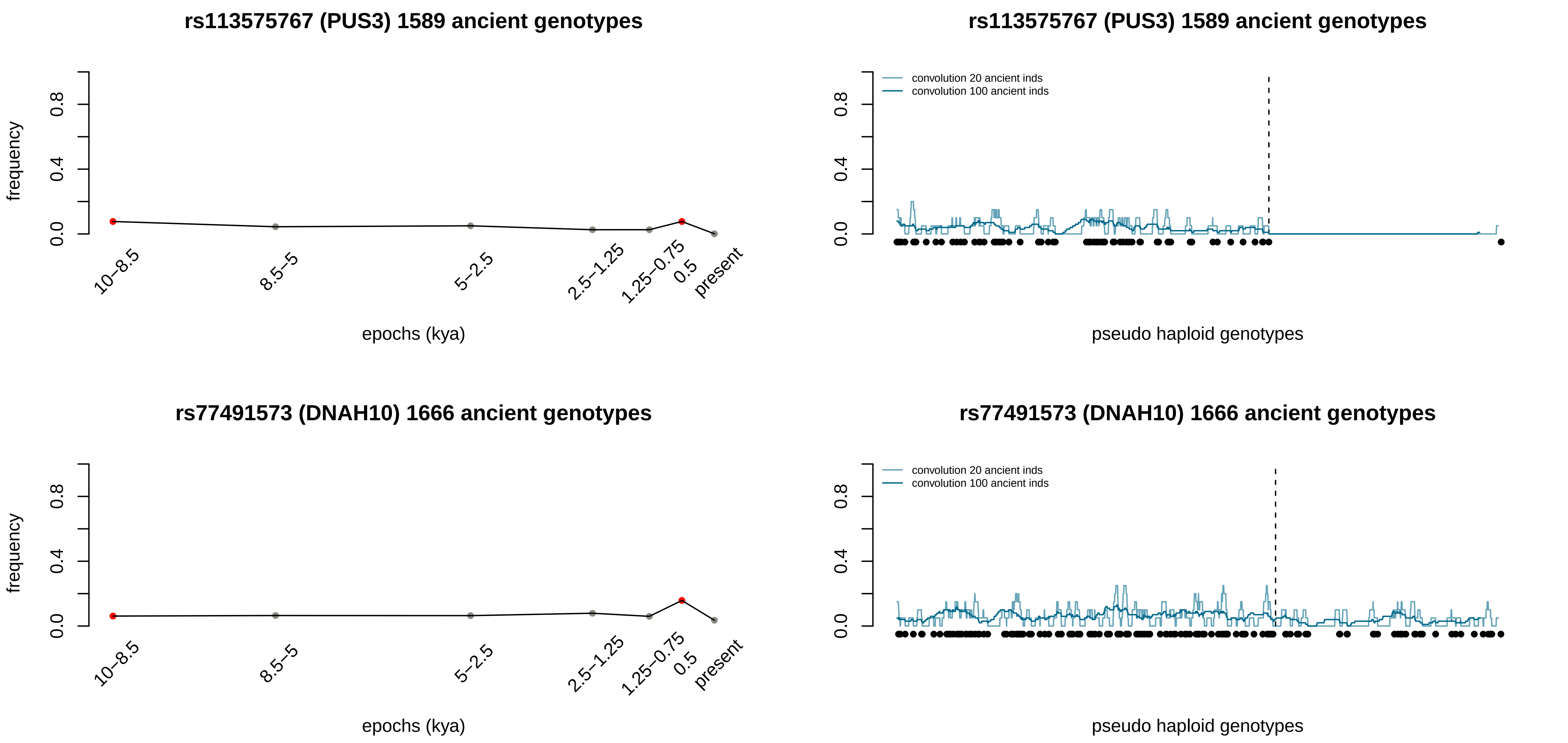
rs113575767 (PUS3) 1589 ancient genotypes
frequency
0.0
0.4
0.8
10–8.5
8.5–5
5–2.5
2.5–1.25
1.25–0.75
0.5
present
epochs (kya)
rs113575767 (PUS3) 1589 ancient genotypes
convolution 20 ancient inds
convolution 100 ancient inds
pseudo haploid genotypes
rs77491573 (DNAH10) 1666 ancient genotypes
frequency
epochs (kya)
rs77491573 (DNAH10) 1666 ancient genotypes
convolution 20 ancient inds
convolution 100 ancient inds
pseudo haploid genotypes

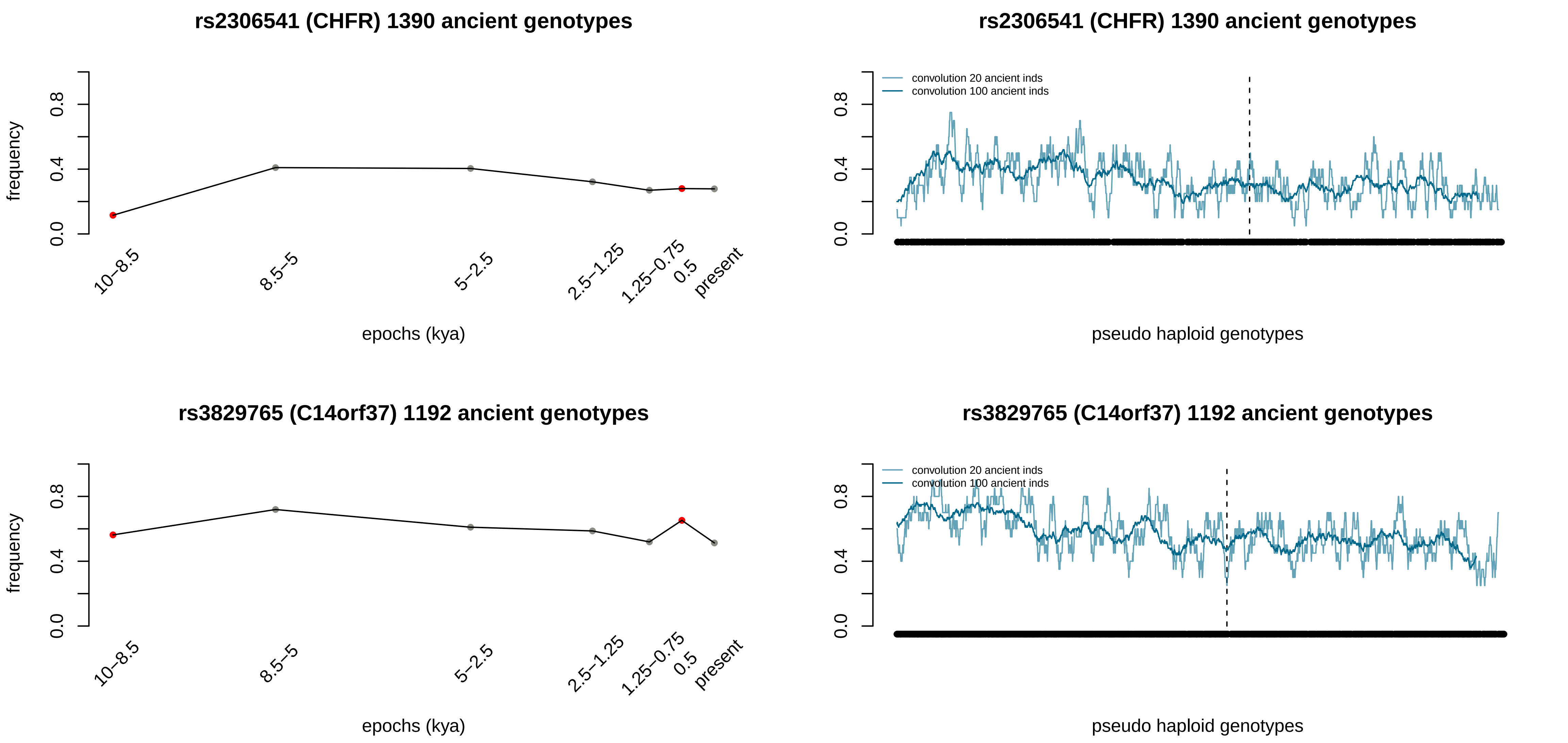
rs2306541 (CHFR) 1390 ancient genotypes
frequency
0.0
0.4
0.8
10–8.5
8.5–5
5–2.5
2.5–1.25
1.25–0.75
0.5
present
epochs (kya)
rs2306541 (CHFR) 1390 ancient genotypes
convolution 20 ancient inds
convolution 100 ancient inds
pseudo haploid genotypes
rs3829765 (C14orf37) 1192 ancient genotypes
frequency
epochs (kya)
rs3829765 (C14orf37) 1192 ancient genotypes
convolution 20 ancient inds
convolution 100 ancient inds
pseudo haploid genotypes

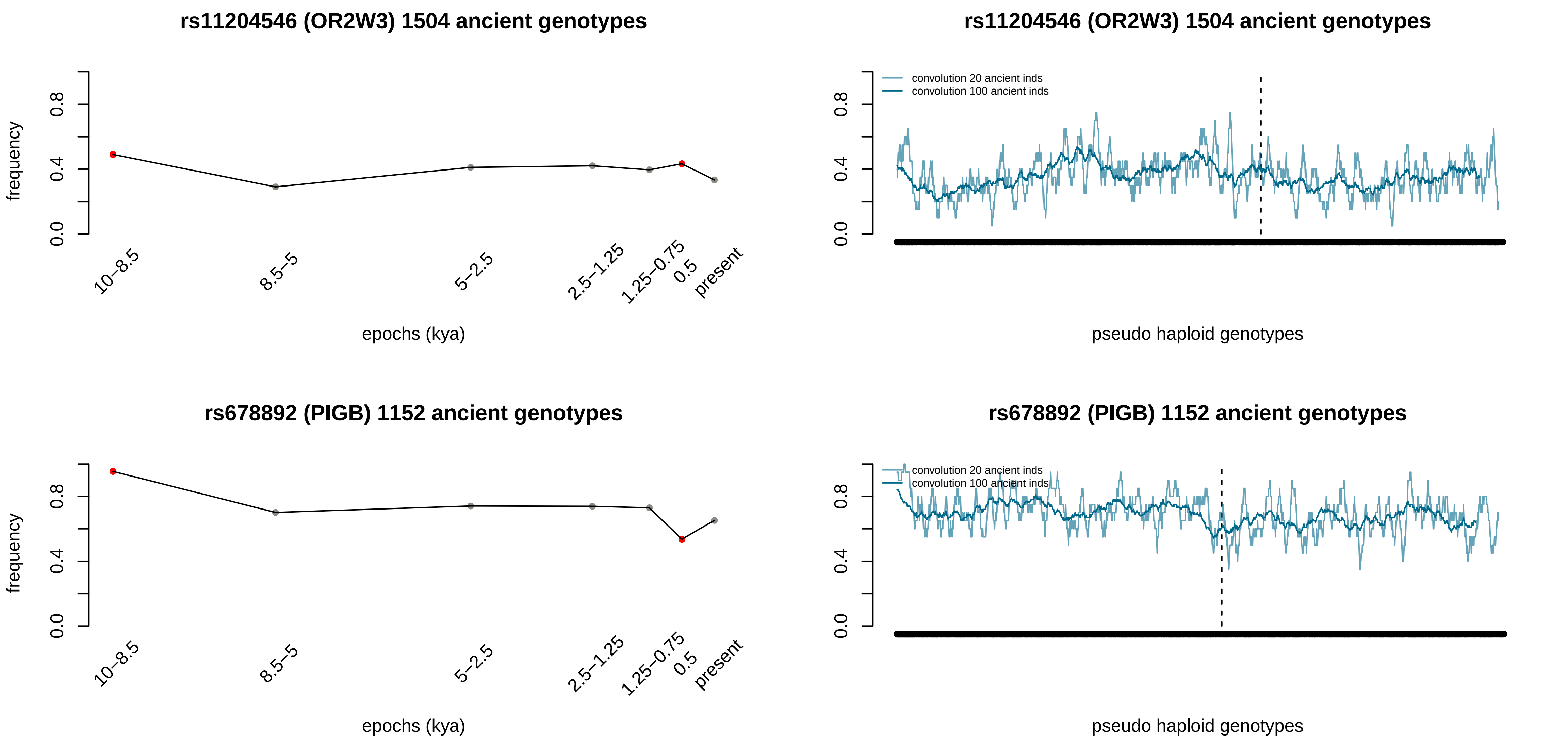
rs11204546 (OR2W3) 1504 ancient genotypes
frequency
0.0
0.4
0.8
10–8.5
8.5–5
5–2.5
2.5–1.25
1.25–0.75
0.5
present
epochs (kya)
rs11204546 (OR2W3) 1504 ancient genotypes
convolution 20 ancient inds
convolution 100 ancient inds
0.0
0.4
0.8
pseudo haploid genotypes
rs678892 (PIGB) 1152 ancient genotypes
frequency
0.0
0.4
0.8
10–8.5
8.5–5
5–2.5
2.5–1.25
1.25–0.75
0.5
present
epochs (kya)
rs678892 (PIGB) 1152 ancient genotypes
convolution 20 ancient inds
convolution 100 ancient inds
0.0
0.4
0.8
pseudo haploid genotypes

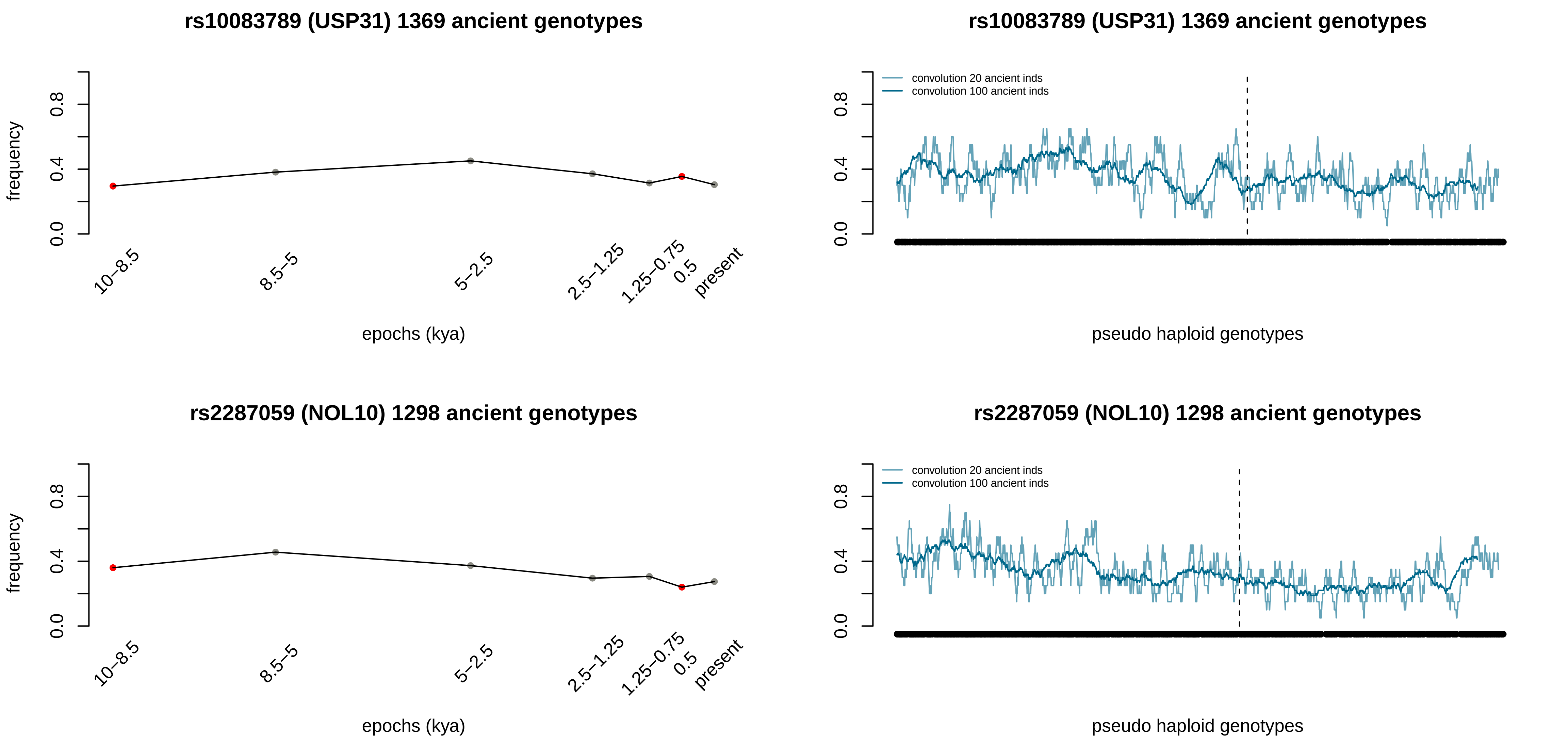

rs10083789 (USP31) 1369 ancient genotypes
frequency
0.0
0.4
0.8
10–8.5
8.5–5
5–2.5
2.5–1.25
1.25–0.75
0.5
present
epochs (kya)
rs10083789 (USP31) 1369 ancient genotypes
convolution 20 ancient inds
convolution 100 ancient inds
pseudo haploid genotypes
rs2287059 (NOL10) 1298 ancient genotypes
frequency
epochs (kya)
rs2287059 (NOL10) 1298 ancient genotypes
convolution 20 ancient inds
convolution 100 ancient inds
pseudo haploid genotypes

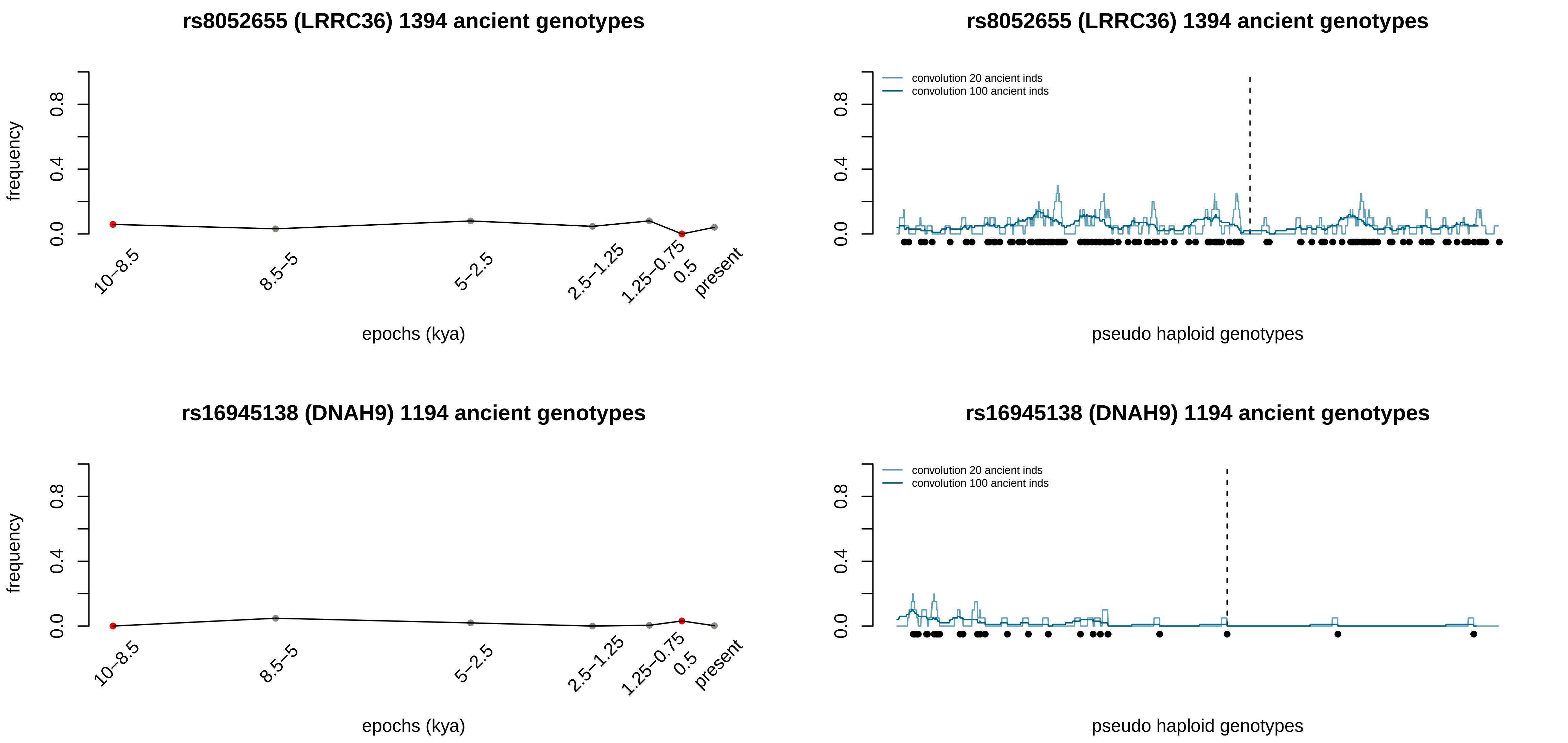
rs8052655 (LRRC36) 1394 ancient genotypes
frequency
0.0
0.4
0.8
10–8.5
8.5–5
5–2.5
2.5–1.25
1.25–0.75
0.5
present
epochs (kya)
rs8052655 (LRRC36) 1394 ancient genotypes
convolution 20 ancient inds
convolution 100 ancient inds
0.0
0.4
0.8
pseudo haploid genotypes
rs16945138 (DNAH9) 1194 ancient genotypes
frequency
0.0
0.4
0.8
10–8.5
8.5–5
5–2.5
2.5–1.25
1.25–0.75
0.5
present
epochs (kya)
rs16945138 (DNAH9) 1194 ancient genotypes
convolution 20 ancient inds
convolution 100 ancient inds
0.0
0.4
0.8
pseudo haploid genotypes

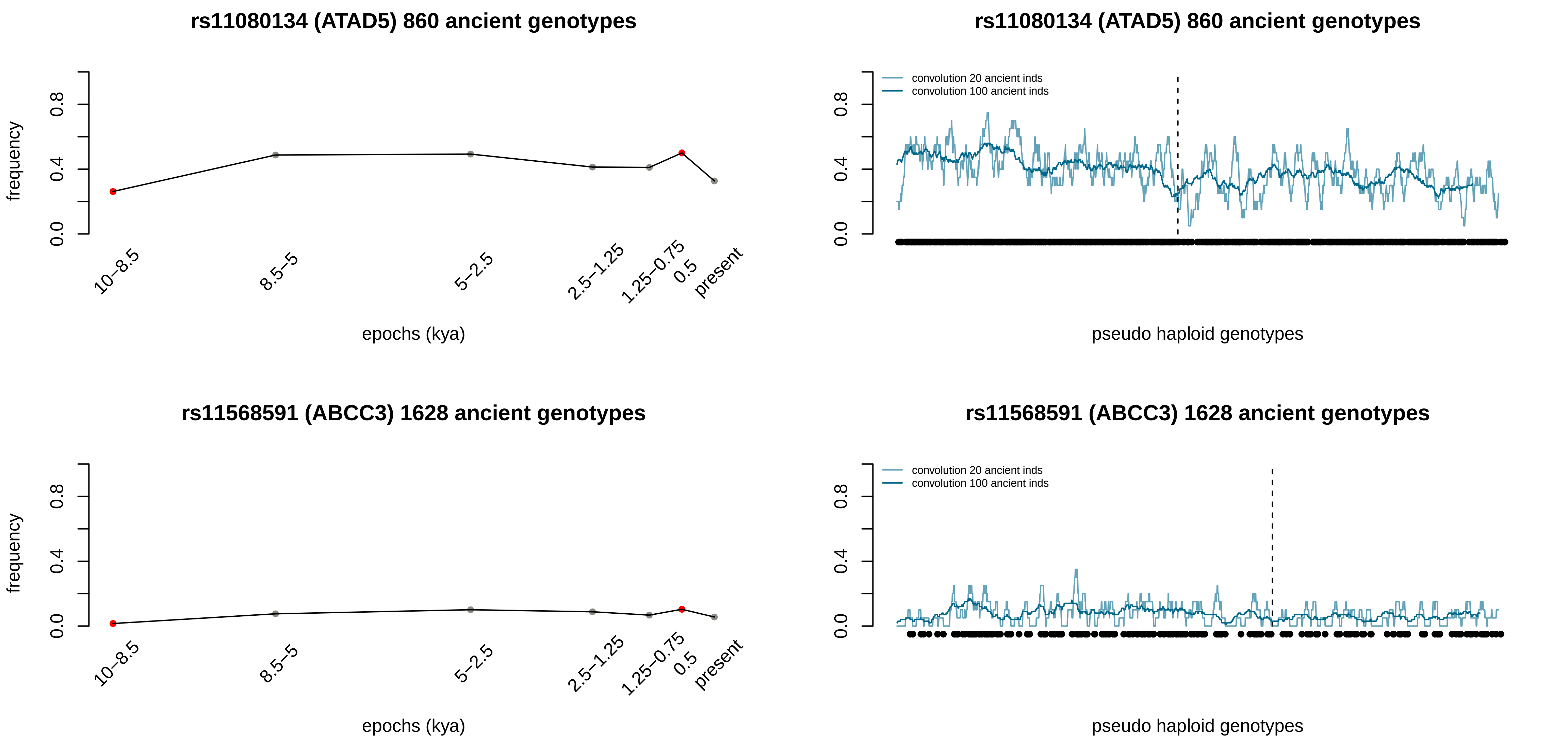
rs11080134 (ATAD5) 860 ancient genotypes
frequency
0.0
0.4
0.8
10–8.5
8.5–5
5–2.5
2.5–1.25
1.25–0.75
0.5
present
epochs (kya)
rs11080134 (ATAD5) 860 ancient genotypes
convolution 20 ancient inds
convolution 100 ancient inds
pseudo haploid genotypes
rs11568591 (ABCC3) 1628 ancient genotypes
frequency
epochs (kya)
rs11568591 (ABCC3) 1628 ancient genotypes
convolution 20 ancient inds
convolution 100 ancient inds
pseudo haploid genotypes

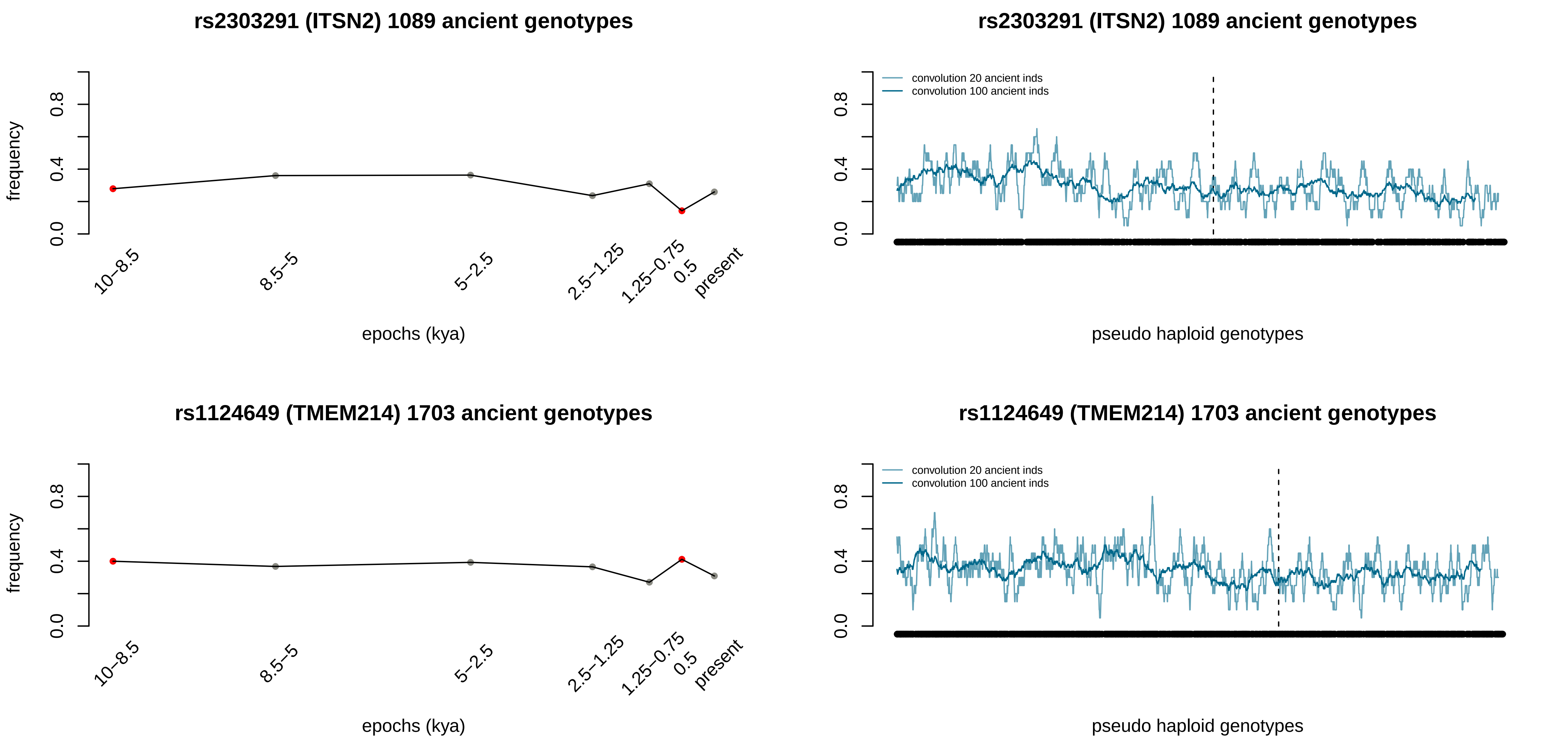
rs2303291 (ITSN2) 1089 ancient genotypes
frequency
0.0
0.4
0.8
10–8.5
8.5–5
5–2.5
2.5–1.25
1.25–0.75
0.5
present
epochs (kya)
rs2303291 (ITSN2) 1089 ancient genotypes
convolution 20 ancient inds
convolution 100 ancient inds
pseudo haploid genotypes
rs1124649 (TMEM214) 1703 ancient genotypes
frequency
epochs (kya)
rs1124649 (TMEM214) 1703 ancient genotypes
convolution 20 ancient inds
convolution 100 ancient inds
pseudo haploid genotypes

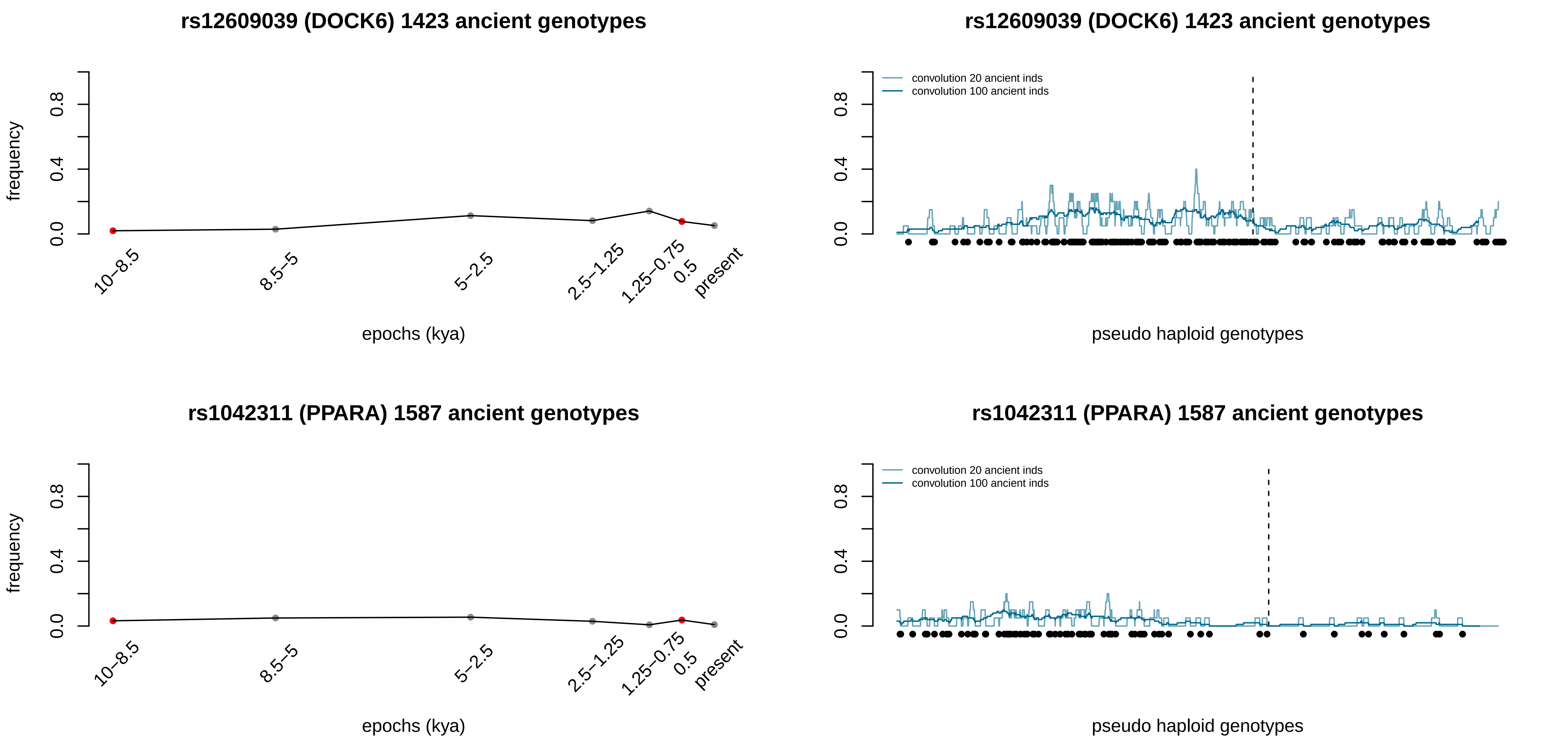
rs12609039 (DOCK6) 1423 ancient genotypes
frequency
0.0
0.4
0.8
10–8.5
8.5–5
5–2.5
2.5–1.25
1.25–0.75
0.5
present
epochs (kya)
rs12609039 (DOCK6) 1423 ancient genotypes
convolution 20 ancient inds
convolution 100 ancient inds
pseudo haploid genotypes
rs1042311 (PPARA) 1587 ancient genotypes
frequency
epochs (kya)
rs1042311 (PPARA) 1587 ancient genotypes
convolution 20 ancient inds
convolution 100 ancient inds
pseudo haploid genotypes

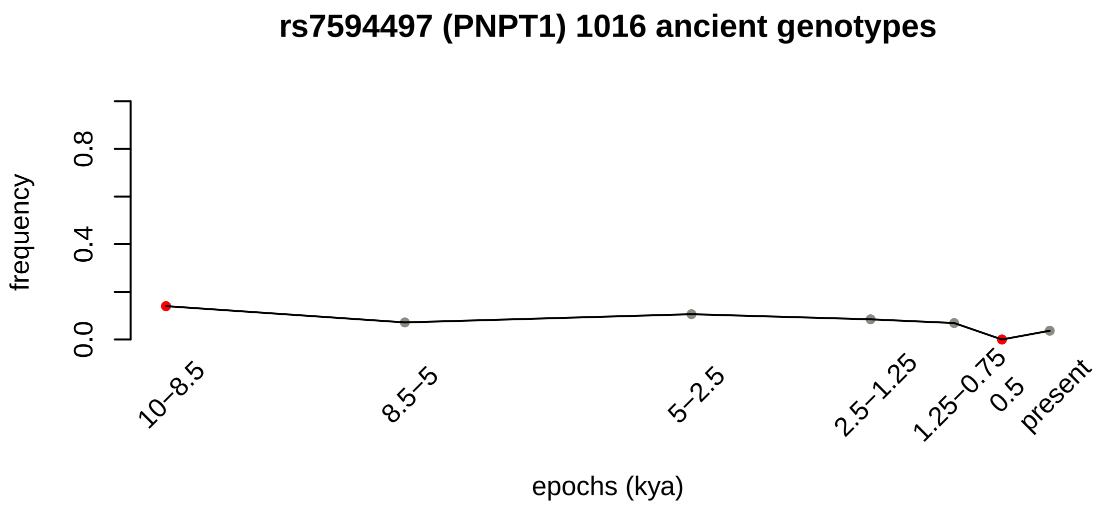
rs7594497 (PNPT1) 1016 ancient genotypes
frequency
0.0
0.4
0.8
10–8.5
8.5–5
5–2.5
2.5–1.25
1.25–0.75
0.5
present
epochs (kya)

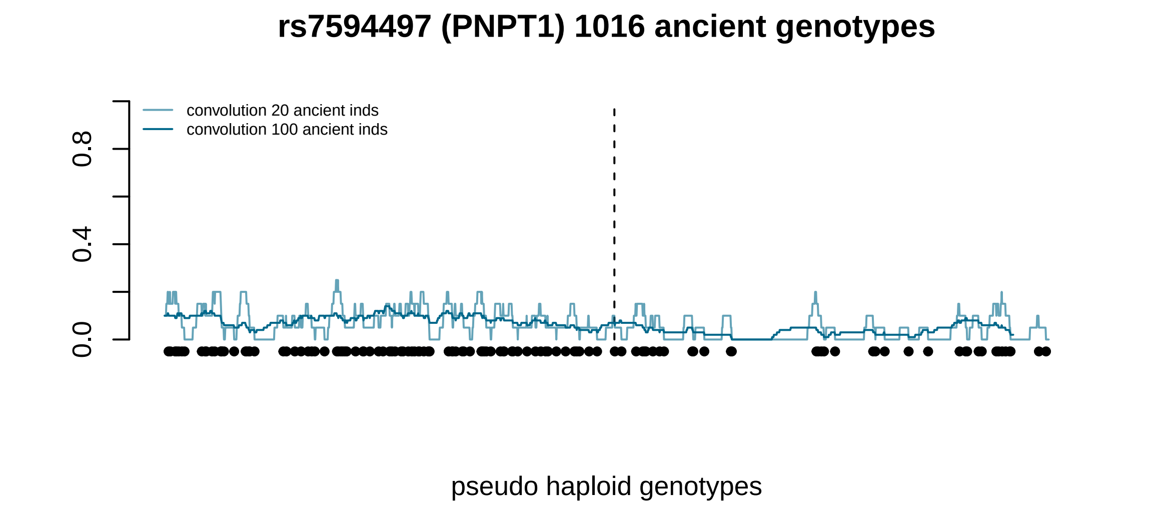
rs7594497 (PNPT1) 1016 ancient genotypes
convolution 20 ancient inds
convolution 100 ancient inds
0.0
0.4
0.8
pseudo haploid genotypes
